# Evolutionary dynamics of *de novo* mutations and mutant lineages arising in a simple, constant environment

**DOI:** 10.1101/540625

**Authors:** Margie Kinnersley, Katja Schwartz, Jacob Boswell, Dong-Dong Yang, Gavin Sherlock, Frank Rosenzweig

**Affiliations:** Division of Biological Sciences, The University of Montana, Missoula, MT 59812; School of Biological Sciences, Georgia Institute of Technology, Atlanta, GA 30332; Department of Genetics, Stanford University School of Medicine, Stanford, CA 94305-5120

**Author notes:** These authors contributed equally to this work (Margie Kinnersley, Katja Schwartz).

**Keywords:** *E. coli*, adaptive evolution, chemostat, metagenomics, population sequencing, clone sequencing, clonal interference, parallelism

## Abstract

A large, asexual population founded by a single clone evolves into a population teeming with many, whether or not its environment is structured, and whether or not resource levels are constant or fluctuating. The maintenance of genetic complexity in such populations has been attributed to balancing selection, or to either clonal interference or clonal reinforcement, arising from antagonistic or synergistic interactions, respectively. To distinguish among these possibilities, to identify targets of selection and establish when and how often they are hit, as well as to gain insight into how *de novo* mutations interact, we carried out 300-500 generation glucose-limited chemostat experiments founded by an *E. coli* mutator. To discover all *de novo* mutations reaching ≥1% frequency, we performed whole-genome, whole-population sequencing at ∼1000X-coverage every 50 generations. To establish linkage relationships among these mutations and depict the dynamics of evolving lineages we sequenced the genomes of 96 clones from each population when allelic diversity was greatest. Operon-specific mutations that enhance glucose uptake arose to high frequency first, followed by global regulatory mutations. Late-arising mutations were related to energy conservation as well as to mitigating pleiotropic effects wrought by earlier regulatory changes. We discovered extensive polymorphism at relatively few loci, with identical mutations arising independently in different lineages, both between and within replicate populations. Out of more than 3,000 SNPs detected in nearly 1,800 genes or intergenic regions, only 17 reached a frequency ≥ 98%, indicating that the evolutionary dynamics of adaptive lineages was dominated by clonal interference. Finally, our data show that even when mutational input is increased by an ancestral defect in DNA repair, the spectrum of beneficial mutations that reach high frequency in a simple, constant resource-limited environment is narrow, resulting in extreme parallelism where many adaptive mutations arise but few ever go to fixation.

**Author Summary:** Microbial evolution experiments open a window on the tempo and dynamics of evolutionary change in asexual populations. High-throughput sequencing can be used to catalog *de novo* mutations, determine in which lineages they arise, and assess allelic interactions by tracking the fate of those lineages. This *adaptive genetics* approach makes it possible to discover whether clonal interactions are antagonistic or synergistic, and complements genetic screens of induced deleterious/loss-of-function mutants. We carried out glucose-limited chemostat experiments founded by an *E. coli* mutator and performed whole-genome, whole-population sequencing on 300-500 generation evolutions, cataloging 3,346 *de novo* mutations that reached ≥1% frequency. Mutations enhancing glucose uptake rose to high frequency first, followed by global regulatory changes that modulate growth rate and limiting resource assimilation, then by mutations that favor energy conservation or mitigate pleiotropic effects of earlier regulatory changes. We discovered that a few loci were highly polymorphic, with identical mutations arising independently in different lineages, both between and within replicate populations. Thus, when mutational input is increased by an ancestral defect in DNA repair, the spectrum of beneficial mutations that arises under constant resource-limitation is narrow, resulting in extreme parallelism where many adaptive mutations arise but few ever become fixed.

## Introduction

Evolution experiments using microbes have enlarged our understanding of the tempo and dynamics of evolutionary change, as well as how selection, drift and historical contingency influence evolutionary trajectories. Combined with high throughput sequencing, experimental microbial evolution (EME) can now be used to identify substantial numbers of *de novo* beneficial mutations in laboratory populations, to determine in which lineages they arise and the fate of those lineages, and to evaluate the sign and strength of possible epistatic interactions [1-3]. This approach, adaptive genetics, based on analyzing cohorts of spontaneous beneficial mutations to determine how their frequencies fluctuate over time, constitutes a mode of inquiry that complements traditional genetic screening of induced deleterious/loss-of-function mutants (e.g., [4] and [5] among others). Adaptive genetics also expands the possibilities for discovering constraints on protein structure and function and for discerning the architecture and malleability of networks that regulate nutrient-sensing and cell division.

Microbial populations were once thought to evolve by periodic selection as a succession of adaptive clones, each fitter than its antecedent, replacing one another over time [6-9]. This model was consistent with Muller and Haldane’s view of how beneficial mutations spread in large asexual populations [10-12] under conditions governed by competitive exclusion [13]. Today we know that large, initially clonal populations rapidly accumulate and retain genetic variation, much of which is beneficial [14-18]. In fact, the amount of adaptive genetic variation observed in EME populations can be enormous, owing large population sizes with a continuous input of beneficial mutations and the subsequent competition among new adaptive lineages, which gives rise to clonal interference [14,16,19,20].

Clonal interference can occur within a larger framework of stable subpopulation structure [21] when microbial lineages come under balancing selection [22-25] or specialize to exploit niches created either by the culture conditions [23,26,27], or by the organisms themselves [28-30]. In a simple constant environment like a chemostat the persistence of subpopulations likely depends on founder genotype, the emergence of specific key mutations, and availability of the limiting nutrient [31]. Ferenci and colleagues never observed stable subpopulation structure in glucose-limited evolutions originating from *E. coli* K12 strain BW2952 [32], whereas Adams and colleagues, using a different strain, often did [30,33]. Unlike BW2952, the K12-derived ancestor used by Adams, JA122 [30] harbors a supE44 *glnX* tRNA nonsense suppressor as well as nonsense mutations in housekeeping and stationary-phase transcription factors, RpoD and RpoS respectively, and mismatch repair enzyme MutY. The JA122 ancestor’s defect in DNA repair increases mutational load on its descendants [28,34], while the nonsense suppressor mitigates the effect of mutations that create premature stop codons. Such a suppressor would likely make the blunt instrument of *de novo* nonsense mutations a less effective agent of adaptive change, possibly resulting in a more nuanced spectrum of beneficial mutations than would otherwise occur among mutators.

To understand the impact that a mutator/suppressor founder has on the spectrum and fate of new beneficial mutations, and on the dynamics of population structure, we repeated Adams *et al*. classic evolution experiments using the same ancestral strain and culture conditions [30]. We monitored, at 50-generation intervals, the incidence of mutations that reached at least 1% frequency over the course of 300-500 generations, identifying mutations that were either transiently beneficial or hitch-hiking with mutations that were. To determine which mutations co-occurred within a given lineage we sequenced 96 clones from each population at the time-point where we observed greatest allelic diversity. We uncovered no evidence for stable sub-population structure, but instead saw pervasive clonal interference, with only 17 out of 3,346 mutations going to near fixation across replicate experiments. The temporal order in which certain mutations rose to high frequency was predictable, reflecting a high degree of parallelism both within and between replicates. In general, mutations that enhanced glucose assimilation arose early, followed by mutations in global regulators and mutations that either increase efficiency of limiting resource utilization or mitigate the deleterious effects of certain earlier mutations. Altogether, our results show that even in bacterial populations founded by an ancestor having a high mutation rate and the capacity to tolerate many *de novo* mutations, the spectrum of genomic changes that rise to appreciable frequency and the adaptive outcome of replicate evolutions are limited when those populations evolve in a simple constant environment.

## Results

### Experimental design

Evolution experiments were carried out in triplicate under continuous nutrient limitation using Davis Minimal Medium [30], with glucose (0.0125% w/v) as the sole source of carbon for energy and growth. Chemostats (300 mL working volume) were run under aerobic conditions for 300-500 generations at constant temperature (30°C) and at constant dilution rate (D=0.2 hr^−1^). Under these conditions, population density reaches ∼10^8^ cells mL^−1^ at steady state. The *E. coli* strain used to initiate these experiments, JA122, is distinguished from *E. coli* K12 by alleles likely to influence the spectrum of mutations arising during adaptive evolution (**Table S1**; [28]). Among these is a nonsense mutation in MutY (Leu299*) that results in a 10-fold greater mutation rate and GC→TA transversion bias [28], nonsense mutations in the genes that encode stationary phase sigma factor RpoS (Gln33*) [35] and ‘housekeeping’ sigma factor RpoD (Glu26*), as well as a suppressor mutation in the *glnX* tRNA known to suppress amber, ochre and opal mutations (**Table S1)** [36].

To identify the mutations that arose during the evolutions, we performed whole genome, whole population sequencing every 50 generations on each of the three chemostat populations. We generated approximately 50 million 2×100bp paired end reads per sample, yielding coverage of up to ∼1000x for each time point (inserts were selected to be short enough such that forward and reverse reads overlapped, which while reducing coverage, increases quality; see Methods). Based on this level of coverage, we were able to identify mutations that rose to an allele frequency of ∼1% of greater. Given an effective population size of >10^10^ and 300-500 generations of selection it is highly improbable that any allele could reach such a frequency by drift alone [23]. We can therefore assume that every mutation recovered was either under positive selection or hitch-hiking along with one that was.

### Population sequencing shows general patterns of mutation that are consistent across independent evolutions

Across all samples, 3,326 SNPs were detected in 2,083 unique genes or intergenic regions (**File S1)**. The overwhelming majority (97.5%) of these SNPs were GC→TA transversions, as expected given the ancestral strain’s defect in the mismatch repair protein MutY, which encodes adenine glycosylase [37]. Consistent with the protein coding density of *E. coli* (87.8%) [38], 85% (2,854) of SNPs occurred in coding regions. On average, 69.2% of these created a missense mutation, 23.4% resulted in a synonymous mutation and 7.4% caused a nonsense mutation (**Fig. 1**). Relative to proportions observed in mutation accumulation experiments carried out using wild-type *E. coli* [39], we observed more nonsynonymous and nonsense mutations. Small deletions were rarely detected (one single-nucleotide deletion in each of vessel 1 and vessel 2, and none detected in vessel 3), but we observed a single large ∼150kb duplication in vessel 2. The overall number of mutations in each population increased linearly over time and at approximately the same rate across replicates (**Fig. 1**), as would be expected with a mutator phenotype.

**Fig. 1.**
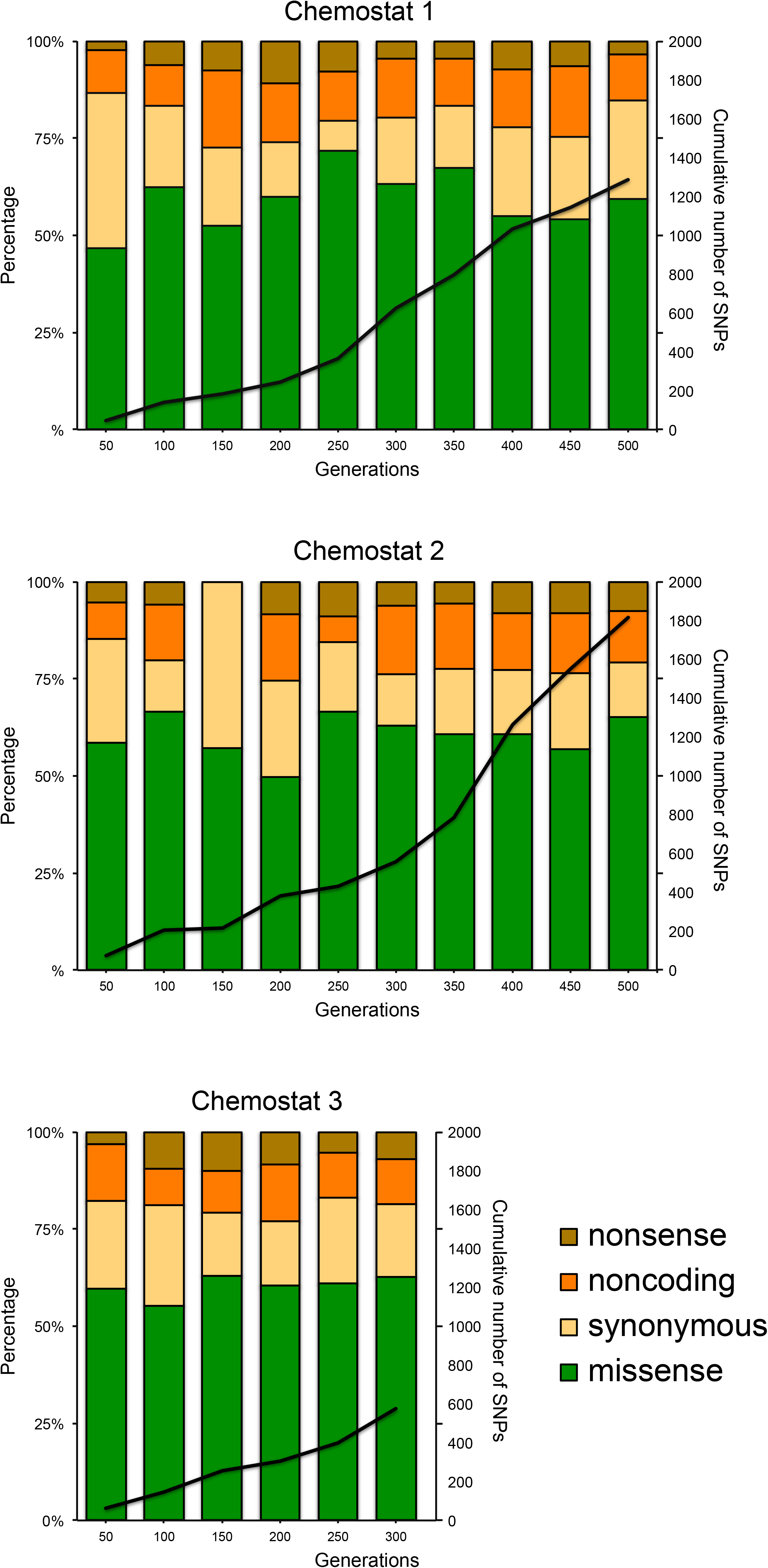
Input of de novo mutations. The rate at which novel alleles appear, and the proportion of synonymous, missense and nonsense mutations, and non-coding mutations are consistent across all three replicate evolutions.

### Comparison of population level mutations reveals clonal interference and widespread parallelism

Despite the large number of SNPs detected, only 17 alleles arose above a frequency of 98% across replicate evolutions ranging from 300-500 generations. Moreover, the maximum frequency of most alleles never exceeded 10% (**Fig. 2A**), and the vast majority of alleles were present at lower frequency in the final time-point than they were at their maximum (**Fig. 2B**). Together, the foregoing observations suggest that in each evolution experiment population dynamics was largely driven by clonal interference [40]. A small number of loci were recurrently mutated above what would be expected by chance, indicating that variants at these loci were likely beneficial (**Table 1**, **Table S2**). For example, a total of 212 mutations arose in the 10 most significantly mutated genes in the population sequencing data, with each gene receiving at least five mutations (**Table 1**). Moreover, 30 and 14 distinct allelic variants were discovered in just two: the genes encoding the DNA binding repressor GalS and the RNA-binding protein Hfq, respectively (**Table S3**). High-resolution population sequencing also revealed that 13 SNPs not present at the start of the experiment reached at least 1% frequency in all three vessels at various time-points, while 52 SNPs recurred in two out of three chemostats (**Table S4**). Thus, our data also provide compelling evidence for substantial parallel evolution at the genic level.

**Table 1.**
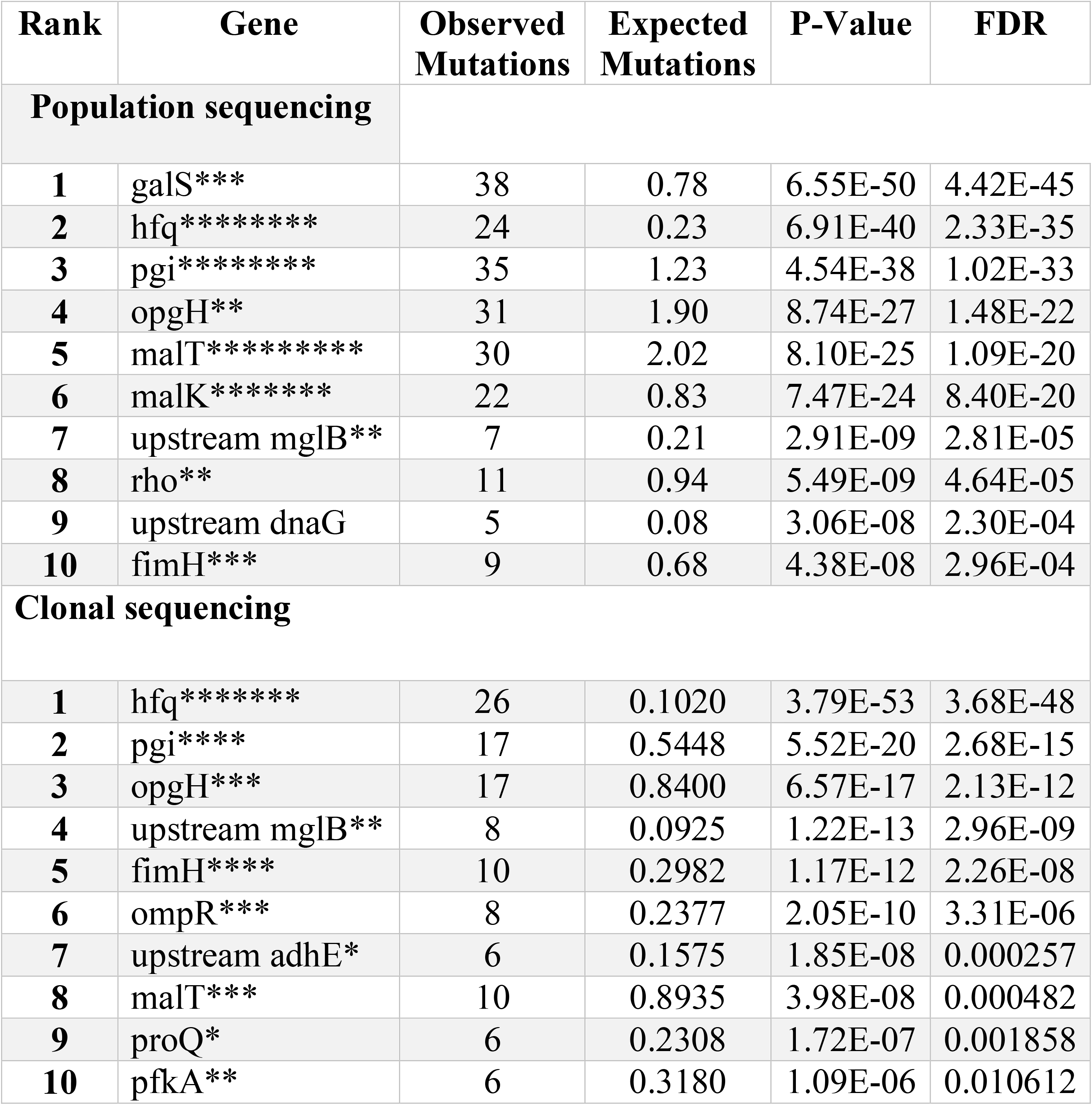
Characteristics of frequently mutated genes. Each asterisk indicates an allele that arose more than once independently, either within or between vessels.

**Fig. 2.**
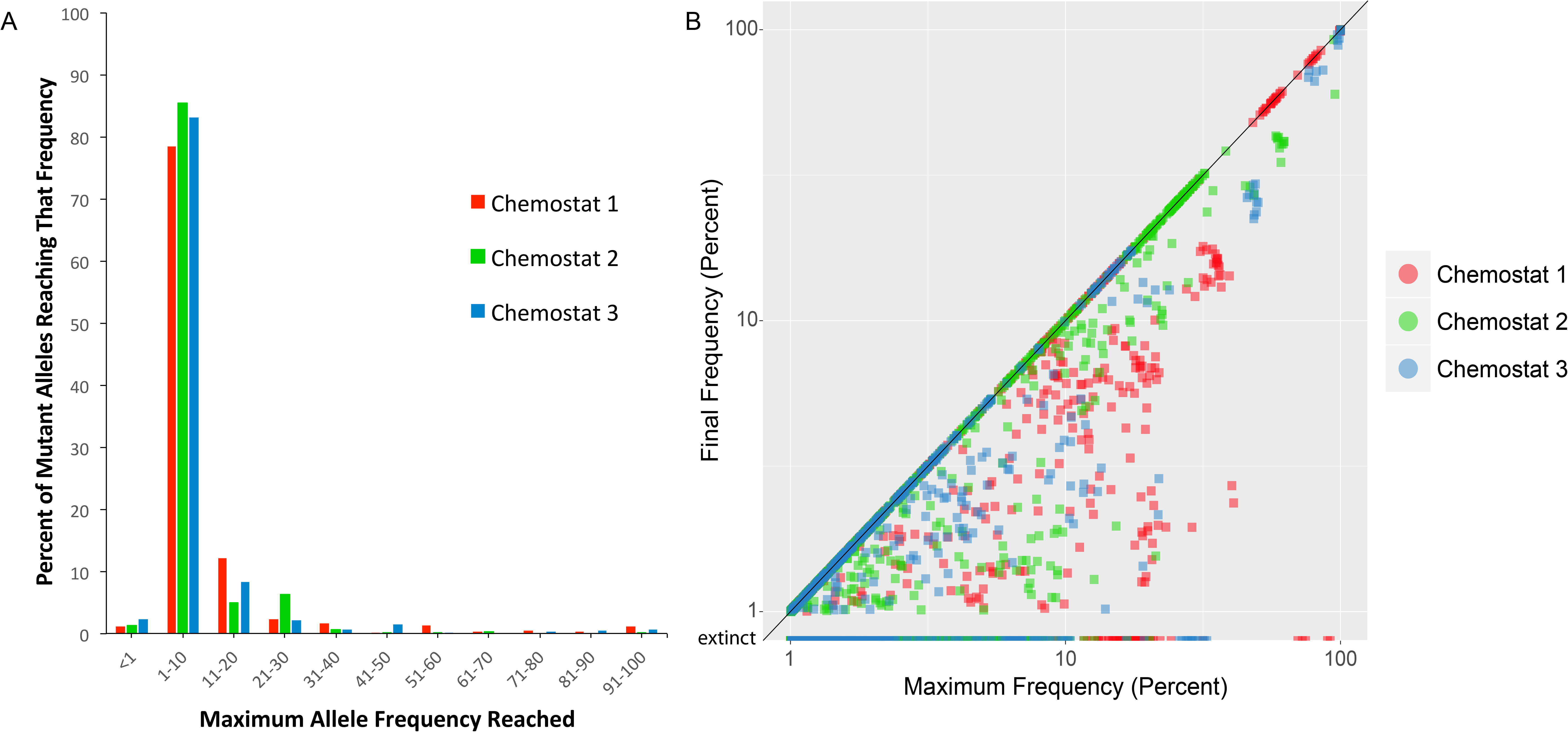
Most *de novo* mutations only reach low allele frequencies, and experience pervasive clonal interference. (A) Histogram of maximum allele frequencies from three replicate evolutions, (B) Final versus maximum allele frequency for each *de novo* mutation, shows most mutations are at a lower frequency at the end of the experiment than they were at their maximum.

### Clonal sequencing further clarifies lineage relationships and parallelism

To determine linkage relationships between the novel alleles, we sequenced 96 individual clones from each vessel. In each case, the 96 clones were isolated at random from the time-point at which we detected the greatest number of mutant alleles at ≥ 5% frequency. To assess whether the frequency estimates from population sequencing were reasonable, and whether the isolated clones constituted a reasonable subsample, we compared frequencies of mutations identified in both datasets at the corresponding time-point and found that they correlate well (**Fig. 3**).

**Fig. 3.**
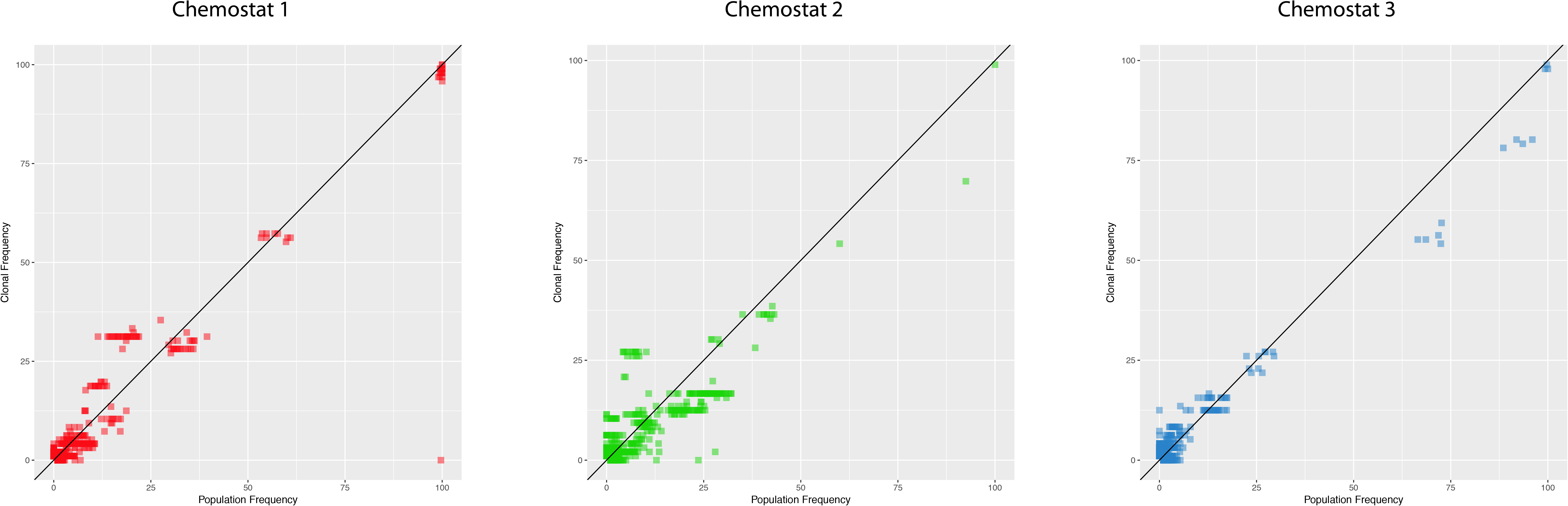
Isolated clones are representative of the populations from which they are drawn. Mutation frequencies for population and clonal sequencing for mutations identified in both datasets at the same time-point shows similar frequencies.

For each set of 96 clones, we constructed a phylogeny to represent their putative evolutionary relationships (**Fig. 4**). Inspection of the mutations and trees from each vessel (i.e. each independent evolution) revealed several instances in which exactly the same mutation arose not only in different vessels, but often more than once in the same vessel on distinct branches of a given tree. In the most extreme case, 6 of the 11 *hfq* alleles detected via clone sequencing were identified in clones from different vessels, indicating independent parallel origins (**Fig. 4, File S2**). Furthermore, 7 of the 11 appear to have arisen more than once within the same vessel.

**Fig. 4.**
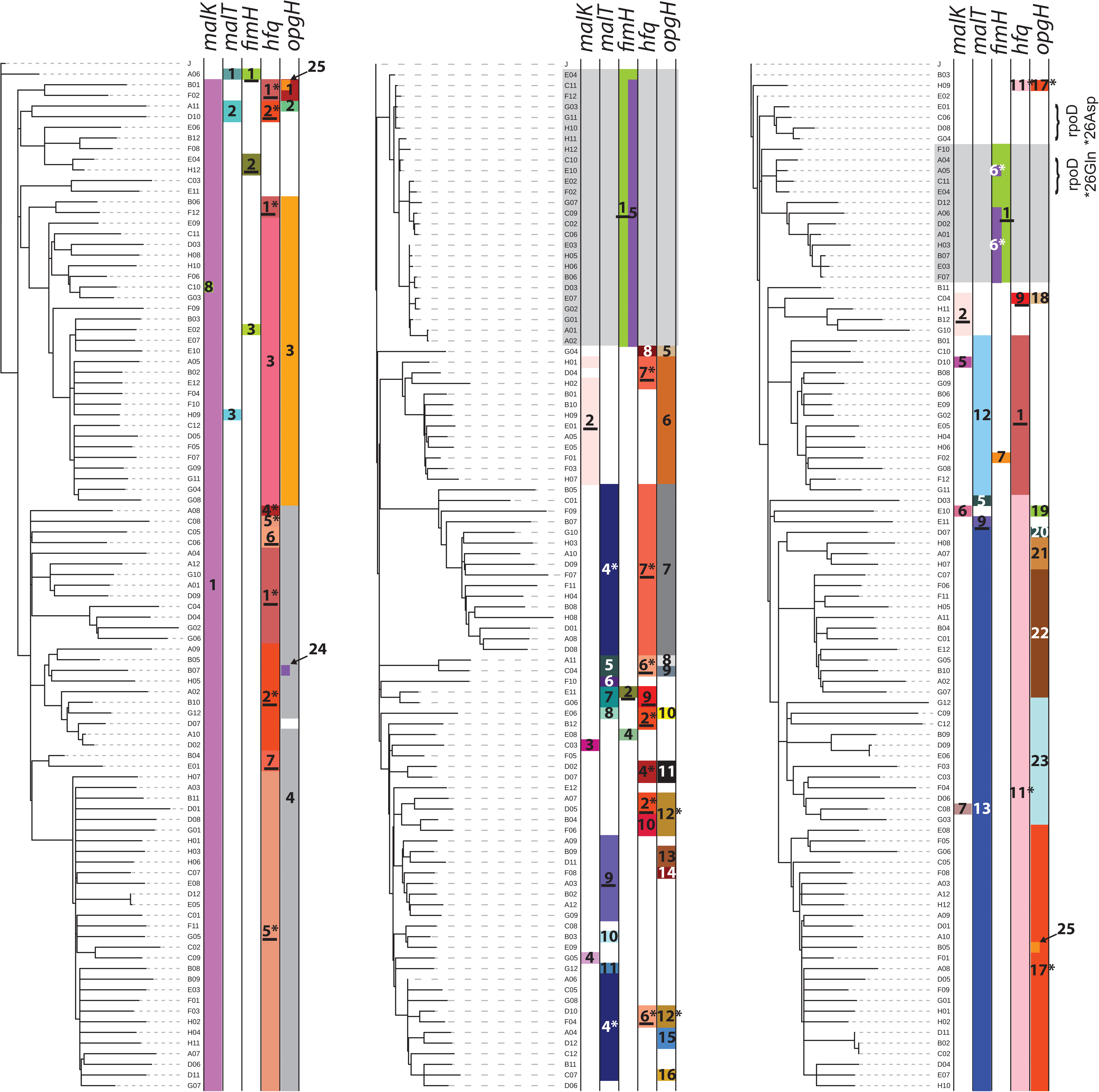
Clone phylogenies. Phylogenies depicting relationships among sequenced clones isolated from chemostats when allelic diversity attained its maximum. Distributions of different *malK, malT, fimH, hfq* and *opgH* alleles are indicated by colored bars. For each gene, all alleles observed in the dataset are numbered (see File S2 for details of which number corresponds to which allele for each gene). Underlined numbers denote alleles independently observed in more than one chemostat, while numbers marked with an asterisk appear to have arisen more than once within the same vessel. Grey shading delineates clades comprised of clones that have not acquired the standard mutations related to enhanced glucose uptake and instead carry variant *fimH* alleles that contribute to biofilm formation. Bracketed clones in chemostat 3 exhibited mutations expected to revert the ancestral nonsense mutations in the housekeeping gene encoding sigma factor RpoD.

### Clonal dynamics are shaped by relationships among de novo alleles, hard and soft selective sweeps, and absence of periodic selection

Combining population allele frequency data with linkage information derived from clonal sequencing makes it possible to depict lineage dynamics using Muller diagrams (**Fig. 5, Files S3-S5**). In general, we observe early, hard sweeps of highly beneficial mutations related to limiting nutrient influx, followed by soft sweeps [41-43] and multiple-origin soft sweeps that may fine-tune glucose uptake or utilization later in the experiment when diversity was higher [44-46]. Hard sweeps consistently involved mutations in regulators (*galS* in chemostat 1, transcriptional terminator *rho* in chemostats 1 and 3) or regulatory regions (upstream of *dnaG* in chemostat 1, upstream of *mglB* in chemostats 1, 2 and 3 – See supplementary Files 2, 3 and 4 for detailed dynamics), while soft sweeps were comprised of both regulatory and operon-specific mutations (e.g. *hfq* and *opgH* in chemostats 1, 2 and 3, upstream of *adhE* in chemostat 1, *pgi* in chemostat 3) (**Fig. 5, S1, Files S3-S5)** [42,47]. Here, we note that multiple-origin soft sweeps may be especially prevalent in our experiments due to the ancestral mutator allele at *mutY*, as the likelihood of concurrent identical mutations in the same gene should increase with mutation rate.

**Fig. 5.**
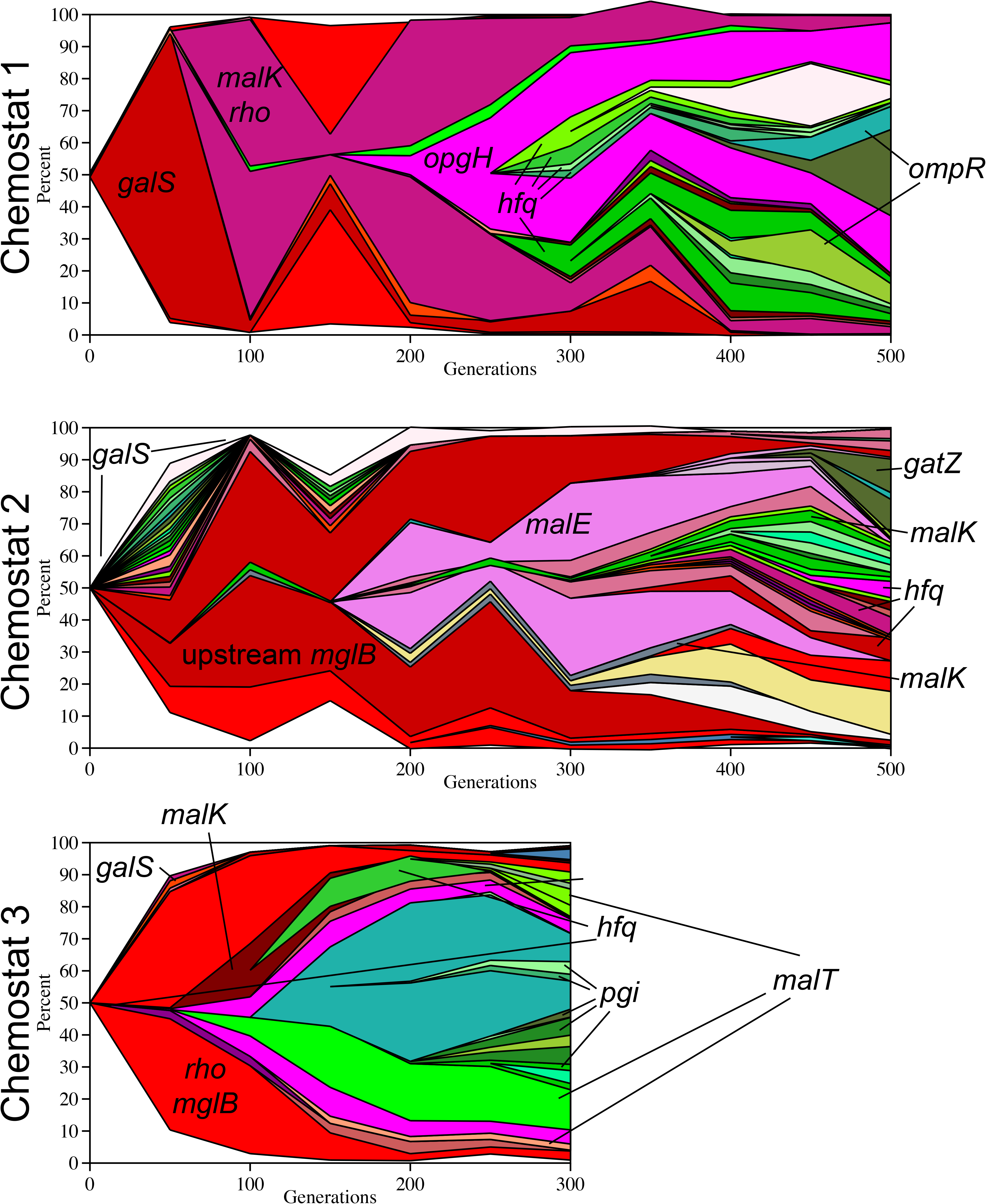
Muller diagrams. Evolutionary dynamics of adaptive lineages, deduced from combining whole-population whole-genome sequence data and whole-genome sequence data of individual clones isolated from each chemostat at the time-point where allelic diversity reached its maximum value. Select genes are indicated in the plots – see Figure S1 and Supplementary Files S3-S5 for additional details. Also note, most mutations that went extinct by the sampling timepoint are not shown. See Figure S1 for their relative frequencies.

With regard to periodic selection, rather than favorable alleles arising within a set of lineages that successively replace one another over time, we observe groups or cohorts of mutations co-evolving, with widespread clonal interference among lineages that carry different beneficial mutations [48]. For example, in chemostat 1, a spreading lineage with a cohort of mutations upstream of *mglB*/*lptA*/*opgH* (pink) is checked by the emergence of lineages carrying mutations in *hfq* (green) (**File S3)**. All of these phenomena – hard and soft sweeps, cohorts of mutations that increase or decrease in frequency together, and clonal interference – have been observed in yeast [14,19,49] and *E. coli* [23] populations evolving in the laboratory, as well as in *Pseudomonas aeruginosa* evolving in the cystic fibrosis lung [50].

### Early sweeps are related to influx of the limiting nutrient glucose

For specific growth rates between ∼ μ =0.1 hr^−1^ and μ =0.9 hr^−1^, glucose is most efficiently transported using a combination of the maltoporin LamB and the galactose transporter MglBAC, and glucose limitation frequently selects for mutations that increase expression of these proteins [45,51-59]. As expected, 7 of the top 10 frequently mutated genes/gene regions we observed (*galS*, upstream *mglB, malT, malK, hfq, rho* and upstream *dnaG*) play a role in transcriptional regulation of *lamB* or *mglBAC*, either directly or through their interactions with global regulators (**Table 1**, **Fig. 6**).

**Fig. 6.**
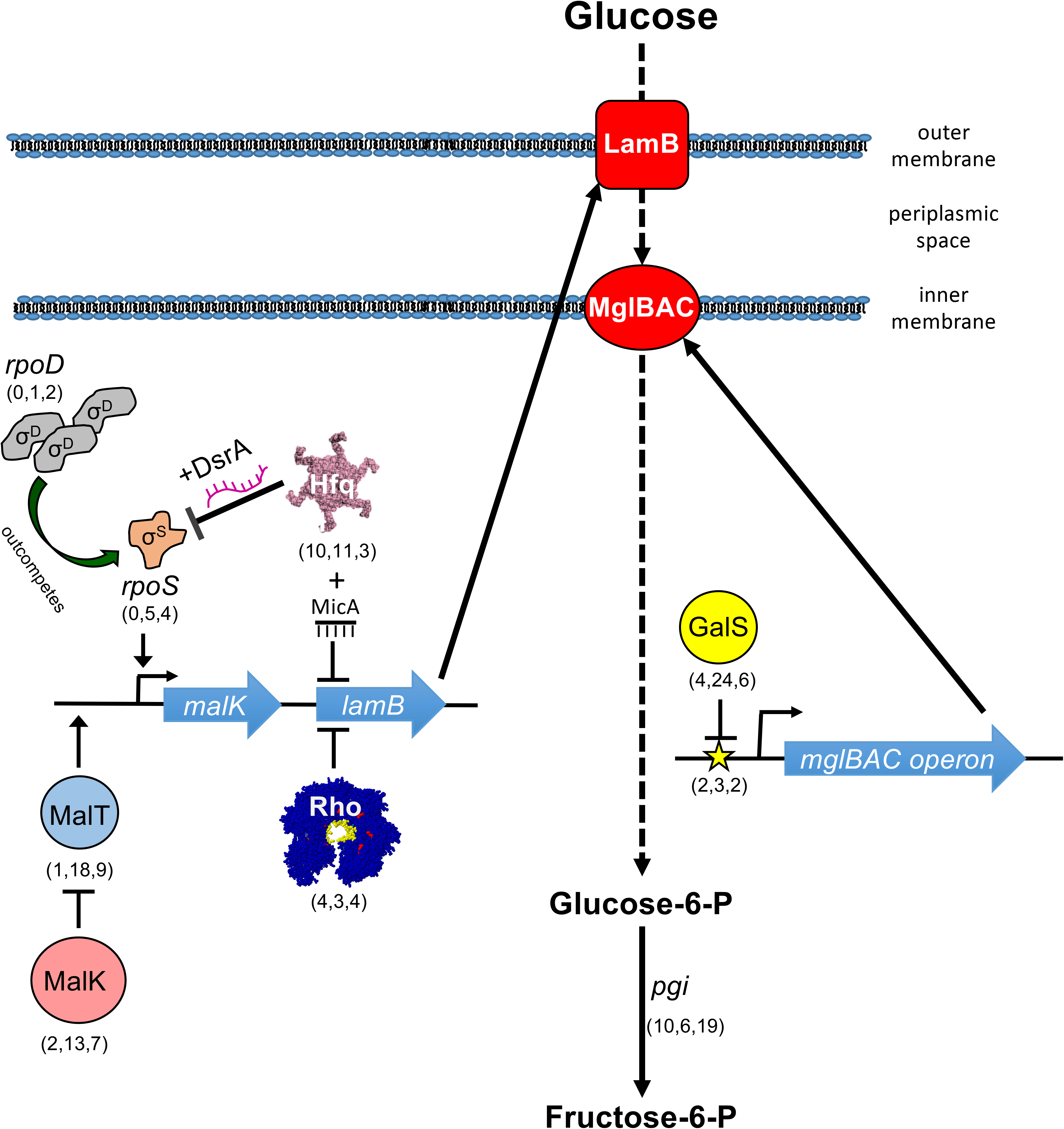
Overview of pathways relating some of the most frequently mutated genes to glucose transport and metabolism. Numbers in parentheses next to protein/gene names denote the number of mutant alleles found in each chemostat population over the course of 300-500 generations (also see Table S3).

Mutations in *gal*S and upstream of *mglB.* Thirty different alleles of *galS* (encoding GalS, a negative regulator of *mglBAC* transcription) were detected over the course of our experiments. These spanned the length of the gene, and the majority caused missense amino acid changes likely to disrupt *mglBAC* transcriptional repression and augment glucose flux across the inner membrane (**Fig. 6**) [60]. Despite the large number of alleles we observed, few persisted beyond generation 50 or attained a final frequency greater than 5%, demonstrating high clonal diversity early in the experiment. In chemostats 2 and 3, no clear “winner” *galS* genotype emerged, though in chemostat 1, a GalS allele (Arg146Leu) swept to near fixation (89.6% of the population at generation 50).

Early-arising *GalS* mutant genotypes were rapidly displaced by clones carrying highly-beneficial mutations in the *mgl* operator sequence upstream of *mglB* (**Fig. 5**). This sequence of events, like the mutations themselves, has been observed elsewhere [45,61]. The most successful mutation upstream of *mglB* (bp 2,238,647 C→A) occurred early in every vessel, and in every case increased in frequency to over 90% of the population (**Table S5**, **File S3-S5**). Notably, this same mutation was observed in chemostat-grown *E. coli* by Notley-McRobb et al. (1999) as well as in the experimental population described by Helling et al. (1987), where it was found to be the only SNP shared by all members of a cross-feeding consortium [34,54].

The dynamics of *galS* replacement illustrates the effect that clonal interference can have on the fate of different alleles. In chemostat 1, clones carrying GalS Arg146Leu rapidly dropped in frequency when lineages emerged with a mutation upstream of *mglB* (position 2,238,647), but were not completely displaced until generation 400 and even enjoyed brief periods of expansion. In chemostat 2, clones with the same mutation upstream of *mglB* were present by generation 50, but did not surpass a 90% threshold for another 250 generations due to competition from 22 different *galS* lineages and a lineage carrying a different upstream of *mglB* allele (2,238,648 G→T) (**Fig. 5, S1B, File S4**). By contrast, in chemostat 3, a lineage with the upstream *mglB* mutation (2,238,647) experienced little competition and was almost fixed by generation 150 **(Table S5).**

Over the remainder of the experiment only three other mutations upstream of *mglB* mutations reached the threshold for detection: two were within 2 base-pairs of the first mutation and did not rise to high frequency, while the third (chemostat 1, 2,238,630 C→A) located in the CRP activator binding site, co-occurred with 2,238,647 C→A and increased to ca. 80% frequency by generation 500 (**Figs. 5, S1A, File S3**). This dynamic suggests additional mutations that affect GalS repressor binding are not of great benefit after the preferred allele has swept the population, whereas mutations that modulate the activity of other regulators (i.e. CRP) can act synergistically.

The dynamics of lamB regulation. LamB glycoporin overexpression is a hallmark feature of *E. coli* populations adapted to glucose-limited chemostat growth [34,45,53,55,56,62]. Previous experiments have shown that under glucose limitation, overexpression of LamB can be the result of any one of the following: constitutive activation of transcriptional regulator MalT, disruption of the MalT inhibitor MalK, mutation of the RNA chaperone Hfq, alteration of sigma factor dynamics (σ^S^/ σ^D^ ratio), or mutation of the *malT* repressor Mlc, (**Fig. 6**) [34,45,53-56,62,63].

Across the three replicate evolutions, we observed 19 unique *malT* alleles and 14 unique *malK* alleles (**Figs. 6, S2, Table S3**). Over half of the mutations in *malT* (10 out of 19) are known either to cause MalT to become constitutively active, or to occur in amino acids involved in MalT/MalK interaction [45,64,65]. A single MalK mutation (Ala296Asp) rose to high frequency early (94% by generation 100) in chemostat 1 (**Fig. S2**, **Table S5**). This SNP is in a regulatory domain likely to be the site of MalK/MalT interaction [66]. Alteration of a neighboring residue (Asp297) has been previously shown to allow unregulated transcription of the *mal* operon [66].

*malT*/*malK* allele dynamics differ among experimental populations. As mentioned above and shown in **Fig. S2**, MalK Ala296Asp sweeps early in chemostat 1, whereas in chemostat 3, early MalK mutations (blue) are displaced by later mutations in MalT (green). In chemostat 2, the picture is quite different: clones with either *malK* or *malT* mutations co-exist through all 500 generations. The reason for this contrast in allele frequency dynamics cannot be attributed to emergence of a single “most fit” allele, as the majority types from chemostats 1 and 3 arose independently in chemostat 2, but did not sweep. Despite the importance of MalT and MalK as high-value targets for selection during adaptation to glucose limitation, other advantageous mutations (upstream *mglB, rho* and *hfq*, discussed below) may have ultimately carried “winning” *mal* alleles in chemostats 1 and 3 to higher frequency, purging allelic diversity at this locus. Interestingly, although we observed 30 *malT* and 22 *malK* mutations in the population sequencing data (**Table 1**), in only 5 out of the 288 sequenced clones do mutant alleles of these two genes co-occur, suggesting that there may be no additional advantage or even some disadvantage to having both. In the Helling et al. evolution experiments [30], which were founded by the same ancestor used here, secondary resource specialists share a mutation in MalT, whereas the primary resource (glucose) specialist that feeds those clones carries a mutation in MalK [34].

selection of mutations in RNA chaperone Hfq that affect translation of lamB and stationary phase transcription factor RpoS. Hfq is a global regulatory protein that facilitates translation and/or RNA degradation by mediating ncRNA-mRNA interactions. It participates in a diverse range of cellular processes including nutrient uptake, motility and metabolism [67]. *hfq* mutations identified in other glucose-limited evolution experiments exhibit pleiotropic physiological effects: they appear to increase translation of LamB glycoporin, reduce levels of stationary phase transcription factor RpoS, inhibit cellular aggregation, and enhance glucose transport via PtsG [63,68].

*Hfq* is one of the most frequently mutated genes observed in our experiments: 24 *hfq* mutations, resulting in 14 distinct *hfq* alleles, were detected via population sequencing; by the end of our evolutions >50% of each population carried a mutation in *hfq* (**Table S3**). (**Table 1**). Two of these alleles arose independently in all three vessels (same nucleotide position, same SNP), and six additional alleles were observed in two of three vessels (**Table S4**). The frequency of and parallelism exhibited in *hfq* mutations is particularly curious in the context of experiments by Maharjan et al. in which *hfq* mutations arise, but are at low frequency and subject to negative frequency-dependent selection and epistatic interaction with mutations in *rpoS* [32,52,62,63].

The dynamics of hfq mutations are variable across evolutions and may depend on which other beneficial alleles are present in the same lineage or in the same population. In chemostats 1 and 2, a large number of *hfq* alleles (10 in chemostat 1 and 11 in chemostat 2) appear after generation 250 and are preceded by mutations in *malK* or *malE* and the *opg* operon. The most successful Hfq allele in chemostat 1, Val62Phe, occurred in a sweeping lineage with a secondary mutation upstream of *mglB* (discussed above) and may have been carried along by association. In chemostat 3, a single *hfq* mutation arises early (Ser60Tyr, present by generation 100), sweeps to near fixation alongside MalT Met311Ile and is closely followed by mutations in *opgH* (**Fig. S1, File S5**).

Recurrent mutations occur in Rho. Early-arising mutations in the *rho* termination factor are a conspicuous feature of chemostats 1 and 3 (**Figs. 5, S1**). Rho is required for transcriptional termination of up to 50% of cellular mRNAs [69,70] and can participate in gene regulation via intragenic terminators [71]. Mutagenesis and ChIP-chip analyses have identified Rho-dependent terminators within multiple genes relevant to glucose limitation, specifically *lamB, mglA*, and *mglC* and downstream of *malT* and *mglC* [71,72]. In fact, it has long been known that defective LamB expression in MalT activator mutants can be restored via compensatory mutations in *rho* [73]. In chemostats 1 and 3, Rho mutations fix or nearly fix early and do so in concert with mutations in MalK (chemostat 1 Ala296Asp) and mutations upstream of *mglB* (chemostats 1 and 3, bp 2,238,647) (**Figs. 5, S1, Table S3, Table S5, Supplementary Files 2 and 4**). Conversely, in chemostat 2 only three *rho* alleles were detected, none of which rose in frequency to >6% of the population (**Fig. S1, Table S3**).

### Mutations that impact energy conservation, membrane biogenesis and cell adhesion are late arising targets of selection

Phosphoglucoseisomerase (Pgi) is an abundantly expressed central metabolic enzyme responsible for converting glucose-6-phosphate into fructose-6-phosphate. Knockdown of *pgi* mRNA alleviates catabolite repression [74], favoring increased expression of CRP-regulated genes such as *lamB* and *mglBAC*. Twenty-four unique *pgi* alleles were detected over the course of our three replicate evolutions. However, few rose to appreciable frequency before generation 200, suggesting their benefit may be contingent on the presence of other mutations or some aspect of the chemostat environment that consistently changed after this time point. *Pgi* alleles were least successful in chemostat 1, which was also the only replicate in which a large fraction of clones (79% by generation 500) acquired a second mutation upstream of *mglB*. This observation suggests that *pgi* mutations and mutations in the CRP-binding site of the *mglBAC* promoter may be functionally redundant.

Membrane glycosyltransferase OpgH is involved in the synthesis of periplasmic glucans, highly branched oligosaccharides made from β-linked glucose monomers. While we do not observe *opgH* mutations earlier than generation 100, they rapidly increase in frequency once they appear, usually either just before or just after *hfq* mutations (**Fig. S1, S4, File S3-S5**). Novel *opgH* alleles, especially the nonsense mutations that we frequently observe, may constrain glucan production and serve as a glucose conservation measure. A “moonlighting” function has also recently been reported for OpgH: the glucosyltransferase interacts with the tubulin-like cell division protein FtsZ to delay cell division when levels of UDP-glucose are low [75]. Thus, mutations in OpgH may augment the rate of cell division, and thereby provide a fitness advantage under slow-growth chemostat conditions. The only *opg* operon mutation identified among strains in previous Adams et al. experiments occurred in *opgG* of the glucose scavenger, CV103 (E487*) [28].

Mutations in Rho-independent terminator T_1_ that allow run-through transcription may tip the balance between competing sigma factors. Sigma factor RpoD (σ^70^) is the predominant sigma factor associated with RNA polymerase during exponential growth. As cells enter stationary phase, transcription of the gene for alternate sigma factor RpoS (σ^S^) increases [76]. *rpoS* mutations are often selected for under continuous glucose limitation as they allow continued transcription from promoters negatively regulated by σ^S^ but required for glucose uptake and metabolism (e.g. [77,78]).

In chemostat 1, a mutation in the *rpsU-dnaG-rpoD* macromolecular synthesis operon upstream of *dnaG* (bp 3,209,081 G→T) was present in over 90% of the population by generation 50 (**Table S5**). This SNP decreases the stability of the rho-independent terminator T_1_ situated between *rpsU* and *dnaG*, and thus may be expected to increase expression of RpoD [79] and as a result operons positively controlled by σ^70^ (e.g. *mglBAC* and *malK-lamB-malM*). A T1 mutation (bp 3,209,075 C→A) was also shared among Helling et al. strains, defining the lineage that gave rise to three of four consortium members [30,34]. In addition, in Chemostat 2, we observed an ∼150kb duplication that included *rpoD* and in chemostat 3, eight clones out of 96 carried intragenic suppressor mutations of the ancestral nonsense allele (*26Asp and *26Gln) in RpoD.

Fimbrial protein genes (*fim*) Genes associated with production/function of type 1 fimbriae, particularly *fimH* (fimbrial adhesion), were an unexpected and frequent target of mutation in all three chemostats (**Table 1**, **Figs. 4**, **S1, Table S3, Files S3-S5**). Though novel *fim* alleles were transient in vessels 2 and 3, in chemostat 1 a FimH Asn54Lys variant rose to a frequency of 70% by generation 150, temporarily displacing high-fitness alleles in *rho, malK* and upstream *mglB* (**File S3**). Because *fimH* mutants demonstrated an increased capacity for biofilm formation (data not presented), a recurrent issue in chemostat experiments, but did not acquire any of the mutations expected to enhance glucose metabolism, *fimH* mutations were likely related to chemostat persistence rather than to competition for limiting substrate.

## Discussion and Conclusions

### History matters: ancestry influences evolutionary trajectory

The tempo and trajectory of a clonal population depend on its genetic point of departure. Our departure point was a founder that harbored nonsense mutations in mismatch repair (MutY, Leu299*), and in housekeeping and stationary phase sigma factors (RpoD, Glu26* and RpoS, Gln33*), but also carried an amber/ochre/opal nonsense tRNA suppressor. Populations originating from such a founder would not only have an increased mutational load but also the capacity to tolerate those mutations, in particular nonsense mutations that would otherwise result in complete loss-of-function.

Laboratory evolution studies have borne out the idea that loss-of-function mutations can be significant drivers of adaptation [20,80-82]. Metabolic network re-programming via modulation of existing function can occur much faster than the evolution of new pathways via mutation [81], and in many cases nonsense mutations or deletions confer greater fitness benefit than missense mutations affecting the same gene [82]. However, loss of function often comes at the expense of metabolic flexibility, limiting the ability of evolved clones to compete in alternative environments [20]. RpoS has been shown to be a high-value target of selection under nutrient limitation: under low-nutrient conditions RpoS normally outcompetes RpoD for binding to RNA polymerase, repressing genes required for growth and cell division and activating those required to enter stationary phase [78,83]. *rpoS* mutants thus continue to divide under conditions where wild-type cells arrest. In this respect, our genetic ‘point of departure’ could be viewed as being pre-adapted to life under glucose limitation. However, the combined phenotypic effect of ancestral *rpoS* and *rpoD* nonsense mutations in a suppressor background is murky and raises the question of whether this combination of mutations is favorable under glucose limitation, merely tolerated or detrimental. Despite the fact that many changes we observed (*galS*, upstream *mglB, hfq*) enhance glucose assimilation, are predictable, occur repeatedly and rise to high frequency, we also saw the persistence of clones with none of these mutations that instead carry intragenic suppressors of the nonsense mutation in *rpoD* (*26→Asp and *26→Gln, chemostat 3 **Fig. 4**) or a duplication that includes *rpoD*. If rapid adaptation can be driven by loss-of-function but occurs at the expense of metabolic flexibility, nonsense mutations have a distinct advantage over deletions in that reversion or suppression is possible should environmental conditions change [20].

Another ancestral allele that we expected to influence evolutionary trajectories was an A→T CRP binding site mutation 224 bp upstream of the acetate scavenging enzyme, *acs* (acetyl-CoA synthetase). This mutation alters regulation of the *acs-pta* operon such that the ancestor poorly assimilates acetate excreted during growth under continuous glucose-limitation, opening up a secondary resource for novel mutants that can [34]. Here, we uncovered no evidence for the type of cross-feeding described in prior reports [29,30,33]. This result was not unanticipated, as evidence for cross-feeding polymorphisms was observed in only half the evolution experiments founded by this ancestor or its close relatives [33]. Moreover, a recent model [31] defining the boundary conditions for cross-feeding to evolve in a chemostat showed that such an outcome is sensitive to variation in dilution rate as well as to the relative fitness of *de novo* mutants that gain access to secondary metabolites. Subtle differences in either of these parameters may account for why we saw no evidence for acetate/glycerol/formate cross-feeding in our experiments. The absence of such interactions may also be due to the fact that no variants arose at loci where mutations have been implicated in cross-feeding evolution: *acs* (acetyl CoA synthetase), *lpd* (lipoamide dehydrogenase) and *ptsI* (phosphoenolpyruvate phosphotransferase).

As expected, the ancestral *acs-pta* defect resulted in appreciable levels of residual acetate (∼45-90 μM) at the onset of our experiments (**Fig. S3**). While we uncovered no evidence for the evolution of secondary resource specialists [30,31,33] and refs therein), residual acetate levels consistently fell below detection limit by generation 200. Thus, adaptive mutants arising here found other ways than cross-feeding to metabolize all available carbon. One possible work-around may involve the *pgi* locus, which was second only to *galS* in the total number of mutations recovered (**Table S2**). Generation 200 coincides with the emergence of mutant *pgi* alleles in all three populations. In chemostat experiments with *pgi* deletion mutants, Yao et al. found that in the absence of Pgi, glucose uptake rate drops slightly compared to wild-type, but no overflow acetate is produced and biomass yield is unchanged [84].

### Population and clone sequencing open up a detailed view of the full spectrum of beneficial mutations and how that spectrum changes over time

High-coverage, whole-genome, whole-population sequencing makes it possible to discover every new allele reaching >1% frequency in a population of >10^10^ cells. Because alleles are highly unlikely to reach such frequencies by draft, all were either transiently beneficial or hitchhiking with alleles that were. This depth of analysis opens up a richly-detailed view of the spectrum of beneficial mutations arising in *E. coli* under constant resource limitation. Periodic whole-population sequencing allows patterns to be discerned as to how these spectra change over time, while clone sequencing makes it possible to establish linkage relations among novel alleles and represent their collective fate as evolving lineages. Multiple patterns emerge from these analyses. *First,* new alleles accumulate in replicate populations at similar rates, and the proportion of alleles that are missense, nonsense, synonymous, or noncoding remains fairly constant; the great majority is either missense (60-70%) or nonsense (5-10%). *Second,* the distribution of new mutations across the genome is skewed, with only a few dozen of the more than 1,000 mutated genes having a significant number of mutations; yet even among these most frequently mutated genes, few *de novo* mutations fix. *Third,* by clonal sequencing we are able to determine that many, independent linages co-exist and compete within the culture. Thus, evolutionary dynamics in these populations is governed by clonal interference rather than by clonal replacement or reinforcement.

A *fourth* pattern to emerge is widespread parallelism in regulatory evolution. Both across and within populations, the same genes are mutated again and again, often at exactly the same nucleotide position in independent replicates, and sometimes in independent lineages co-evolving in the same vessel. Many of these genes (*galS, malT, malK*, upstream *mglB, hfq, rho*) act in processes related to the transport and assimilation of the limiting nutrient, glucose. However, in most cases the mutations recovered alter regulation of these processes, and not the structural proteins that carry them out.

A *fifth* pattern relates to the order of beneficial mutations and the influence that order has on dynamics. Consistent with previous reports, mutations that increase glucose flux across the inner membrane (*galS*, upstream *mglB*) occur early and precede those that increase flux across the outer membrane (*malK*/*malT, hfq, rho*). In both cases, mutations in binding partners (GalS/upstream *mglB* and MalT/MalK) rarely occur in the same clone, and the order in which they occur can lead to either a sweep (upstream *mglB* clones quickly displace *galS* clones) or clonal interference (*malT* and *malK* clones can co-exist). Other alleles appear to emerge later in the experiment and nearly always together: clones with existing mutations in the *mal* operon acquire subsequent mutations in *hfq* and *opgH*, regardless of which gene is altered first and which alleles are already present in the population **(Fig. S1)**. These patterns are echoed by the genotypes reported by Kinnersley et al. [22] in which glucose scavenger CV103 has mutations in *malK, opgG* and *hfq* while acetate specialist CV101 only carries a mutation in *malT.*

Similar experiments carried out by Maharjan et al. [32] using *E. coli* BW2952 demonstrated that under continuous glucose limitation population-level phenotypic changes are often the result of multiple soft sweeps by combinations of beneficial mutations. While we did not assay clone phenotypes, multiple alleles of *galS, hfq* and *opgH* appear to sweep our populations in concert suggesting a similar pattern in which a phenotypic effect (reduced expression of a particular gene) is favored, but has different genetic bases in co-existing lineages. At the clone level, BW2952 also exhibits sign epistasis between mutations in *rpoS*/*hfq* and *galS*/*malT* [32,52]. In our experiments, we did not uncover evidence of sign epistasis between the ancestral *rpoS* allele and *hfq*: by generation 250, over 50% of clones in populations 1 and 3 carry mutations in both genes. Maharjan et al (2013) proposed that fitness deficits exhibited by *rpoS*/*hfq* double mutants may be the result of altered cell division [62,85] and that *hfq* mutations enhance glucose uptake during slow growth, but diminish viability when cells are dividing rapidly. Hfq deletion mutants exhibit cell division anomalies due to elevated expression of cell division proteins, including FtsZ [86,87]. Recent work by Hill et al. (2013) has shown that during fast growth OpgH (which in our experiments is nearly always mutated alongside *hfq*) binds FtsZ to postpone cell division. Thus, it may be that negative fitness effects experienced by *hfq-rpoS* double mutants are the result of cell division anomalies mitigated by mutations in *opgH*. It is noteworthy in this regard that whereas cells in the Maharjan et al. experiments experienced a dilution rate of D=0.1 hr^−1^, those in evolutions performed by Adams, Helling and colleagues were doubling twice as fast (D=0.2 h^−1^). Thus, this discrepancy may be a manifestation of trade-offs between glucose uptake and cell viability.

Finally, some mutations occur repeatedly and are likely beneficial, but their dynamics are unpredictable (e.g. beneficial mutations in transcriptional terminator *rho* sweep when they co-occur with beneficial mutations upstream of *mglB,* but otherwise remain at low frequency (**Table S3, Fig. S1**). This dependence on genetic context, or “quasi-hitchhiking”, of beneficial mutations was previously observed by Lang et al. (2013) in evolving yeast populations and may be consistent feature of microbial evolution experiments that becomes observable when populations are sequenced at high depth of coverage and sufficient temporal resolution [14].

### The evolution of population genetic complexity

Szostak, Hazen and others [88,89] argue that a biological system’s complexity should be evaluated in terms of its functional information content. Although the total number of alleles in an evolving population at any given time-point is information content, it is functional only in how it is integrated among the lineages co-existing at that time-point. Our approach of integrating population sequencing with clone sequencing makes it possible to estimate the pace and extent with which complexity, measured as lineage-specific functional information [88,89], emerges in replicate evolving populations originating from a common ancestor. Implicit in our perspective is the assumption that the sequence differences by which lineages can be distinguished have physiological and fitness consequences. For each population, we calculated at 50-generation intervals three measures of complexity: Shannon’s Entropy (H), equitability (H/H_max_) and normalized population richness (Lineage counts) (**Fig. S5**). All three measures increased during the course of evolution, but with a different tempo in each population. Complexity increased in population 3 with no indication of reaching an asymptote by generation 300 when the experiment was terminated. Populations 1 and 2 reached asymptotes after ∼400 generations, following a steady increase in population 1, but a nearly-300 generation period of stasis in Population 2. While longer experiments are clearly called for, our finding that longer-term experiments appear to reach an asymptote in complexity is consistent with theoretical [90] and empirical [91]observations that fitness plateaus when microbes are cultured by serial transfer or in chemostats, even starting with mutator strains [92], and that complexity itself may plateau when its evolution is simulated using RNA-like replicators [93]. It is intriguing to contemplate the possibility that there may be a limit to the level of clonal interference that can be sustained in asexual populations once all avenues for large fitness gains have been exhausted. Indeed, it was recently shown using lineage tracking, that while fixation of an adaptive mutant causes a stochastic crash in diversity, the generation of new adaptive mutants within such a fixing lineage is expected to generate new diversity, such that a longer term steady state level of diversity will be achieved [94].

Previous evolution experiments founded with the ancestor used here often resulted in stable sub-populations supported by cross-feeding. In the present experiments neither the spectrum of observed mutations nor the structure of clone phylogenies provides evidence for the evolution of this type of trophic interaction. Instead, the possibility of a plateau in complexity, coupled with the finding that every population has driven residual metabolites close to their detection limit, suggest that these populations converge on an adaptive peak by diverse mechanisms but that clonal interference keeps adaptive lineages off the summit, confined to exploring the many roads by which the summit can be approached.

## Materials and Methods

### Strains, media and culture conditions

*Escherichia coli* JA122, population samples and clones were maintained as permanent frozen stocks and stored at −80°C in 20% glycerol. Davis minimal medium was used for all liquid cultures with 0.025% glucose added for batch cultures and 0.0125% for chemostats, as previously described [34]. Chemostat cultures were initiated using colonies picked from Tryptone Agar (TA) plate inoculated with JA122, and outgrown in Davis minimal batch medium overnight. Chemostats were maintained at 30°C with a dilution rate of ≈ 0.2 hr^−1^ for 300-500 generations. Every other day culture density was assessed by measuring absorbance spectrophotometrically at A_550_. Every other day, population samples were archived at −80°C, and assayed for purity by plating serial dilutions on TA and examining Colony Forming Units (CFU) that arose following 24-hr incubation at 30°C. When necessary, chemostats were re-started from frozen stocks (chemostat 1: generation 217; chemostat 2: generation 410; chemostat 3 generation 251). At each sequencing time-point, 50 mL of culture was pelleted then stored at −80°C for DNA extraction. For clone sequencing, entire colonies were picked from TA plates inoculated from glycerol stocks, and re-archived in 96-well plate format.

### Metabolite assays

10 mL of sterile, cell-free chemostat filtrate was concentrated 20-fold by lyophilization (Labconco 4.5 Liter Freeze Dry System), then re-suspended in 0.5 mL sterile Millipore water. Residual glucose and residual acetate concentrations were determined on concentrated filtrate. Glucose was assayed enzymatically using the High Sensitivity Glucose Assay Kit (Sigma-Aldrich, Cat# MAK181), while acetate concentration was determined using the Acetate Colorimetric Assay Kit (Sigma-Aldrich, Cat# MAK086). Results presented in **Fig. S3** represent means ± SEM of duplicate assays.

### Population sequencing

Bacterial DNA was prepared using the DNeasy Blood and Tissue Kit (Qiagen, cat. 69504) following the manufacturer’s guidelines. For population sequencing, 5 × 10^10^ cells, collected from every 50 generations in three chemostat vessels (up to 500 generations in vessels 1 and 2, and up to 300 generations in vessel 3, 29 samples total) and frozen as pellets, yielded 10-20µg of DNA. Following Proteinase K treatment, RNaseA treatment was used (20µL 10mg/mL RNAse A, 2 min at room temperature) to avoid degraded RNA from visually obscuring size selection during library preparation. Samples were split into two columns to avoid overloading. Bacterial DNA was sheared to a 150-200bp fragment size using a Covaris S2 series sonicator (6min, Duty=5%, Intensity=3, Cycles/Burst=200), and was then ligated to barcoded adapters as described [95], except that 200bp fragments were size selected after adapter ligation (to maximize the fidelity of sequencing, by reading each fragment in both directions). Six barcoded libraries were combined and sequenced on each lane of HiSeq 2000 Illumina Sequencer.

### Variant calling from population sequencing with CLC Genomics Workbench 7.5

Illumina reads were trimmed (removing adapters on both ends) and stringently mapped (Mismatch cost 2, Insertion cost 3, Deletion cost 3, Length fraction 1.0, Similarity fraction 0.97) to the reference sequence (WIS_MG1655_m56). Variants were called with the following parameters: minimum frequency 1%, minimal coverage 100, minimum count 2, and base quality filtering (neighborhood radius 5, minimum central quality 15 and minimum neighborhood quality 20). Sequencing data uncovered low-level contamination of whole population samples with *Serratia liquifaciensis.* We therefore first determined proportion of contaminating reads by mapping population sequencing to *S. liquifaciensis* genome and then removed SNPs with frequency closely tracking percentage of contamination (between 1 and 5%) that matched *S. liquifaciensis* sequence.

### Selection of clones for sequencing

Allele frequencies for each chemostat were examined at each time point, and the time-point at which there was the largest number of alleles present at 5% or greater frequency was chosen for the isolation of clones for whole genome sequencing. The rationale for this was that it would afford us the greatest opportunity to phase as many high frequency alleles as possible.

### Clonal DNA preparation

A colony was re-suspended in 300µL of sterile ddH_2_0 with 17% glycerol and stored in three aliquots at −80°C. 100 µL of glycerol stock were used for DNA preparation. After removing glycerol (using MultiScreen High Volume Filter Plates with 0.45 μm Durapore membrane, Millipore MVHVN4525), cell were resuspended in 500µL LB and grown overnight at 30°C without shaking in deep well plates. Cells were collected again using filter plates and subjected to DNeasy 96 Blood and Tissue Kit (Qiagen 69581) (yielding 4-15μg per strain).

### Clonal libraries preparation and sequencing

Multiplexed sequencing libraries from clones were prepared using the Nextera kit (Illumina catalog # FC-121-1031 and # FC-121-1012) as described in [96], starting with 1-4ng of genomic DNA. Resulting libraries from each 96-well plate were pooled at an equal volume. Resulting pooled libraries were analyzed on the Qubit and Bioanalyzer platforms and sequenced on HiSeq 2000 (one lane per 96 clone pool). All raw sequencing data are available from the SRA under BioProject ID PRJNA517527.

### Variant calling from clonal sequencing with CLC Genomics Workbench 7.5

Short reads with adapters removed were mapped to the reference with the same parameters as above, except the length fraction was set to 0.5, and the similarity fraction to 0.8. Variants were called with a minimum frequency 80%, minimum count 2 and the same base quality filtering as above.

### Generation of phylogenies

For each chemostat, SNP and indel events for all 96 clones and the ancestor JA122 were concatenated and re-coded as binary characters (i.e. presence/absence with the ancestral state composed of all zeroes) and assembled into character matrices. PAUP ver. 4.0a149 was used to generate Camin-Sokal parsimony trees using the ancestor as the outgroup under the assumption that reversions were extremely unlikely due to the extreme transversion bias [97,98]. Tree files (.tre) were loaded into the Interactive Tree of Life (iTOL) web service for character mapping and figure generation [99].

### Determining genes with an excess of mutations

To identify genes with an excess of mutations, we first determined the overall density of mutations as:

ρ = M/L, where M is the total number of mutations, and L is the length of the genome. The probability of a given mutation landing in a segment of length *l*, is:

*λ* = ρ × *l*

To calculate the p-value of n mutations landing in a segment of length *l*, we assume a Poisson sampling process of a mutation landing in a given segment, and thus use:

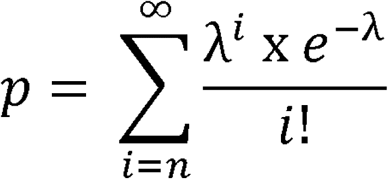

though in practice, we capped *i* arbitrarily at 50, as continually summing at *i* >50 does not appreciably affect the calculated p-value. For a given segment, we calculated the number of segments that would be expected to have p-value as good or better, as the number of tested segments multiplied by the p-value. From this, we also determined a false positive rate.

### Generation of Muller diagrams

Based on both the clonal sequencing we were able to determine which mutations were in which lineages together, and from both the clonal and population sequencing an approximate order of those mutations (though this was not exhaustive for all mutations). Using these data, we developed a lineage file format that described which mutations occurred in which lineages, and which lineages descended from one another, and used a custom Perl script that combined this information with the allele frequencies over time from the population sequencing to generate a graphical representation of the evolutionary dynamics, often referred to as a Muller diagram.

## Acknowledgements

The authors thank Matthew Herron and Eugene Kroll for their careful reading of the manuscript and their thoughtful suggestions for its improvement.

## Author Contributions

Conceived and designed the experiments: GS MK FR. Performed the experiments: MK and KS. Analyzed the data: GS KS MK JB DY FR. Contributed reagents/materials/analysis tools: GS. Wrote the paper: MK GS FR

## Supplementary Figure Legends

**Fig. S1. Population-level dynamics of mutations in 10 frequently hit genes show consistent patterns.** For each panel, elapsed generations are depicted on the x-axis and the height of each grey box represents a frequency of 100%. Cumulative frequencies for all alleles of a given gene present in the population at each time point were calculated and are represented as colored plots. (A) chemostat 1 (B) chemostat 2 (C) chemostat 3.

**Fig. S2. MalK/MalT population dynamics.** Mutant alleles of both LamB regulators, *malT* and *malK*, seldom co-occur in the same lineage, and when they do, those lineages fail to go to high frequency.

**Fig. S3**. **Cell density and residual metabolite concentrations**. Chemostat populations at steady state exhibit balanced growth, where population size remains constant and the limiting substrate, glucose is near its detection limit. As expected, populations initially produce the overflow metabolite acetate, as the founder carries a mutation that dysregulates acetyl CoA synthetase, the chief route by which *E. coli* assimilates low levels of this metabolite.

**Fig. S4. Mutations in glucosyltransferase *opgH* occur repeatedly and go to high frequency.**

**Fig. S5. Patterns of change in population genetic complexity.** Shannon’s Entropy [H], Equitability [H/Hmax] and normalized population Richness [Lineage counts] were calculated at 50 generation intervals for each of three replicate evolution experiments. Shannon’s entropy is an effective metric of population diversity as it accounts for both lineage richness (the number of lineages observed) and the relative abundance of different lineages (evenness). Lineage richness was normalized between zero and one by dividing the number of observed lineages, *S*, by the maximum *S* observed over the course of each experiment.

### Supplementary Tables

**Table S1.** Key mutations that distinguish ancestral strain JA122 from K12 (MG1655)

**Table S2.** Beneficial alleles

**Table S3**. Population allele frequencies for frequently mutated genes

**Table S4.** Identical mutations arise within and among replicate evolution experiments.

**Table S5.** Fixed alleles among replicate populations (“fixed” defined as >98% at any time point between generation 50 and 500).

### Supplementary Data Files

**File S1.** Identity and frequencies of mutations detected via population sequencing.

**File S2**. Alleles mapped onto clone phylogenies represented in main Fig. 4.

**File S3.** Muller diagrams for novel alleles arising in chemostat 1, showing details for each lineage

**File S4.** Muller diagrams for novel alleles arising in chemostat 2, showing details for each lineage

**File S5.** Muller diagrams for novel alleles arising in chemostat 3, showing details for each lineage.

