## Supplementary figures and images for "Evolutionary dynamics of *de novo* mutations and mutant lineages arising in a simple, constant environment"

### Supplementary Fig. S1

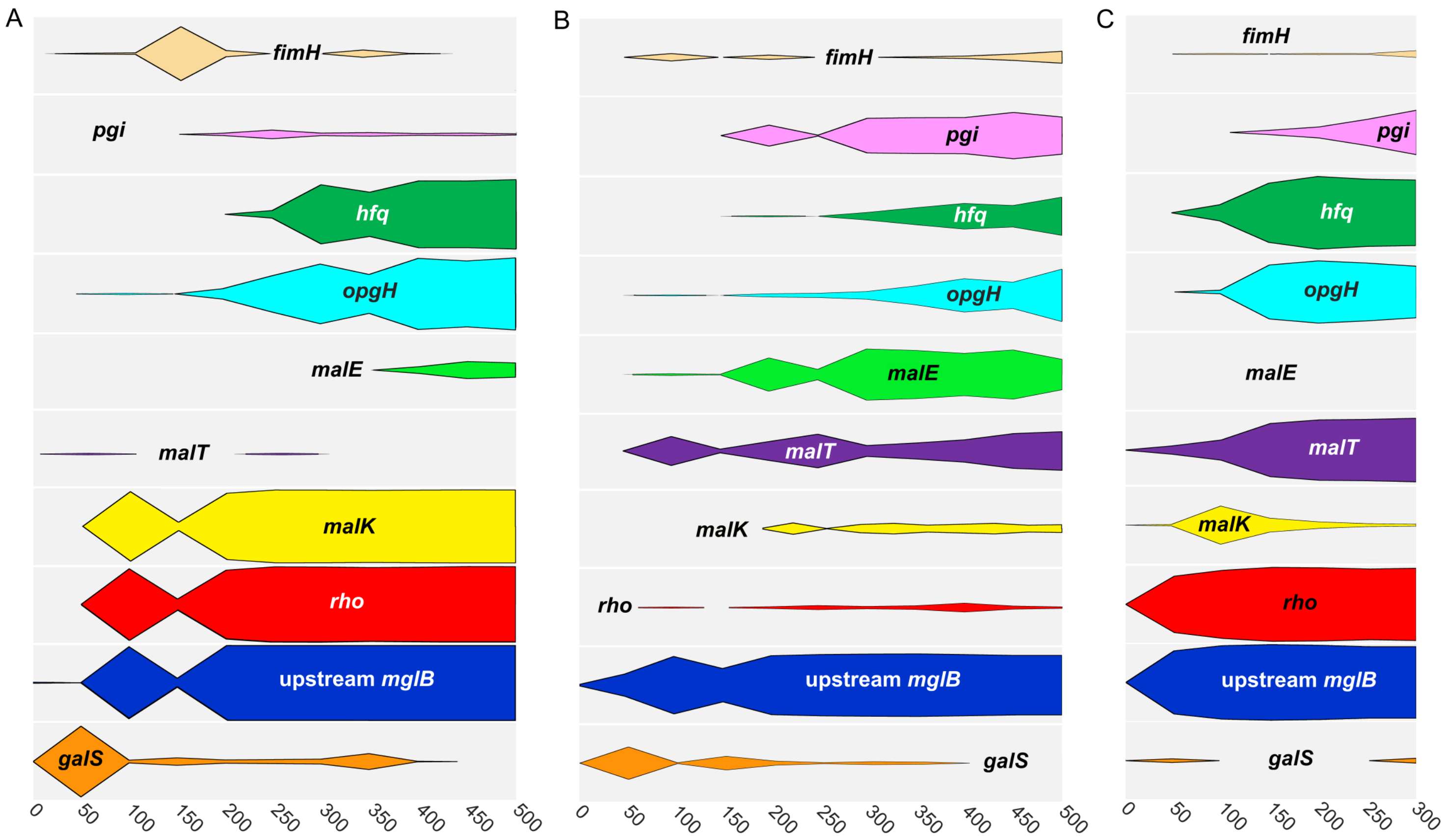

### Supplementary Fig. S2

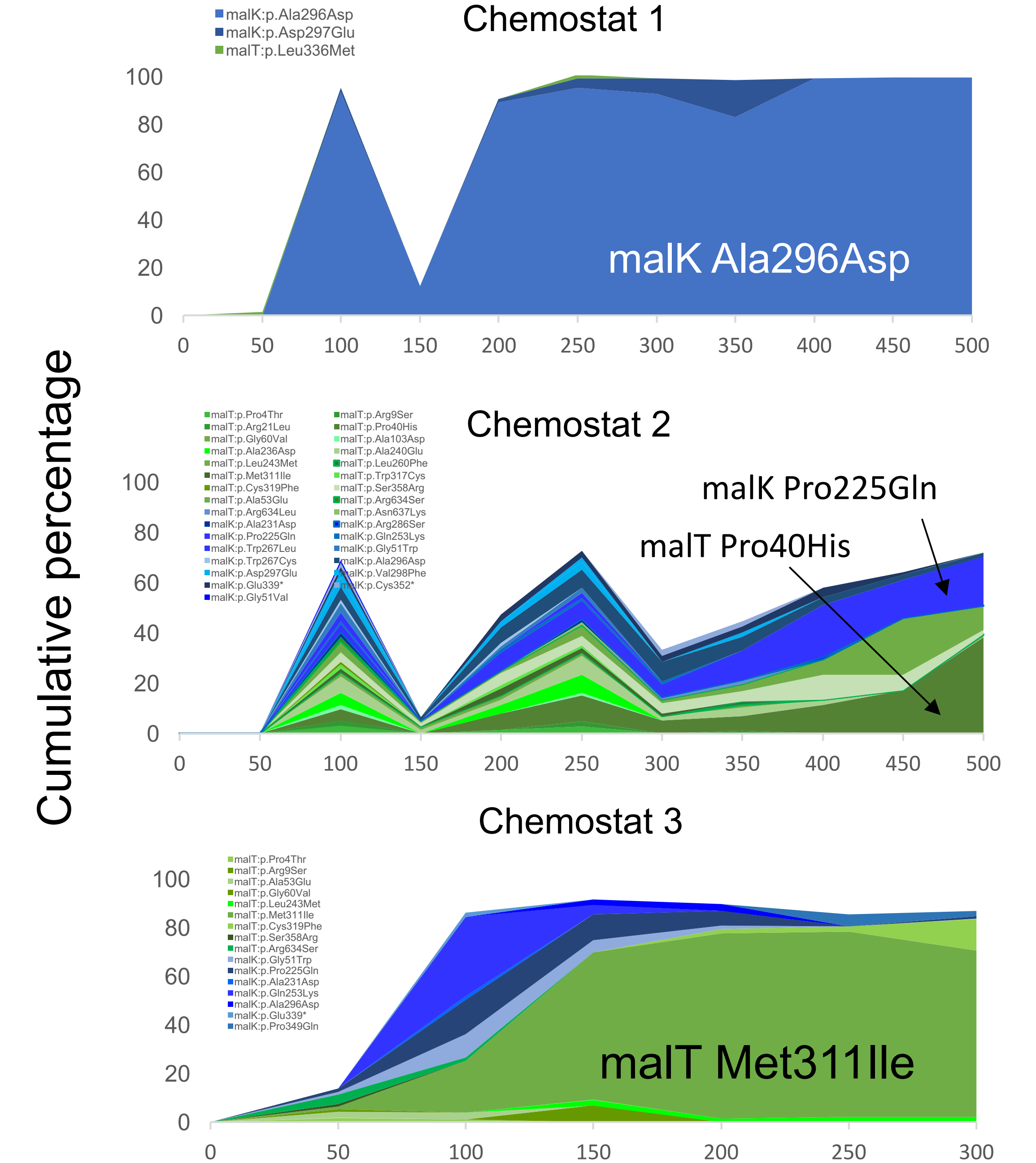

### Supplementary Fig. S3

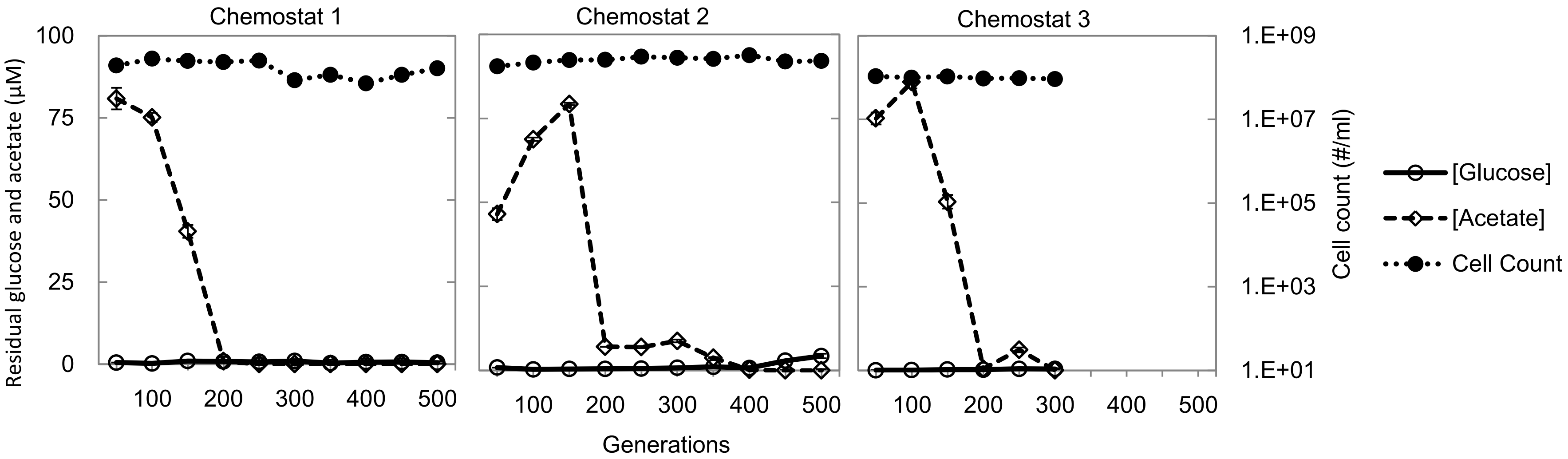

### Supplementary Fig. S4

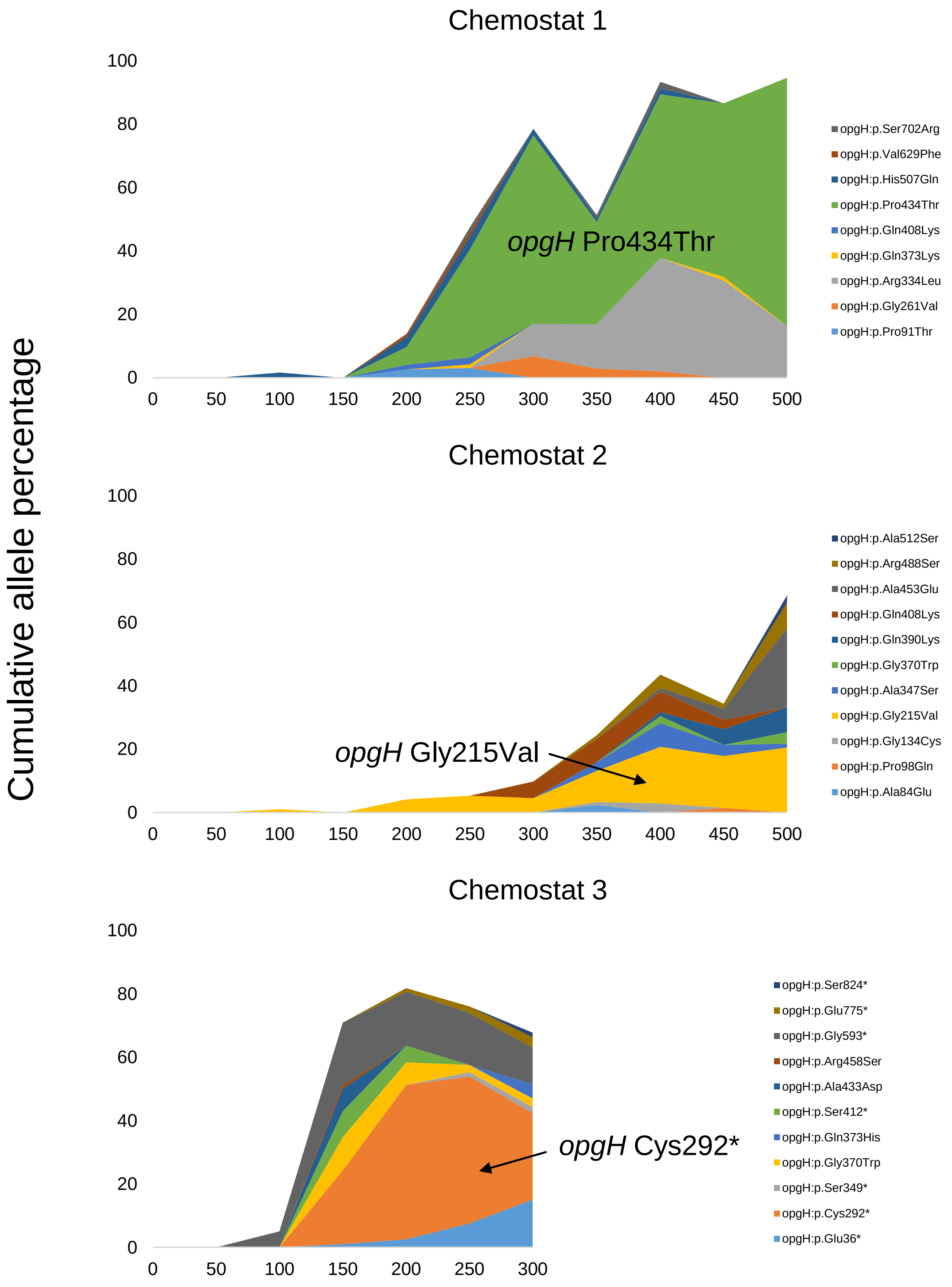

### Supplementary Fig. S5

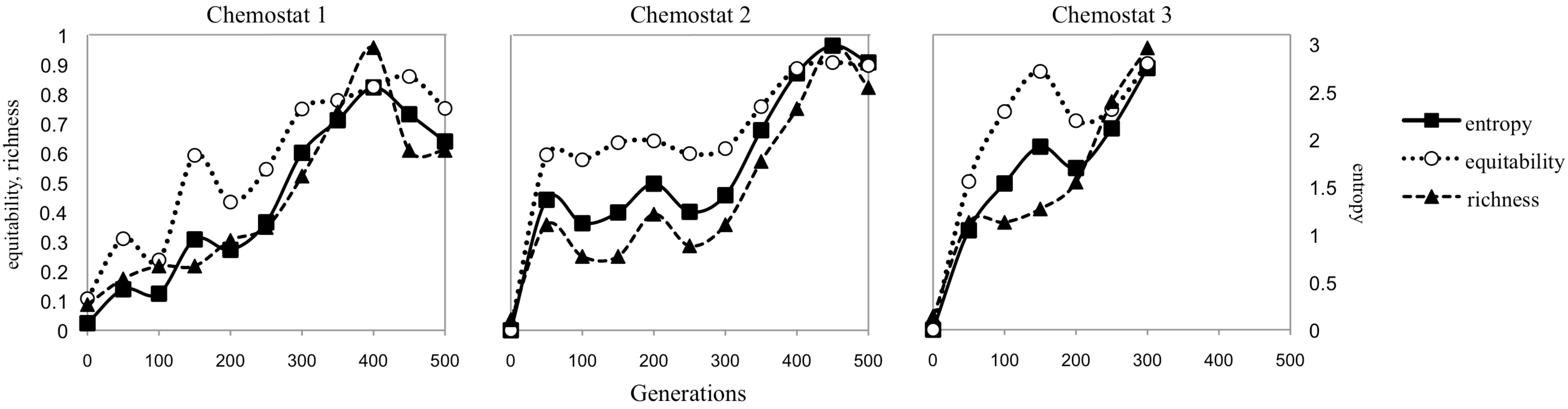
