## Supplementary Table S1 for "Evolutionary dynamics of *de novo* mutations and mutant lineages arising in a simple, constant environment"

**Table S1.** Key mutations that distinguish ancestral strain JA122 from K12 (MG1655)

| Gene | Amino acid change | Phenotype |
|---|---|---|
| *mutY (*adenine glycosylase) | Leu299* (ochre) | Increased rate of GC→TA transversions |
| *rpoD* (“exponential phase” sigma factor RpoD/$\sigma^{70}$) | Glu26* (amber) | Unknown |
| *rpoS* (“stationary phase” sigma factor RpoS/$\sigma^{38}$) | Gln33* | Largely unknown but involves suppression of “stationary-phase” gene transcription |
| *glnX* amber suppressor; reported in parent of JA122, RH101 | N/A | Allows translational read-through of amber, opal and ochre nonsense codons |
| *fimH* (fimbrial adhesion) | Pro33His | unknown |
| *fimE* (recombinase, catalyzes *fim* operon switch inversion in the on-to-off direction) | IS1 insertion +8bp target site duplication | Unknown but recombinase likely inactive |
| *lsrR* (transcriptional repressor related to quorum sensing, biofilm formation, sRNA production) | A251S | Unknown; deletion upregulates sRNAs DsrA and DicF among others |
| *acs* (acetyl-CoA synthetase (AMP-forming) A→T, position -93 from START in CRP binding site | N/A | Dysregulation of the *acs-yjcG* operon, resulting in diminished capacity to scavenge limiting acetate |
