## Supplementary Table S2 for "Evolutionary dynamics of *de novo* mutations and mutant lineages arising in a simple, constant environment"

### **Table S2** Beneficial alleles

| Rank | Gene | Expected | P-Value | Observed | Length | -LOG10 I | NUM_EX | FDR |
|---|---|---|---|---|---|---|---|---|
| 1 | galS*** | 0.778336 | 6.55E-50 | 38 | 1041 | 49.18344 | 4.42E-47 | 4.42E-45 |
| 2 | hfq****** | 0.231034 | 6.91E-40 | 24 | 309 | 39.16081 | 4.66E-37 | 2.33E-35 |
| 3 | pgi****** | 1.233675 | 4.54E-38 | 35 | 1650 | 37.34286 | 3.07E-35 | 1.02E-33 |
| 4 | opgH** | 1.902102 | 8.74E-27 | 31 | 2544 | 26.05829 | 5.90E-24 | 1.48E-22 |
| 5 | malT**** | 2.023226 | 8.10E-25 | 30 | 2706 | 24.09176 | 5.46E-22 | 1.09E-20 |
| 6 | malK**** | 0.834413 | 7.47E-24 | 22 | 1116 | 23.12675 | 5.04E-21 | 8.40E-20 |
| 7 | upstream n | 0.209351 | 2.91E-09 | 7 | 280 | 8.535746 | 1.97E-06 | 2.81E-05 |
| 8 | rho** | 0.942079 | 5.49E-09 | 11 | 1260 | 8.260065 | 3.71E-06 | 4.64E-05 |
| 9 | upstream d | 0.082993 | 3.06E-08 | 5 | 111 | 7.513989 | 2.07E-05 | 0.00023 |
| 10 | fimH*** | 0.675156 | 4.38E-08 | 9 | 903 | 7.358184 | 2.96E-05 | 0.000296 |
| 11 | rpoS* | 0.742448 | 9.71E-08 | 9 | 993 | 7.012954 | 6.55E-05 | 0.000596 |
| 12 | upstream r | 0.124115 | 2.21E-07 | 5 | 166 | 6.65491 | 0.000149 | 0.001245 |
| 13 | gatZ | 0.944322 | 7.06E-07 | 9 | 1263 | 6.151132 | 0.000477 | 0.003666 |
| 14 | pfkA* | 0.720017 | 9.47E-07 | 8 | 963 | 6.023582 | 0.000639 | 0.004567 |
| 15 | rpoA* | 0.740205 | 1.27E-05 | 7 | 990 | 4.89675 | 0.008562 | 0.057077 |
| 16 | lptD* | 1.76079 | 1.61E-05 | 10 | 2355 | 4.792867 | 0.010875 | 0.06797 |
| 17 | proQ | 0.522629 | 1.81E-05 | 6 | 699 | 4.741786 | 0.012233 | 0.071956 |
| 18 | lptG | 0.809739 | 2.24E-05 | 7 | 1083 | 4.649919 | 0.015114 | 0.083968 |
| 19 | downstrear | 0.056824 | 2.93E-05 | 3 | 76 | 4.533043 | 0.019782 | 0.104114 |
| 20 | yciM* | 0.874787 | 3.64E-05 | 7 | 1170 | 4.439428 | 0.02454 | 0.1227 |
| 21 | downstrear | 0.064301 | 4.22E-05 | 3 | 86 | 4.374416 | 0.028503 | 0.135728 |
| 22 | slt* | 1.449007 | 0.000134 | 8 | 1938 | 3.871502 | 0.090741 | 0.412457 |
| 23 | gabD* | 1.083391 | 0.000136 | 7 | 1449 | 3.867359 | 0.09161 | 0.398306 |
| 24 | wzzE | 0.782823 | 0.000164 | 6 | 1047 | 3.784636 | 0.110833 | 0.461802 |
| 25 | yobF* | 0.107666 | 0.000192 | 3 | 144 | 3.716888 | 0.129544 | 0.518174 |
| 26 | opgG* | 1.148439 | 0.000193 | 7 | 1536 | 3.714401 | 0.130287 | 0.501105 |
| 27 | fliH | 0.513657 | 0.000195 | 5 | 687 | 3.710543 | 0.13145 | 0.486852 |
| 28 | ompR | 0.538331 | 0.000241 | 5 | 720 | 3.617477 | 0.162865 | 0.581659 |
| 29 | malE | 0.890489 | 0.000325 | 6 | 1191 | 3.488285 | 0.21929 | 0.756173 |
| 30 | ybaL* | 1.253862 | 0.000326 | 7 | 1677 | 3.486737 | 0.220073 | 0.733576 |
| 31 | upstream a | 0.356644 | 0.000508 | 4 | 477 | 3.29443 | 0.342668 | 1.10538 |
| 32 | yiaO | 0.737962 | 0.000992 | 5 | 987 | 3.003644 | 0.669359 | 2.091748 |
| 33 | lptC | 0.430665 | 0.001018 | 4 | 576 | 2.992182 | 0.687261 | 2.082609 |
| 34 | fliG* | 0.744691 | 0.001032 | 5 | 996 | 2.986323 | 0.696595 | 2.04881 |
| 35 | lpxD | 0.767121 | 0.001175 | 5 | 1026 | 2.929845 | 0.793339 | 2.266681 |
| 36 | valY tRNA | 0.056824 | 0.001555 | 2 | 76 | 2.808383 | 1.049352 | 2.914866 |
| 37 | fliP | 0.551789 | 0.002495 | 4 | 738 | 2.603011 | 1.683809 | 4.550836 |
| 38 | hyaB | 1.341341 | 0.0026 | 6 | 1794 | 2.584989 | 1.755154 | 4.618827 |
| 39 | glpR | 0.56749 | 0.002757 | 4 | 759 | 2.559618 | 1.860742 | 4.771133 |
| 40 | upstream n | 0.085983 | 0.003491 | 2 | 115 | 2.457006 | 2.356663 | 5.891658 |
| 41 | ydgI* | 1.034044 | 0.004209 | 5 | 1383 | 2.375839 | 2.840958 | 6.929167 |
| 42 | rbsB | 0.666184 | 0.004846 | 4 | 891 | 2.314621 | 3.27102 | 7.788143 |
| 43 | ytfR | 1.123765 | 0.005933 | 5 | 1503 | 2.226713 | 4.004888 | 9.313694 |
| 44 | yphE | 1.130494 | 0.00608 | 5 | 1512 | 2.216113 | 4.103845 | 9.32692 |
| 45 | lpxM | 0.726746 | 0.006547 | 4 | 972 | 2.183989 | 4.418901 | 9.819781 |
| 46 | ydcT | 0.758149 | 0.007566 | 4 | 1014 | 2.121125 | 5.107146 | 11.10249 |
| 47 | phnD | 0.760392 | 0.007643 | 4 | 1017 | 2.116752 | 5.15884 | 10.97625 |
| 48 | dppA | 1.202272 | 0.007805 | 5 | 1608 | 2.107623 | 5.26842 | 10.97588 |
| 49 | pgsA | 0.410477 | 0.0085 | 3 | 549 | 2.070582 | 5.737489 | 11.70916 |

| | | | | | | | | |
|---|---|---|---|---|---|---|---|---|
| 50 | sgrR | 1.238161 | 0.008784 | 5 | 1656 | 2.056323 | 5.928996 | 11.85799 |
| 51 | lptA | 0.417206 | 0.008881 | 3 | 558 | 2.051541 | 5.994632 | 11.75418 |
| 52 | recF | 0.80301 | 0.009196 | 4 | 1074 | 2.036413 | 6.207128 | 11.93679 |
| 53 | yohD | 0.432908 | 0.009808 | 3 | 579 | 2.00841 | 6.620548 | 12.4916 |
| 54 | downstrear | 0.15477 | 0.01081 | 2 | 207 | 1.966176 | 7.296728 | 13.51246 |
| 55 | cspE | 0.157013 | 0.011109 | 2 | 210 | 1.954319 | 7.498686 | 13.63397 |
| 56 | flhB | 0.859086 | 0.011533 | 4 | 1149 | 1.938054 | 7.784847 | 13.90151 |
| 57 | nagA | 0.859086 | 0.011533 | 4 | 1149 | 1.938054 | 7.784847 | 13.65763 |
| 58 | dgoD | 0.859086 | 0.011533 | 4 | 1149 | 1.938054 | 7.784847 | 13.42215 |
| 59 | yqgC | 0.161499 | 0.011718 | 2 | 216 | 1.931131 | 7.90993 | 13.40666 |
| 60 | ydhP | 0.874787 | 0.01225 | 4 | 1170 | 1.911876 | 8.268523 | 13.78087 |
| 61 | yhdV | 0.165985 | 0.012342 | 2 | 222 | 1.908614 | 8.330866 | 13.65716 |
| 62 | sspA | 0.477768 | 0.012758 | 3 | 639 | 1.89421 | 8.611802 | 13.89 |
| 63 | emrD | 0.886003 | 0.012779 | 4 | 1185 | 1.893519 | 8.625505 | 13.69128 |
| 64 | yfaY | 0.899461 | 0.013432 | 4 | 1203 | 1.871856 | 9.066671 | 14.16667 |
| 65 | secD | 1.381715 | 0.013544 | 5 | 1848 | 1.868261 | 9.142042 | 14.06468 |
| 66 | asmA | 1.386202 | 0.013716 | 5 | 1854 | 1.862785 | 9.258029 | 14.02732 |
| 67 | ycjU | 0.49347 | 0.013897 | 3 | 660 | 1.857066 | 9.380764 | 14.00114 |
| 68 | ydhI | 0.177201 | 0.013963 | 2 | 237 | 1.855023 | 9.424986 | 13.86027 |
| 69 | upstream y | 0.189163 | 0.015787 | 2 | 253 | 1.801689 | 10.65651 | 15.44422 |
| 70 | tolA | 0.946565 | 0.015885 | 4 | 1266 | 1.799005 | 10.72258 | 15.31797 |
| 71 | efeB | 0.951051 | 0.016133 | 4 | 1272 | 1.792296 | 10.88949 | 15.33731 |
| 72 | upstream d | 0.192902 | 0.016377 | 2 | 258 | 1.785755 | 11.05474 | 15.3538 |
| 73 | ndh* | 0.975724 | 0.017536 | 4 | 1305 | 1.756076 | 11.83661 | 16.21454 |
| 74 | lptB* | 0.542817 | 0.017843 | 3 | 726 | 1.748526 | 12.04418 | 16.27592 |
| 75 | menE | 1.013856 | 0.01985 | 4 | 1356 | 1.70225 | 13.39843 | 17.86457 |
| 76 | upstream p | 0.214585 | 0.01998 | 2 | 287 | 1.699401 | 13.48662 | 17.74555 |
| 77 | yedV | 1.016099 | 0.019991 | 4 | 1359 | 1.699161 | 13.49405 | 17.52474 |
| 78 | aroD | 0.56749 | 0.020026 | 3 | 759 | 1.698414 | 13.51729 | 17.32986 |
| 79 | hyi | 0.580949 | 0.021275 | 3 | 777 | 1.672128 | 14.3607 | 18.1781 |
| 80 | flgG | 0.585435 | 0.021701 | 3 | 783 | 1.663523 | 14.6481 | 18.31012 |
| 81 | fhuF | 0.589921 | 0.022131 | 3 | 789 | 1.654993 | 14.93863 | 18.44275 |
| 82 | mdtD | 1.058717 | 0.022802 | 4 | 1416 | 1.642034 | 15.3911 | 18.76963 |
| 83 | upstream p | 0.231034 | 0.022913 | 2 | 309 | 1.63992 | 15.46619 | 18.63397 |
| 84 | asr | 0.231034 | 0.022913 | 2 | 309 | 1.63992 | 15.46619 | 18.41213 |
| 85 | ptrA | 2.160052 | 0.023057 | 6 | 2889 | 1.637201 | 15.56333 | 18.3098 |
| 86 | qseE | 1.067689 | 0.023422 | 4 | 1428 | 1.630371 | 15.81004 | 18.38377 |
| 87 | hsrA | 1.067689 | 0.023422 | 4 | 1428 | 1.630371 | 15.81004 | 18.17246 |
| 88 | yggL | 0.244492 | 0.025435 | 2 | 327 | 1.59456 | 17.16895 | 19.51017 |
| 89 | yjfO | 0.246735 | 0.025866 | 2 | 330 | 1.587264 | 17.45984 | 19.6178 |
| 90 | wzxC | 1.105821 | 0.026174 | 4 | 1479 | 1.582133 | 17.66732 | 19.63036 |
| 91 | gspE | 1.108064 | 0.026341 | 4 | 1482 | 1.579361 | 17.78046 | 19.53897 |
| 92 | gatY | 0.639268 | 0.027172 | 3 | 855 | 1.56588 | 18.34104 | 19.93592 |
| 93 | cdsA | 0.641511 | 0.027414 | 3 | 858 | 1.562022 | 18.50468 | 19.89751 |
| 94 | relA | 1.671068 | 0.027811 | 5 | 2235 | 1.555778 | 18.77267 | 19.97092 |
| 95 | cysW* | 0.654969 | 0.028893 | 3 | 876 | 1.539206 | 19.50285 | 20.52932 |
| 96 | upstream y | 0.268418 | 0.030183 | 2 | 359 | 1.520243 | 20.37326 | 21.22214 |
| 97 | yfgD | 0.269165 | 0.030336 | 2 | 360 | 1.518039 | 20.47694 | 21.11025 |
| 98 | ygfK | 2.317065 | 0.030903 | 6 | 3099 | 1.510005 | 20.85924 | 21.28494 |
| 99 | putA | 2.963062 | 0.031681 | 7 | 3963 | 1.499205 | 21.38451 | 21.60051 |
| 100 | lpxC | 0.686372 | 0.032505 | 3 | 918 | 1.488056 | 21.94055 | 21.94055 |
| 101 | adrB | 1.195543 | 0.033384 | 4 | 1599 | 1.476467 | 22.53394 | 22.31084 |

| | | | | | | | | |
|---|---|---|---|---|---|---|---|---|
| 102 | rihA | 0.69983 | 0.034121 | 3 | 936 | 1.46698 | 23.0316 | 22.58 |
| 103 | flgJ | 0.704316 | 0.034669 | 3 | 942 | 1.460061 | 23.40143 | 22.71984 |
| 104 | aidB | 1.21573 | 0.03515 | 4 | 1626 | 1.45408 | 23.72596 | 22.81342 |
| 105 | yqeB | 1.21573 | 0.03515 | 4 | 1626 | 1.45408 | 23.72596 | 22.59615 |
| 106 | downstrear | 0.711045 | 0.035499 | 3 | 951 | 1.449782 | 23.96192 | 22.60558 |
| 107 | ygiT | 0.296082 | 0.03607 | 2 | 396 | 1.44286 | 24.34693 | 22.75414 |
| 108 | fumA | 1.231431 | 0.03656 | 4 | 1647 | 1.436996 | 24.67786 | 22.84987 |
| 109 | upstream n | 0.299073 | 0.03673 | 2 | 400 | 1.434974 | 24.79304 | 22.74591 |
| 110 | hslR | 0.300568 | 0.037063 | 2 | 402 | 1.431064 | 25.01727 | 22.74297 |
| 111 | rnk | 0.307297 | 0.038572 | 2 | 411 | 1.41373 | 26.03596 | 23.45582 |
| 112 | dusA | 0.742448 | 0.039508 | 3 | 993 | 1.40331 | 26.66818 | 23.81088 |
| 113 | fruR | 0.75142 | 0.040694 | 3 | 1005 | 1.390466 | 27.46866 | 24.30854 |
| 114 | rhsC | 3.135776 | 0.040824 | 7 | 4194 | 1.389082 | 27.55633 | 24.17222 |
| 115 | upstream y | 0.321503 | 0.041833 | 2 | 430 | 1.37848 | 28.23734 | 24.5542 |
| 116 | upstream n | 0.325241 | 0.042708 | 2 | 435 | 1.369491 | 28.82788 | 24.85162 |
| 117 | tdh | 0.767121 | 0.042812 | 3 | 1026 | 1.36843 | 28.89838 | 24.69947 |
| 118 | mdtN | 0.771607 | 0.043428 | 3 | 1032 | 1.362235 | 29.3136 | 24.84203 |
| 119 | yagN* | 0.329728 | 0.043767 | 2 | 441 | 1.358855 | 29.5426 | 24.82571 |
| 120 | yhbP | 0.331971 | 0.0443 | 2 | 444 | 1.353597 | 29.90242 | 24.91868 |
| 121 | tas | 0.778336 | 0.044359 | 3 | 1041 | 1.353023 | 29.942 | 24.74545 |
| 122 | yejM | 1.316667 | 0.044777 | 4 | 1761 | 1.348947 | 30.22436 | 24.77406 |
| 123 | fbaB | 0.787309 | 0.045615 | 3 | 1053 | 1.340891 | 30.79024 | 25.03272 |
| 124 | phnG | 0.3387 | 0.045913 | 2 | 453 | 1.33806 | 30.99161 | 24.99323 |
| 125 | matP | 0.3387 | 0.045913 | 2 | 453 | 1.33806 | 30.99161 | 24.79329 |
| 126 | yfjX | 0.343186 | 0.047001 | 2 | 459 | 1.327892 | 31.72576 | 25.17917 |
| 127 | pyrI | 0.345429 | 0.047548 | 2 | 462 | 1.322864 | 32.09521 | 25.27182 |
| 128 | narX | 1.343584 | 0.047569 | 4 | 1797 | 1.32268 | 32.10879 | 25.08499 |
| 129 | ulaC | 0.347672 | 0.048098 | 2 | 465 | 1.317872 | 32.46623 | 25.16762 |
| 130 | yiaL | 0.349915 | 0.04865 | 2 | 468 | 1.312916 | 32.83882 | 25.26063 |
| 131 | ybiC | 0.811982 | 0.049161 | 3 | 1086 | 1.30838 | 33.1836 | 25.331 |
| 132 | recD | 1.366014 | 0.049967 | 4 | 1827 | 1.301317 | 33.72774 | 25.55132 |
| 133 | anmK | 0.829926 | 0.051822 | 3 | 1110 | 1.285489 | 34.97957 | 26.30043 |
| 134 | hypD | 0.838899 | 0.053178 | 3 | 1122 | 1.274272 | 35.89485 | 26.7872 |
| 135 | yhhJ | 0.841142 | 0.053519 | 3 | 1125 | 1.27149 | 36.12545 | 26.7596 |
| 136 | yfdN | 0.370102 | 0.05372 | 2 | 495 | 1.269863 | 36.26109 | 26.66256 |
| 137 | yebR | 0.372345 | 0.054295 | 2 | 498 | 1.265244 | 36.64884 | 26.75098 |
| 138 | yafM | 0.372345 | 0.054295 | 2 | 498 | 1.265244 | 36.64884 | 26.55713 |
| 139 | recX | 0.374588 | 0.054871 | 2 | 501 | 1.260655 | 37.03806 | 26.64609 |
| 140 | phnM | 0.850114 | 0.054896 | 3 | 1137 | 1.260457 | 37.05501 | 26.46787 |
| 141 | downstrear | 0.375336 | 0.055064 | 2 | 502 | 1.259133 | 37.16813 | 26.36038 |
| 142 | ydeR | 0.376831 | 0.05545 | 2 | 504 | 1.256098 | 37.42875 | 26.35827 |
| 143 | mug | 0.379075 | 0.056031 | 2 | 507 | 1.251572 | 37.82088 | 26.44817 |
| 144 | yheS* | 1.431062 | 0.05729 | 4 | 1914 | 1.241918 | 38.67104 | 26.85489 |
| 145 | ygbK | 0.872544 | 0.058412 | 3 | 1167 | 1.233494 | 39.42841 | 27.19201 |
| 146 | yhhZ | 0.881517 | 0.059848 | 3 | 1179 | 1.222951 | 40.39738 | 27.66944 |
| 147 | tsgA | 0.88376 | 0.060209 | 3 | 1182 | 1.220336 | 40.64136 | 27.64719 |
| 148 | tufA | 0.886003 | 0.060572 | 3 | 1185 | 1.217729 | 40.88603 | 27.6257 |
| 149 | ybjK | 0.401505 | 0.061956 | 2 | 537 | 1.207917 | 41.82024 | 28.06728 |
| 150 | yagF | 1.471437 | 0.062109 | 4 | 1968 | 1.206846 | 41.92349 | 27.94899 |
| 151 | dnaT | 0.403748 | 0.06256 | 2 | 540 | 1.203705 | 42.2278 | 27.96543 |
| 152 | dacC | 0.899461 | 0.062768 | 3 | 1203 | 1.20226 | 42.36853 | 27.87403 |
| 153 | aegA | 1.480409 | 0.063208 | 4 | 1980 | 1.19923 | 42.66524 | 27.88578 |

| | | | | | | | | |
|---|---|---|---|---|---|---|---|---|
| 154 | mdtH | 0.903947 | 0.063508 | 3 | 1209 | 1.197169 | 42.86818 | 27.83648 |
| 155 | hyfH | 0.408234 | 0.063773 | 2 | 546 | 1.195361 | 43.04695 | 27.77223 |
| 156 | dgsA | 0.912919 | 0.065001 | 3 | 1221 | 1.18708 | 43.87564 | 28.12541 |
| 157 | phnN | 0.417206 | 0.066224 | 2 | 558 | 1.178984 | 44.70124 | 28.47213 |
| 158 | hyfB | 1.509569 | 0.066849 | 4 | 2019 | 1.174904 | 45.12321 | 28.559 |
| 159 | cca* | 0.926377 | 0.06727 | 3 | 1239 | 1.17218 | 45.4071 | 28.55792 |
| 160 | ytfN | 2.826236 | 0.067422 | 6 | 3780 | 1.171198 | 45.50988 | 28.44367 |
| 161 | spr | 0.423935 | 0.068082 | 2 | 567 | 1.166965 | 45.95567 | 28.54389 |
| 162 | yfeO | 0.939836 | 0.069574 | 3 | 1257 | 1.157551 | 46.96263 | 28.98928 |
| 163 | yjeH* | 0.939836 | 0.069574 | 3 | 1257 | 1.157551 | 46.96263 | 28.81143 |
| 164 | nfuA | 0.430665 | 0.069958 | 2 | 576 | 1.155163 | 47.22163 | 28.79367 |
| 165 | rpoE | 0.430665 | 0.069958 | 2 | 576 | 1.155163 | 47.22163 | 28.61917 |
| 166 | upstream n | 0.43216 | 0.070377 | 2 | 578 | 1.152569 | 47.50449 | 28.61716 |
| 167 | lhgO | 0.948808 | 0.07113 | 3 | 1269 | 1.147946 | 48.01289 | 28.75023 |
| 168 | gabT | 0.95778 | 0.072702 | 3 | 1281 | 1.138456 | 49.07364 | 29.2105 |
| 169 | yeaH | 0.960023 | 0.073097 | 3 | 1284 | 1.136101 | 49.34046 | 29.19554 |
| 170 | yajL | 0.44188 | 0.073121 | 2 | 591 | 1.135959 | 49.35659 | 29.03329 |
| 171 | pinR | 0.44188 | 0.073121 | 2 | 591 | 1.135959 | 49.35659 | 28.8635 |
| 172 | ydiS | 0.964509 | 0.07389 | 3 | 1290 | 1.131412 | 49.87604 | 28.9977 |
| 173 | ynbD | 0.966752 | 0.074289 | 3 | 1293 | 1.129078 | 50.1448 | 28.98543 |
| 174 | upstream p | 0.447114 | 0.074612 | 2 | 598 | 1.127189 | 50.36339 | 28.94448 |
| 175 | upstream h | 0.452347 | 0.076114 | 2 | 605 | 1.118537 | 51.37673 | 29.35813 |
| 176 | yrfF | 1.597048 | 0.078407 | 4 | 2136 | 1.105645 | 52.9248 | 30.07091 |
| 177 | rhtB | 0.46431 | 0.079581 | 2 | 621 | 1.099192 | 53.71696 | 30.34856 |
| 178 | fdhF | 1.60602 | 0.079645 | 4 | 2148 | 1.098839 | 53.76063 | 30.2026 |
| 179 | yaiV | 0.466553 | 0.080236 | 2 | 624 | 1.09563 | 54.15941 | 30.25665 |
| 180 | gudX | 1.002641 | 0.080788 | 3 | 1341 | 1.092654 | 54.53177 | 30.29543 |
| 181 | ydhY | 0.468796 | 0.080893 | 2 | 627 | 1.092087 | 54.60299 | 30.1674 |
| 182 | yagK | 0.468796 | 0.080893 | 2 | 627 | 1.092087 | 54.60299 | 30.00164 |
| 183 | fdnG | 2.278933 | 0.081307 | 5 | 3048 | 1.089871 | 54.88235 | 29.99035 |
| 184 | nudF | 0.471039 | 0.081552 | 2 | 630 | 1.088565 | 55.04771 | 29.91723 |
| 185 | upstream y | 0.471787 | 0.081772 | 2 | 631 | 1.087395 | 55.19619 | 29.83578 |
| 186 | dhaL | 0.473282 | 0.082213 | 2 | 633 | 1.085061 | 55.49354 | 29.83524 |
| 187 | ftsK | 2.983249 | 0.082239 | 6 | 3990 | 1.084924 | 55.51105 | 29.68505 |
| 188 | yfdX | 0.475525 | 0.082875 | 2 | 636 | 1.081578 | 55.94049 | 29.75558 |
| 189 | argO | 0.475525 | 0.082875 | 2 | 636 | 1.081578 | 55.94049 | 29.59814 |
| 190 | hcaE | 1.018342 | 0.083706 | 3 | 1362 | 1.077243 | 56.50156 | 29.73766 |
| 191 | eda | 0.480012 | 0.084204 | 2 | 642 | 1.074667 | 56.8377 | 29.75796 |
| 192 | adk | 0.482255 | 0.084871 | 2 | 645 | 1.07124 | 57.28795 | 29.83748 |
| 193 | cheZ | 0.482255 | 0.084871 | 2 | 645 | 1.07124 | 57.28795 | 29.68288 |
| 194 | aroP | 1.027314 | 0.085394 | 3 | 1374 | 1.068575 | 57.64067 | 29.71169 |
| 195 | fabR | 0.484498 | 0.08554 | 2 | 648 | 1.067832 | 57.73929 | 29.60989 |
| 196 | rcsB | 0.486741 | 0.08621 | 2 | 651 | 1.064443 | 58.1917 | 29.68964 |
| 197 | elbB | 0.488984 | 0.086882 | 2 | 654 | 1.061071 | 58.64519 | 29.76913 |
| 198 | hyuA | 1.036287 | 0.087096 | 3 | 1386 | 1.060004 | 58.7895 | 29.69167 |
| 199 | sad | 1.03853 | 0.087523 | 3 | 1389 | 1.057876 | 59.07822 | 29.68755 |
| 200 | upstream y | 0.491227 | 0.087555 | 2 | 657 | 1.057718 | 59.09974 | 29.54987 |
| 201 | ybiO | 1.664339 | 0.087928 | 4 | 2226 | 1.055872 | 59.35157 | 29.52814 |
| 202 | xanP | 1.040773 | 0.087952 | 3 | 1392 | 1.055755 | 59.36754 | 29.38987 |
| 203 | pcnB | 1.045259 | 0.088812 | 3 | 1398 | 1.051529 | 59.94796 | 29.53102 |
| 204 | yedK | 0.500199 | 0.090264 | 2 | 669 | 1.044484 | 60.92843 | 29.86688 |
| 205 | upstream y | 0.500947 | 0.090491 | 2 | 670 | 1.043394 | 61.08158 | 29.79589 |

| | | | | | | | | |
|---|---|---|---|---|---|---|---|---|
| 206 | ybiH | 0.502442 | 0.090945 | 2 | 672 | 1.041219 | 61.3882 | 29.8001 |
| 207 | yiiM | 0.504685 | 0.091628 | 2 | 675 | 1.037971 | 61.84899 | 29.87874 |
| 208 | treB | 1.063203 | 0.092286 | 3 | 1422 | 1.034862 | 62.29338 | 29.94874 |
| 209 | lpd | 1.065446 | 0.092725 | 3 | 1425 | 1.032804 | 62.5892 | 29.94699 |
| 210 | narV | 0.509171 | 0.092998 | 2 | 681 | 1.031527 | 62.7736 | 29.89219 |
| 211 | rnc | 0.509171 | 0.092998 | 2 | 681 | 1.031527 | 62.7736 | 29.75052 |
| 212 | upstream e | 1.069185 | 0.093457 | 3 | 1430 | 1.029388 | 63.08353 | 29.75638 |
| 213 | cmk | 0.511414 | 0.093685 | 2 | 684 | 1.02833 | 63.23742 | 29.68893 |
| 214 | ydiU | 1.074418 | 0.094486 | 3 | 1437 | 1.024631 | 63.7783 | 29.80294 |
| 215 | bglA | 1.076661 | 0.094929 | 3 | 1440 | 1.022601 | 64.07702 | 29.80327 |
| 216 | aldA | 1.076661 | 0.094929 | 3 | 1440 | 1.022601 | 64.07702 | 29.66529 |
| 217 | paoA | 0.5159 | 0.095064 | 2 | 690 | 1.021985 | 64.16802 | 29.57052 |
| 218 | modB | 0.5159 | 0.095064 | 2 | 690 | 1.021985 | 64.16802 | 29.43487 |
| 219 | nanE* | 0.5159 | 0.095064 | 2 | 690 | 1.021985 | 64.16802 | 29.30047 |
| 220 | mutH | 0.5159 | 0.095064 | 2 | 690 | 1.021985 | 64.16802 | 29.16728 |
| 221 | yieP | 0.518143 | 0.095755 | 2 | 693 | 1.018837 | 64.6348 | 29.24652 |
| 222 | torR | 0.518143 | 0.095755 | 2 | 693 | 1.018837 | 64.6348 | 29.11477 |
| 223 | napG | 0.520386 | 0.096448 | 2 | 696 | 1.015706 | 65.10255 | 29.19397 |
| 224 | frvB | 1.085634 | 0.096708 | 3 | 1452 | 1.014539 | 65.27763 | 29.1418 |
| 225 | yjfJ | 0.522629 | 0.097143 | 2 | 699 | 1.01259 | 65.57127 | 29.14279 |
| 226 | yhgF | 1.736116 | 0.098666 | 4 | 2322 | 1.005832 | 66.5996 | 29.46885 |
| 227 | upstream tl | 0.527863 | 0.098768 | 2 | 706 | 1.005382 | 66.66868 | 29.36946 |
| 228 | guaB | 1.096849 | 0.09895 | 3 | 1467 | 1.004585 | 66.7912 | 29.29439 |
| 229 | upstream g | 0.530106 | 0.099468 | 2 | 709 | 1.002319 | 67.14058 | 29.31903 |
| 230 | modF | 1.101335 | 0.099853 | 3 | 1473 | 1.00064 | 67.40057 | 29.30459 |
| 231 | ispD | 0.531602 | 0.099934 | 2 | 711 | 1.000285 | 67.4557 | 29.2016 |
| 232 | ampG | 1.103578 | 0.100305 | 3 | 1476 | 0.998676 | 67.70609 | 29.18366 |
| 233 | sanA | 0.538331 | 0.102043 | 2 | 720 | 0.991218 | 68.87886 | 29.56174 |
| 234 | puuC | 1.11255 | 0.102124 | 3 | 1488 | 0.990872 | 68.93374 | 29.45886 |
| 235 | amyA | 1.11255 | 0.102124 | 3 | 1488 | 0.990872 | 68.93374 | 29.33351 |
| 236 | baeR | 0.540574 | 0.102748 | 2 | 723 | 0.988225 | 69.35509 | 29.38775 |
| 237 | yafS | 0.540574 | 0.102748 | 2 | 723 | 0.988225 | 69.35509 | 29.26375 |
| 238 | pyrH | 0.542817 | 0.103455 | 2 | 726 | 0.985248 | 69.83223 | 29.34127 |
| 239 | yfiH | 0.547303 | 0.104873 | 2 | 732 | 0.979337 | 70.7892 | 29.61891 |
| 240 | putP | 1.128251 | 0.105338 | 3 | 1509 | 0.977414 | 71.10328 | 29.62637 |
| 241 | rhsD | 3.200825 | 0.105502 | 6 | 4281 | 0.976739 | 71.21393 | 29.54935 |
| 242 | pyrF | 0.551789 | 0.106296 | 2 | 738 | 0.973484 | 71.74972 | 29.64865 |
| 243 | yjbG | 0.551789 | 0.106296 | 2 | 738 | 0.973484 | 71.74972 | 29.52663 |
| 244 | yifB | 1.137224 | 0.107193 | 3 | 1521 | 0.969836 | 72.35494 | 29.65367 |
| 245 | yacF | 0.556275 | 0.107724 | 2 | 744 | 0.967687 | 72.71376 | 29.67908 |
| 246 | wcaE | 0.558518 | 0.10844 | 2 | 747 | 0.96481 | 73.19707 | 29.75491 |
| 247 | emrB | 1.150682 | 0.109998 | 3 | 1539 | 0.958616 | 74.24844 | 30.0601 |
| 248 | agaI | 0.565247 | 0.110596 | 2 | 756 | 0.956261 | 74.65214 | 30.10167 |
| 249 | znuC | 0.565247 | 0.110596 | 2 | 756 | 0.956261 | 74.65214 | 29.98078 |
| 250 | yagI | 0.56749 | 0.111317 | 2 | 759 | 0.953439 | 75.13885 | 30.05554 |
| 251 | phnK | 0.56749 | 0.111317 | 2 | 759 | 0.953439 | 75.13885 | 29.9358 |
| 252 | yjjI | 1.159654 | 0.111883 | 3 | 1551 | 0.951234 | 75.52129 | 29.96876 |
| 253 | ydiQ | 0.571976 | 0.112763 | 2 | 765 | 0.947835 | 76.11476 | 30.08489 |
| 254 | thiG | 0.576462 | 0.114213 | 2 | 771 | 0.942283 | 77.09396 | 30.35195 |
| 255 | srlR | 0.578705 | 0.11494 | 2 | 774 | 0.939527 | 77.58477 | 30.4254 |
| 256 | uxuR | 0.578705 | 0.11494 | 2 | 774 | 0.939527 | 77.58477 | 30.30655 |
| 257 | garD | 1.175355 | 0.115213 | 3 | 1572 | 0.938499 | 77.76869 | 30.26019 |

| | | | | | | | | |
|---|---|---|---|---|---|---|---|---|
| 258 | ygfM | 0.583192 | 0.116398 | 2 | 780 | 0.934054 | 78.56879 | 30.45302 |
| 259 | ylbA | 0.587678 | 0.117861 | 2 | 786 | 0.928631 | 79.55598 | 30.71659 |
| 260 | ycjR | 0.589921 | 0.118594 | 2 | 789 | 0.925938 | 80.05074 | 30.78875 |
| 261 | nanR | 0.592164 | 0.119328 | 2 | 792 | 0.923258 | 80.54627 | 30.86064 |
| 262 | ssuC | 0.592164 | 0.119328 | 2 | 792 | 0.923258 | 80.54627 | 30.74285 |
| 263 | agaD | 0.592164 | 0.119328 | 2 | 792 | 0.923258 | 80.54627 | 30.62596 |
| 264 | otsB | 0.598893 | 0.121537 | 2 | 801 | 0.915292 | 82.03745 | 31.07479 |
| 265 | bcsC | 2.597446 | 0.122216 | 5 | 3474 | 0.912874 | 82.49547 | 31.13037 |
| 266 | suhB | 0.601136 | 0.122276 | 2 | 804 | 0.91266 | 82.53601 | 31.02858 |
| 267 | rna | 0.603379 | 0.123015 | 2 | 807 | 0.910041 | 83.03532 | 31.09937 |
| 268 | nikE | 0.603379 | 0.123015 | 2 | 807 | 0.910041 | 83.03532 | 30.98333 |
| 269 | mhpD | 0.605622 | 0.123756 | 2 | 810 | 0.907433 | 83.53536 | 31.05404 |
| 270 | yaeI | 0.607865 | 0.124498 | 2 | 813 | 0.904838 | 84.03613 | 31.12449 |
| 271 | yegX | 0.612351 | 0.125985 | 2 | 819 | 0.899681 | 85.03985 | 31.38002 |
| 272 | ybhA | 0.612351 | 0.125985 | 2 | 819 | 0.899681 | 85.03985 | 31.26465 |
| 273 | rsmA | 0.614594 | 0.12673 | 2 | 822 | 0.89712 | 85.54278 | 31.33435 |
| 274 | upstream e | 1.229936 | 0.127069 | 3 | 1645 | 0.895961 | 85.77137 | 31.30342 |
| 275 | mqo | 1.231431 | 0.127399 | 3 | 1647 | 0.894832 | 85.99465 | 31.27078 |
| 276 | gadX | 0.616837 | 0.127476 | 2 | 825 | 0.894571 | 86.04642 | 31.17624 |
| 277 | tauC | 0.61908 | 0.128223 | 2 | 828 | 0.892033 | 86.55076 | 31.24576 |
| 278 | ushA* | 1.235918 | 0.128394 | 3 | 1653 | 0.891456 | 86.66571 | 31.17472 |
| 279 | glpG | 0.621323 | 0.128972 | 2 | 831 | 0.889506 | 87.0558 | 31.2028 |
| 280 | fliF | 1.240404 | 0.129391 | 3 | 1659 | 0.888097 | 87.33864 | 31.19237 |
| 281 | nrfE | 1.240404 | 0.129391 | 3 | 1659 | 0.888097 | 87.33864 | 31.08137 |
| 282 | csgG | 0.623566 | 0.129721 | 2 | 834 | 0.88699 | 87.56153 | 31.05019 |
| 283 | prmC | 0.623566 | 0.129721 | 2 | 834 | 0.88699 | 87.56153 | 30.94047 |
| 284 | asnB | 1.24489 | 0.13039 | 3 | 1665 | 0.884755 | 88.01342 | 30.99064 |
| 285 | hdfR | 0.628052 | 0.131222 | 2 | 840 | 0.881992 | 88.57504 | 31.07896 |
| 286 | purU | 0.630296 | 0.131975 | 2 | 843 | 0.87951 | 89.08281 | 31.14783 |
| 287 | ligB | 1.258348 | 0.133406 | 3 | 1683 | 0.874826 | 90.04874 | 31.37587 |
| 288 | panC | 0.637025 | 0.134237 | 2 | 852 | 0.872127 | 90.6101 | 31.46184 |
| 289 | upstream y | 0.637772 | 0.134489 | 2 | 853 | 0.871313 | 90.78017 | 31.41182 |
| 290 | fruA | 1.265077 | 0.134922 | 3 | 1692 | 0.869917 | 91.0725 | 31.40431 |
| 291 | ampE | 0.639268 | 0.134993 | 2 | 855 | 0.869688 | 91.12051 | 31.31289 |
| 292 | ybbN | 0.639268 | 0.134993 | 2 | 855 | 0.869688 | 91.12051 | 31.20565 |
| 293 | ydaU | 0.641511 | 0.13575 | 2 | 858 | 0.867259 | 91.63157 | 31.27357 |
| 294 | ybaE | 1.271806 | 0.136445 | 3 | 1701 | 0.865043 | 92.10027 | 31.32662 |
| 295 | yfjP | 0.645997 | 0.137268 | 2 | 864 | 0.862432 | 92.65559 | 31.40868 |
| 296 | upstream p | 0.645997 | 0.137268 | 2 | 864 | 0.862432 | 92.65559 | 31.30257 |
| 297 | yebT | 1.969393 | 0.137396 | 4 | 2634 | 0.862025 | 92.74251 | 31.22643 |
| 298 | upstream z | 1.279283 | 0.138144 | 3 | 1711 | 0.859669 | 93.24689 | 31.2909 |
| 299 | rluF | 0.652726 | 0.13955 | 2 | 873 | 0.85527 | 94.19635 | 31.5038 |
| 300 | cvrA | 1.298723 | 0.142593 | 3 | 1737 | 0.8459 | 96.2506 | 32.08353 |
| 301 | yghA | 0.661698 | 0.142606 | 2 | 885 | 0.845861 | 96.25926 | 31.97982 |
| 302 | adhE | 2.000796 | 0.14302 | 4 | 2676 | 0.844603 | 96.53861 | 31.96643 |
| 303 | arnD | 0.666184 | 0.14414 | 2 | 891 | 0.841216 | 97.29429 | 32.11033 |
| 304 | malG | 0.666184 | 0.14414 | 2 | 891 | 0.841216 | 97.29429 | 32.0047 |
| 305 | argP | 0.668427 | 0.144908 | 2 | 894 | 0.838909 | 97.81269 | 32.06973 |
| 306 | ilvY | 0.668427 | 0.144908 | 2 | 894 | 0.838909 | 97.81269 | 31.96493 |
| 307 | yfeX | 0.672913 | 0.146446 | 2 | 900 | 0.834322 | 98.8512 | 32.19909 |
| 308 | ompG | 0.677399 | 0.147988 | 2 | 906 | 0.829773 | 99.89198 | 32.43246 |
| 309 | nhaR | 0.677399 | 0.147988 | 2 | 906 | 0.829773 | 99.89198 | 32.3275 |

| | | | | | | | | |
|---|---|---|---|---|---|---|---|---|
| 310 | gltI | 0.679643 | 0.14876 | 2 | 909 | 0.827513 | 100.4132 | 32.39136 |
| 311 | fucI | 1.327882 | 0.149356 | 3 | 1776 | 0.825777 | 100.8153 | 32.41649 |
| 312 | upstream p | 0.683381 | 0.150049 | 2 | 914 | 0.823767 | 101.2831 | 32.46254 |
| 313 | gcl | 1.332368 | 0.150405 | 3 | 1782 | 0.822737 | 101.5237 | 32.43567 |
| 314 | oxyR | 0.686372 | 0.151082 | 2 | 918 | 0.820788 | 101.9802 | 32.47776 |
| 315 | ydcI | 0.690858 | 0.152633 | 2 | 924 | 0.816351 | 103.0275 | 32.70715 |
| 316 | ddpF | 0.693101 | 0.15341 | 2 | 927 | 0.814145 | 103.552 | 32.76961 |
| 317 | alsK | 0.695344 | 0.154188 | 2 | 930 | 0.811949 | 104.0769 | 32.83185 |
| 318 | mmuM | 0.697587 | 0.154967 | 2 | 933 | 0.809762 | 104.6024 | 32.89385 |
| 319 | fixB | 0.704316 | 0.157307 | 2 | 942 | 0.803253 | 106.182 | 33.28589 |
| 320 | hycC | 1.366014 | 0.15835 | 3 | 1827 | 0.800381 | 106.8864 | 33.402 |
| 321 | paaX | 0.711045 | 0.159653 | 2 | 951 | 0.796822 | 107.766 | 33.57196 |
| 322 | gshB | 0.711045 | 0.159653 | 2 | 951 | 0.796822 | 107.766 | 33.4677 |
| 323 | corA | 0.711045 | 0.159653 | 2 | 951 | 0.796822 | 107.766 | 33.36408 |
| 324 | rpoD | 1.377229 | 0.161027 | 3 | 1842 | 0.793102 | 108.693 | 33.54722 |
| 325 | fecD | 0.715531 | 0.161221 | 2 | 957 | 0.792577 | 108.8244 | 33.48443 |
| 326 | yjiA | 0.715531 | 0.161221 | 2 | 957 | 0.792577 | 108.8244 | 33.38172 |
| 327 | paoB | 0.715531 | 0.161221 | 2 | 957 | 0.792577 | 108.8244 | 33.27964 |
| 328 | aes | 0.717774 | 0.162006 | 2 | 960 | 0.790468 | 109.3543 | 33.33974 |
| 329 | ssuA | 0.717774 | 0.162006 | 2 | 960 | 0.790468 | 109.3543 | 33.2384 |
| 330 | stfQ | 0.720017 | 0.162792 | 2 | 963 | 0.788366 | 109.8847 | 33.2984 |
| 331 | yhbE | 0.72226 | 0.163579 | 2 | 966 | 0.786273 | 110.4156 | 33.35818 |
| 332 | iaaA | 0.72226 | 0.163579 | 2 | 966 | 0.786273 | 110.4156 | 33.2577 |
| 333 | glk | 0.72226 | 0.163579 | 2 | 966 | 0.786273 | 110.4156 | 33.15783 |
| 334 | sapB | 0.72226 | 0.163579 | 2 | 966 | 0.786273 | 110.4156 | 33.05856 |
| 335 | ycjG | 0.72226 | 0.163579 | 2 | 966 | 0.786273 | 110.4156 | 32.95987 |
| 336 | fcl | 0.72226 | 0.163579 | 2 | 966 | 0.786273 | 110.4156 | 32.86178 |
| 337 | dxs | 1.392931 | 0.164796 | 3 | 1863 | 0.783052 | 111.2376 | 33.00819 |
| 338 | kefC | 1.392931 | 0.164796 | 3 | 1863 | 0.783052 | 111.2376 | 32.91053 |
| 339 | manX | 0.726746 | 0.165154 | 2 | 972 | 0.782112 | 111.4787 | 32.88456 |
| 340 | ybeQ | 0.731233 | 0.166731 | 2 | 978 | 0.777983 | 112.5435 | 33.10104 |
| 341 | rcsC | 2.130892 | 0.167234 | 4 | 2850 | 0.776675 | 112.8832 | 33.10357 |
| 342 | cld | 0.733476 | 0.167521 | 2 | 981 | 0.775931 | 113.0766 | 33.06334 |
| 343 | ydaN | 0.735719 | 0.168311 | 2 | 984 | 0.773887 | 113.6101 | 33.12249 |
| 344 | kdsD | 0.737962 | 0.169102 | 2 | 987 | 0.77185 | 114.1441 | 33.18143 |
| 345 | flk | 0.744691 | 0.171479 | 2 | 996 | 0.765788 | 115.7485 | 33.5503 |
| 346 | idnR | 0.746934 | 0.172273 | 2 | 999 | 0.763783 | 116.2842 | 33.60814 |
| 347 | yphD | 0.746934 | 0.172273 | 2 | 999 | 0.763783 | 116.2842 | 33.51128 |
| 348 | yoaA | 1.428819 | 0.173509 | 3 | 1911 | 0.760678 | 117.1186 | 33.65478 |
| 349 | dppF | 0.75142 | 0.173862 | 2 | 1005 | 0.759796 | 117.3566 | 33.62655 |
| 350 | fliM | 0.75142 | 0.173862 | 2 | 1005 | 0.759796 | 117.3566 | 33.53047 |
| 351 | hflC | 0.75142 | 0.173862 | 2 | 1005 | 0.759796 | 117.3566 | 33.43494 |
| 352 | ynhG | 0.75142 | 0.173862 | 2 | 1005 | 0.759796 | 117.3566 | 33.33996 |
| 353 | hypE | 0.755906 | 0.175453 | 2 | 1011 | 0.755839 | 118.4307 | 33.54978 |
| 354 | yhjD | 0.758149 | 0.176249 | 2 | 1014 | 0.753872 | 118.9684 | 33.60688 |
| 355 | rnb | 1.446764 | 0.177913 | 3 | 1935 | 0.749792 | 120.0914 | 33.82857 |
| 356 | yjjN | 0.764878 | 0.178642 | 2 | 1023 | 0.748016 | 120.5835 | 33.87178 |
| 357 | ytfT | 0.767121 | 0.179441 | 2 | 1026 | 0.746078 | 121.1227 | 33.92792 |
| 358 | purR | 0.767121 | 0.179441 | 2 | 1026 | 0.746078 | 121.1227 | 33.83314 |
| 359 | yfiF | 0.776093 | 0.182641 | 2 | 1038 | 0.738401 | 123.2828 | 34.34063 |
| 360 | yghZ | 0.778336 | 0.183443 | 2 | 1041 | 0.7365 | 123.8238 | 34.39549 |
| 361 | yraQ | 0.778336 | 0.183443 | 2 | 1041 | 0.7365 | 123.8238 | 34.30021 |

| | | | | | | | | |
|---|---|---|---|---|---|---|---|---|
| 362 | cheA | 1.469194 | 0.183461 | 3 | 1965 | 0.736456 | 123.8361 | 34.20887 |
| 363 | guaC | 0.78058 | 0.184245 | 2 | 1044 | 0.734605 | 124.3651 | 34.26035 |
| 364 | ypfG | 0.78058 | 0.184245 | 2 | 1044 | 0.734605 | 124.3651 | 34.16622 |
| 365 | galT | 0.782823 | 0.185047 | 2 | 1047 | 0.732718 | 124.9067 | 34.22101 |
| 366 | rsxD | 0.791795 | 0.188261 | 2 | 1059 | 0.725239 | 127.0765 | 34.72035 |
| 367 | yphC | 0.794038 | 0.189066 | 2 | 1062 | 0.723386 | 127.6197 | 34.77377 |
| 368 | ycjF | 0.794038 | 0.189066 | 2 | 1062 | 0.723386 | 127.6197 | 34.67928 |
| 369 | queA | 0.800767 | 0.191484 | 2 | 1071 | 0.717868 | 129.2514 | 35.02748 |
| 370 | ugpC | 0.800767 | 0.191484 | 2 | 1071 | 0.717868 | 129.2514 | 34.93281 |
| 371 | glpQ | 0.805253 | 0.193097 | 2 | 1077 | 0.714224 | 130.3407 | 35.13227 |
| 372 | rep | 1.511812 | 0.194124 | 3 | 2022 | 0.711921 | 131.0337 | 35.22412 |
| 373 | malS | 1.518541 | 0.195822 | 3 | 2031 | 0.708139 | 132.1796 | 35.43688 |
| 374 | rlpA | 0.814225 | 0.19633 | 2 | 1089 | 0.707013 | 132.5228 | 35.43391 |
| 375 | ybdH | 0.814225 | 0.19633 | 2 | 1089 | 0.707013 | 132.5228 | 35.33942 |
| 376 | yaiW | 0.818711 | 0.197949 | 2 | 1095 | 0.703447 | 133.6156 | 35.53606 |
| 377 | dcp | 1.529756 | 0.198659 | 3 | 2046 | 0.701892 | 134.0947 | 35.56889 |
| 378 | mltA | 0.820954 | 0.198759 | 2 | 1098 | 0.701673 | 134.1624 | 35.49269 |
| 379 | downstrear | 0.821702 | 0.199029 | 2 | 1099 | 0.701083 | 134.3447 | 35.44715 |
| 380 | upstream y | 1.531999 | 0.199227 | 3 | 2049 | 0.700651 | 134.4786 | 35.38909 |
| 381 | lptF | 0.823197 | 0.19957 | 2 | 1101 | 0.699906 | 134.7094 | 35.35681 |
| 382 | chaA | 0.823197 | 0.19957 | 2 | 1101 | 0.699906 | 134.7094 | 35.26425 |
| 383 | helD | 1.536486 | 0.200366 | 3 | 2055 | 0.698176 | 135.247 | 35.31253 |
| 384 | asd | 0.82544 | 0.20038 | 2 | 1104 | 0.698145 | 135.2567 | 35.22311 |
| 385 | ribD | 0.82544 | 0.20038 | 2 | 1104 | 0.698145 | 135.2567 | 35.13162 |
| 386 | lacZ | 2.299121 | 0.200474 | 4 | 3075 | 0.697942 | 135.3201 | 35.05702 |
| 387 | ybiR | 0.836656 | 0.20444 | 2 | 1119 | 0.689434 | 137.9972 | 35.65818 |
| 388 | fes | 0.841142 | 0.206067 | 2 | 1125 | 0.685992 | 139.095 | 35.84923 |
| 389 | smf | 0.841142 | 0.206067 | 2 | 1125 | 0.685992 | 139.095 | 35.75708 |
| 390 | yhcG | 0.843385 | 0.20688 | 2 | 1128 | 0.684281 | 139.6443 | 35.80623 |
| 391 | fliK | 0.843385 | 0.20688 | 2 | 1128 | 0.684281 | 139.6443 | 35.71466 |
| 392 | ybhS | 0.847871 | 0.208509 | 2 | 1134 | 0.680875 | 140.7436 | 35.90397 |
| 393 | norW | 0.847871 | 0.208509 | 2 | 1134 | 0.680875 | 140.7436 | 35.81261 |
| 394 | potG | 0.847871 | 0.208509 | 2 | 1134 | 0.680875 | 140.7436 | 35.72172 |
| 395 | fepE | 0.847871 | 0.208509 | 2 | 1134 | 0.680875 | 140.7436 | 35.63128 |
| 396 | yhhI | 0.850114 | 0.209324 | 2 | 1137 | 0.679181 | 141.2935 | 35.68018 |
| 397 | nlpD | 0.852357 | 0.210139 | 2 | 1140 | 0.677494 | 141.8437 | 35.72889 |
| 398 | caiA | 0.8546 | 0.210954 | 2 | 1143 | 0.675812 | 142.3941 | 35.77741 |
| 399 | carA | 0.859086 | 0.212586 | 2 | 1149 | 0.672465 | 143.4955 | 35.96379 |
| 400 | agaS | 0.863572 | 0.214219 | 2 | 1155 | 0.669142 | 144.5977 | 36.14944 |
| 401 | metB | 0.868058 | 0.215853 | 2 | 1161 | 0.665842 | 145.7007 | 36.33435 |
| 402 | ynjB | 0.872544 | 0.217488 | 2 | 1167 | 0.662564 | 146.8045 | 36.51853 |
| 403 | nrdE | 1.603777 | 0.217618 | 3 | 2145 | 0.662305 | 146.8922 | 36.44967 |
| 404 | torC | 0.87703 | 0.219124 | 2 | 1173 | 0.659309 | 147.909 | 36.61114 |
| 405 | iadA | 0.87703 | 0.219124 | 2 | 1173 | 0.659309 | 147.909 | 36.52074 |
| 406 | hybB | 0.881517 | 0.220762 | 2 | 1179 | 0.656076 | 149.0142 | 36.703 |
| 407 | glcB | 1.623964 | 0.222853 | 3 | 2172 | 0.651982 | 150.4255 | 36.95959 |
| 408 | bcr | 0.890489 | 0.224039 | 2 | 1191 | 0.649676 | 151.2265 | 37.06532 |
| 409 | hofC | 0.899461 | 0.22732 | 2 | 1203 | 0.643362 | 153.4413 | 37.5162 |
| 410 | yhiN | 0.899461 | 0.22732 | 2 | 1203 | 0.643362 | 153.4413 | 37.4247 |
| 411 | proV | 0.899461 | 0.22732 | 2 | 1203 | 0.643362 | 153.4413 | 37.33364 |
| 412 | paaJ | 0.901704 | 0.228141 | 2 | 1206 | 0.641796 | 153.9953 | 37.3775 |
| 413 | gltS | 0.901704 | 0.228141 | 2 | 1206 | 0.641796 | 153.9953 | 37.287 |

| | | | | | | | | |
|---|---|---|---|---|---|---|---|---|
| 414 | dacA | 0.90619 | 0.229783 | 2 | 1212 | 0.638681 | 155.1038 | 37.46469 |
| 415 | ydhC | 0.90619 | 0.229783 | 2 | 1212 | 0.638681 | 155.1038 | 37.37441 |
| 416 | sucB | 0.910676 | 0.231426 | 2 | 1218 | 0.635587 | 156.2128 | 37.55116 |
| 417 | ybdN | 0.912919 | 0.232248 | 2 | 1221 | 0.634048 | 156.7675 | 37.59413 |
| 418 | cusB | 0.915162 | 0.23307 | 2 | 1224 | 0.632513 | 157.3223 | 37.63692 |
| 419 | deoB | 0.915162 | 0.23307 | 2 | 1224 | 0.632513 | 157.3223 | 37.54709 |
| 420 | eutH | 0.917405 | 0.233892 | 2 | 1227 | 0.630984 | 157.8772 | 37.58982 |
| 421 | yhfW | 0.917405 | 0.233892 | 2 | 1227 | 0.630984 | 157.8772 | 37.50053 |
| 422 | pepT | 0.917405 | 0.233892 | 2 | 1227 | 0.630984 | 157.8772 | 37.41167 |
| 423 | yqeG | 0.919648 | 0.234714 | 2 | 1230 | 0.62946 | 158.4323 | 37.45444 |
| 424 | serA | 0.921891 | 0.235537 | 2 | 1233 | 0.627941 | 158.9874 | 37.49703 |
| 425 | yfgF | 1.677797 | 0.236928 | 3 | 2244 | 0.625384 | 159.9261 | 37.62968 |
| 426 | ptsP | 1.68004 | 0.237517 | 3 | 2247 | 0.624305 | 160.3243 | 37.6348 |
| 427 | intA | 0.92862 | 0.238005 | 2 | 1242 | 0.623414 | 160.6535 | 37.62377 |
| 428 | ybhO | 0.92862 | 0.238005 | 2 | 1242 | 0.623414 | 160.6535 | 37.53586 |
| 429 | hypF | 1.684526 | 0.238698 | 3 | 2253 | 0.622152 | 161.121 | 37.55735 |
| 430 | ftsW | 0.930863 | 0.238828 | 2 | 1245 | 0.621914 | 161.209 | 37.49047 |
| 431 | frsA | 0.930863 | 0.238828 | 2 | 1245 | 0.621914 | 161.209 | 37.40349 |
| 432 | yjeP | 2.485293 | 0.239285 | 4 | 3324 | 0.621085 | 161.5172 | 37.38825 |
| 433 | mscM | 0.933107 | 0.239651 | 2 | 1248 | 0.62042 | 161.7647 | 37.35905 |
| 434 | fryC | 0.933107 | 0.239651 | 2 | 1248 | 0.62042 | 161.7647 | 37.27297 |
| 435 | entS | 0.93535 | 0.240475 | 2 | 1251 | 0.618931 | 162.3204 | 37.31504 |
| 436 | ycjT | 1.695742 | 0.241653 | 3 | 2268 | 0.616808 | 163.1159 | 37.41189 |
| 437 | hflK | 0.942079 | 0.242945 | 2 | 1260 | 0.614491 | 163.9881 | 37.52588 |
| 438 | rhaA | 0.942079 | 0.242945 | 2 | 1260 | 0.614491 | 163.9881 | 37.44021 |
| 439 | ydeP | 1.704714 | 0.244022 | 3 | 2280 | 0.612572 | 164.7146 | 37.52042 |
| 440 | puuE | 0.946565 | 0.244593 | 2 | 1266 | 0.611556 | 165.1003 | 37.52279 |
| 441 | uidC | 0.946565 | 0.244593 | 2 | 1266 | 0.611556 | 165.1003 | 37.43771 |
| 442 | sufD | 0.951051 | 0.246241 | 2 | 1272 | 0.608639 | 166.2128 | 37.6047 |
| 443 | hisS | 0.953294 | 0.247065 | 2 | 1275 | 0.607188 | 166.7691 | 37.64539 |
| 444 | yggP | 0.955537 | 0.24789 | 2 | 1278 | 0.605742 | 167.3255 | 37.68592 |
| 445 | livM | 0.955537 | 0.24789 | 2 | 1278 | 0.605742 | 167.3255 | 37.60123 |
| 446 | bioA | 0.964509 | 0.251188 | 2 | 1290 | 0.600002 | 169.5516 | 38.01605 |
| 447 | dcuA | 0.973481 | 0.254487 | 2 | 1302 | 0.594335 | 171.7785 | 38.42919 |
| 448 | ybbY | 0.973481 | 0.254487 | 2 | 1302 | 0.594335 | 171.7785 | 38.34341 |
| 449 | upstream y | 3.393726 | 0.254653 | 5 | 4539 | 0.594051 | 171.8909 | 38.28306 |
| 450 | yhfT | 0.975724 | 0.255312 | 2 | 1305 | 0.59293 | 172.3353 | 38.29673 |
| 451 | gltP | 0.982454 | 0.257786 | 2 | 1314 | 0.58874 | 174.0059 | 38.58223 |
| 452 | murD | 0.984697 | 0.258612 | 2 | 1317 | 0.587352 | 174.5628 | 38.62008 |
| 453 | brnQ | 0.98694 | 0.259437 | 2 | 1320 | 0.585969 | 175.1197 | 38.65777 |
| 454 | ykgC | 0.991426 | 0.261087 | 2 | 1326 | 0.583215 | 176.2336 | 38.81798 |
| 455 | yliF | 0.993669 | 0.261912 | 2 | 1329 | 0.581845 | 176.7906 | 38.85508 |
| 456 | hslU | 0.995912 | 0.262737 | 2 | 1332 | 0.580479 | 177.3476 | 38.89201 |
| 457 | qseF | 0.998155 | 0.263562 | 2 | 1335 | 0.579117 | 177.9046 | 38.92879 |
| 458 | glmM | 1.000398 | 0.264387 | 2 | 1338 | 0.577759 | 178.4616 | 38.9654 |
| 459 | frlA | 1.000398 | 0.264387 | 2 | 1338 | 0.577759 | 178.4616 | 38.88051 |
| 460 | gudD | 1.002641 | 0.265213 | 2 | 1341 | 0.576406 | 179.0185 | 38.91707 |
| 461 | rarA | 1.004884 | 0.266038 | 2 | 1344 | 0.575057 | 179.5755 | 38.95348 |
| 462 | chbF | 1.011613 | 0.268513 | 2 | 1353 | 0.571034 | 181.2464 | 39.23083 |
| 463 | yeeF | 1.016099 | 0.270163 | 2 | 1359 | 0.568373 | 182.3603 | 39.38668 |
| 464 | yegQ | 1.018342 | 0.270988 | 2 | 1362 | 0.567049 | 182.9172 | 39.42181 |
| 465 | allB | 1.018342 | 0.270988 | 2 | 1362 | 0.567049 | 182.9172 | 39.33704 |

| | | | | | | | | |
|---|---|---|---|---|---|---|---|---|
| 466 | ydbK | 2.635577 | 0.271759 | 4 | 3525 | 0.565815 | 183.4377 | 39.3643 |
| 467 | sdaB | 1.022828 | 0.272638 | 2 | 1368 | 0.564413 | 184.031 | 39.40706 |
| 468 | pgaA | 1.81238 | 0.272696 | 3 | 2424 | 0.564321 | 184.07 | 39.3312 |
| 469 | cpsG | 1.025071 | 0.273463 | 2 | 1371 | 0.563101 | 184.5878 | 39.35774 |
| 470 | glmU | 1.025071 | 0.273463 | 2 | 1371 | 0.563101 | 184.5878 | 39.274 |
| 471 | dbpA | 1.027314 | 0.274288 | 2 | 1374 | 0.561793 | 185.1446 | 39.30883 |
| 472 | fliI | 1.027314 | 0.274288 | 2 | 1374 | 0.561793 | 185.1446 | 39.22555 |
| 473 | ybiW | 1.819109 | 0.274501 | 3 | 2433 | 0.561456 | 185.2884 | 39.17302 |
| 474 | ygeH | 1.029557 | 0.275113 | 2 | 1377 | 0.560489 | 185.7014 | 39.1775 |
| 475 | patA | 1.0318 | 0.275938 | 2 | 1380 | 0.559189 | 186.2581 | 39.21223 |
| 476 | radA | 1.034044 | 0.276763 | 2 | 1383 | 0.557893 | 186.8148 | 39.24681 |
| 477 | atpD | 1.034044 | 0.276763 | 2 | 1383 | 0.557893 | 186.8148 | 39.16453 |
| 478 | fadE | 1.828081 | 0.27691 | 3 | 2445 | 0.557662 | 186.9141 | 39.10337 |
| 479 | narU | 1.03853 | 0.278412 | 2 | 1389 | 0.555312 | 187.9281 | 39.23341 |
| 480 | galP | 1.043016 | 0.280061 | 2 | 1395 | 0.552747 | 189.0411 | 39.38357 |
| 481 | hsdS | 1.043016 | 0.280061 | 2 | 1395 | 0.552747 | 189.0411 | 39.30169 |
| 482 | yhjA | 1.045259 | 0.280885 | 2 | 1398 | 0.551471 | 189.5976 | 39.3356 |
| 483 | gadB | 1.047502 | 0.28171 | 2 | 1401 | 0.550198 | 190.154 | 39.36936 |
| 484 | leuC | 1.047502 | 0.28171 | 2 | 1401 | 0.550198 | 190.154 | 39.28802 |
| 485 | fumC | 1.049745 | 0.282534 | 2 | 1404 | 0.548929 | 190.7104 | 39.32172 |
| 486 | ydcR | 1.051988 | 0.283358 | 2 | 1407 | 0.547665 | 191.2667 | 39.35528 |
| 487 | gnd | 1.051988 | 0.283358 | 2 | 1407 | 0.547665 | 191.2667 | 39.27447 |
| 488 | melB | 1.054231 | 0.284182 | 2 | 1410 | 0.546403 | 191.8229 | 39.30797 |
| 489 | yfjI | 1.054231 | 0.284182 | 2 | 1410 | 0.546403 | 191.8229 | 39.22758 |
| 490 | pykF | 1.056474 | 0.285006 | 2 | 1413 | 0.545146 | 192.379 | 39.26103 |
| 491 | araE | 1.06096 | 0.286654 | 2 | 1419 | 0.542643 | 193.4912 | 39.40757 |
| 492 | fryA | 1.866213 | 0.287167 | 3 | 2496 | 0.541865 | 193.8379 | 39.39794 |
| 493 | ymdC | 1.063203 | 0.287477 | 2 | 1422 | 0.541397 | 194.0471 | 39.36047 |
| 494 | miaB | 1.065446 | 0.288301 | 2 | 1425 | 0.540154 | 194.603 | 39.39333 |
| 495 | degP | 1.065446 | 0.288301 | 2 | 1425 | 0.540154 | 194.603 | 39.31374 |
| 496 | paaH | 1.067689 | 0.289124 | 2 | 1428 | 0.538916 | 195.1588 | 39.34654 |
| 497 | yehI | 2.716327 | 0.289511 | 4 | 3633 | 0.538336 | 195.4196 | 39.31985 |
| 498 | ybjT | 1.069932 | 0.289948 | 2 | 1431 | 0.537681 | 195.7146 | 39.30011 |
| 499 | dacB | 1.072175 | 0.290771 | 2 | 1434 | 0.536449 | 196.2702 | 39.33271 |
| 500 | cusS | 1.078904 | 0.293239 | 2 | 1443 | 0.532778 | 197.9366 | 39.58733 |
| 501 | uxaB | 1.085634 | 0.295707 | 2 | 1452 | 0.529138 | 199.6023 | 39.84077 |
| 502 | pepD | 1.09012 | 0.297351 | 2 | 1458 | 0.52673 | 200.7122 | 39.98251 |
| 503 | ydfI | 1.092363 | 0.298173 | 2 | 1461 | 0.525531 | 201.267 | 40.01332 |
| 504 | phoQ | 1.092363 | 0.298173 | 2 | 1461 | 0.525531 | 201.267 | 39.93393 |
| 505 | ttdT | 1.094606 | 0.298995 | 2 | 1464 | 0.524336 | 201.8217 | 39.96469 |
| 506 | rng | 1.099092 | 0.300638 | 2 | 1470 | 0.521956 | 202.9308 | 40.1049 |
| 507 | narZ | 2.797077 | 0.307412 | 4 | 3741 | 0.512279 | 207.5032 | 40.92766 |
| 508 | acnB | 1.942477 | 0.307758 | 3 | 2598 | 0.511791 | 207.7367 | 40.89305 |
| 509 | topA | 1.942477 | 0.307758 | 3 | 2598 | 0.511791 | 207.7367 | 40.81271 |
| 510 | ravA | 1.119279 | 0.308023 | 2 | 1497 | 0.511416 | 207.9158 | 40.76781 |
| 511 | lyxK | 1.119279 | 0.308023 | 2 | 1497 | 0.511416 | 207.9158 | 40.68803 |
| 512 | elfC | 1.94472 | 0.308365 | 3 | 2601 | 0.510935 | 208.1462 | 40.65355 |
| 513 | glcD | 1.121522 | 0.308843 | 2 | 1500 | 0.510262 | 208.4691 | 40.63725 |
| 514 | yjgR | 1.123765 | 0.309663 | 2 | 1503 | 0.509111 | 209.0222 | 40.6658 |
| 515 | rbsA | 1.126008 | 0.310482 | 2 | 1506 | 0.507964 | 209.5752 | 40.69421 |
| 516 | casA | 1.128251 | 0.311301 | 2 | 1509 | 0.50682 | 210.1281 | 40.72249 |
| 517 | bcsA | 1.958178 | 0.312005 | 3 | 2619 | 0.505838 | 210.6035 | 40.73569 |

| | | | | | | | | |
|---|---|---|---|---|---|---|---|---|
| 518 | araG | 1.132738 | 0.312938 | 2 | 1515 | 0.504541 | 211.2333 | 40.77864 |
| 519 | lysU | 1.134981 | 0.313757 | 2 | 1518 | 0.503407 | 211.7858 | 40.8065 |
| 520 | ycgG | 1.139467 | 0.315393 | 2 | 1524 | 0.501148 | 212.8902 | 40.94041 |
| 521 | ybcK | 1.14171 | 0.316211 | 2 | 1527 | 0.500024 | 213.4421 | 40.96778 |
| 522 | narH | 1.150682 | 0.319479 | 2 | 1539 | 0.495557 | 215.6485 | 41.31197 |
| 523 | tyrR | 1.152925 | 0.320296 | 2 | 1542 | 0.494449 | 216.1997 | 41.33837 |
| 524 | aceE | 1.991824 | 0.32111 | 3 | 2664 | 0.493346 | 216.7496 | 41.36442 |
| 525 | ilvA | 1.155168 | 0.321112 | 2 | 1545 | 0.493343 | 216.7507 | 41.28584 |
| 526 | ygcB | 1.994067 | 0.321718 | 3 | 2667 | 0.492525 | 217.1594 | 41.28505 |
| 527 | caiC | 1.161897 | 0.32356 | 2 | 1554 | 0.490046 | 218.4027 | 41.44265 |
| 528 | rtn | 1.16414 | 0.324375 | 2 | 1557 | 0.488953 | 218.9531 | 41.46839 |
| 529 | cydA | 1.173112 | 0.327634 | 2 | 1569 | 0.484612 | 221.1527 | 41.80581 |
| 530 | entF | 2.9025 | 0.330936 | 4 | 3882 | 0.480256 | 223.3816 | 42.14747 |
| 531 | ybiP | 1.184328 | 0.331701 | 2 | 1584 | 0.479253 | 223.8981 | 42.16537 |
| 532 | prfC | 1.188814 | 0.333326 | 2 | 1590 | 0.477131 | 224.995 | 42.29229 |
| 533 | lsrK | 1.191057 | 0.334138 | 2 | 1593 | 0.476074 | 225.5431 | 42.31578 |
| 534 | ygiS | 1.202272 | 0.338194 | 2 | 1608 | 0.470834 | 228.2808 | 42.74921 |
| 535 | entE | 1.204515 | 0.339004 | 2 | 1611 | 0.469795 | 228.8277 | 42.77153 |
| 536 | thiP | 1.204515 | 0.339004 | 2 | 1611 | 0.469795 | 228.8277 | 42.69173 |
| 537 | mppA | 1.206758 | 0.339814 | 2 | 1614 | 0.468759 | 229.3744 | 42.71404 |
| 538 | pck | 1.213487 | 0.342242 | 2 | 1623 | 0.465667 | 231.0132 | 42.93926 |
| 539 | ubiB | 1.226945 | 0.347089 | 2 | 1641 | 0.459559 | 234.2851 | 43.46662 |
| 540 | yojI | 1.229188 | 0.347896 | 2 | 1644 | 0.458551 | 234.8296 | 43.48697 |
| 541 | yidC | 1.231431 | 0.348702 | 2 | 1647 | 0.457545 | 235.3739 | 43.5072 |
| 542 | actP | 1.233675 | 0.349508 | 2 | 1650 | 0.456543 | 235.918 | 43.52731 |
| 543 | yfaQ | 1.233675 | 0.349508 | 2 | 1650 | 0.456543 | 235.918 | 43.44715 |
| 544 | yjcE | 1.233675 | 0.349508 | 2 | 1650 | 0.456543 | 235.918 | 43.36728 |
| 545 | lldP | 1.238161 | 0.351119 | 2 | 1656 | 0.454545 | 237.0055 | 43.48724 |
| 546 | opgD | 1.238161 | 0.351119 | 2 | 1656 | 0.454545 | 237.0055 | 43.40759 |
| 547 | yqiK | 1.242647 | 0.352729 | 2 | 1662 | 0.452559 | 238.092 | 43.52687 |
| 548 | tar | 1.242647 | 0.352729 | 2 | 1662 | 0.452559 | 238.092 | 43.44744 |
| 549 | hyfG | 1.247133 | 0.354337 | 2 | 1668 | 0.450583 | 239.1775 | 43.56603 |
| 550 | hscC | 1.249376 | 0.355141 | 2 | 1671 | 0.4496 | 239.7199 | 43.58544 |
| 551 | glnE | 2.124163 | 0.356889 | 3 | 2841 | 0.447467 | 240.9001 | 43.72053 |
| 552 | bcsG | 1.256105 | 0.357549 | 2 | 1680 | 0.446664 | 241.3457 | 43.72205 |
| 553 | ybjL | 1.260591 | 0.359153 | 2 | 1686 | 0.44472 | 242.4284 | 43.83876 |
| 554 | fadD | 1.260591 | 0.359153 | 2 | 1686 | 0.44472 | 242.4284 | 43.75963 |
| 555 | yddA | 1.260591 | 0.359153 | 2 | 1686 | 0.44472 | 242.4284 | 43.68079 |
| 556 | ilvB | 1.262834 | 0.359955 | 2 | 1689 | 0.443752 | 242.9693 | 43.69951 |
| 557 | yedQ | 1.26732 | 0.361556 | 2 | 1695 | 0.441824 | 244.0504 | 43.81516 |
| 558 | xdhD | 2.146594 | 0.362934 | 3 | 2871 | 0.440173 | 244.9803 | 43.90327 |
| 559 | narQ | 1.271806 | 0.363156 | 2 | 1701 | 0.439906 | 245.1305 | 43.85161 |
| 560 | gcvP | 2.148837 | 0.363538 | 3 | 2874 | 0.439451 | 245.3879 | 43.81927 |
| 561 | hybC | 1.274049 | 0.363956 | 2 | 1704 | 0.438951 | 245.6702 | 43.79147 |
| 562 | ilvI | 1.289751 | 0.369541 | 2 | 1725 | 0.432337 | 249.4404 | 44.38441 |
| 563 | ptsI | 1.291994 | 0.370338 | 2 | 1728 | 0.431402 | 249.9779 | 44.40106 |
| 564 | yijP | 1.29648 | 0.371929 | 2 | 1734 | 0.42954 | 251.0522 | 44.5128 |
| 565 | dnaG | 1.305452 | 0.375107 | 2 | 1746 | 0.425845 | 253.1974 | 44.8137 |
| 566 | msbA | 1.307695 | 0.375901 | 2 | 1749 | 0.424927 | 253.733 | 44.82915 |
| 567 | ftsI | 1.321153 | 0.380653 | 2 | 1767 | 0.419471 | 256.9406 | 45.3158 |
| 568 | ade | 1.321153 | 0.380653 | 2 | 1767 | 0.419471 | 256.9406 | 45.23602 |
| 569 | nfrA | 2.222857 | 0.383409 | 3 | 2973 | 0.416338 | 258.8009 | 45.48345 |

| | | | | | | | | |
|---|---|---|---|---|---|---|---|---|
| 570 | ygeV | 1.330125 | 0.383812 | 2 | 1779 | 0.415881 | 259.0732 | 45.45143 |
| 571 | nuoC | 1.339098 | 0.386964 | 2 | 1791 | 0.412329 | 261.201 | 45.74449 |
| 572 | cysJ | 1.345827 | 0.389324 | 2 | 1800 | 0.409689 | 262.7937 | 45.94296 |
| 573 | malZ | 1.359285 | 0.394031 | 2 | 1818 | 0.40447 | 265.9709 | 46.41726 |
| 574 | typA | 1.363771 | 0.395596 | 2 | 1824 | 0.402748 | 267.0275 | 46.52046 |
| 575 | uvrC | 1.3705 | 0.397941 | 2 | 1833 | 0.400182 | 268.6099 | 46.71477 |
| 576 | yfbS | 1.3705 | 0.397941 | 2 | 1833 | 0.400182 | 268.6099 | 46.63367 |
| 577 | btuB | 1.379472 | 0.40106 | 2 | 1845 | 0.396791 | 270.7154 | 46.91775 |
| 578 | sppA | 1.388445 | 0.404171 | 2 | 1857 | 0.393434 | 272.8157 | 47.19995 |
| 579 | ppiD | 1.39966 | 0.40805 | 2 | 1872 | 0.389287 | 275.4336 | 47.57058 |
| 580 | bglF | 1.404146 | 0.409598 | 2 | 1878 | 0.387642 | 276.4785 | 47.6687 |
| 581 | mdtF | 2.32828 | 0.411446 | 3 | 3114 | 0.385687 | 277.7264 | 47.80144 |
| 582 | acrD | 2.32828 | 0.411446 | 3 | 3114 | 0.385687 | 277.7264 | 47.71931 |
| 583 | flu | 2.332766 | 0.412631 | 3 | 3120 | 0.384438 | 278.526 | 47.77462 |
| 584 | deaD | 1.413118 | 0.412688 | 2 | 1890 | 0.384379 | 278.5641 | 47.69933 |
| 585 | acrB | 2.355197 | 0.418543 | 3 | 3150 | 0.37826 | 282.5167 | 48.29344 |
| 586 | nagE | 1.455736 | 0.427252 | 2 | 1947 | 0.369316 | 288.3951 | 49.21418 |
| 587 | gspD | 1.460222 | 0.428774 | 2 | 1953 | 0.367772 | 289.4225 | 49.30536 |
| 588 | topB | 1.466951 | 0.431053 | 2 | 1962 | 0.365469 | 290.9608 | 49.48313 |
| 589 | aaeB | 1.471437 | 0.43257 | 2 | 1968 | 0.363944 | 291.9846 | 49.57293 |
| 590 | speA | 1.478166 | 0.434841 | 2 | 1977 | 0.36167 | 293.5174 | 49.74871 |
| 591 | gmr | 1.484895 | 0.437107 | 2 | 1986 | 0.359413 | 295.0469 | 49.92333 |
| 592 | cirA | 1.489382 | 0.438614 | 2 | 1992 | 0.357917 | 296.0647 | 50.01093 |
| 593 | yghJ | 3.411671 | 0.444193 | 4 | 4563 | 0.352429 | 299.83 | 50.56155 |
| 594 | yphG | 2.453891 | 0.444306 | 3 | 3282 | 0.352318 | 299.9064 | 50.4893 |
| 595 | fadH | 1.509569 | 0.445372 | 2 | 2019 | 0.351277 | 300.6262 | 50.52541 |
| 596 | upstream i | 1.511812 | 0.44612 | 2 | 2022 | 0.350548 | 301.1311 | 50.52536 |
| 597 | uvrB | 1.511812 | 0.44612 | 2 | 2022 | 0.350548 | 301.1311 | 50.44073 |
| 598 | kdpB | 1.531999 | 0.452827 | 2 | 2049 | 0.344068 | 305.6582 | 51.11341 |
| 599 | lhr | 3.452046 | 0.452985 | 4 | 4617 | 0.343916 | 305.7648 | 51.04588 |
| 600 | yhjG | 1.540972 | 0.455793 | 2 | 2061 | 0.341233 | 307.6601 | 51.27668 |
| 601 | stfR | 2.514453 | 0.459889 | 3 | 3363 | 0.337347 | 310.425 | 51.65142 |
| 602 | kefA | 2.514453 | 0.459889 | 3 | 3363 | 0.337347 | 310.425 | 51.56562 |
| 603 | yjbH | 1.567888 | 0.464634 | 2 | 2097 | 0.332889 | 313.6278 | 52.01125 |
| 604 | yncD | 1.572374 | 0.466099 | 2 | 2103 | 0.331522 | 314.6169 | 52.08888 |
| 605 | ydhV | 1.572374 | 0.466099 | 2 | 2103 | 0.331522 | 314.6169 | 52.00279 |
| 606 | yhfK | 1.572374 | 0.466099 | 2 | 2103 | 0.331522 | 314.6169 | 51.91697 |
| 607 | rlmL | 1.57686 | 0.467562 | 2 | 2109 | 0.330161 | 315.6043 | 51.99411 |
| 608 | upstream y | 1.58658 | 0.470723 | 2 | 2122 | 0.327234 | 317.7381 | 52.25956 |
| 609 | speC | 1.597048 | 0.474115 | 2 | 2136 | 0.324116 | 320.0275 | 52.54967 |
| 610 | cadA | 1.60602 | 0.477011 | 2 | 2148 | 0.321471 | 321.9828 | 52.78406 |
| 611 | wzc | 1.617235 | 0.480619 | 2 | 2163 | 0.318199 | 324.4175 | 53.09616 |
| 612 | dnaE | 2.604175 | 0.482618 | 3 | 3483 | 0.316397 | 325.7671 | 53.22992 |
| 613 | katG | 1.630693 | 0.484927 | 2 | 2181 | 0.314324 | 327.3255 | 53.3973 |
| 614 | zntA | 1.644152 | 0.489212 | 2 | 2199 | 0.310503 | 330.2184 | 53.78149 |
| 615 | rsxC | 1.662096 | 0.494892 | 2 | 2223 | 0.30549 | 334.0518 | 54.31737 |
| 616 | yjdA | 1.666582 | 0.496305 | 2 | 2229 | 0.304251 | 335.0059 | 54.38408 |
| 617 | parC | 1.689013 | 0.503334 | 2 | 2259 | 0.298143 | 339.7508 | 55.06496 |
| 618 | ybhJ | 1.691256 | 0.504034 | 2 | 2262 | 0.29754 | 340.2229 | 55.05225 |
| 619 | clpA | 1.702471 | 0.507521 | 2 | 2277 | 0.294546 | 342.577 | 55.34361 |
| 620 | yfhM | 3.709996 | 0.507931 | 4 | 4962 | 0.294195 | 342.8535 | 55.29895 |
| 621 | pflB | 1.706957 | 0.508912 | 2 | 2283 | 0.293357 | 343.5156 | 55.31652 |

| | | | | | | | | |
|---|---|---|---|---|---|---|---|---|
| 622 | nrdA | 1.7092 | 0.509606 | 2 | 2286 | 0.292765 | 343.9842 | 55.30293 |
| 623 | bglX | 1.718172 | 0.512377 | 2 | 2298 | 0.29041 | 345.8544 | 55.51435 |
| 624 | upstream a | 1.721163 | 0.513298 | 2 | 2302 | 0.28963 | 346.4762 | 55.52504 |
| 625 | bisC | 1.745089 | 0.520627 | 2 | 2334 | 0.283474 | 351.4229 | 56.22766 |
| 626 | polB | 1.758547 | 0.524716 | 2 | 2352 | 0.280076 | 354.1833 | 56.57881 |
| 627 | pps | 1.778734 | 0.530806 | 2 | 2379 | 0.275064 | 358.294 | 57.14418 |
| 628 | gcd | 1.787707 | 0.533495 | 2 | 2391 | 0.272869 | 360.1094 | 57.34226 |
| 629 | malP | 1.78995 | 0.534166 | 2 | 2394 | 0.272324 | 360.5621 | 57.32307 |
| 630 | ybbP | 1.805651 | 0.538842 | 2 | 2415 | 0.268538 | 363.7186 | 57.73311 |
| 631 | gyrB | 1.805651 | 0.538842 | 2 | 2415 | 0.268538 | 363.7186 | 57.64161 |
| 632 | plsB | 1.81238 | 0.540836 | 2 | 2424 | 0.266934 | 365.0646 | 57.76339 |
| 633 | rnr | 1.825838 | 0.544807 | 2 | 2442 | 0.263758 | 367.7446 | 58.09551 |
| 634 | glgP | 1.830324 | 0.546125 | 2 | 2448 | 0.262708 | 368.6342 | 58.1442 |
| 635 | thrA | 1.84154 | 0.549408 | 2 | 2463 | 0.260105 | 370.8506 | 58.40166 |
| 636 | napA | 1.859484 | 0.554627 | 2 | 2487 | 0.255999 | 374.3731 | 58.8637 |
| 637 | purL | 2.906986 | 0.555651 | 3 | 3888 | 0.255198 | 375.0647 | 58.87985 |
| 638 | hrpA | 2.918201 | 0.558236 | 3 | 3903 | 0.253182 | 376.8094 | 59.06103 |
| 639 | copA | 1.872942 | 0.558512 | 2 | 2505 | 0.252967 | 376.9959 | 58.9978 |
| 640 | yraJ | 1.881914 | 0.561089 | 2 | 2517 | 0.250968 | 378.7354 | 59.1774 |
| 641 | nirB | 1.902102 | 0.566848 | 2 | 2544 | 0.246533 | 382.6224 | 59.69148 |
| 642 | torA | 1.904345 | 0.567484 | 2 | 2547 | 0.246046 | 383.052 | 59.66542 |
| 643 | mutS | 1.91556 | 0.570657 | 2 | 2562 | 0.243625 | 385.1932 | 59.90563 |
| 644 | clpB | 1.924532 | 0.573182 | 2 | 2574 | 0.241707 | 386.8979 | 60.07731 |
| 645 | leuS | 1.931261 | 0.575069 | 2 | 2583 | 0.24028 | 388.1716 | 60.18164 |
| 646 | sfmD | 1.946963 | 0.579448 | 2 | 2604 | 0.236985 | 391.1275 | 60.54606 |
| 647 | rpoB | 3.012409 | 0.579584 | 3 | 4029 | 0.236883 | 391.2194 | 60.46667 |
| 648 | fimD | 1.971636 | 0.586262 | 2 | 2637 | 0.231908 | 395.7272 | 61.06901 |
| 649 | ydbH | 1.973879 | 0.586878 | 2 | 2640 | 0.231452 | 396.1426 | 61.03891 |
| 650 | ppc | 1.982851 | 0.589333 | 2 | 2652 | 0.22964 | 397.7995 | 61.19993 |
| 651 | glnD | 1.998553 | 0.593602 | 2 | 2673 | 0.226504 | 400.6815 | 61.54862 |
| 652 | mgtA | 2.016497 | 0.598441 | 2 | 2697 | 0.222979 | 403.9477 | 61.95517 |
| 653 | secA | 2.023226 | 0.600244 | 2 | 2706 | 0.221672 | 405.1649 | 62.04669 |
| 654 | nuoG | 2.038928 | 0.604428 | 2 | 2727 | 0.218655 | 407.989 | 62.38364 |
| 655 | torS | 2.052386 | 0.607988 | 2 | 2745 | 0.216105 | 410.3918 | 62.65523 |
| 656 | yeeJ | 5.291342 | 0.609059 | 5 | 7077 | 0.21534 | 411.1151 | 62.66999 |
| 657 | rpoC | 3.158207 | 0.611306 | 3 | 4224 | 0.213742 | 412.6313 | 62.80538 |
| 658 | uvrA | 2.110705 | 0.62313 | 2 | 2823 | 0.205421 | 420.6129 | 63.92293 |
| 659 | mukB | 3.335407 | 0.647635 | 3 | 4461 | 0.18867 | 437.1534 | 66.33588 |
| 660 | upstream y | 2.269961 | 0.66216 | 2 | 3036 | 0.179037 | 446.958 | 67.72091 |
| 661 | ydiJ | 2.285662 | 0.665826 | 2 | 3057 | 0.176639 | 449.4327 | 67.99284 |
| 662 | sbcC | 2.352954 | 0.681175 | 2 | 3147 | 0.166741 | 459.7929 | 69.45513 |
| 663 | rne | 2.382113 | 0.687644 | 2 | 3186 | 0.162636 | 464.1599 | 70.00903 |
| 664 | carB | 2.40903 | 0.69352 | 2 | 3222 | 0.158941 | 468.1259 | 70.50088 |
| 665 | yegE | 2.480807 | 0.708741 | 2 | 3318 | 0.149512 | 478.4003 | 71.9399 |
| 666 | mfd | 2.577258 | 0.728193 | 2 | 3447 | 0.137754 | 491.53 | 73.8033 |
| 667 | hsdR | 2.626605 | 0.737711 | 2 | 3513 | 0.132114 | 497.9549 | 74.65591 |
| 668 | recB | 2.649036 | 0.741942 | 2 | 3543 | 0.12963 | 500.8112 | 74.97173 |
| 669 | evgS | 2.687167 | 0.749002 | 2 | 3594 | 0.125517 | 505.5761 | 75.57191 |
| 670 | metH | 2.754459 | 0.761053 | 2 | 3684 | 0.118585 | 513.7109 | 76.67328 |
| 671 | yfaL | 2.806049 | 0.76995 | 2 | 3753 | 0.113538 | 519.716 | 77.45395 |
| 672 | upstream y | 2.950351 | 0.793312 | 2 | 3946 | 0.100556 | 535.4859 | 79.6854 |
| 673 | rhsA | 3.090915 | 0.814026 | 2 | 4134 | 0.089362 | 549.4673 | 81.64448 |

| | | | | | | | | |
|---|---|---|---|---|---|---|---|---|
| 674 | gltB | 3.404942 | 0.853717 | 2 | 4554 | 0.068686 | 576.2592 | 85.4984 |
| 675 | ypjA | 3.425129 | 0.855984 | 2 | 4581 | 0.067534 | 577.7891 | 85.59838 |
| | | | | 1939 | | | | |
| | | | | 0.579498 | | | | |

| TP | FN | |
|---|---|---|
| 1.00 | 0.00 | 276 |
| 2.00 | 0.00 | 0.082487 |
| 3.00 | 0.00 | |
| 4.00 | 0.00 | |
| 5.00 | 0.00 | |
| 6.00 | 0.00 | |
| 7.00 | 0.00 | |
| 8.00 | 0.00 | |
| 9.00 | 0.00 | |
| 10.00 | 0.00 | |
| 11.00 | 0.00 | |
| 12.00 | 0.00 | |
| 13.00 | 0.00 | |
| 14.00 | 0.00 | |
| 14.99 | 0.01 | |
| 15.99 | 0.01 | |
| 16.99 | 0.01 | |
| 17.98 | 0.02 | |
| 18.98 | 0.02 | |
| 19.98 | 0.02 | |
| 20.97 | 0.03 | |
| 21.91 | 0.09 | |
| 22.91 | 0.09 | |
| 23.89 | 0.11 | |
| 24.87 | 0.13 | |
| 25.87 | 0.13 | |
| 26.87 | 0.13 | |
| 27.84 | 0.16 | |
| 28.78 | 0.22 | |
| 29.78 | 0.22 | |
| 30.66 | 0.34 | |
| 31.33 | 0.67 | |
| 32.31 | 0.69 | |
| 33.30 | 0.70 | |
| 34.21 | 0.79 | |
| 34.95 | 1.05 | |
| 35.32 | 1.68 | |
| 36.24 | 1.76 | |
| 37.14 | 1.86 | |
| 37.64 | 2.36 | |
| 38.16 | 2.84 | |
| 38.73 | 3.27 | |
| 39.00 | 4.00 | |
| 39.90 | 4.10 | |
| 40.58 | 4.42 | |
| 40.89 | 5.11 | |
| 41.84 | 5.16 | |
| 42.73 | 5.27 | |
| 43.26 | 5.74 | |

| | |
|---|---|
| 44.07 | 5.93 |
| 45.01 | 5.99 |
| 45.79 | 6.21 |
| 46.38 | 6.62 |
| 46.70 | 7.30 |
| 47.50 | 7.50 |
| 48.22 | 7.78 |
| 49.22 | 7.78 |
| 50.22 | 7.78 |
| 51.09 | 7.91 |
| 51.73 | 8.27 |
| 52.67 | 8.33 |
| 53.39 | 8.61 |
| 54.37 | 8.63 |
| 54.93 | 9.07 |
| 55.86 | 9.14 |
| 56.74 | 9.26 |
| 57.62 | 9.38 |
| 58.58 | 9.42 |
| 58.34 | 10.66 |
| 59.28 | 10.72 |
| 60.11 | 10.89 |
| 60.95 | 11.05 |
| 61.16 | 11.84 |
| 61.96 | 12.04 |
| 61.60 | 13.40 |
| 62.51 | 13.49 |
| 63.51 | 13.49 |
| 64.48 | 13.52 |
| 64.64 | 14.36 |
| 65.35 | 14.65 |
| 66.06 | 14.94 |
| 66.61 | 15.39 |
| 67.53 | 15.47 |
| 68.53 | 15.47 |
| 69.44 | 15.56 |
| 70.19 | 15.81 |
| 71.19 | 15.81 |
| 70.83 | 17.17 |
| 71.54 | 17.46 |
| 72.33 | 17.67 |
| 73.22 | 17.78 |
| 73.66 | 18.34 |
| 74.50 | 18.50 |
| 75.23 | 18.77 |
| 75.50 | 19.50 |
| 75.63 | 20.37 |
| 76.52 | 20.48 |
| 77.14 | 20.86 |
| 77.62 | 21.38 |
| 78.06 | 21.94 |
| 78.47 | 22.53 |

| | |
|---|---|
| 78.97 | 23.03 |
| 79.60 | 23.40 |
| 80.27 | 23.73 |
| 81.27 | 23.73 |
| 82.04 | 23.96 |
| 82.65 | 24.35 |
| 83.32 | 24.68 |
| 84.21 | 24.79 |
| 84.98 | 25.02 |
| 84.96 | 26.04 |
| 85.33 | 26.67 |
| 85.53 | 27.47 |
| 86.44 | 27.56 |
| 86.76 | 28.24 |
| 87.17 | 28.83 |
| 88.10 | 28.90 |
| 88.69 | 29.31 |
| 89.46 | 29.54 |
| 90.10 | 29.90 |
| 91.06 | 29.94 |
| 91.78 | 30.22 |
| 92.21 | 30.79 |
| 93.01 | 30.99 |
| 94.01 | 30.99 |
| 94.27 | 31.73 |
| 94.90 | 32.10 |
| 95.89 | 32.11 |
| 96.53 | 32.47 |
| 97.16 | 32.84 |
| 97.82 | 33.18 |
| 98.27 | 33.73 |
| 98.02 | 34.98 |
| 98.11 | 35.89 |
| 98.87 | 36.13 |
| 99.74 | 36.26 |
| 100.35 | 36.65 |
| 101.35 | 36.65 |
| 101.96 | 37.04 |
| 102.94 | 37.06 |
| 103.83 | 37.17 |
| 104.57 | 37.43 |
| 105.18 | 37.82 |
| 105.33 | 38.67 |
| 105.57 | 39.43 |
| 105.60 | 40.40 |
| 106.36 | 40.64 |
| 107.11 | 40.89 |
| 107.18 | 41.82 |
| 108.08 | 41.92 |
| 108.77 | 42.23 |
| 109.63 | 42.37 |
| 110.33 | 42.67 |

| | |
|---|---|
| 111.13 | 42.87 |
| 111.95 | 43.05 |
| 112.12 | 43.88 |
| 112.30 | 44.70 |
| 112.88 | 45.12 |
| 113.59 | 45.41 |
| 114.49 | 45.51 |
| 115.04 | 45.96 |
| 115.04 | 46.96 |
| 116.04 | 46.96 |
| 116.78 | 47.22 |
| 117.78 | 47.22 |
| 118.50 | 47.50 |
| 118.99 | 48.01 |
| 118.93 | 49.07 |
| 119.66 | 49.34 |
| 120.64 | 49.36 |
| 121.64 | 49.36 |
| 122.12 | 49.88 |
| 122.86 | 50.14 |
| 123.64 | 50.36 |
| 123.62 | 51.38 |
| 123.08 | 52.92 |
| 123.28 | 53.72 |
| 124.24 | 53.76 |
| 124.84 | 54.16 |
| 125.47 | 54.53 |
| 126.40 | 54.60 |
| 127.40 | 54.60 |
| 128.12 | 54.88 |
| 128.95 | 55.05 |
| 129.80 | 55.20 |
| 130.51 | 55.49 |
| 131.49 | 55.51 |
| 132.06 | 55.94 |
| 133.06 | 55.94 |
| 133.50 | 56.50 |
| 134.16 | 56.84 |
| 134.71 | 57.29 |
| 135.71 | 57.29 |
| 136.36 | 57.64 |
| 137.26 | 57.74 |
| 137.81 | 58.19 |
| 138.35 | 58.65 |
| 139.21 | 58.79 |
| 139.92 | 59.08 |
| 140.90 | 59.10 |
| 141.65 | 59.35 |
| 142.63 | 59.37 |
| 143.05 | 59.95 |
| 143.07 | 60.93 |
| 143.92 | 61.08 |

| | |
|---|---|
| 144.61 | 61.39 |
| 145.15 | 61.85 |
| 145.71 | 62.29 |
| 146.41 | 62.59 |
| 147.23 | 62.77 |
| 148.23 | 62.77 |
| 148.92 | 63.08 |
| 149.76 | 63.24 |
| 150.22 | 63.78 |
| 150.92 | 64.08 |
| 151.92 | 64.08 |
| 152.83 | 64.17 |
| 153.83 | 64.17 |
| 154.83 | 64.17 |
| 155.83 | 64.17 |
| 156.37 | 64.63 |
| 157.37 | 64.63 |
| 157.90 | 65.10 |
| 158.72 | 65.28 |
| 159.43 | 65.57 |
| 159.40 | 66.60 |
| 160.33 | 66.67 |
| 161.21 | 66.79 |
| 161.86 | 67.14 |
| 162.60 | 67.40 |
| 163.54 | 67.46 |
| 164.29 | 67.71 |
| 164.12 | 68.88 |
| 165.07 | 68.93 |
| 166.07 | 68.93 |
| 166.64 | 69.36 |
| 167.64 | 69.36 |
| 168.17 | 69.83 |
| 168.21 | 70.79 |
| 168.90 | 71.10 |
| 169.79 | 71.21 |
| 170.25 | 71.75 |
| 171.25 | 71.75 |
| 171.65 | 72.35 |
| 172.29 | 72.71 |
| 172.80 | 73.20 |
| 172.75 | 74.25 |
| 173.35 | 74.65 |
| 174.35 | 74.65 |
| 174.86 | 75.14 |
| 175.86 | 75.14 |
| 176.48 | 75.52 |
| 176.89 | 76.11 |
| 176.91 | 77.09 |
| 177.42 | 77.58 |
| 178.42 | 77.58 |
| 179.23 | 77.77 |

| | |
|---|---|
| 179.43 | 78.57 |
| 179.44 | 79.56 |
| 179.95 | 80.05 |
| 180.45 | 80.55 |
| 181.45 | 80.55 |
| 182.45 | 80.55 |
| 181.96 | 82.04 |
| 182.50 | 82.50 |
| 183.46 | 82.54 |
| 183.96 | 83.04 |
| 184.96 | 83.04 |
| 185.46 | 83.54 |
| 185.96 | 84.04 |
| 185.96 | 85.04 |
| 186.96 | 85.04 |
| 187.46 | 85.54 |
| 188.23 | 85.77 |
| 189.01 | 85.99 |
| 189.95 | 86.05 |
| 190.45 | 86.55 |
| 191.33 | 86.67 |
| 191.94 | 87.06 |
| 192.66 | 87.34 |
| 193.66 | 87.34 |
| 194.44 | 87.56 |
| 195.44 | 87.56 |
| 195.99 | 88.01 |
| 196.42 | 88.58 |
| 196.92 | 89.08 |
| 196.95 | 90.05 |
| 197.39 | 90.61 |
| 198.22 | 90.78 |
| 198.93 | 91.07 |
| 199.88 | 91.12 |
| 200.88 | 91.12 |
| 201.37 | 91.63 |
| 201.90 | 92.10 |
| 202.34 | 92.66 |
| 203.34 | 92.66 |
| 204.26 | 92.74 |
| 204.75 | 93.25 |
| 204.80 | 94.20 |
| 203.75 | 96.25 |
| 204.74 | 96.26 |
| 205.46 | 96.54 |
| 205.71 | 97.29 |
| 206.71 | 97.29 |
| 207.19 | 97.81 |
| 208.19 | 97.81 |
| 208.15 | 98.85 |
| 208.11 | 99.89 |
| 209.11 | 99.89 |

| | |
|---|---|
| 209.59 | 100.41 |
| 210.18 | 100.82 |
| 210.72 | 101.28 |
| 211.48 | 101.52 |
| 212.02 | 101.98 |
| 211.97 | 103.03 |
| 212.45 | 103.55 |
| 212.92 | 104.08 |
| 213.40 | 104.60 |
| 212.82 | 106.18 |
| 213.11 | 106.89 |
| 213.23 | 107.77 |
| 214.23 | 107.77 |
| 215.23 | 107.77 |
| 215.31 | 108.69 |
| 216.18 | 108.82 |
| 217.18 | 108.82 |
| 218.18 | 108.82 |
| 218.65 | 109.35 |
| 219.65 | 109.35 |
| 220.12 | 109.88 |
| 220.58 | 110.42 |
| 221.58 | 110.42 |
| 222.58 | 110.42 |
| 223.58 | 110.42 |
| 224.58 | 110.42 |
| 225.58 | 110.42 |
| 225.76 | 111.24 |
| 226.76 | 111.24 |
| 227.52 | 111.48 |
| 227.46 | 112.54 |
| 228.12 | 112.88 |
| 228.92 | 113.08 |
| 229.39 | 113.61 |
| 229.86 | 114.14 |
| 229.25 | 115.75 |
| 229.72 | 116.28 |
| 230.72 | 116.28 |
| 230.88 | 117.12 |
| 231.64 | 117.36 |
| 232.64 | 117.36 |
| 233.64 | 117.36 |
| 234.64 | 117.36 |
| 234.57 | 118.43 |
| 235.03 | 118.97 |
| 234.91 | 120.09 |
| 235.42 | 120.58 |
| 235.88 | 121.12 |
| 236.88 | 121.12 |
| 235.72 | 123.28 |
| 236.18 | 123.82 |
| 237.18 | 123.82 |

| | |
|---|---|
| 238.16 | 123.84 |
| 238.63 | 124.37 |
| 239.63 | 124.37 |
| 240.09 | 124.91 |
| 238.92 | 127.08 |
| 239.38 | 127.62 |
| 240.38 | 127.62 |
| 239.75 | 129.25 |
| 240.75 | 129.25 |
| 240.66 | 130.34 |
| 240.97 | 131.03 |
| 240.82 | 132.18 |
| 241.48 | 132.52 |
| 242.48 | 132.52 |
| 242.38 | 133.62 |
| 242.91 | 134.09 |
| 243.84 | 134.16 |
| 244.66 | 134.34 |
| 245.52 | 134.48 |
| 246.29 | 134.71 |
| 247.29 | 134.71 |
| 247.75 | 135.25 |
| 248.74 | 135.26 |
| 249.74 | 135.26 |
| 250.68 | 135.32 |
| 249.00 | 138.00 |
| 248.90 | 139.10 |
| 249.90 | 139.10 |
| 250.36 | 139.64 |
| 251.36 | 139.64 |
| 251.26 | 140.74 |
| 252.26 | 140.74 |
| 253.26 | 140.74 |
| 254.26 | 140.74 |
| 254.71 | 141.29 |
| 255.16 | 141.84 |
| 255.61 | 142.39 |
| 255.50 | 143.50 |
| 255.40 | 144.60 |
| 255.30 | 145.70 |
| 255.20 | 146.80 |
| 256.11 | 146.89 |
| 256.09 | 147.91 |
| 257.09 | 147.91 |
| 256.99 | 149.01 |
| 256.57 | 150.43 |
| 256.77 | 151.23 |
| 255.56 | 153.44 |
| 256.56 | 153.44 |
| 257.56 | 153.44 |
| 258.00 | 154.00 |
| 259.00 | 154.00 |

| | |
|---|---|
| 258.90 | 155.10 |
| 259.90 | 155.10 |
| 259.79 | 156.21 |
| 260.23 | 156.77 |
| 260.68 | 157.32 |
| 261.68 | 157.32 |
| 262.12 | 157.88 |
| 263.12 | 157.88 |
| 264.12 | 157.88 |
| 264.57 | 158.43 |
| 265.01 | 158.99 |
| 265.07 | 159.93 |
| 265.68 | 160.32 |
| 266.35 | 160.65 |
| 267.35 | 160.65 |
| 267.88 | 161.12 |
| 268.79 | 161.21 |
| 269.79 | 161.21 |
| 270.48 | 161.52 |
| 271.24 | 161.76 |
| 272.24 | 161.76 |
| 272.68 | 162.32 |
| 272.88 | 163.12 |
| 273.01 | 163.99 |
| 274.01 | 163.99 |
| 274.29 | 164.71 |
| 274.90 | 165.10 |
| 275.90 | 165.10 |
| 275.79 | 166.21 |
| 276.23 | 166.77 |
| 276.67 | 167.33 |
| 277.67 | 167.33 |
| 276.45 | 169.55 |
| 275.22 | 171.78 |
| 276.22 | 171.78 |
| 277.11 | 171.89 |
| 277.66 | 172.34 |
| 276.99 | 174.01 |
| 277.44 | 174.56 |
| 277.88 | 175.12 |
| 277.77 | 176.23 |
| 278.21 | 176.79 |
| 278.65 | 177.35 |
| 279.10 | 177.90 |
| 279.54 | 178.46 |
| 280.54 | 178.46 |
| 280.98 | 179.02 |
| 281.42 | 179.58 |
| 280.75 | 181.25 |
| 280.64 | 182.36 |
| 281.08 | 182.92 |
| 282.08 | 182.92 |

| | |
|---|---|
| 282.56 | 183.44 |
| 282.97 | 184.03 |
| 283.93 | 184.07 |
| 284.41 | 184.59 |
| 285.41 | 184.59 |
| 285.86 | 185.14 |
| 286.86 | 185.14 |
| 287.71 | 185.29 |
| 288.30 | 185.70 |
| 288.74 | 186.26 |
| 289.19 | 186.81 |
| 290.19 | 186.81 |
| 291.09 | 186.91 |
| 291.07 | 187.93 |
| 290.96 | 189.04 |
| 291.96 | 189.04 |
| 292.40 | 189.60 |
| 292.85 | 190.15 |
| 293.85 | 190.15 |
| 294.29 | 190.71 |
| 294.73 | 191.27 |
| 295.73 | 191.27 |
| 296.18 | 191.82 |
| 297.18 | 191.82 |
| 297.62 | 192.38 |
| 297.51 | 193.49 |
| 298.16 | 193.84 |
| 298.95 | 194.05 |
| 299.40 | 194.60 |
| 300.40 | 194.60 |
| 300.84 | 195.16 |
| 301.58 | 195.42 |
| 302.29 | 195.71 |
| 302.73 | 196.27 |
| 302.06 | 197.94 |
| 301.40 | 199.60 |
| 301.29 | 200.71 |
| 301.73 | 201.27 |
| 302.73 | 201.27 |
| 303.18 | 201.82 |
| 303.07 | 202.93 |
| 299.50 | 207.50 |
| 300.26 | 207.74 |
| 301.26 | 207.74 |
| 302.08 | 207.92 |
| 303.08 | 207.92 |
| 303.85 | 208.15 |
| 304.53 | 208.47 |
| 304.98 | 209.02 |
| 305.42 | 209.58 |
| 305.87 | 210.13 |
| 306.40 | 210.60 |

| | |
|---|---|
| 306.77 | 211.23 |
| 307.21 | 211.79 |
| 307.11 | 212.89 |
| 307.56 | 213.44 |
| 306.35 | 215.65 |
| 306.80 | 216.20 |
| 307.25 | 216.75 |
| 308.25 | 216.75 |
| 308.84 | 217.16 |
| 308.60 | 218.40 |
| 309.05 | 218.95 |
| 307.85 | 221.15 |
| 306.62 | 223.38 |
| 307.10 | 223.90 |
| 307.01 | 224.99 |
| 307.46 | 225.54 |
| 305.72 | 228.28 |
| 306.17 | 228.83 |
| 307.17 | 228.83 |
| 307.63 | 229.37 |
| 306.99 | 231.01 |
| 304.71 | 234.29 |
| 305.17 | 234.83 |
| 305.63 | 235.37 |
| 306.08 | 235.92 |
| 307.08 | 235.92 |
| 308.08 | 235.92 |
| 307.99 | 237.01 |
| 308.99 | 237.01 |
| 308.91 | 238.09 |
| 309.91 | 238.09 |
| 309.82 | 239.18 |
| 310.28 | 239.72 |
| 310.10 | 240.90 |
| 310.65 | 241.35 |
| 310.57 | 242.43 |
| 311.57 | 242.43 |
| 312.57 | 242.43 |
| 313.03 | 242.97 |
| 312.95 | 244.05 |
| 313.02 | 244.98 |
| 313.87 | 245.13 |
| 314.61 | 245.39 |
| 315.33 | 245.67 |
| 312.56 | 249.44 |
| 313.02 | 249.98 |
| 312.95 | 251.05 |
| 311.80 | 253.20 |
| 312.27 | 253.73 |
| 310.06 | 256.94 |
| 311.06 | 256.94 |
| 310.20 | 258.80 |

| | |
|---|---|
| 310.93 | 259.07 |
| 309.80 | 261.20 |
| 309.21 | 262.79 |
| 307.03 | 265.97 |
| 306.97 | 267.03 |
| 306.39 | 268.61 |
| 307.39 | 268.61 |
| 306.28 | 270.72 |
| 305.18 | 272.82 |
| 303.57 | 275.43 |
| 303.52 | 276.48 |
| 303.27 | 277.73 |
| 304.27 | 277.73 |
| 304.47 | 278.53 |
| 305.44 | 278.56 |
| 302.48 | 282.52 |
| 297.60 | 288.40 |
| 297.58 | 289.42 |
| 297.04 | 290.96 |
| 297.02 | 291.98 |
| 296.48 | 293.52 |
| 295.95 | 295.05 |
| 295.94 | 296.06 |
| 293.17 | 299.83 |
| 294.09 | 299.91 |
| 294.37 | 300.63 |
| 294.87 | 301.13 |
| 295.87 | 301.13 |
| 292.34 | 305.66 |
| 293.24 | 305.76 |
| 292.34 | 307.66 |
| 290.57 | 310.43 |
| 291.57 | 310.43 |
| 289.37 | 313.63 |
| 289.38 | 314.62 |
| 290.38 | 314.62 |
| 291.38 | 314.62 |
| 291.40 | 315.60 |
| 290.26 | 317.74 |
| 288.97 | 320.03 |
| 288.02 | 321.98 |
| 286.58 | 324.42 |
| 286.23 | 325.77 |
| 285.67 | 327.33 |
| 283.78 | 330.22 |
| 280.95 | 334.05 |
| 280.99 | 335.01 |
| 277.25 | 339.75 |
| 277.78 | 340.22 |
| 276.42 | 342.58 |
| 277.15 | 342.85 |
| 277.48 | 343.52 |

| | |
|---|---|
| 278.02 | 343.98 |
| 277.15 | 345.85 |
| 277.52 | 346.48 |
| 273.58 | 351.42 |
| 271.82 | 354.18 |
| 268.71 | 358.29 |
| 267.89 | 360.11 |
| 268.44 | 360.56 |
| 266.28 | 363.72 |
| 267.28 | 363.72 |
| 266.94 | 365.06 |
| 265.26 | 367.74 |
| 265.37 | 368.63 |
| 264.15 | 370.85 |
| 261.63 | 374.37 |
| 261.94 | 375.06 |
| 261.19 | 376.81 |
| 262.00 | 377.00 |
| 261.26 | 378.74 |
| 258.38 | 382.62 |
| 258.95 | 383.05 |
| 257.81 | 385.19 |
| 257.10 | 386.90 |
| 256.83 | 388.17 |
| 254.87 | 391.13 |
| 255.78 | 391.22 |
| 252.27 | 395.73 |
| 252.86 | 396.14 |
| 252.20 | 397.80 |
| 250.32 | 400.68 |
| 248.05 | 403.95 |
| 247.84 | 405.16 |
| 246.01 | 407.99 |
| 244.61 | 410.39 |
| 244.88 | 411.12 |
| 244.37 | 412.63 |
| 237.39 | 420.61 |
| 221.85 | 437.15 |
| 213.04 | 446.96 |
| 211.57 | 449.43 |
| 202.21 | 459.79 |
| 198.84 | 464.16 |
| 195.87 | 468.13 |
| 186.60 | 478.40 |
| 174.47 | 491.53 |
| 169.05 | 497.95 |
| 167.19 | 500.81 |
| 163.42 | 505.58 |
| 156.29 | 513.71 |
| 151.28 | 519.72 |
| 136.51 | 535.49 |
| 123.53 | 549.47 |

| | |
|---|---|
| 97.74 | 576.26 |
| 97.21 | 577.79 |
