## Supplementary Table S3 for "Evolutionary dynamics of *de novo* mutations and mutant lineages arising in a simple, constant environment"

**Table S3.** Population allele frequencies for frequently mutated genes

| | | Alleles in each population | | | | Cumulative frequency at end of experiment | | |
|---|---|---|---|---|---|---|---|---|
| **Gene** | **Observed Mutations** | **1** | **2** | **3** | **Total unique alleles** | **1** | **2** | **3** |
| ***galS*** | 38 | 4 | 24 | 6 | 30 | 0 | 0 | 0.06 |
| ***hfq*** | 24 | 10 | 11 | 3 | 14 | 0.91 | 0.50 | 0.86 |
| ***pgi*** | 35 | 10 | 6 | 19 | 24 | 0.02 | 0.49 | 0.60 |
| ***opgH*** | 31 | 9 | 11 | 11 | 29 | 0.95 | 0.69 | 0.68 |
| ***malT*** | 30 | 1 | 18 | 9 | 19 | 0 | 0.51 | 0.84 |
| ***malK*** | 22 | 2 | 13 | 7 | 14 | 0.99 | 0.21 | 0.03 |
| **upstream *mglB*** | 7 | 2 | 3 | 2 | 4 | 1.00 | 0.94 | 0.94 |
| ***rho*** | 11 | 4 | 3 | 4 | 9 | 0.99 | 0.02 | 0.95 |
| **upstream *dnaG*** | 5 | 2 | 3 | 0 | 5 | 1.00 | 0.23 | 0 |
| ***fimH*** | 9 | 4 | 3 | 2 | 5 | 0 | 0.16 | 0.09 |
