## Supplementary Table S4 for "Evolutionary dynamics of *de novo* mutations and mutant lineages arising in a simple, constant environment"

**Table S4.** Identical mutations arise within and among replicate evolution experiments.

| Chemostat | Nucleotide position | Ancestral | Derived | Gene/Region | Amino acid change |
|---|---|---|---|---|---|
| all 3 | 55258 | C | A | lptD | Glu618* |
| | 2238647 | C | A | upstream mglB* | none |
| | 2238671 | G | T | galS | Asn340Lys |
| | 2239254 | C | A | galS | Arg146Leu |
| | 2790401 | G | T | gabD | Gln369His |
| | 3481115 | G | T | yheS | Cys602Phe |
| | 4232061 | G | T | pgi | Arg94Leu |
| | 4232255 | G | T | pgi | Asp159Tyr |
| | 4232262 | G | T | pgi | Gly161Val |
| | 4245693 | C | A | malK | Ala296Asp |
| | 4398360 | G | T | hfq | Arg17Leu |
| | 4398395 | G | T | hfq | Gly29Cys |
| | 4547110 | G | T | fimH | Gly94Trp |
| 1 and 2 | 294752 | C | A | yagN | Ala18Ser |
| | 505592 | C | A | ushA | none |
| | 1108798 | G | T | opgG | Glu81* |
| | 1111307 | C | A | opgH | Gln408Lys |
| | 1297594 | C | A | upstream adhE | N/A |
| | 3438088 | G | T | rpoA | Pro322Thr |
| | 3964698 | C | A | rho | Arg87Ser |
| | 4232851 | C | A | pgi | Asn357Lys |
| | 4245697 | C | A | malK | Asp297Glu |
| | 4367603 | G | T | yjeH | Ala278Asp |
| | 4398388 | G | T | hfq | Leu26Phe |
| | 4398403 | G | T | hfq | Lys31Asn |
| | 4398404 | C | A | hfq | Leu32Met |
| | 4398494 | G | T | hfq | Val62Phe |
| | 4398501 | C | A | hfq | Pro64Gln |
| | 4398665 | C | A | downstream hfq/upstream hflX | N/A |
| | 4547014 | C | A | fimH | Gln62Lys |
| | 4629113 | G | T | slt | Ala120Ser |
| 1 and 3 | 501321 | G | T | ybaL | Pro381Gln |
| | 3368984 | C | A | nanE | none |
| | 3965316 | G | T | rho | Ala293Ser |
| | 4185625 | T | C | rpoC | none |
| | 4232066 | G | T | pgi | Val96Leu |
| | 4232436 | C | A | pgi | Ala219Asp |

| | | | | | |
|---|---|---|---|---|---|
| | 4232577 | G | T | pgi | Trp266Leu |
| | 4547002 | G | T | fimH | Asp58Tyr |
| **2 and 3** | 1111193 | G | T | opgH | Gly370Trp |
| | 1338741 | G | T | yciM | Ala54Ser |
| | 1905184 | C | A | upstream rlmA | |
| | 1905560 | C | A | yobF | Arg19Leu |
| | 2012958 | G | T | fliG | Glu19* |
| | 2238648 | G | T | upstream mglB | |
| | 2864679 | G | T | rpoS | Arg299Ser |
| | 3342432 | C | A | lptB | Pro156Gln |
| | 3421312 | A | C | downstream yhdZ/upstream rrfF | |
| | 3551116 | C | A | malT | Pro4Thr |
| | 3551131 | C | A | malT | Arg9Ser |
| | 3551264 | C | A | malT | Ala53Glu |
| | 3551285 | G | T | malT | Gly60Val |
| | 3551833 | C | A | malT | Leu243Met |
| | 3552039 | G | T | malT | Met311Ile |
| | 3552062 | G | T | malT | Cys319Phe |
| | 3552180 | C | A | malT | Ser358Arg |
| | 3553006 | C | A | malT | Arg634Ser |
| | 4105641 | G | T | pfkA | Gly23Trp |
| | 4233075 | C | A | pgi | Ala432Glu |
| | 4244957 | G | T | malK | Gly51Trp |
| | 4245480 | C | A | malK | Pro225Gln |
| | 4245498 | C | A | malK | Ala231Asp |
| | 4245563 | C | A | malK | Gln253Lys |
| | 4245821 | G | T | malK | Glu339* |
| | 4398489 | C | A | hfq | Ser60Tyr |

- present at 1.4% at gen 0 in chemostat 1, but drops to undetectable at g 50.
