## Supplementary Table S5 for "Evolutionary dynamics of *de novo* mutations and mutant lineages arising in a simple, constant environment"

**Table S5.** Fixed alleles among replicate populations ("fixed" defined as >98% at any time point between generation 50 and 500).

| | | | | | | **Generation** | | | | | | | | | | |
|---|---|---|---|---|---|---|---|---|---|---|---|---|---|---|---|---|
| **Chemostat** | **Nucleotide Position** | **Ancestral** | **Derived** | **Region** | **Amino acid change** | **0** | **50** | **100** | **150** | **200** | **250** | **300** | **350** | **400** | **450** | **500** |
| **1** | 438,942 | C | A | *dxs* | Ala154Ser | 0 | 0 | 93 | 12 | 88 | 94 | 94 | 78 | 100 | 100 | 100 |
| **1** | 546,837 | G | T | *fdrA* | Ala312Ser | 0 | 0 | 93 | 11 | 87 | 96 | 93 | 82 | 98 | 100 | 99 |
| **1** | 922,714 | C | A | *clpA* | | 0 | 0 | 93 | 12 | 85 | 95 | 93 | 78 | 99 | 99 | 99 |
| **1** | 1,107,984 | G | T | *opgC* | Gln61Lys | 0 | 0 | 92 | 14 | 85 | 95 | 93 | 79 | 100 | 100 | 100 |
| **1** | 1,361,341 | C | A | *puuC* | Pro192Gln | 0 | 0 | 93 | 13 | 86 | 95 | 93 | 81 | 99 | 100 | 100 |
| **1** | 2,019,771 | G | T | *fliO* | Val82Phe | 0 | 0 | 92 | 11 | 87 | 95 | 93 | 77 | 99 | 99 | 100 |
| **1** | 2,119,008 | C | A | *wcaJ* | Gly191Cys | 0 | 0 | 92 | 15 | 85 | 95 | 93 | 79 | 100 | 100 | 100 |
| **1** | 2,238,647 | C | A | upstream *mglB* | | 1 | 0 | 95 | 12 | 88 | 95 | 92 | 77 | 99 | 100 | 100 |
| **1** | 2,408,047 | G | T | *yfbS* | Ala443Asp | 0 | 0 | 94 | 11 | 87 | 96 | 95 | 81 | 100 | 100 | 100 |
| **1** | 2,500,493 | G | T | *fryA* | Ala672Glu | 0 | 0 | 94 | 12 | 86 | 95 | 93 | 79 | 99 | 100 | 100 |
| **1** | 2,775,348 | G | T | *yfjZ* | Arg71Leu | 0 | 0 | 94 | 13 | 86 | 95 | 90 | 79 | 100 | 100 | 100 |
| **1** | 3,209,081 | G | T | upstream *dnaG* | | 0 | 92 | 98 | 93 | 95 | 99 | 99 | 99 | 99 | 100 | 100 |
| **1** | 3,965,271 | G | T | *rho* | Val278Phe | 0 | 0 | 93 | 13 | 87 | 93 | 91 | 82 | 98 | 99 | 99 |
| **1** | 4,205,962 | G | T | | | 0 | 0 | 93 | 10 | 89 | 95 | 91 | 82 | 99 | 100 | 100 |
| **1** | 4,245,693 | C | A | *malK* | Ala296Asp | 0 | 0 | 94 | 12 | 89 | 95 | 93 | 83 | 99 | 100 | 99 |
| **3** | 82,6641 | C | A | *ybhF* | Glu522* | 9 | 79 | 91 | 97 | 98 | 91 | 89 | | | | |
| **3** | 2,238,647 | C | A | upstream *mglB* | | 0 | 81 | 96 | 99 | 97 | 94 | 94 | | | | |
