## Supplementary Data File S3 for "Evolutionary dynamics of *de novo* mutations and mutant lineages arising in a simple, constant environment"

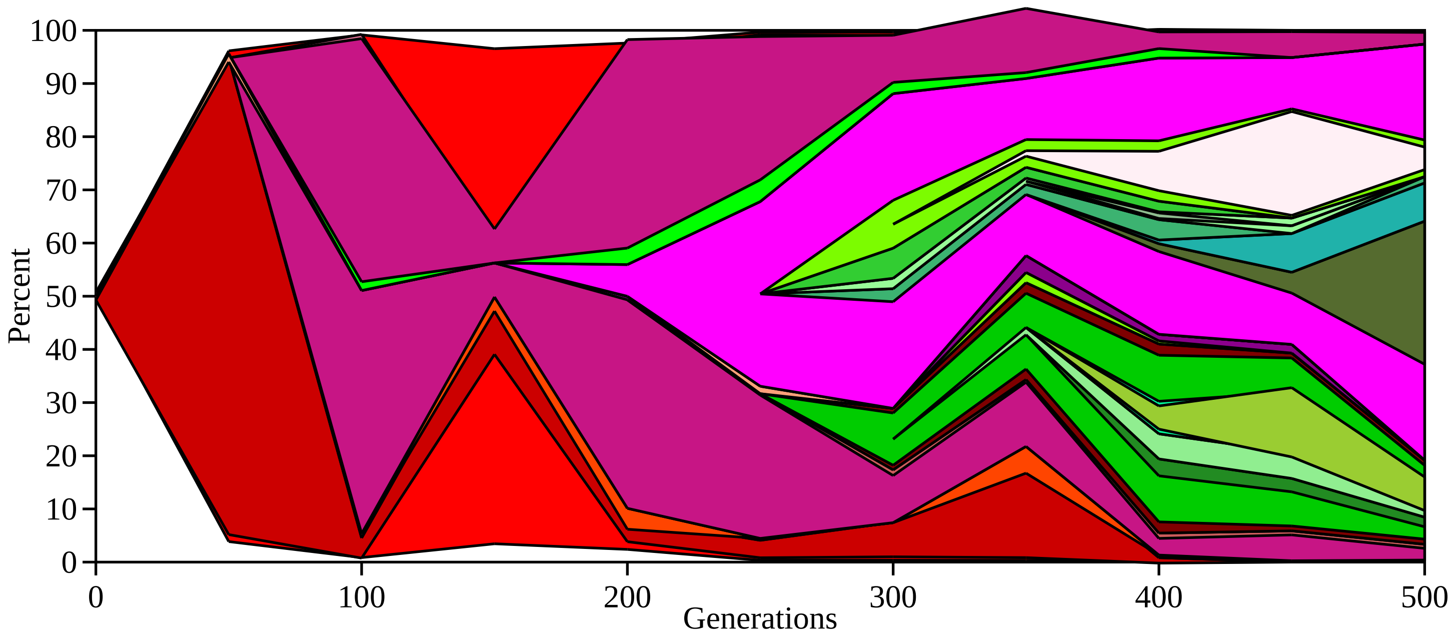
100
90
80
70
60
50
40
30
20
10
0
Percent
0
100
200
300
400
500
Generations

Lineages for downstream hfq

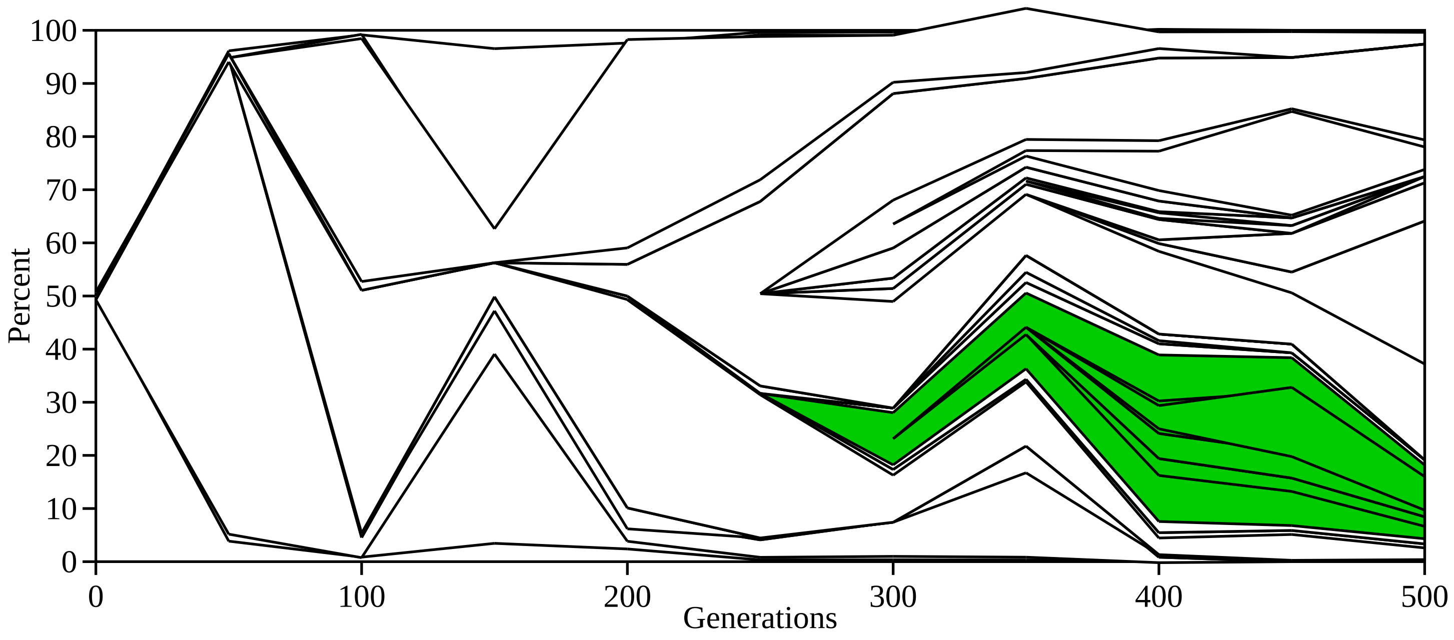

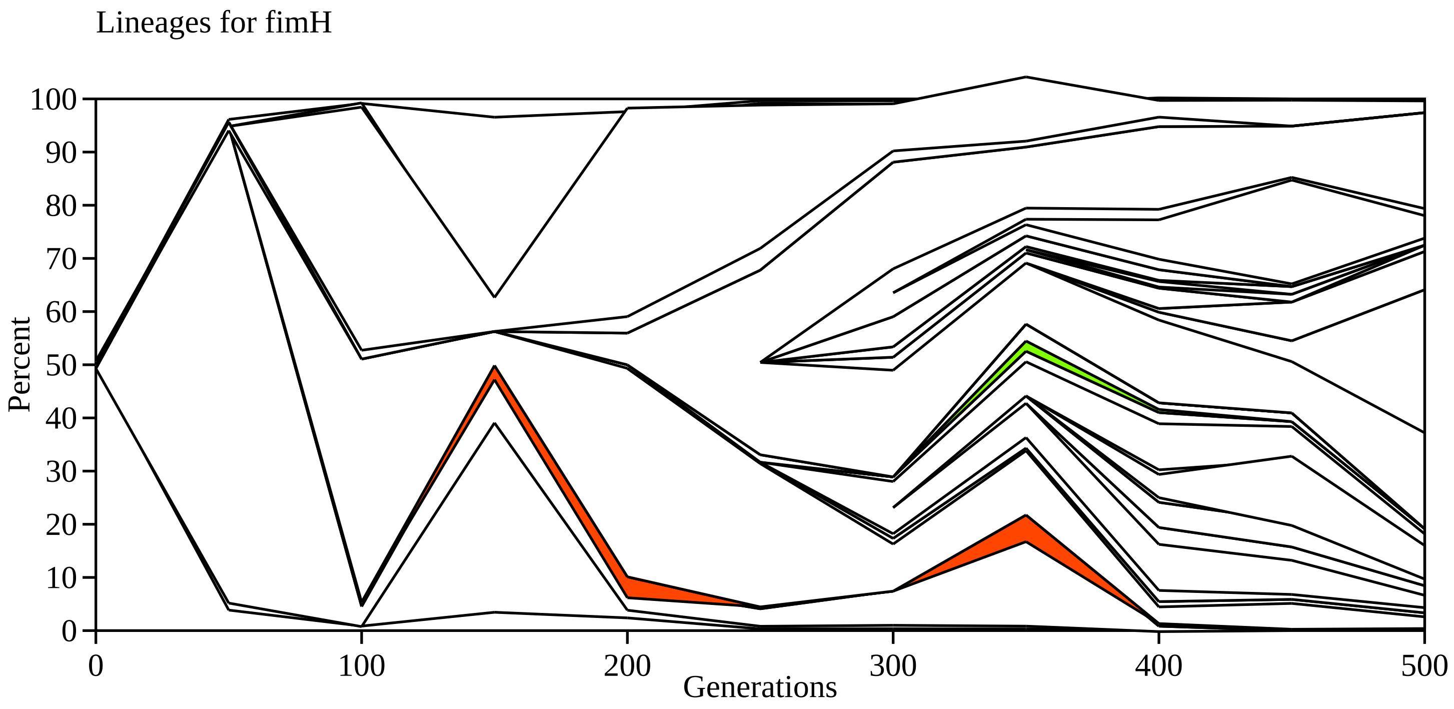

Lineages for fimH
100
90
80
70
60
50
40
30
20
10
0
Percent
0
100
200
300
400
500
Generations

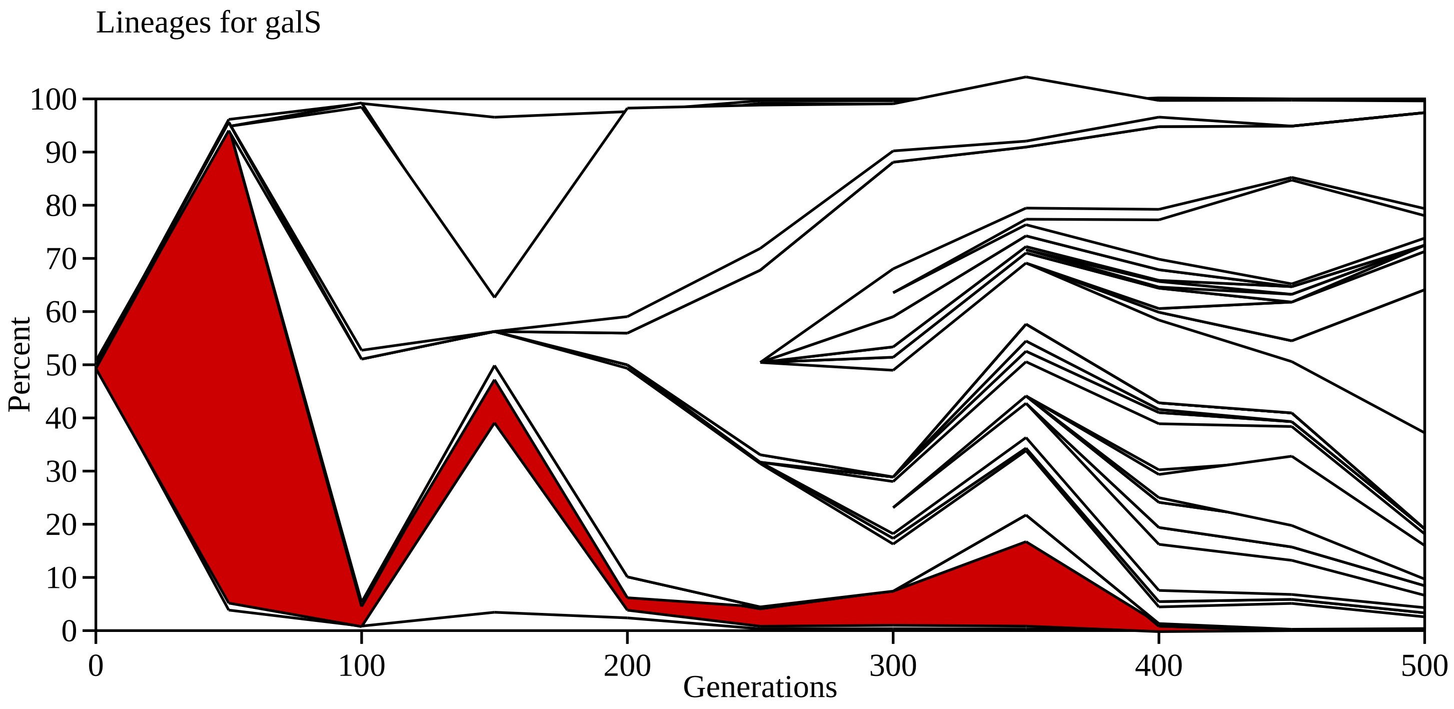
Lineages for galS
100
90
80
70
60
50
40
30
20
10
0
Percent
0
100
200
300
400
500
Generations

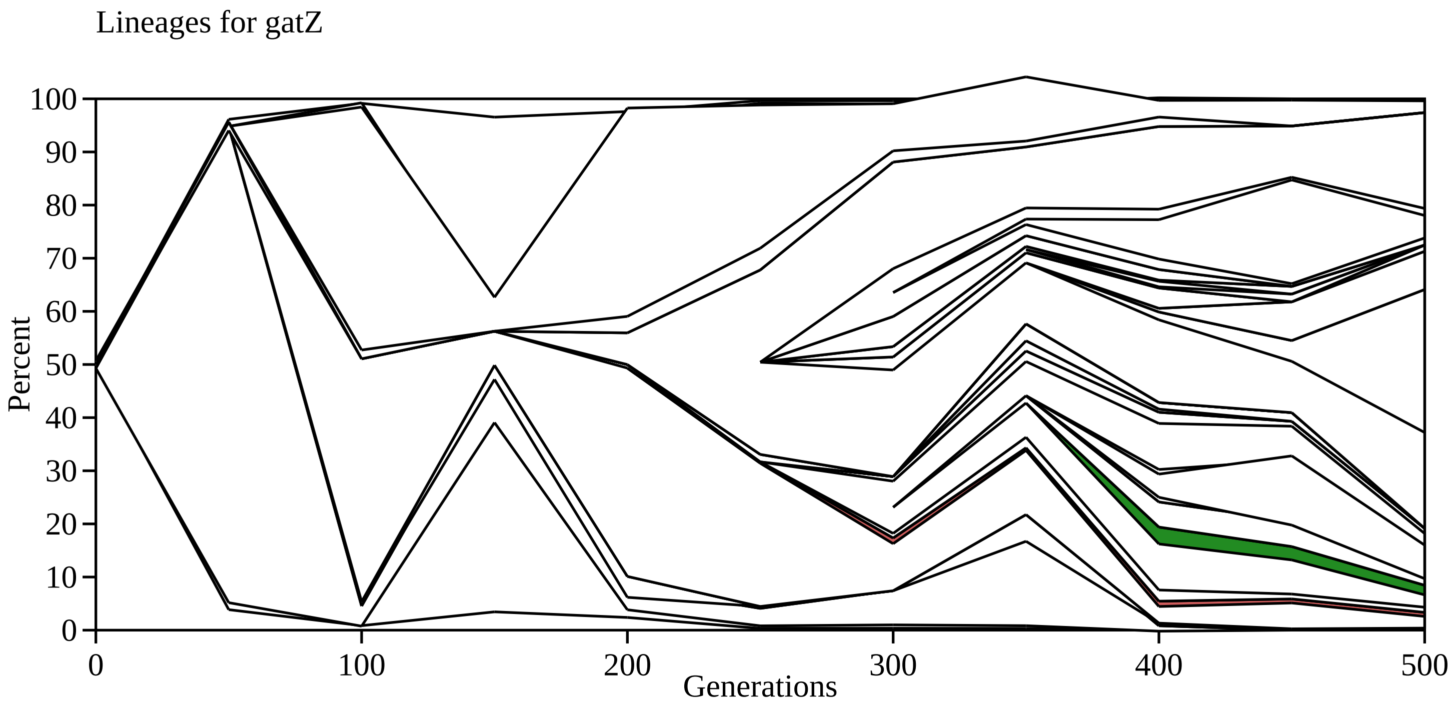

Lineages for gatZ
100
90
80
70
60
50
40
30
20
10
0
Percent
0
100
200
300
400
500
Generations

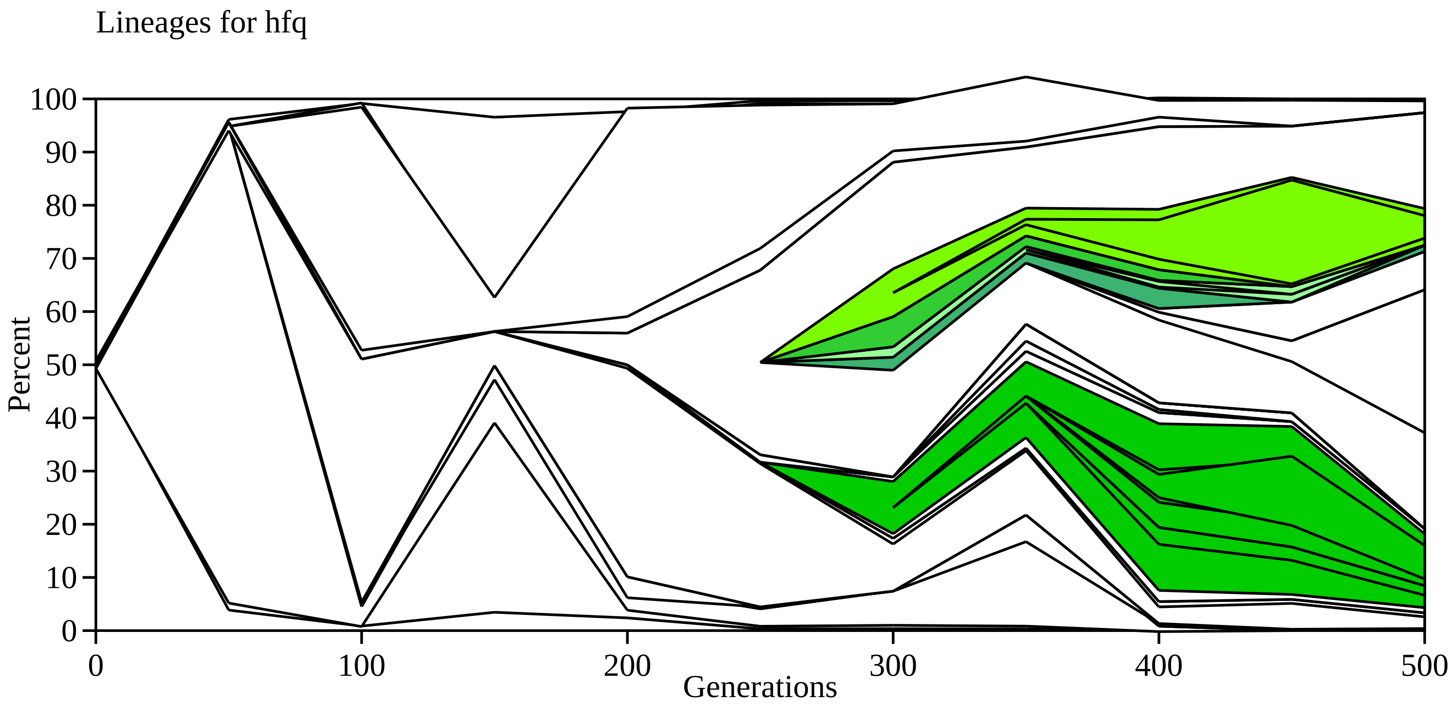

Lineages for hfq
Percent
0
10
20
30
40
50
60
70
80
90
100
0
100
200
300
400
500
Generations

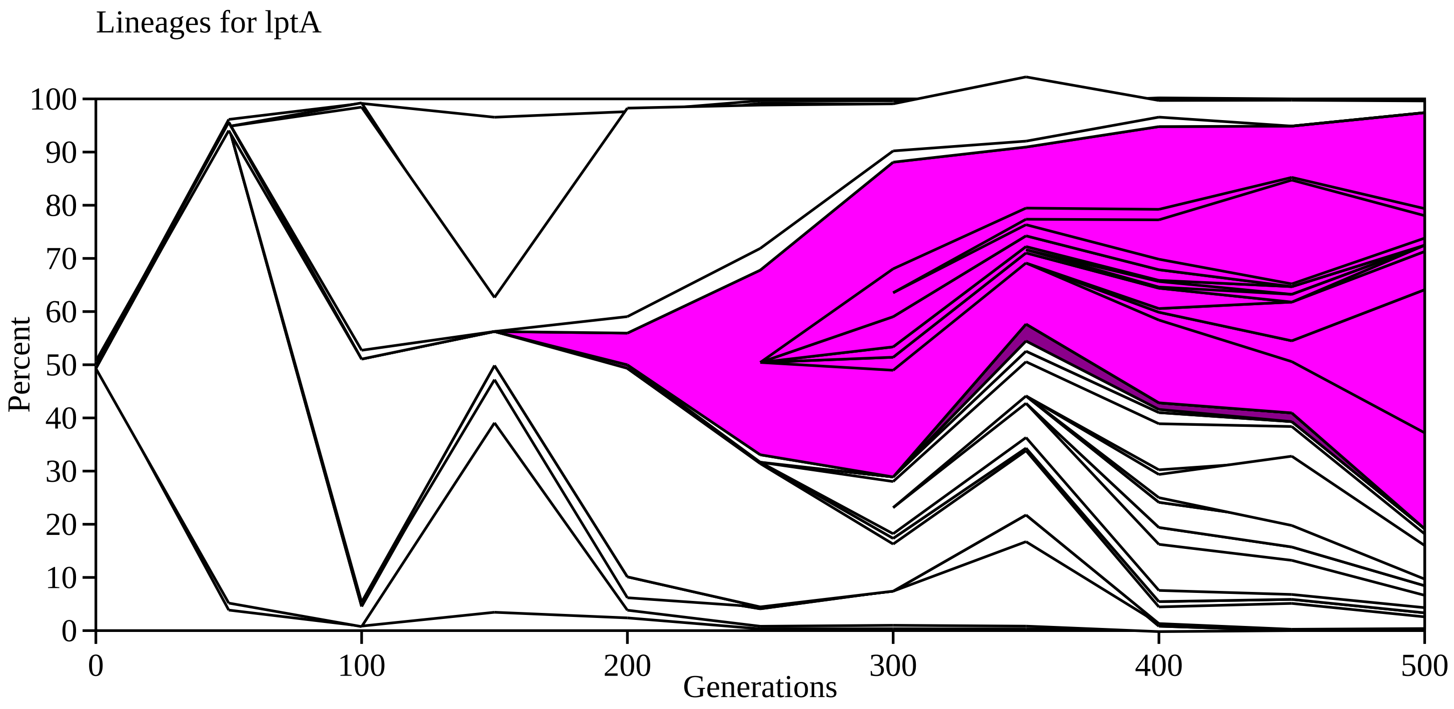

Lineages for lptA
Percent
Generations
100
90
80
70
60
50
40
30
20
10
0
0
100
200
300
400
500

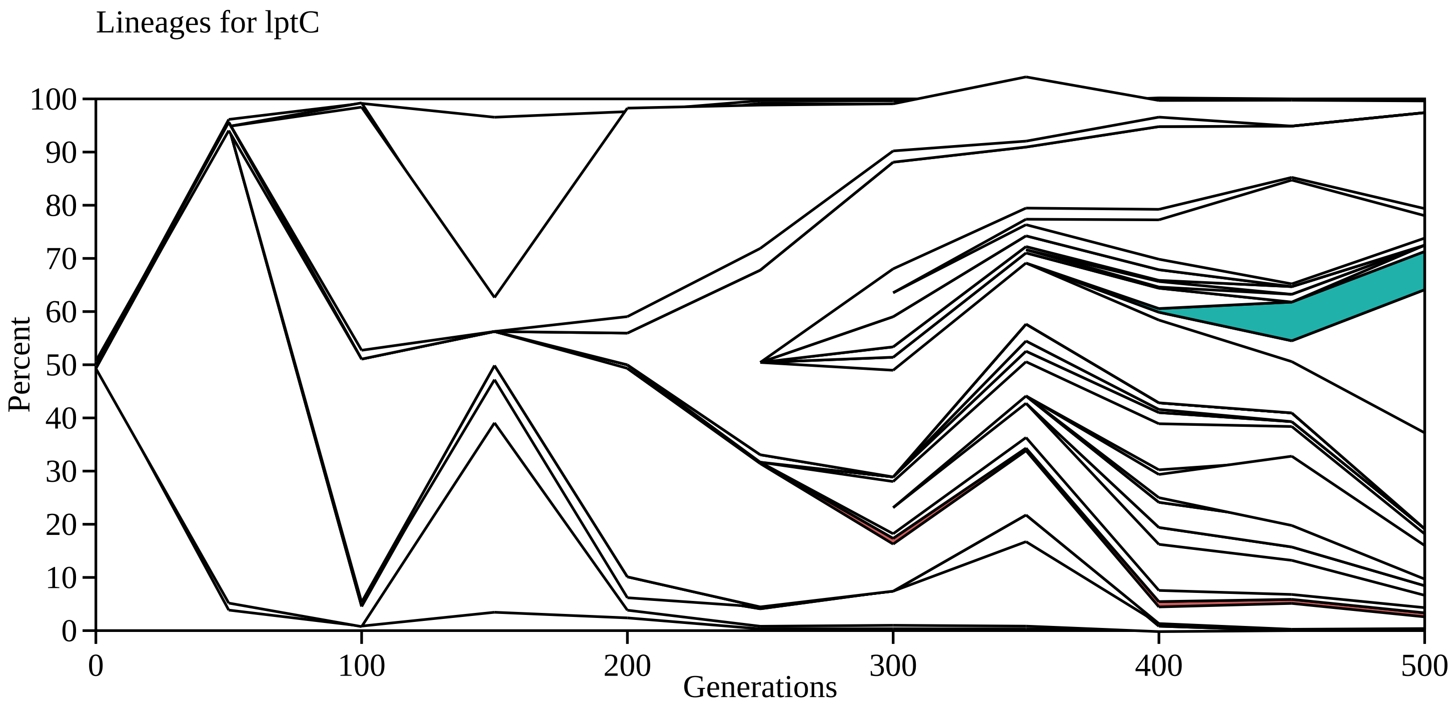

Lineages for lptC
100
90
80
70
60
50
40
30
20
10
0
Percent
0
100
200
300
400
500
Generations

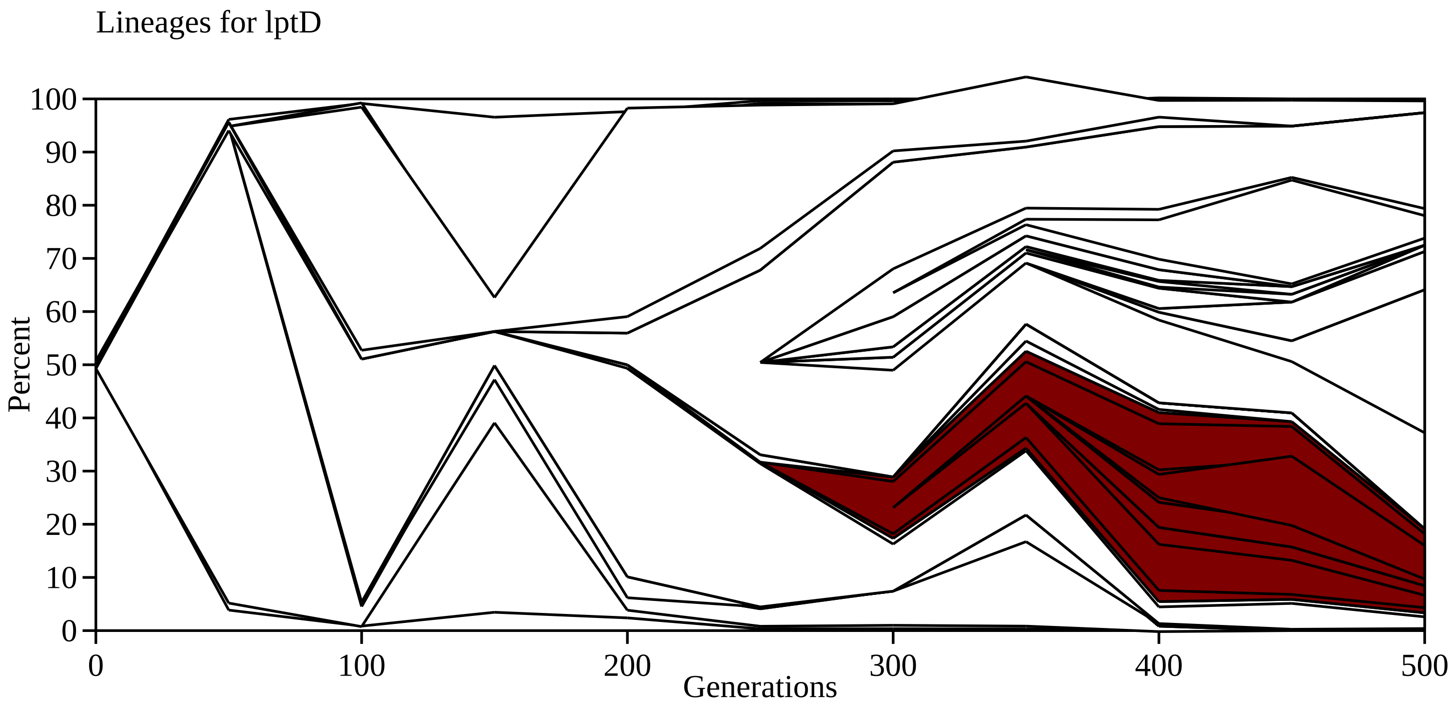

Lineages for lptD
100
90
80
70
60
50
40
30
20
10
0
Percent
0
100
200
300
400
500
Generations

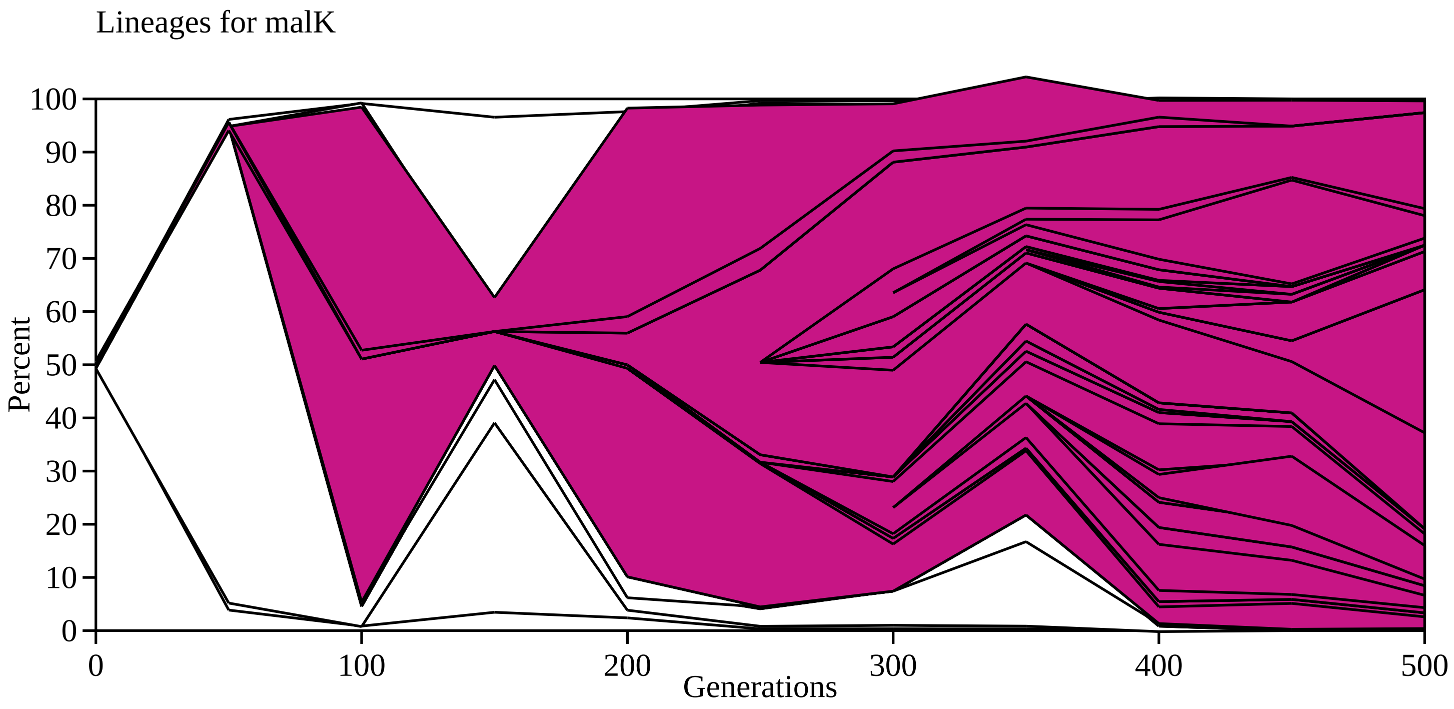

Lineages for malK
Percent
100
90
80
70
60
50
40
30
20
10
0
0
100
200
300
400
500
Generations

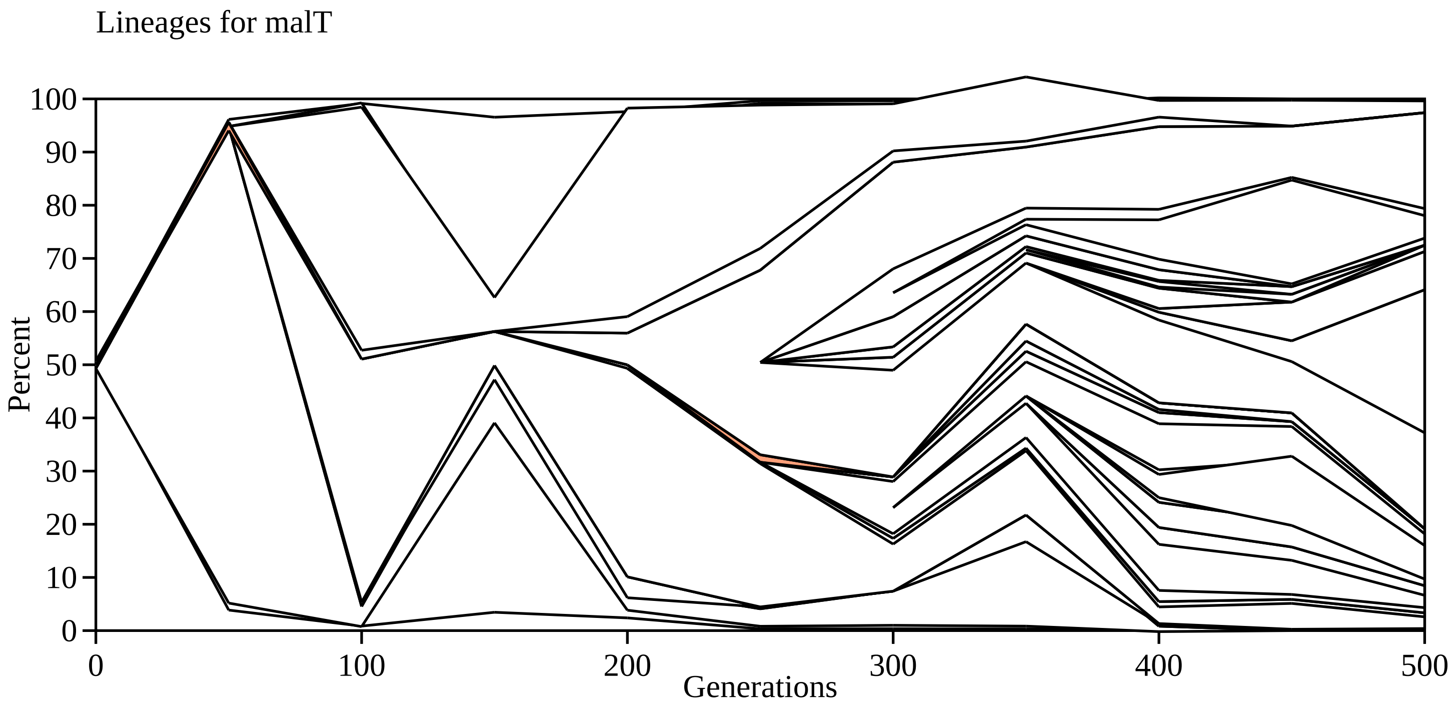

Lineages for malT
Percent
100
90
80
70
60
50
40
30
20
10
0
0
100
200
300
400
500
Generations

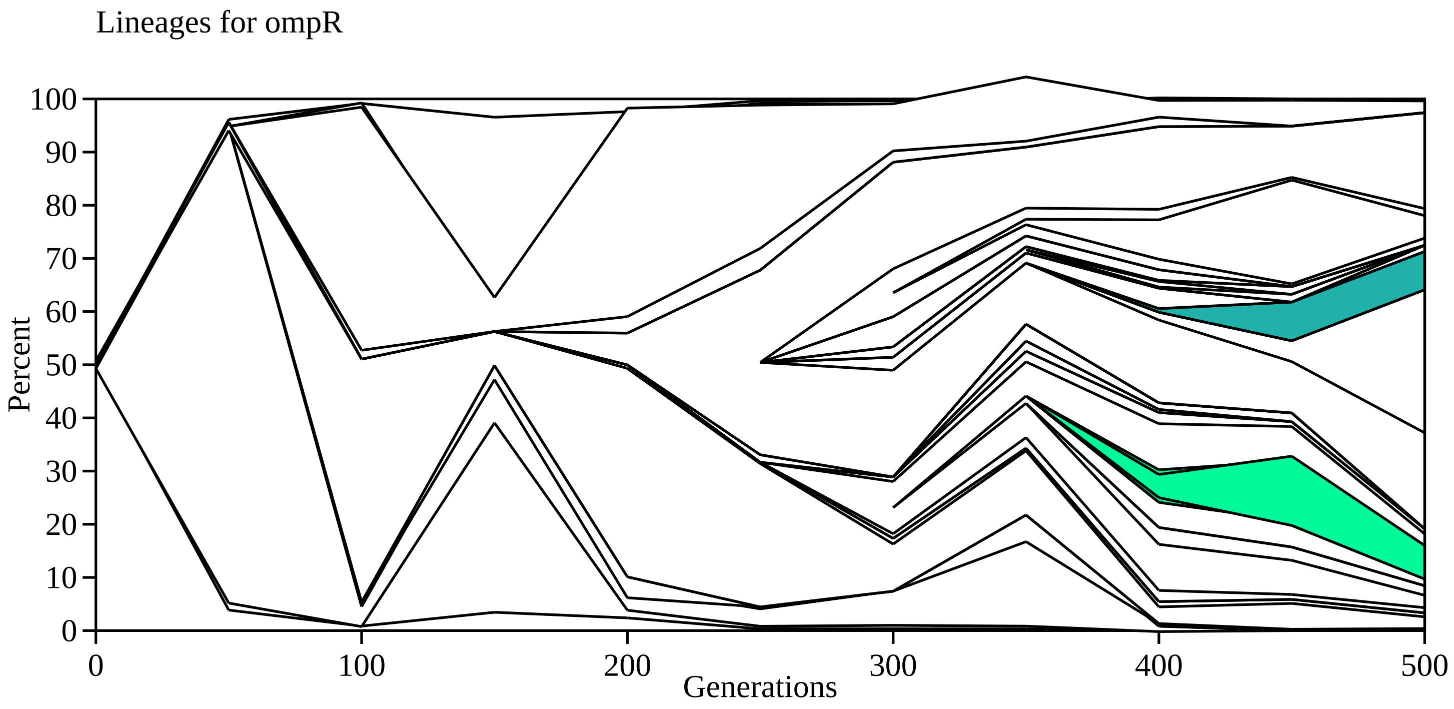

Lineages for ompR
Percent
Generations
0
10
20
30
40
50
60
70
80
90
100
0
100
200
300
400
500

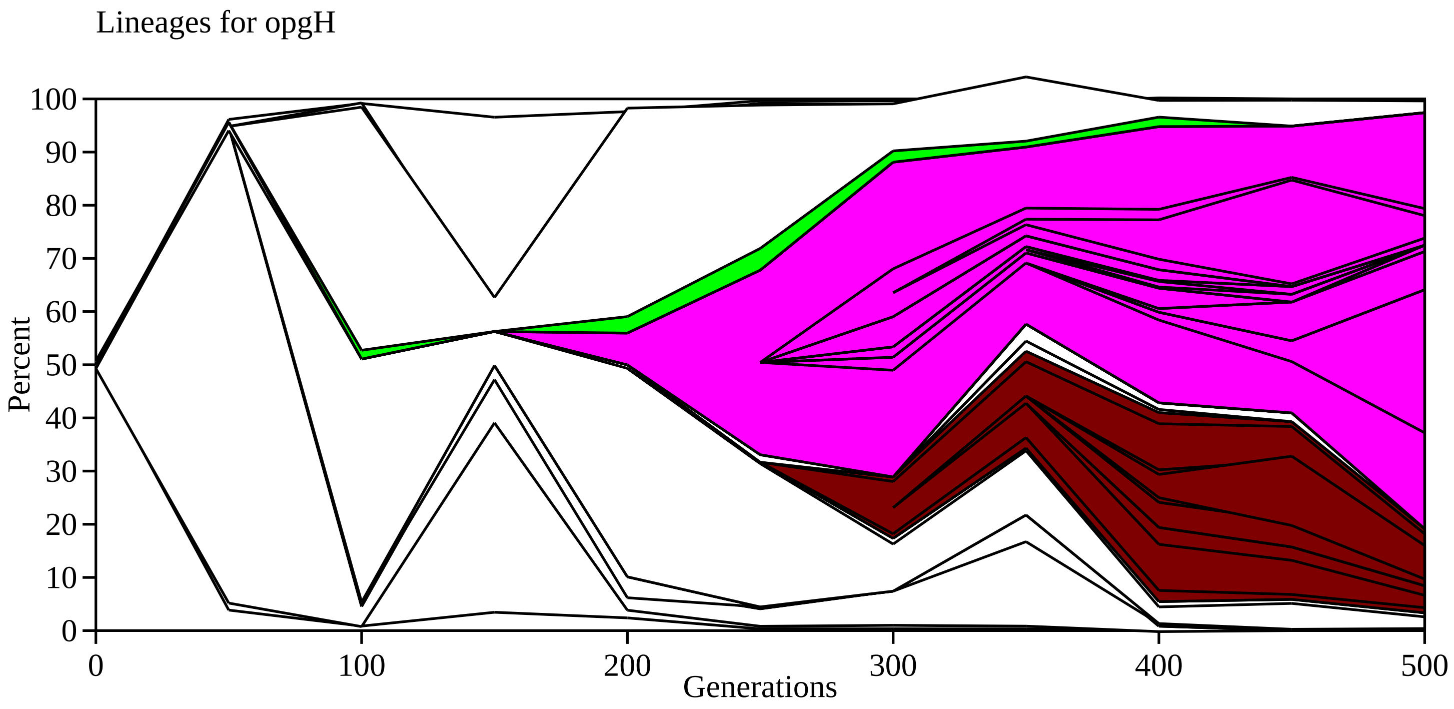

Lineages for opgH
Percent
100
90
80
70
60
50
40
30
20
10
0
0
100
200
300
400
500
Generations

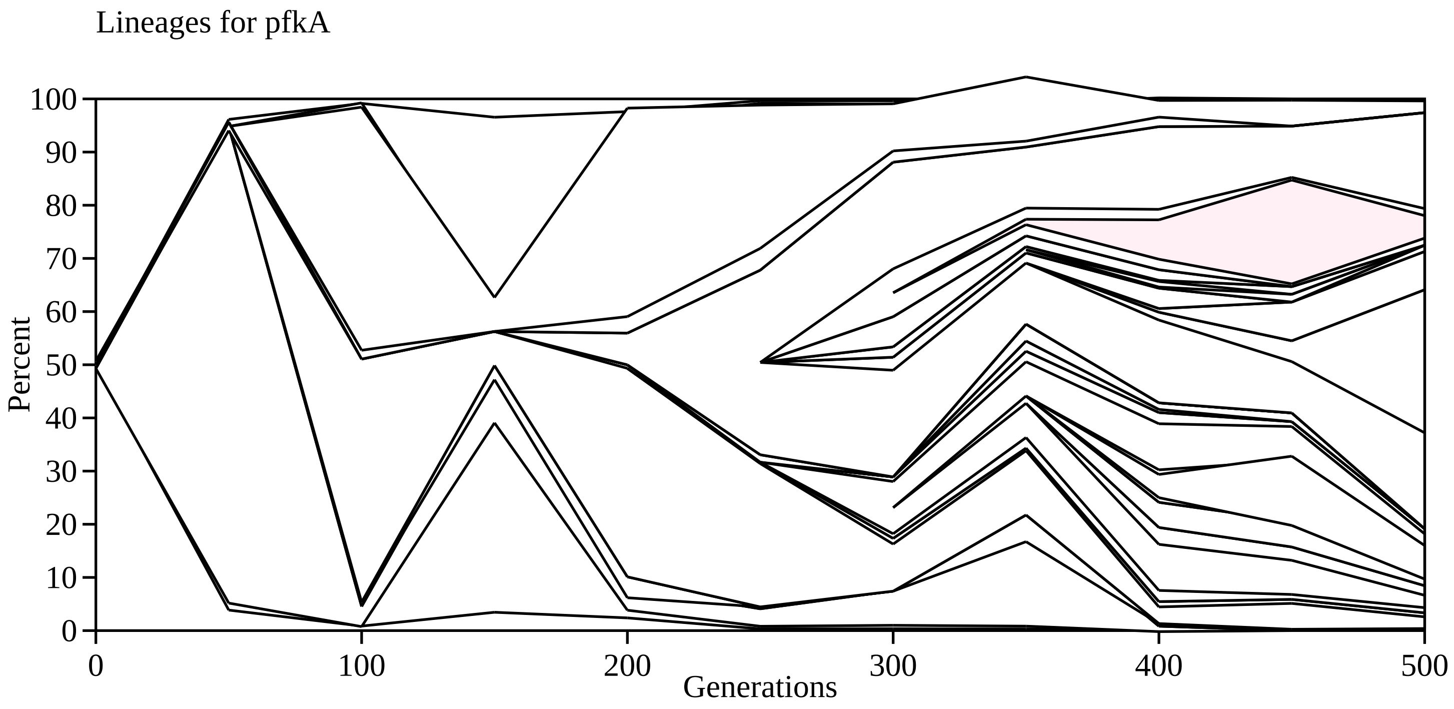
Lineages for pfkA
Percent
0
10
20
30
40
50
60
70
80
90
100
0
100
200
300
400
500
Generations

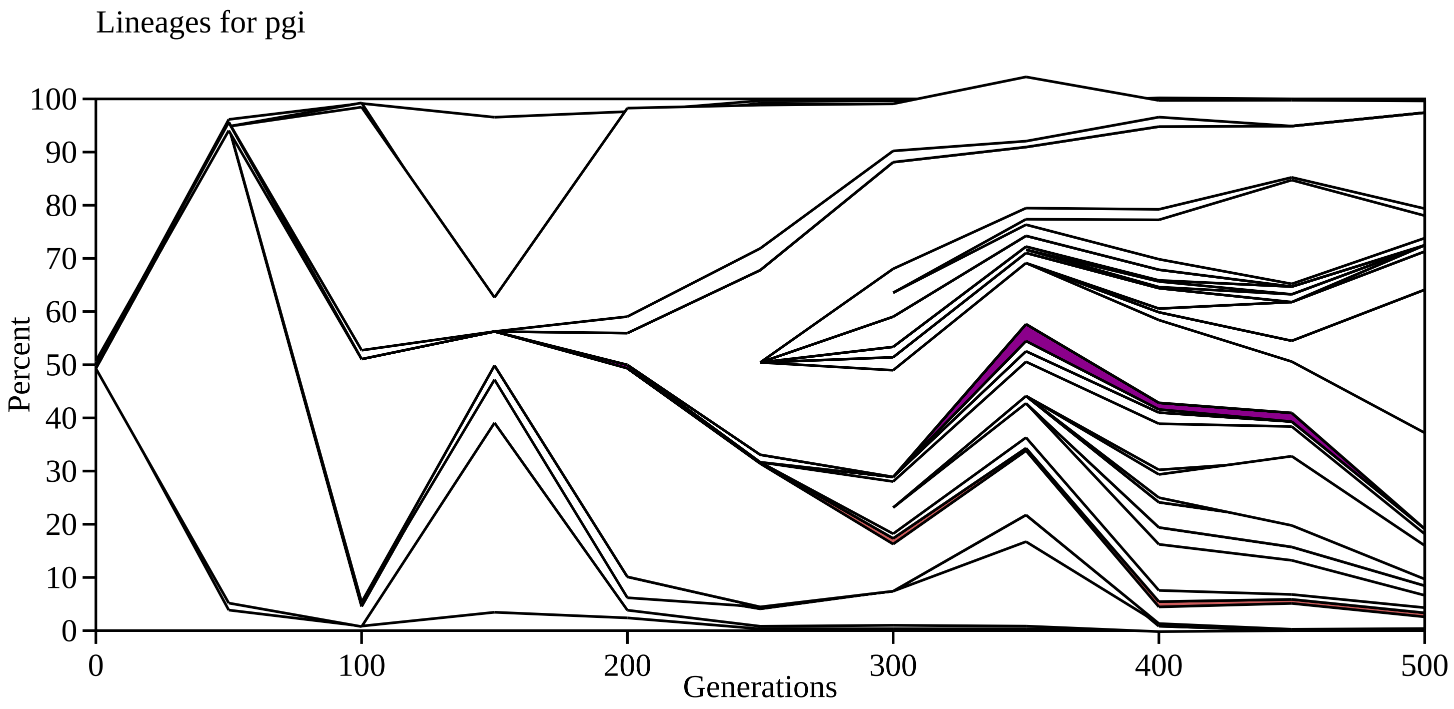
Lineages for pgi
Percent
Generations
0
10
20
30
40
50
60
70
80
90
100
0
100
200
300
400
500

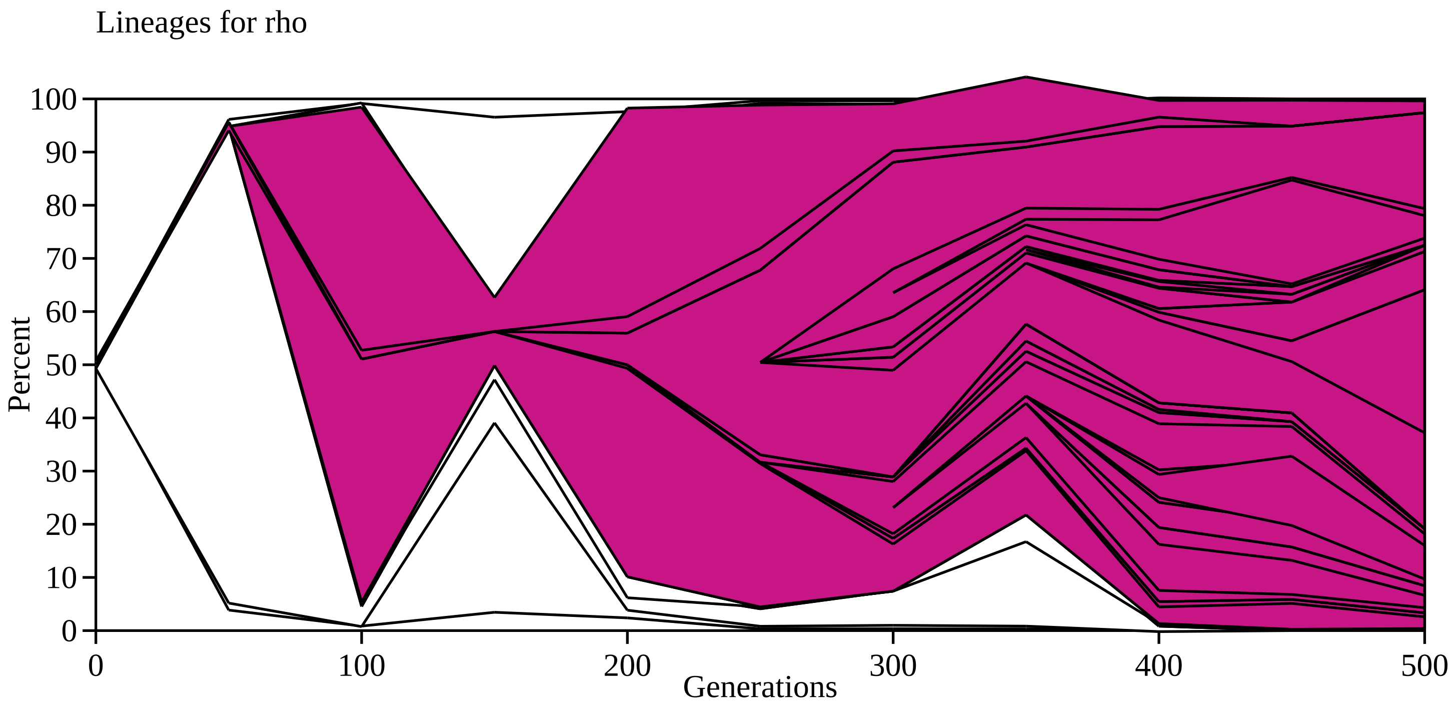

Lineages for rho
100
90
80
70
60
50
40
30
20
10
0
Percent
0
100
200
300
400
500
Generations

Lineages for rpoA

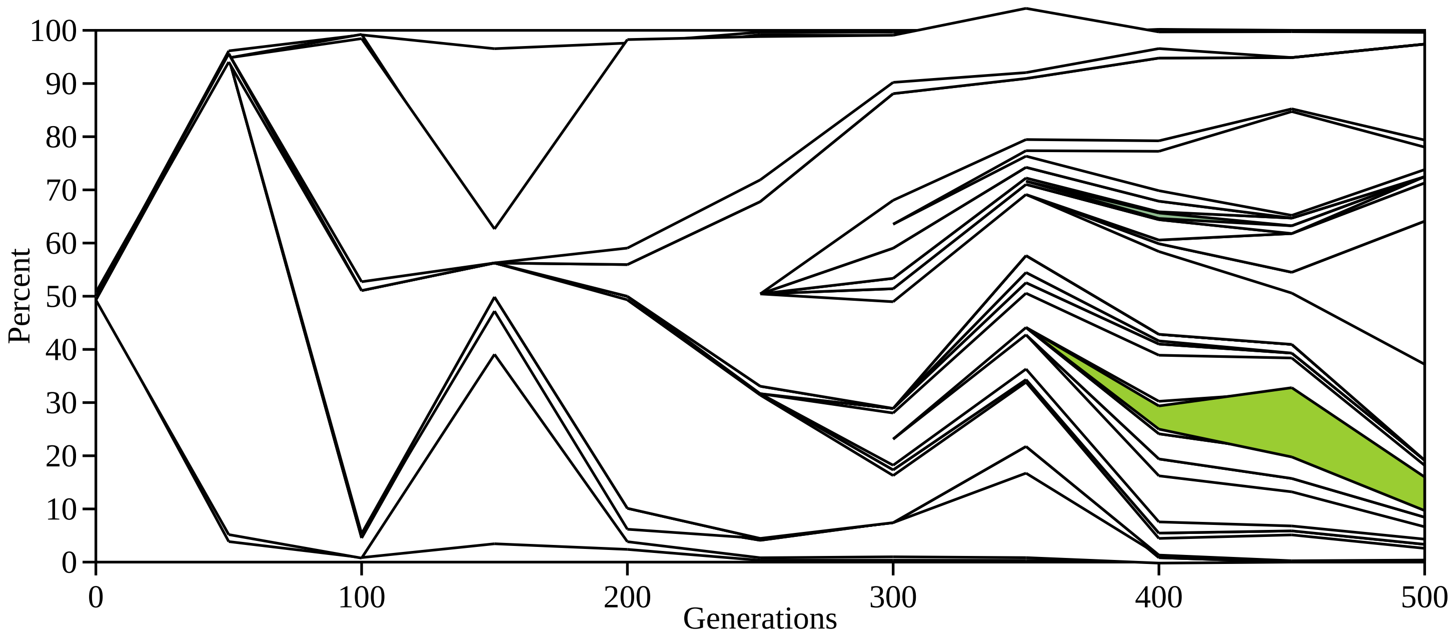

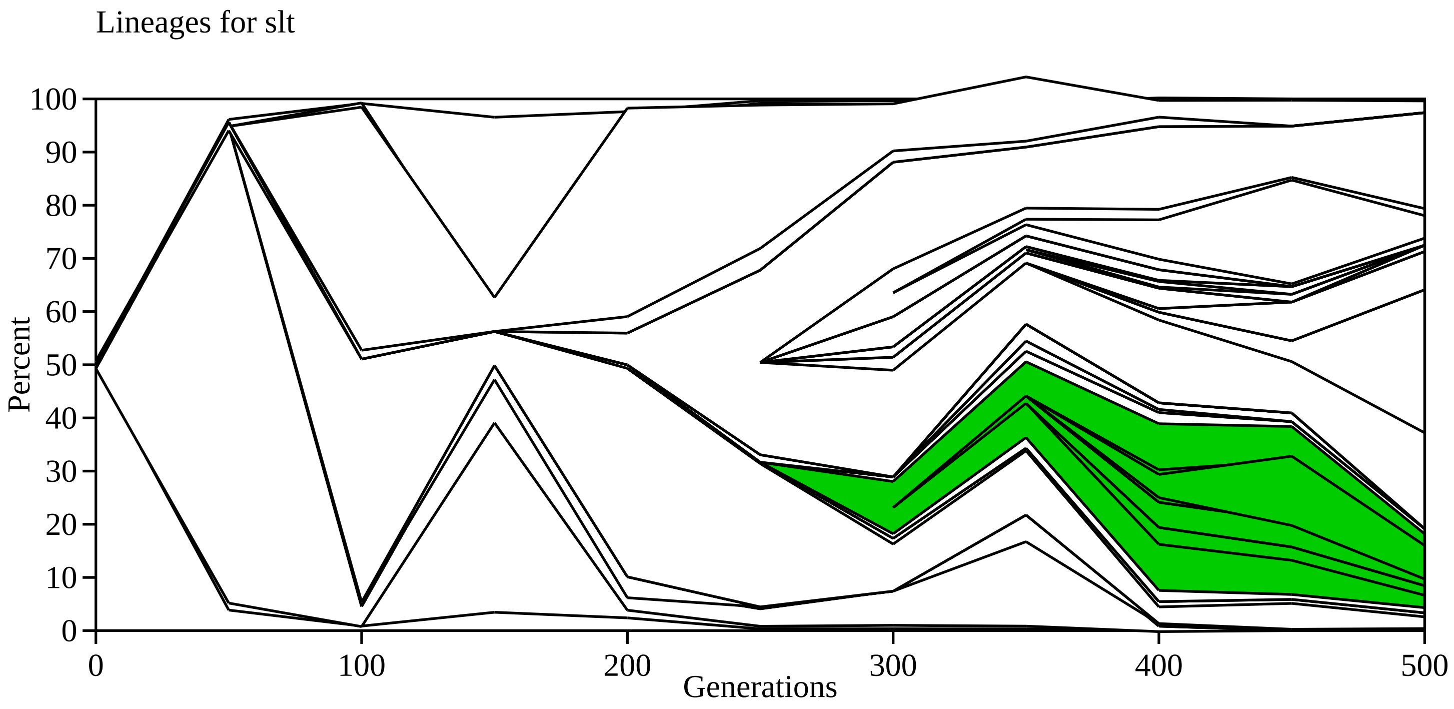

Lineages for slt
Percent
0
10
20
30
40
50
60
70
80
90
100
0
100
200
300
400
500
Generations

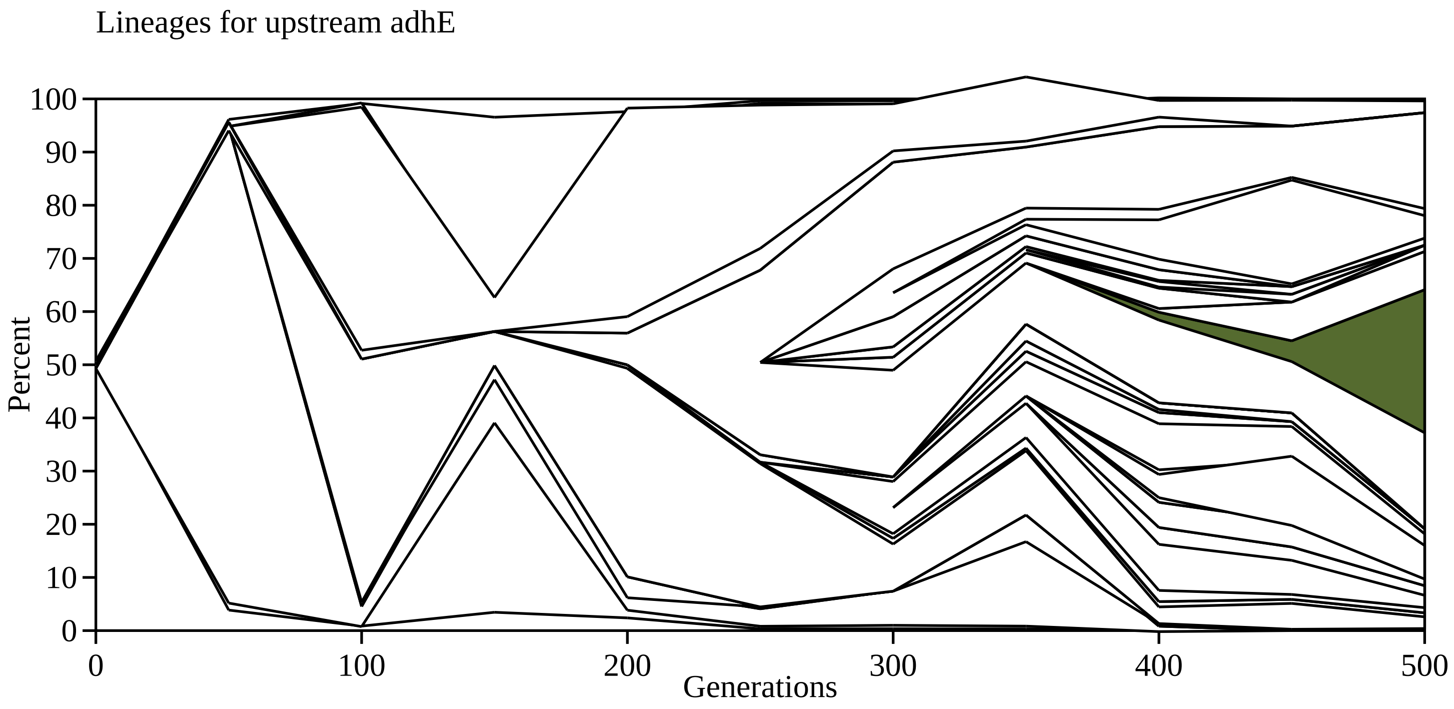

Lineages for upstream adhE
Percent
100
90
80
70
60
50
40
30
20
10
0
0
100
200
300
400
500
Generations

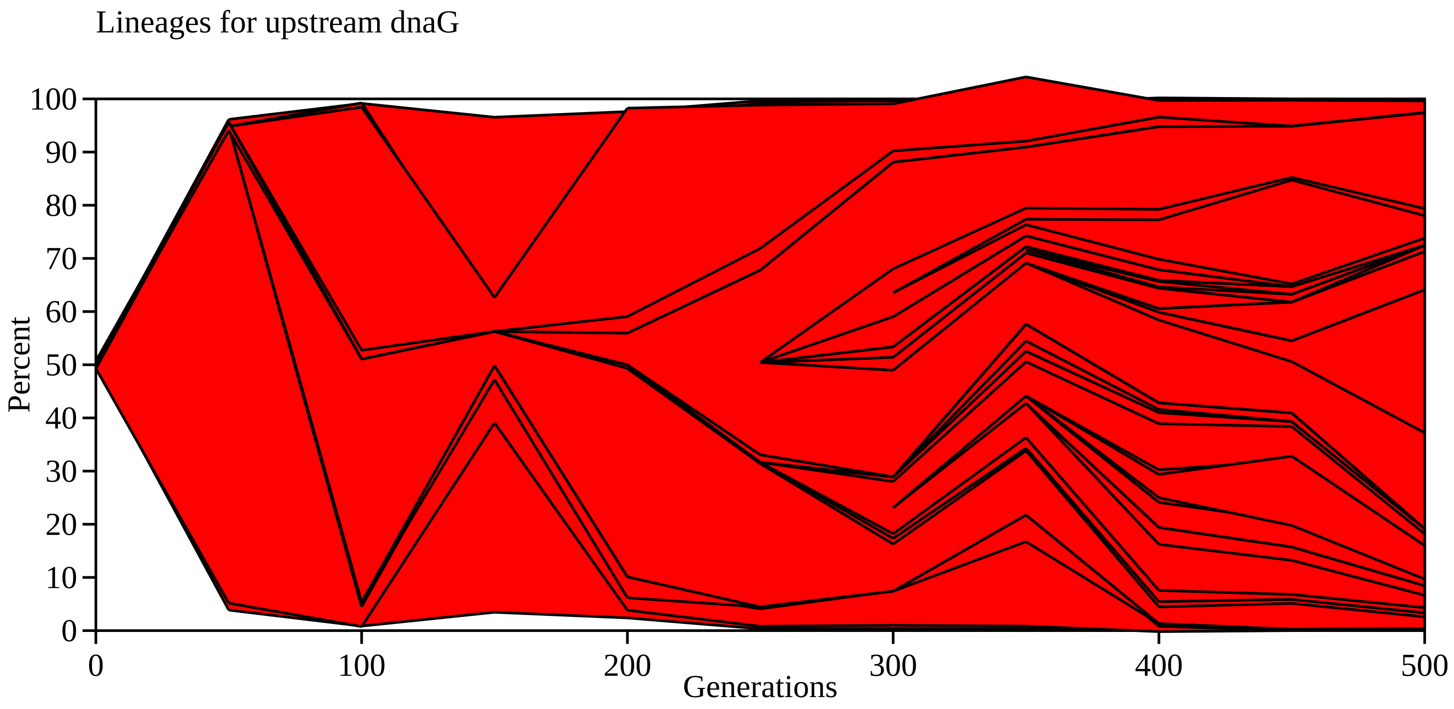

Lineages for upstream dnaG
Percent
100
90
80
70
60
50
40
30
20
10
0
0
100
200
300
400
500
Generations

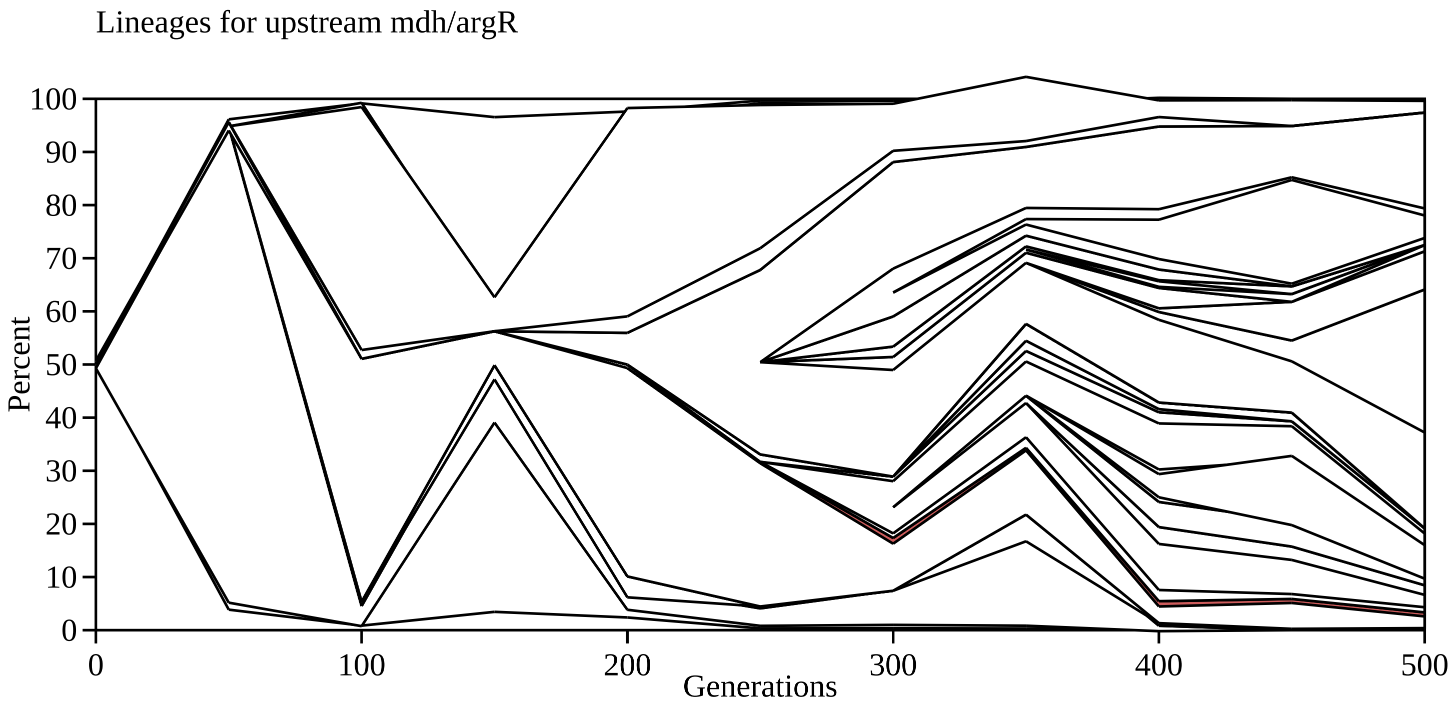

Lineages for upstream mdh/argR
Percent
100
90
80
70
60
50
40
30
20
10
0
0
100
200
300
400
500
Generations

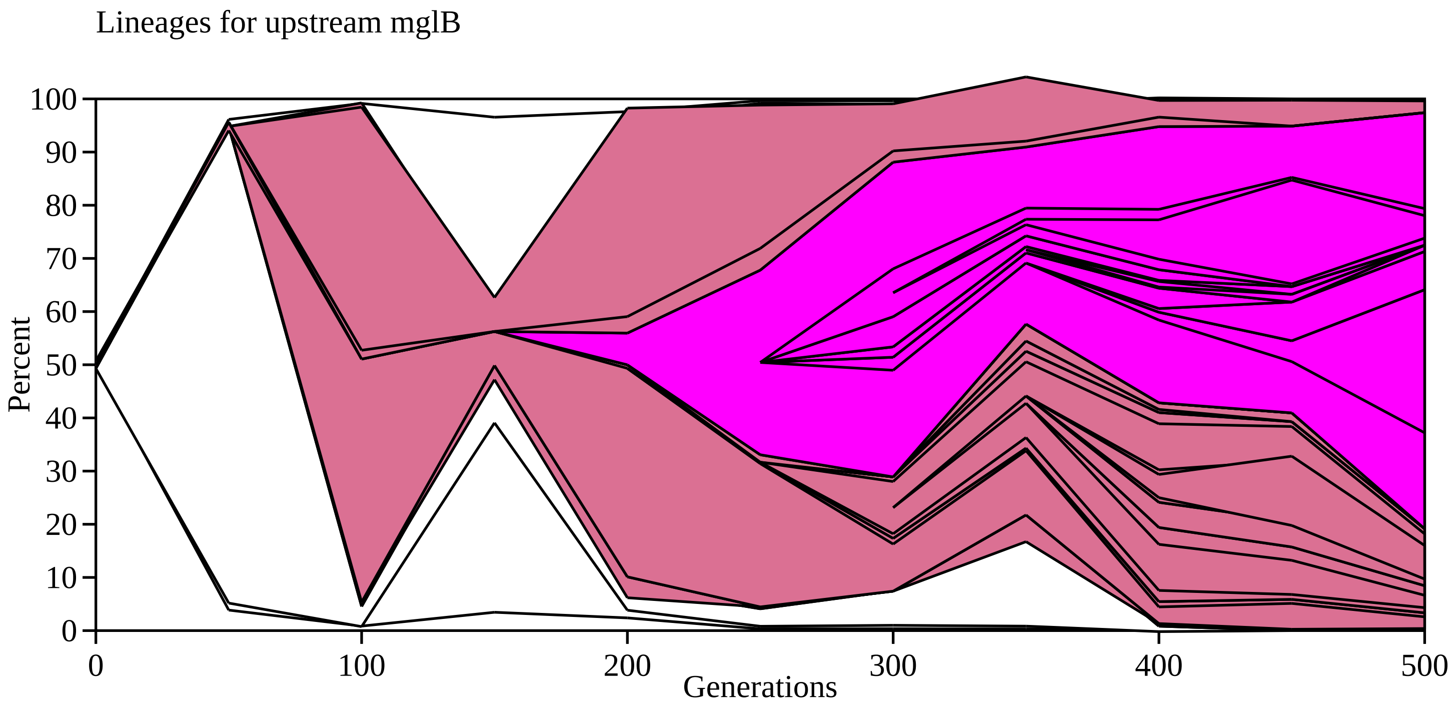

Lineages for upstream mglB
Percent
100
90
80
70
60
50
40
30
20
10
0
0
100
200
300
400
500
Generations

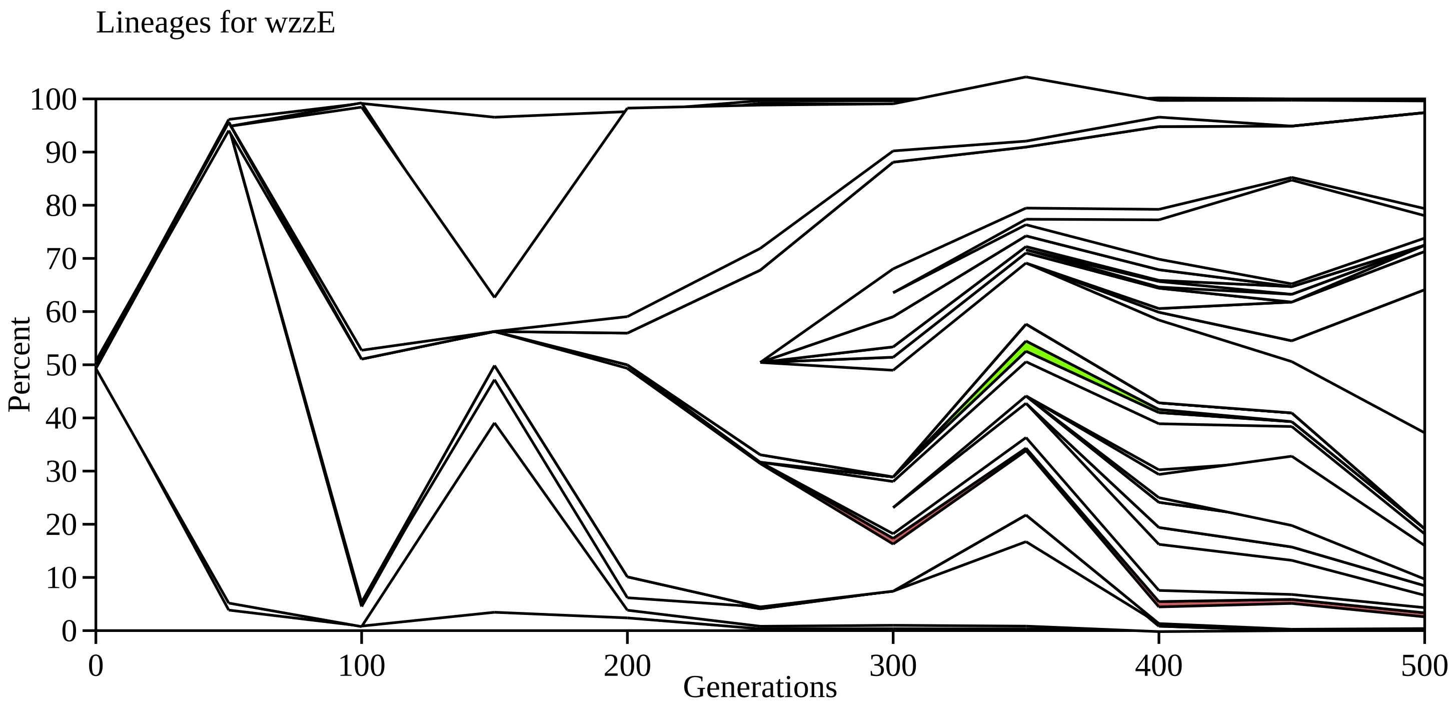

Lineages for wzzE
Percent
Generations
0
10
20
30
40
50
60
70
80
90
100
0
100
200
300
400
500

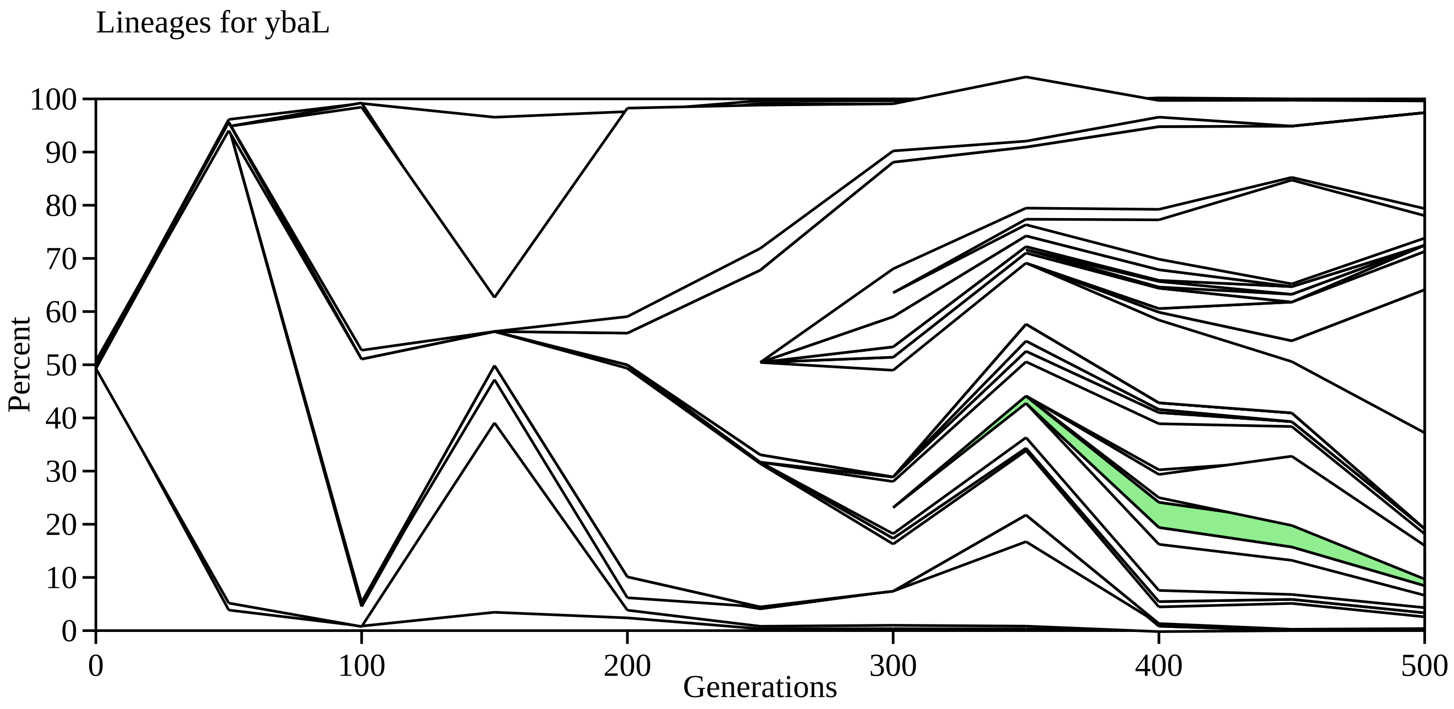

Lineages for ybaL
Percent
100
90
80
70
60
50
40
30
20
10
0
0
100
200
300
400
500
Generations

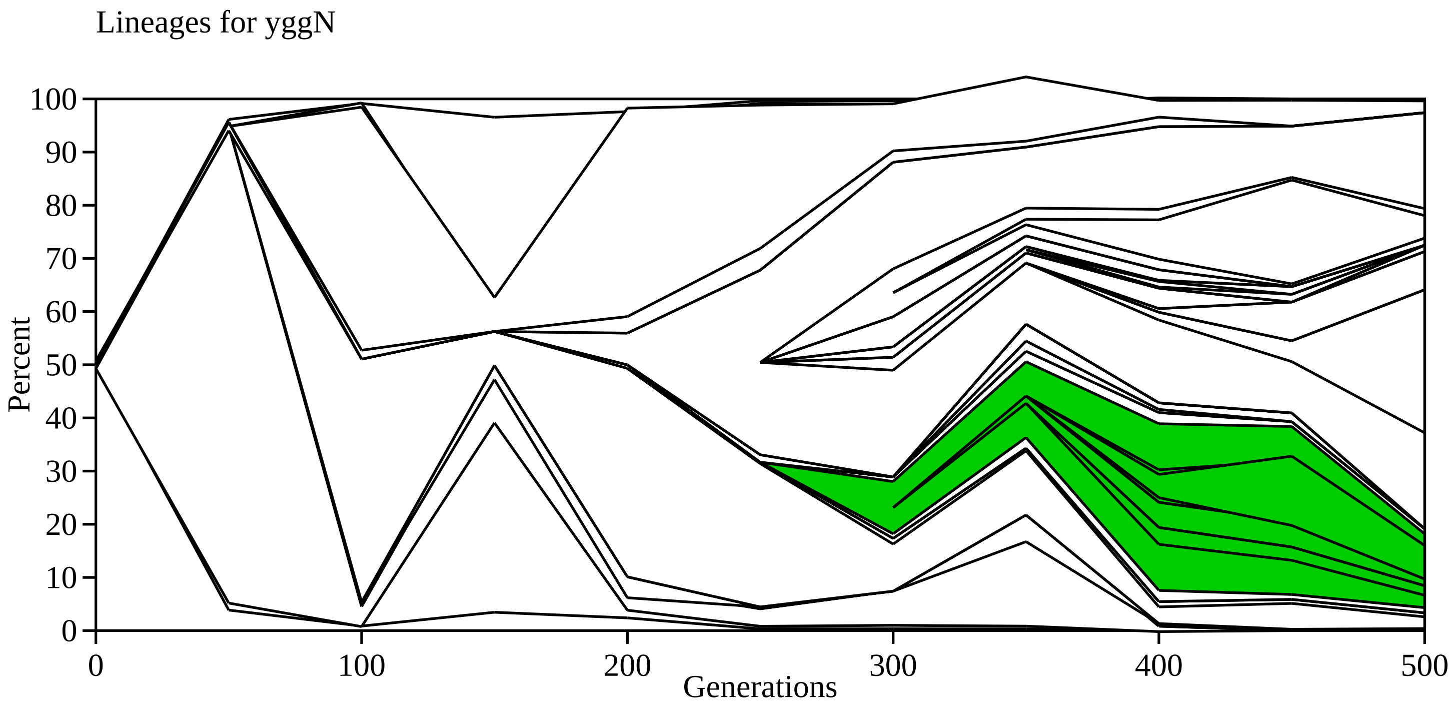

Lineages for yggN
100
90
80
70
60
50
40
30
20
10
0
Percent
0
100
200
300
400
500
Generations

0.1 (upstream dnaG)

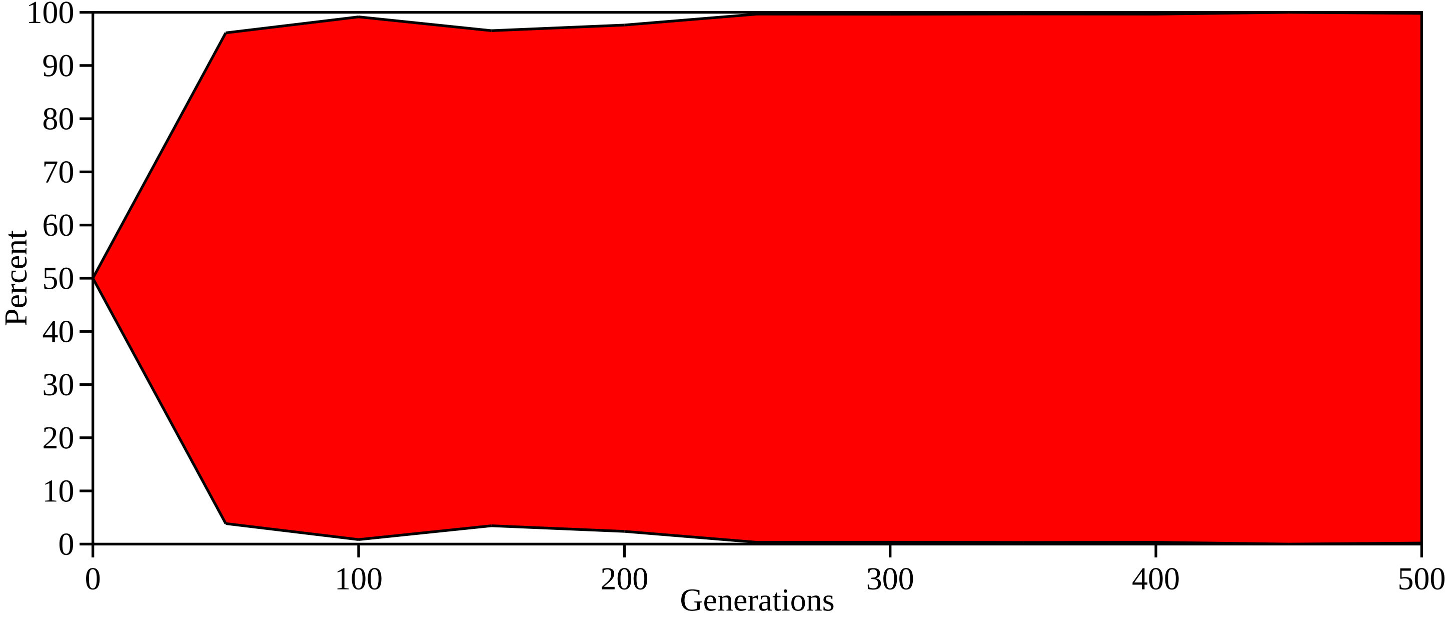

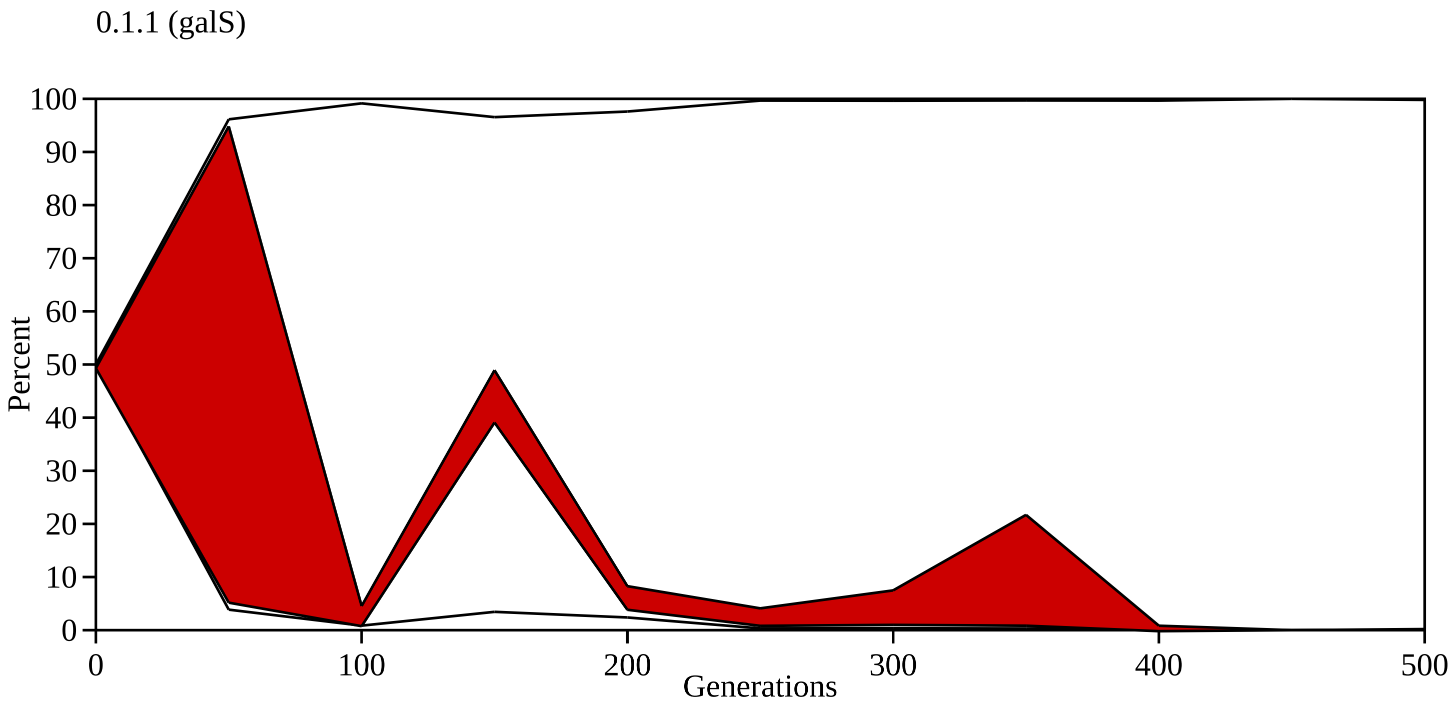

0.1.1 (galS)
Percent
100
90
80
70
60
50
40
30
20
10
0
0
100
200
300
400
500
Generations

0.1.2 (upstream mglB)

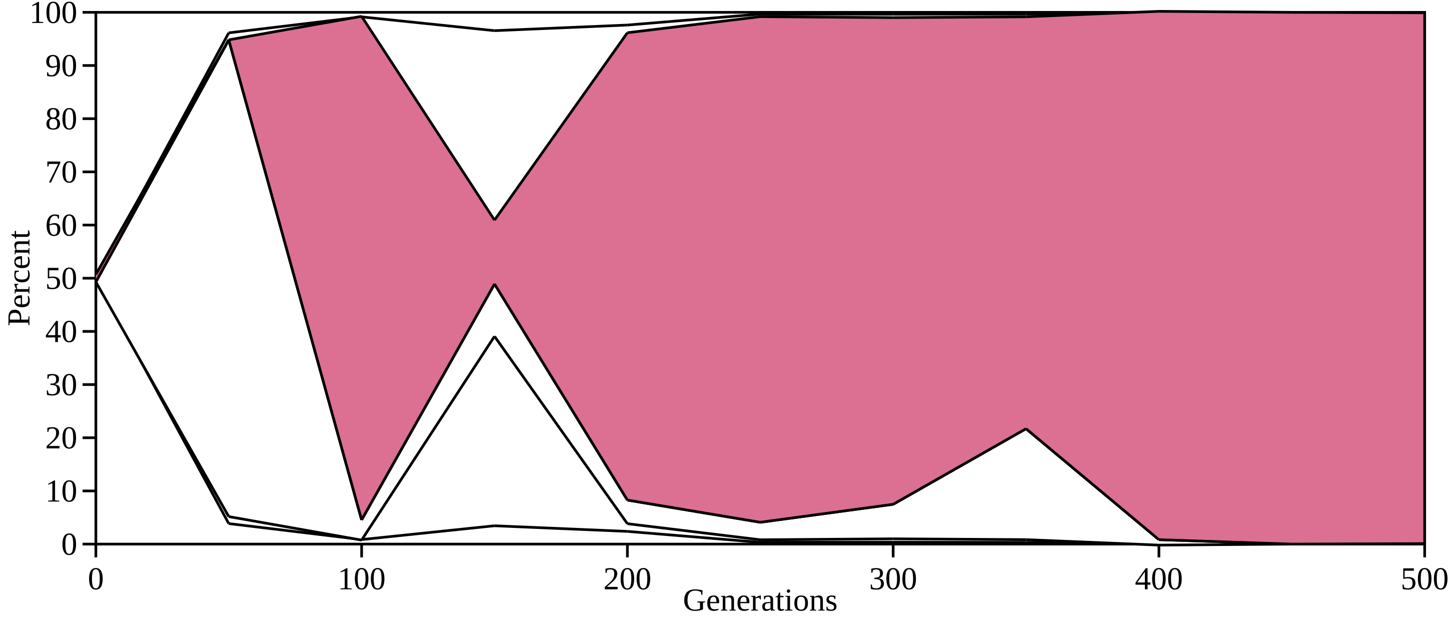

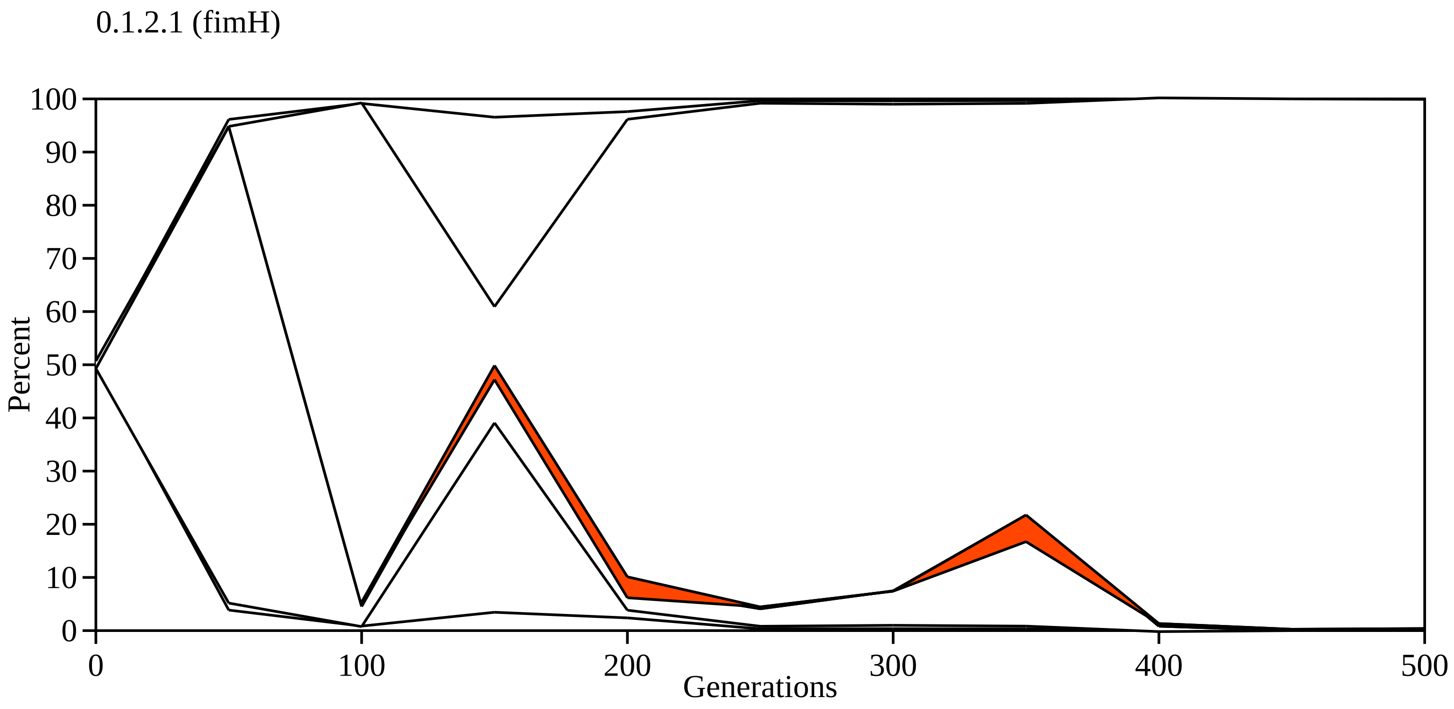
0.1.2.1 (fimH)
100
90
80
70
60
50
40
30
20
10
0
Percent
0
100
200
300
400
500
Generations

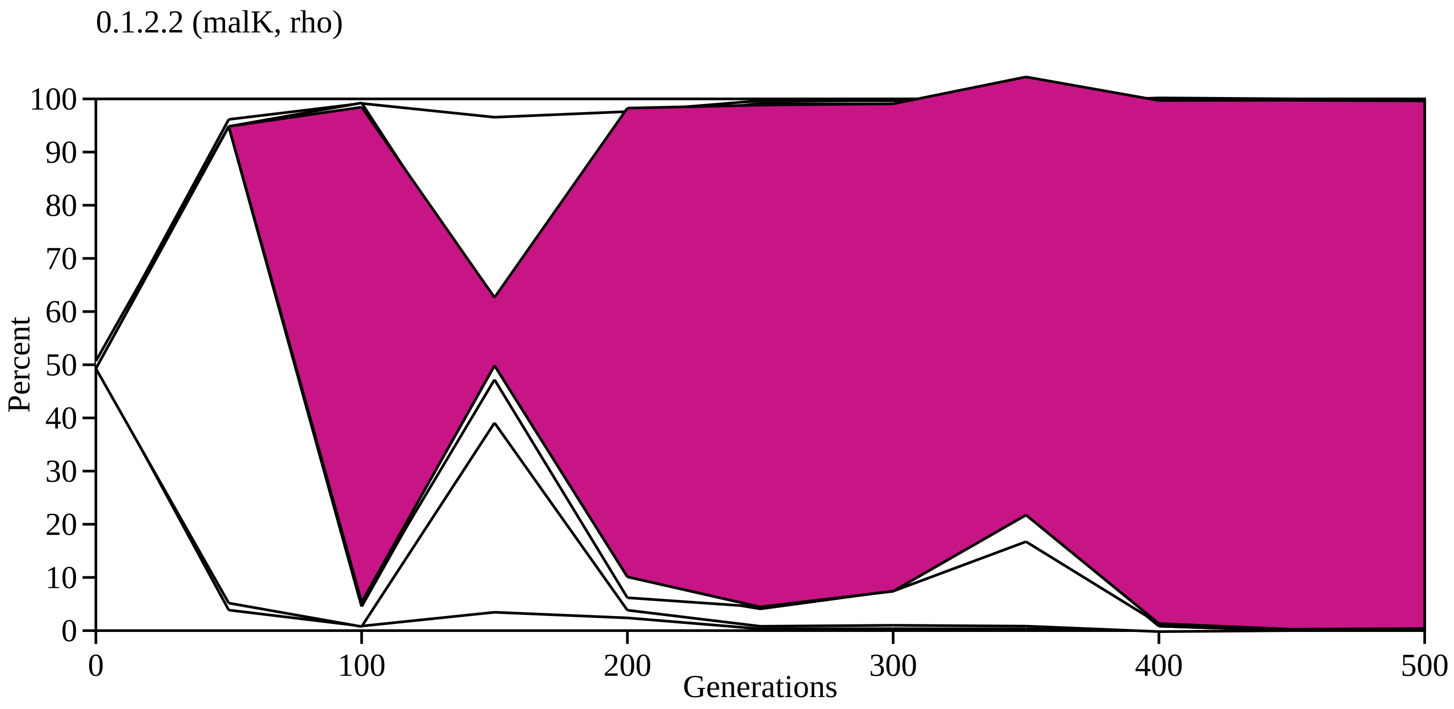

0.1.2.2 (malK, rho)
Percent
100
90
80
70
60
50
40
30
20
10
0
0
100
200
300
400
500
Generations

0.1.2.2.1 (gatZ, wzzE, pgi, lptC, pgi, upstream mdh/argR)

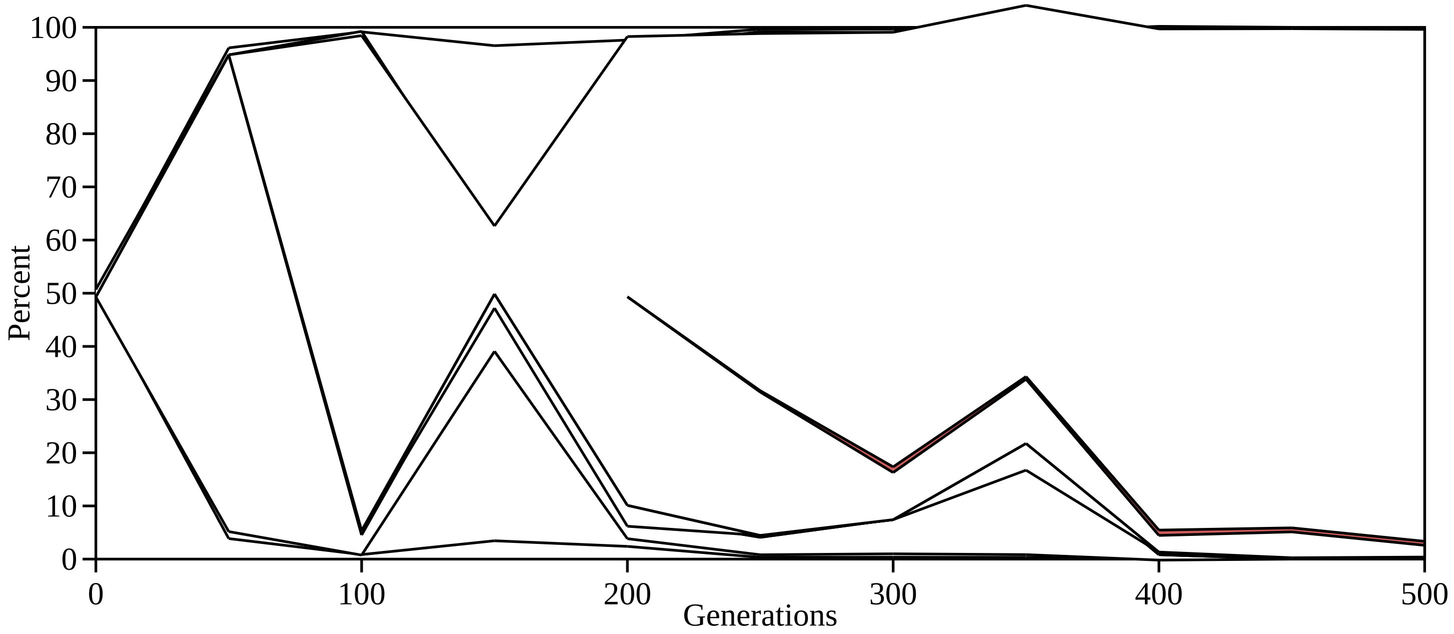

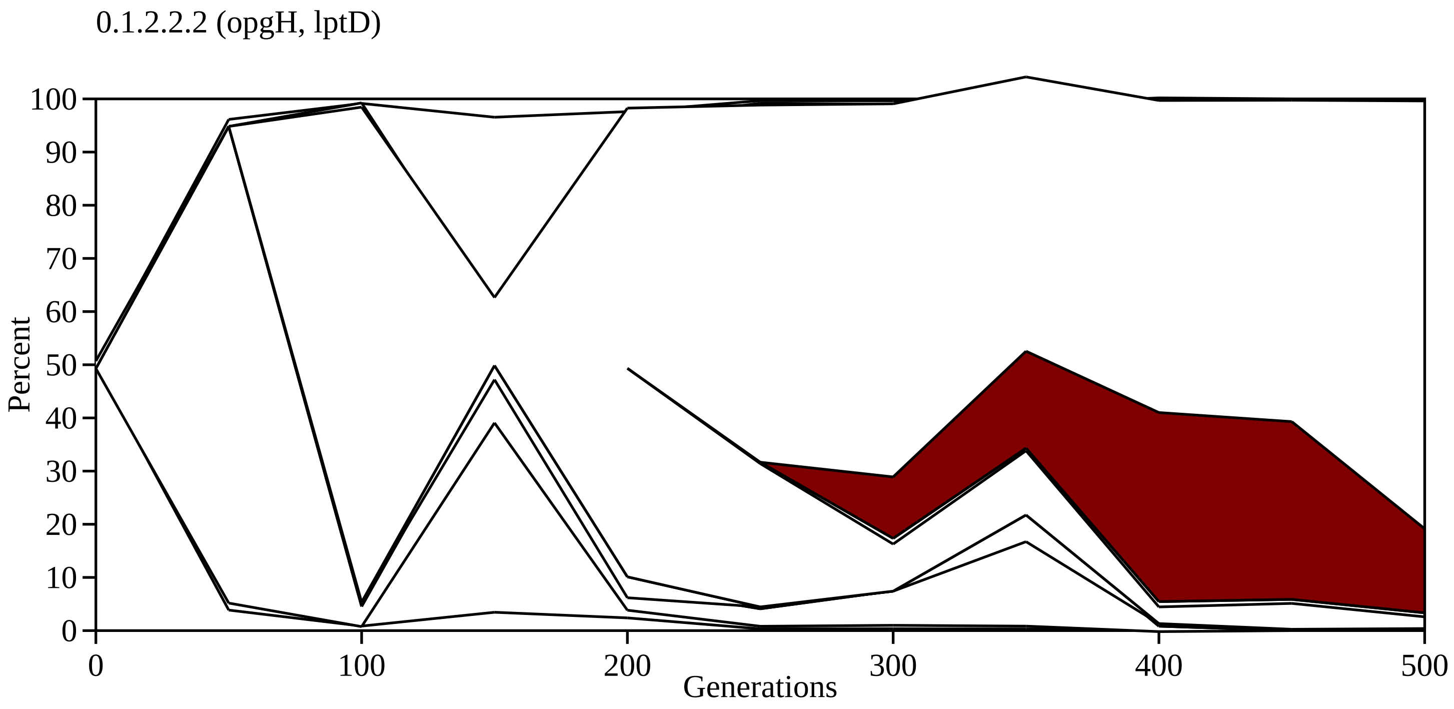
0.1.2.2.2 (opgH, lptD)
Percent
100
90
80
70
60
50
40
30
20
10
0
0
100
200
300
400
500
Generations

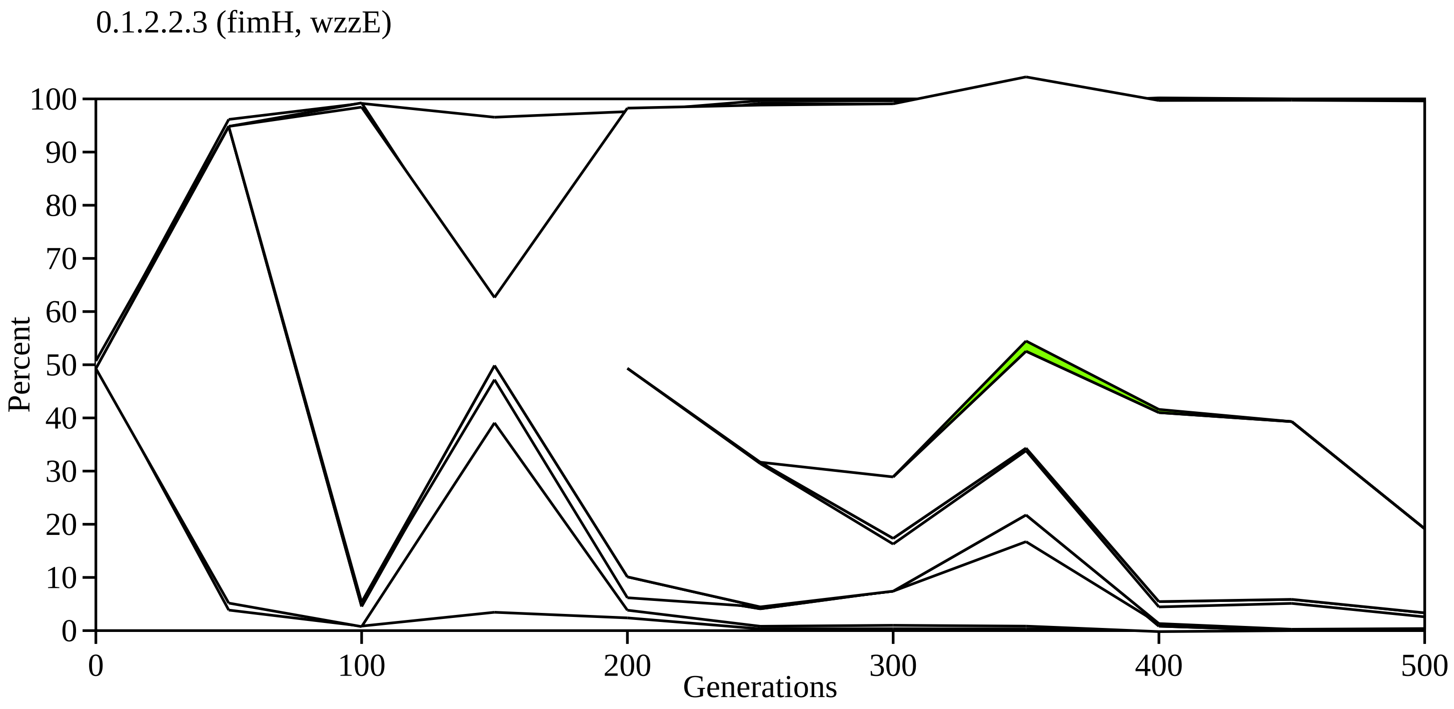
0.1.2.2.3 (fimH, wzzE)
Percent
100
90
80
70
60
50
40
30
20
10
0
0
100
200
300
400
500
Generations

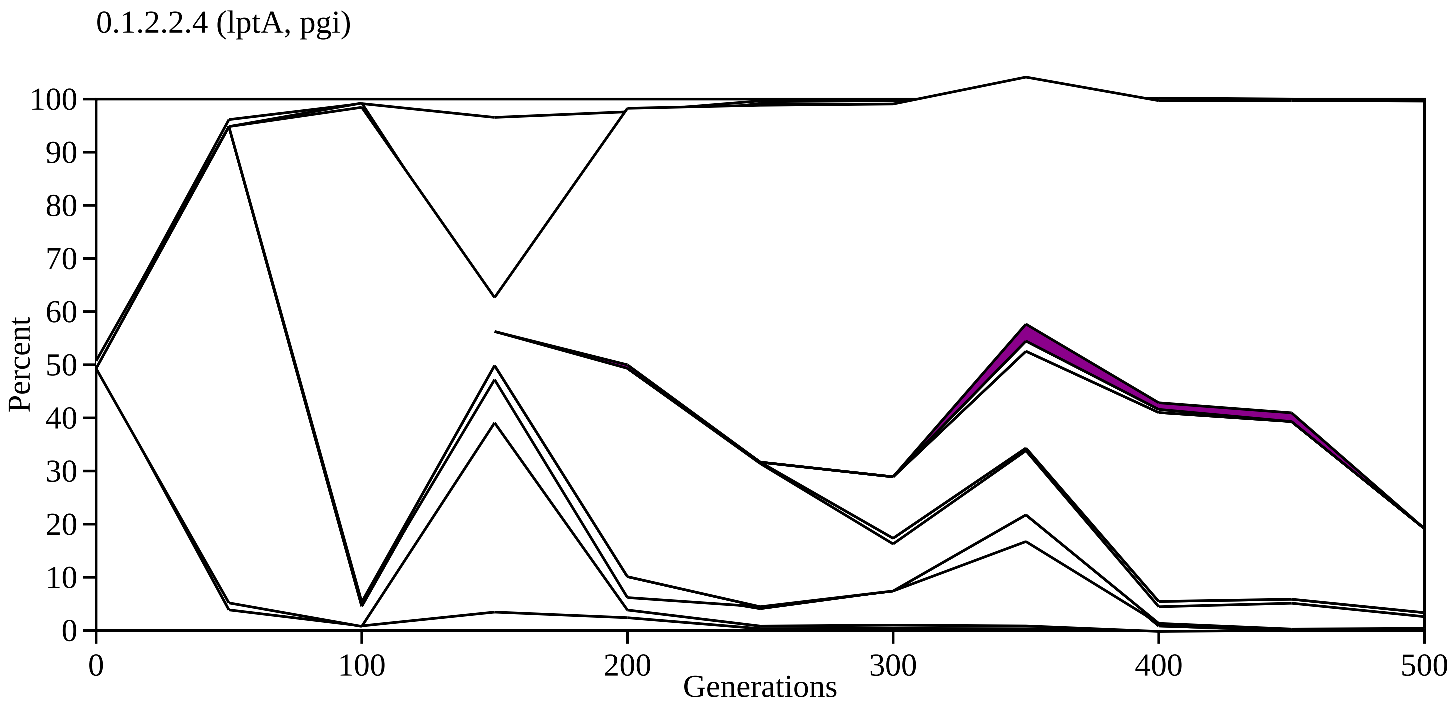
0.1.2.2.4 (lptA, pgi)
Percent
100
90
80
70
60
50
40
30
20
10
0
0
100
200
300
400
500
Generations

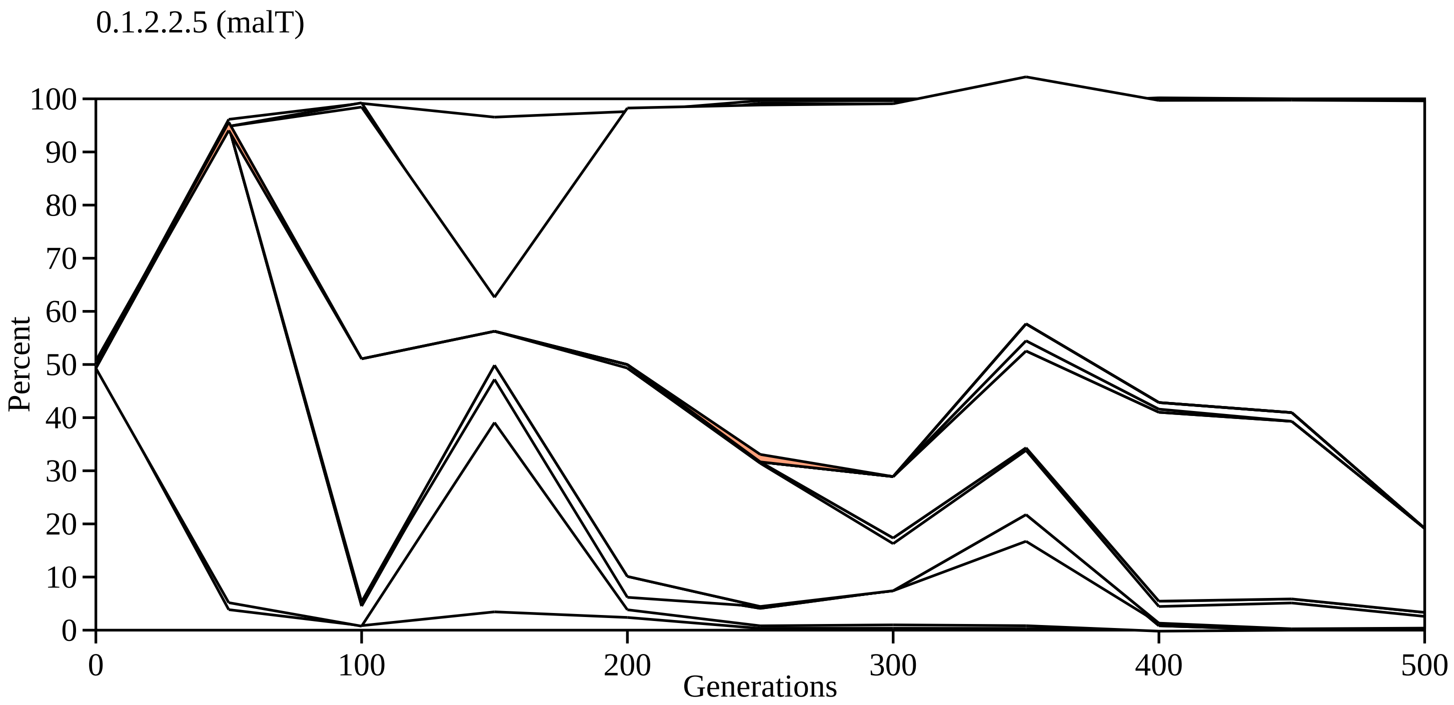
0.1.2.2.5 (malT)
100
90
80
70
60
50
40
30
20
10
0
Percent
0
100
200
300
400
500
Generations

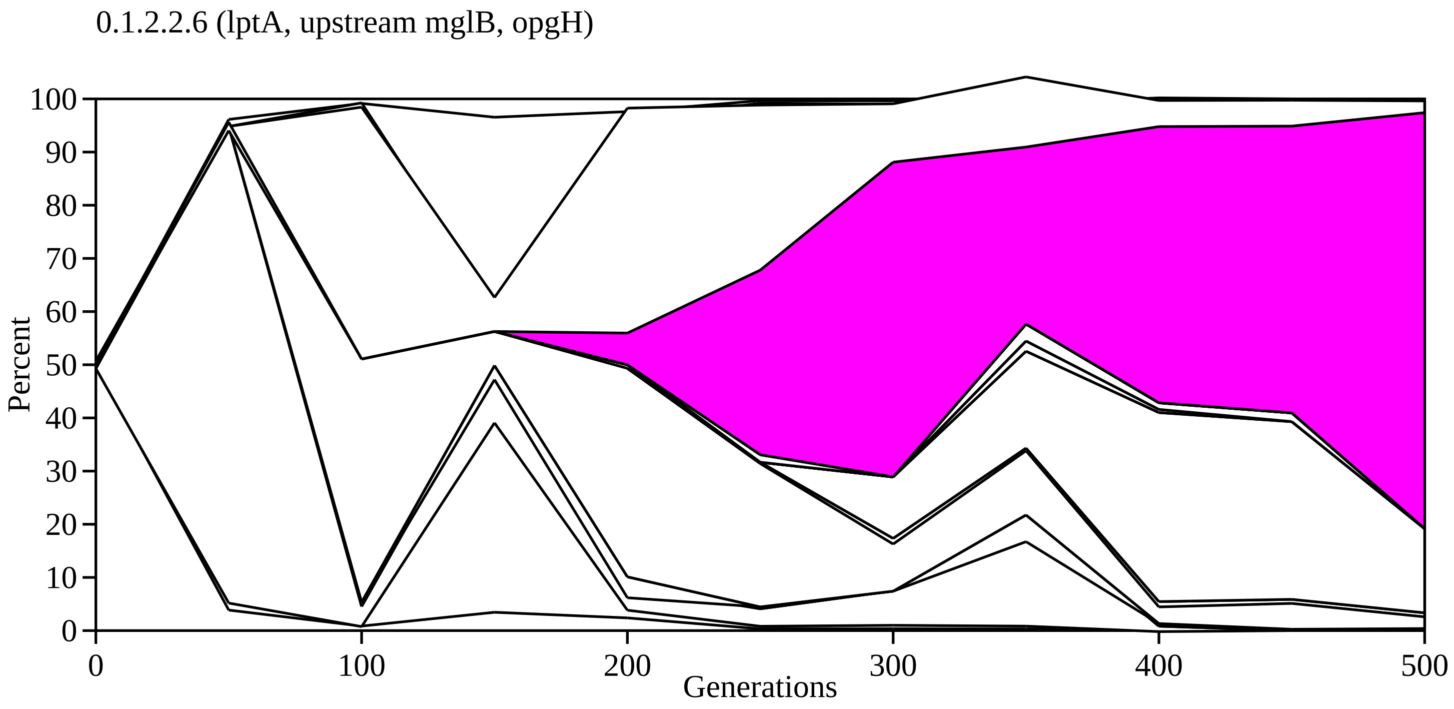
0.1.2.2.6 (lptA, upstream mglB, opgH)
Percent
0
10
20
30
40
50
60
70
80
90
100
0
100
200
300
400
500
Generations

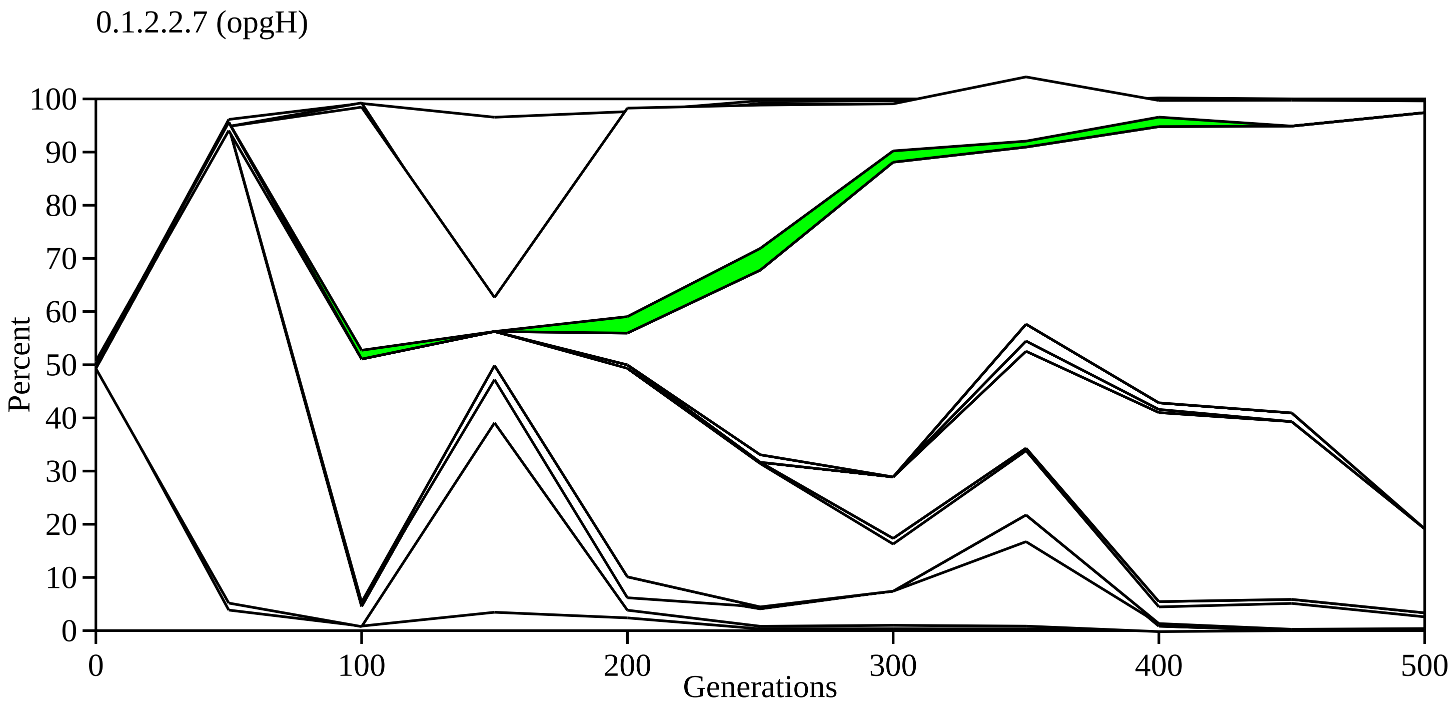
0.1.2.2.7 (opgH)
Percent
100
90
80
70
60
50
40
30
20
10
0
0
100
200
300
400
500
Generations

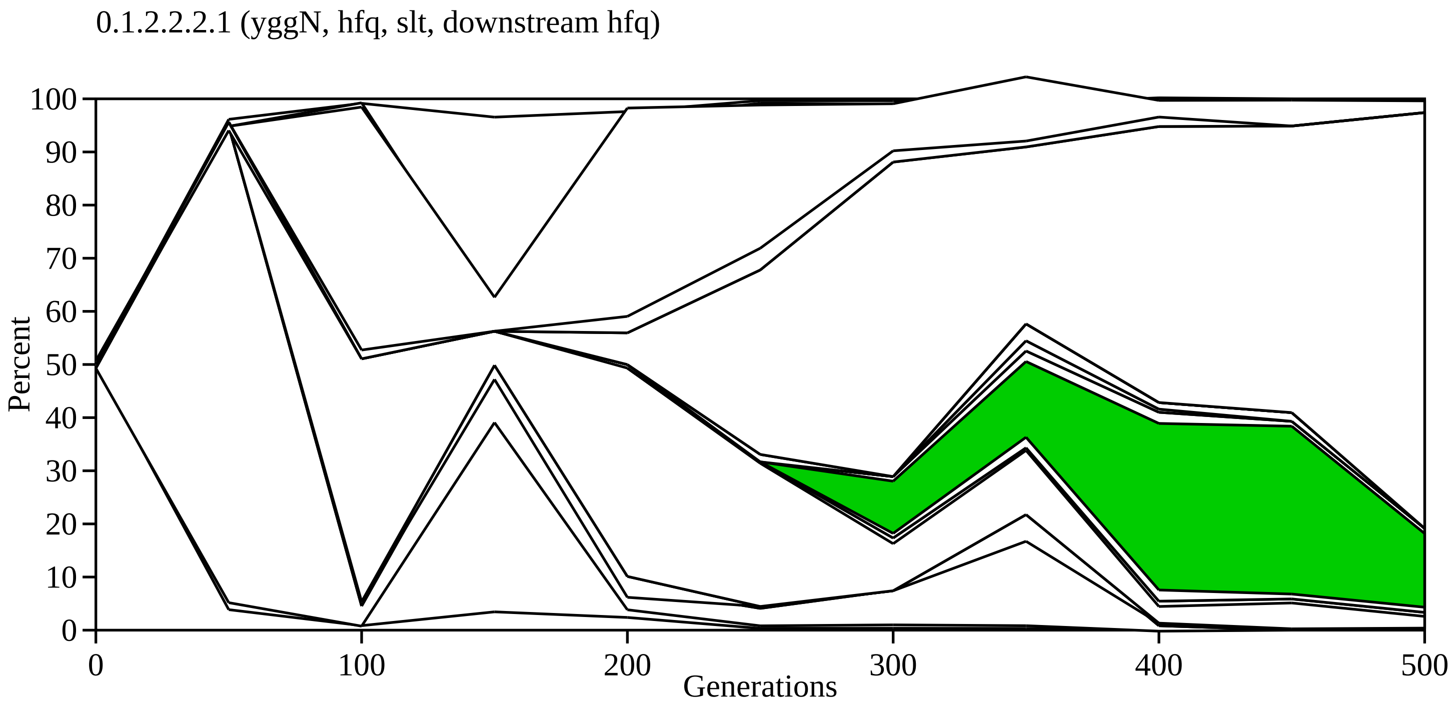

0.1.2.2.2.1 (yggN, hfq, slt, downstream hfq)
Percent
Generations
0
10
20
30
40
50
60
70
80
90
100
0
100
200
300
400
500

0.1.2.2.2.1.1 (gatZ)

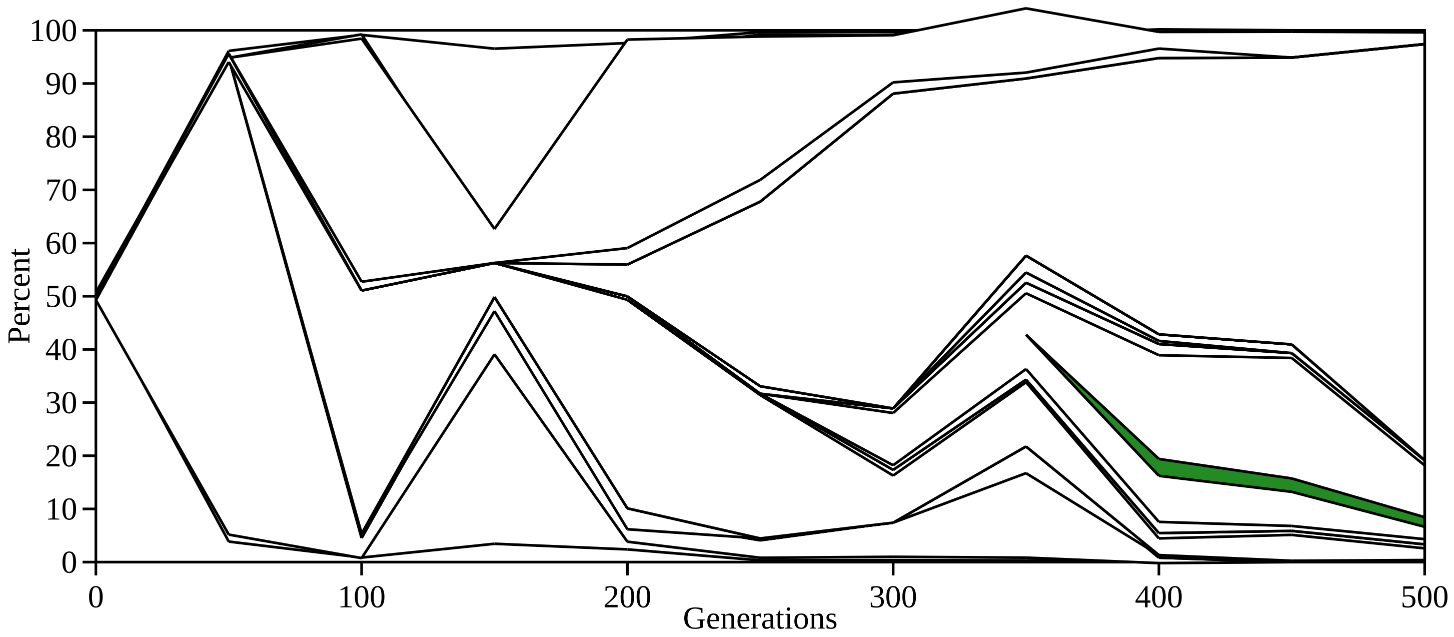

0.1.2.2.2.1.2 (ybaL)

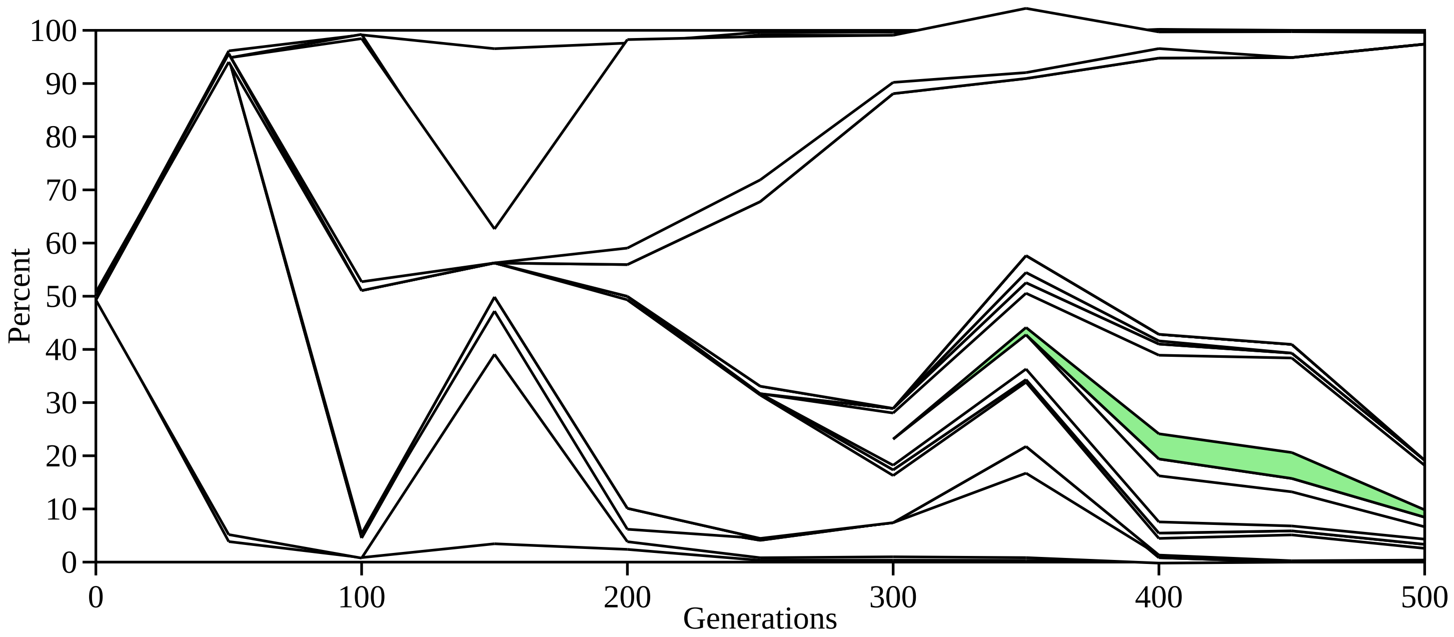

0.1.2.2.2.1.3 (ompR)

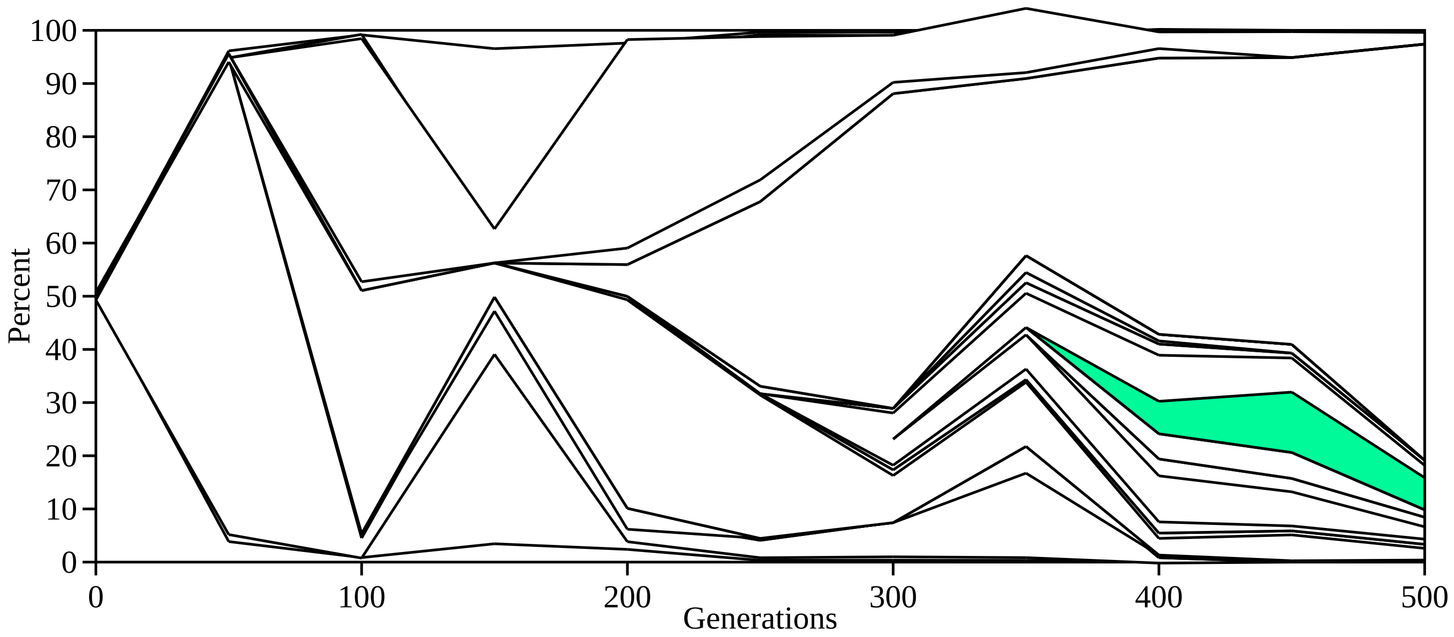

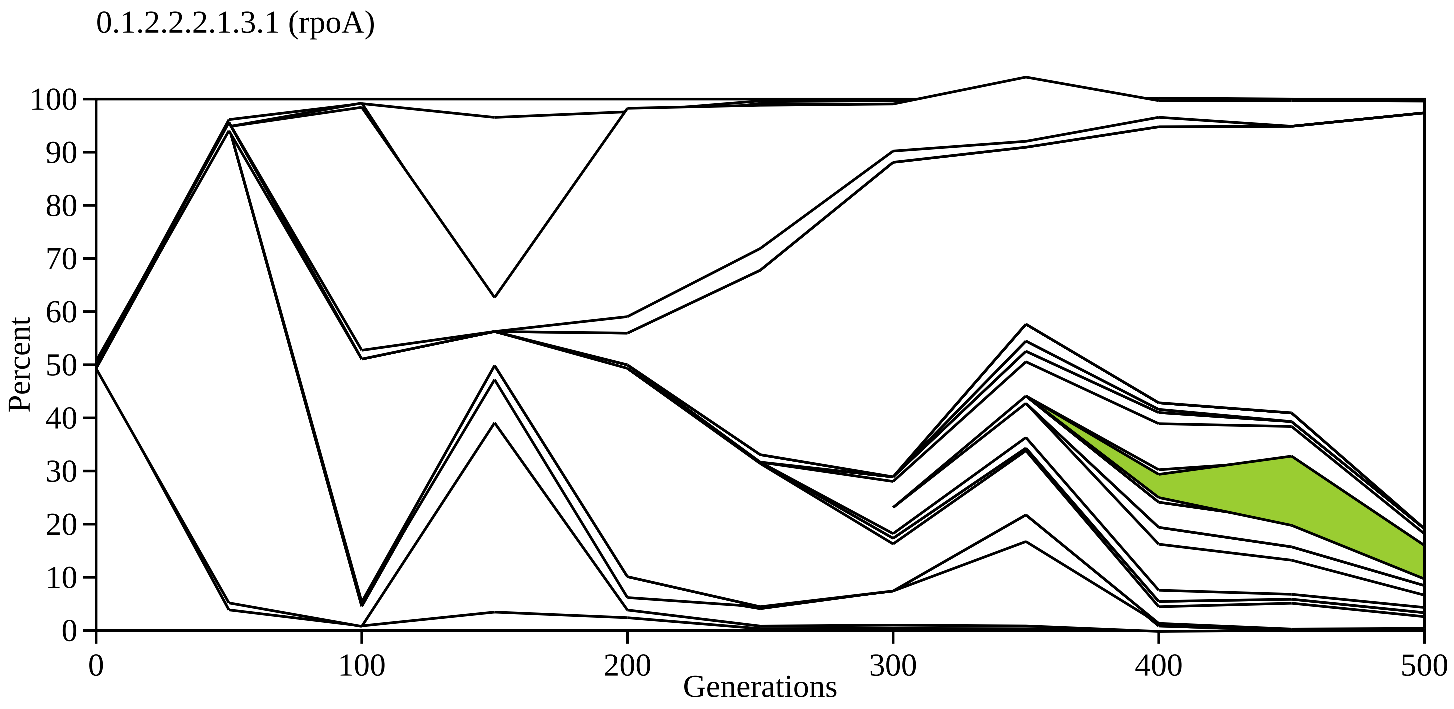
0.1.2.2.2.1.3.1 (rpoA)
Percent
100
90
80
70
60
50
40
30
20
10
0
0
100
200
300
400
500
Generations

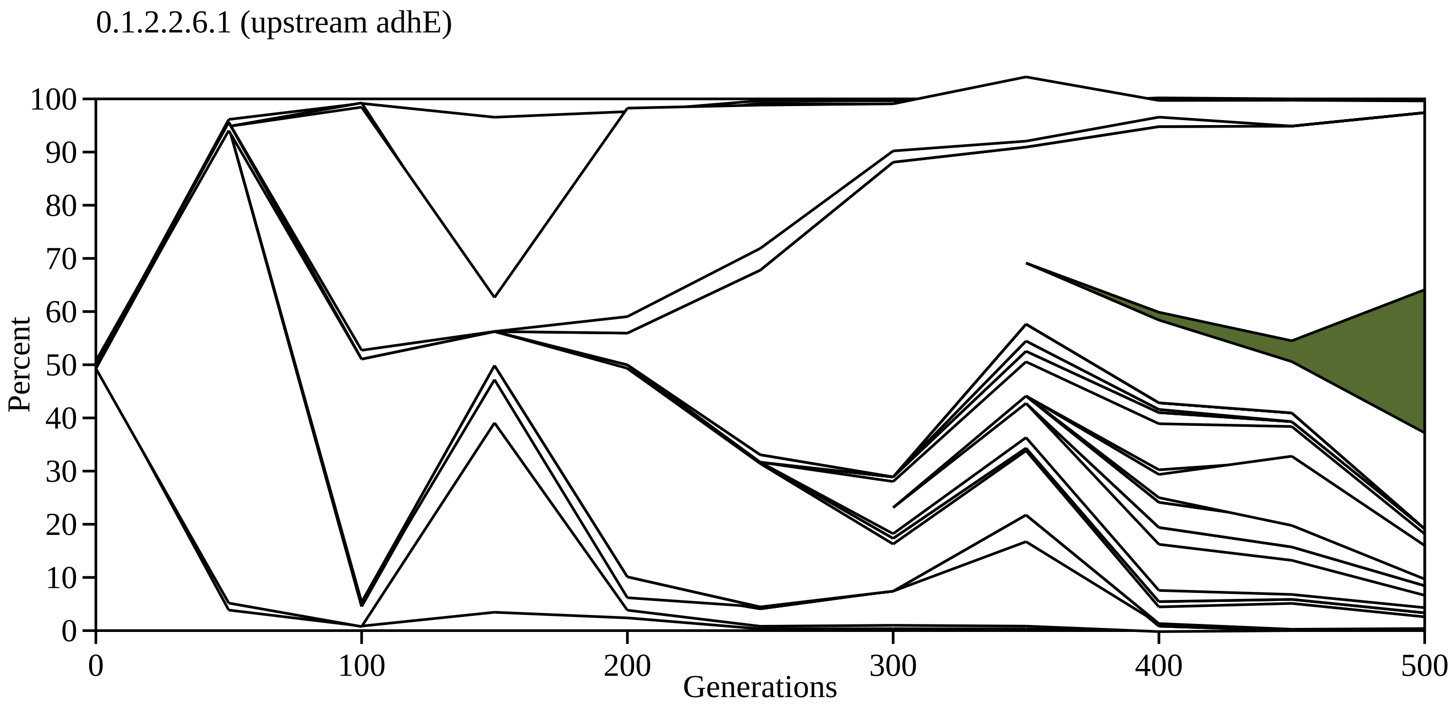
0.1.2.2.6.1 (upstream adhE)
Percent
100
90
80
70
60
50
40
30
20
10
0
0
100
200
300
400
500
Generations

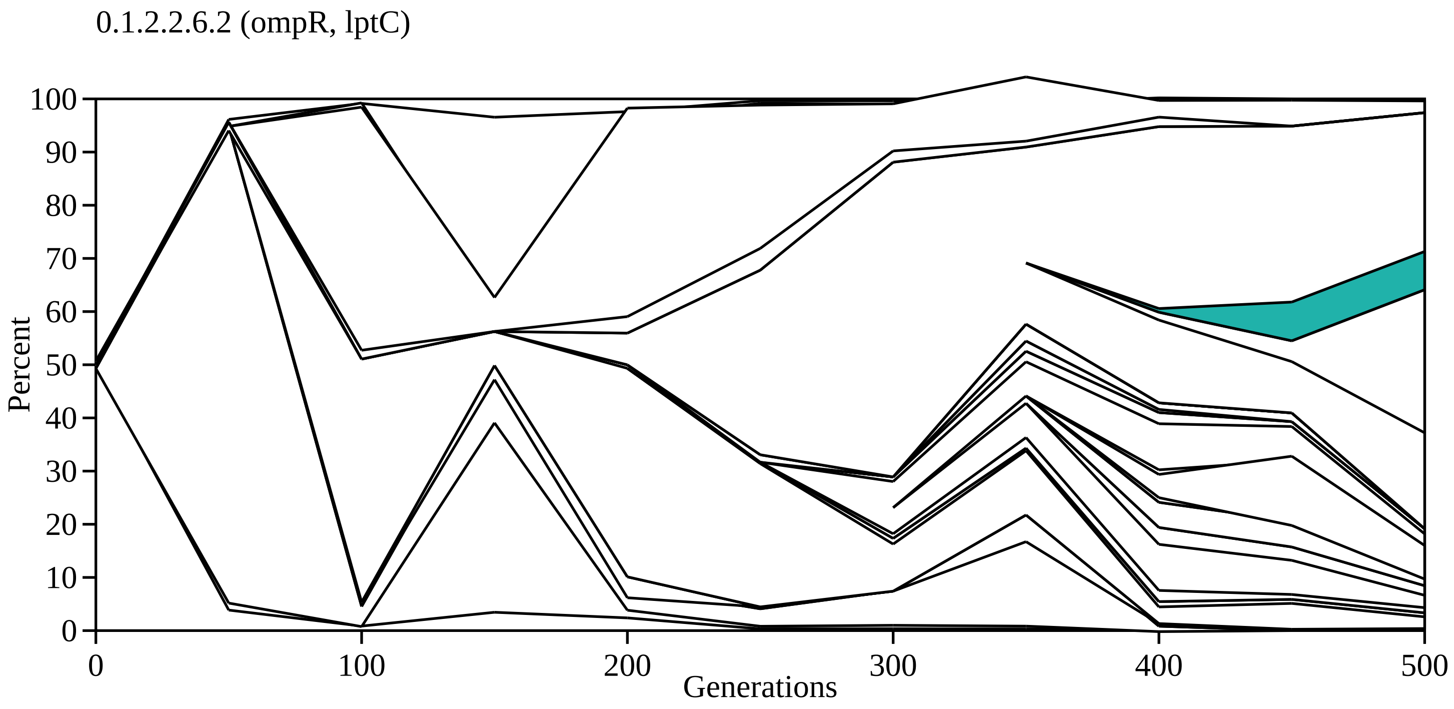

0.1.2.2.6.2 (ompR, lptC)
Percent
100
90
80
70
60
50
40
30
20
10
0
0
100
200
300
400
500
Generations

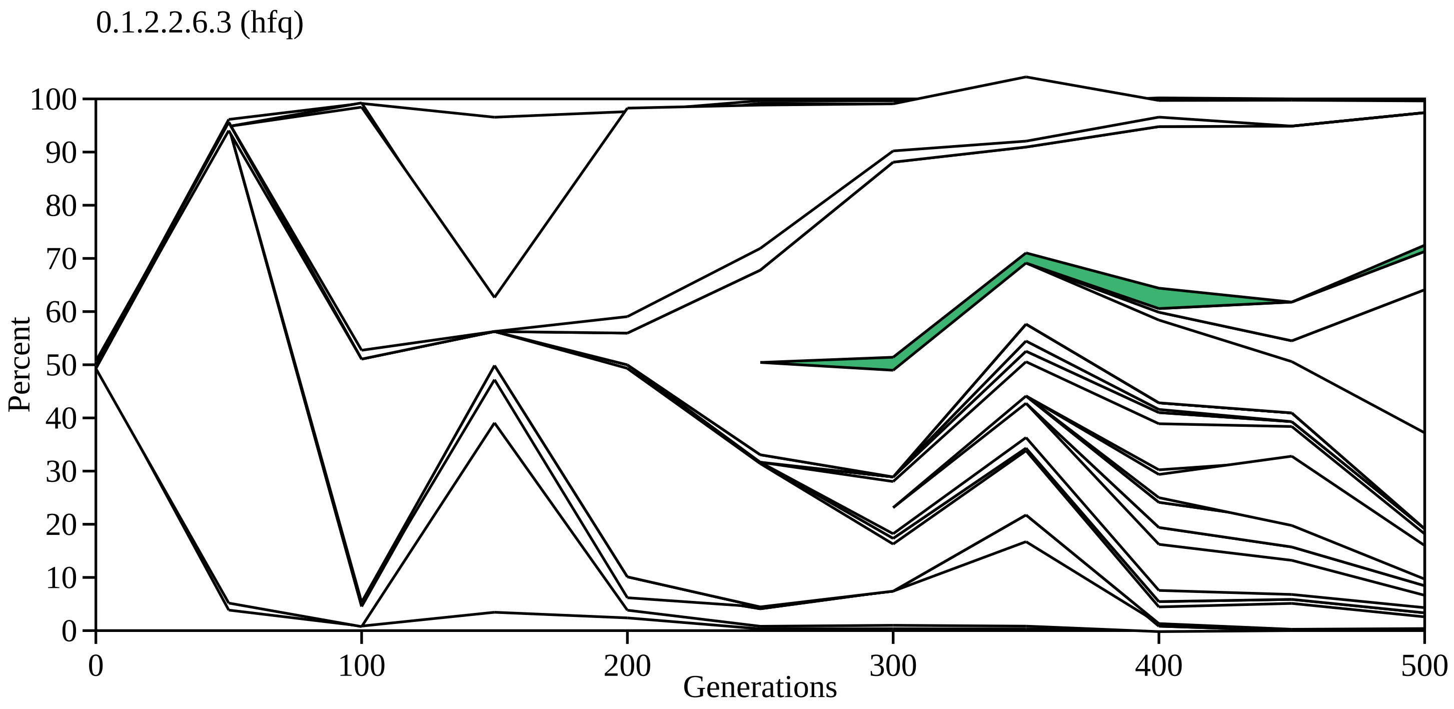
0.1.2.2.6.3 (hfq)
Percent
0
10
20
30
40
50
60
70
80
90
100
0
100
200
300
400
500
Generations

0.1.2.2.6.4 (hfq)

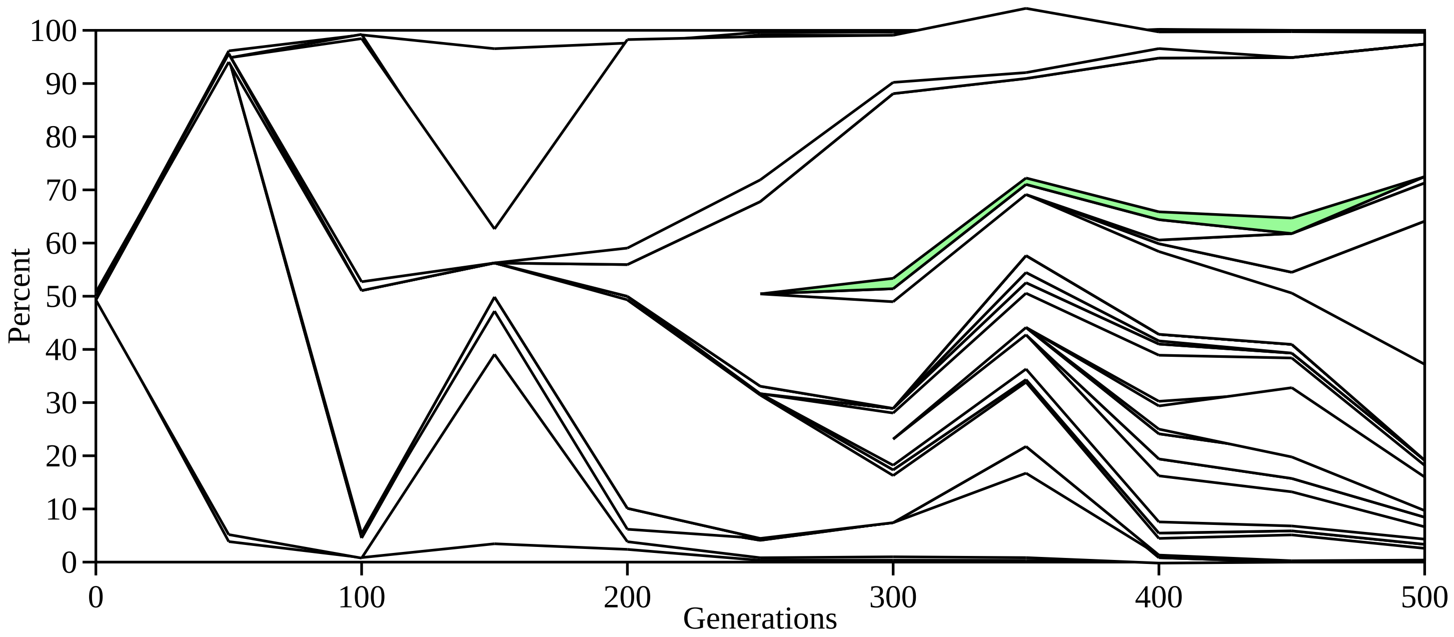

100
90
80
70
60
50
40
30
20
10
0
Percent
0
100
200
300
400
500
Generations

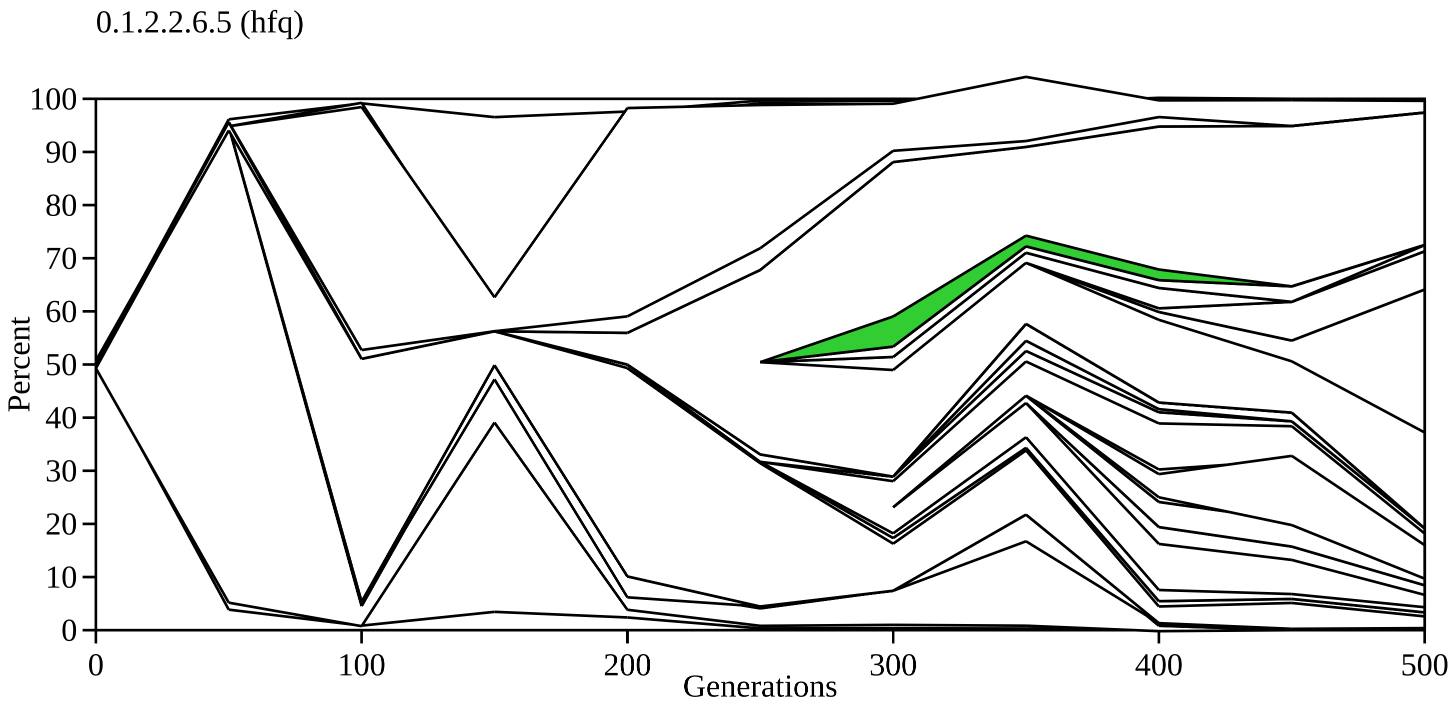

0.1.2.2.6.5 (hfq)
100
90
80
70
60
50
40
30
20
10
0
Percent
0
100
200
300
400
500
Generations

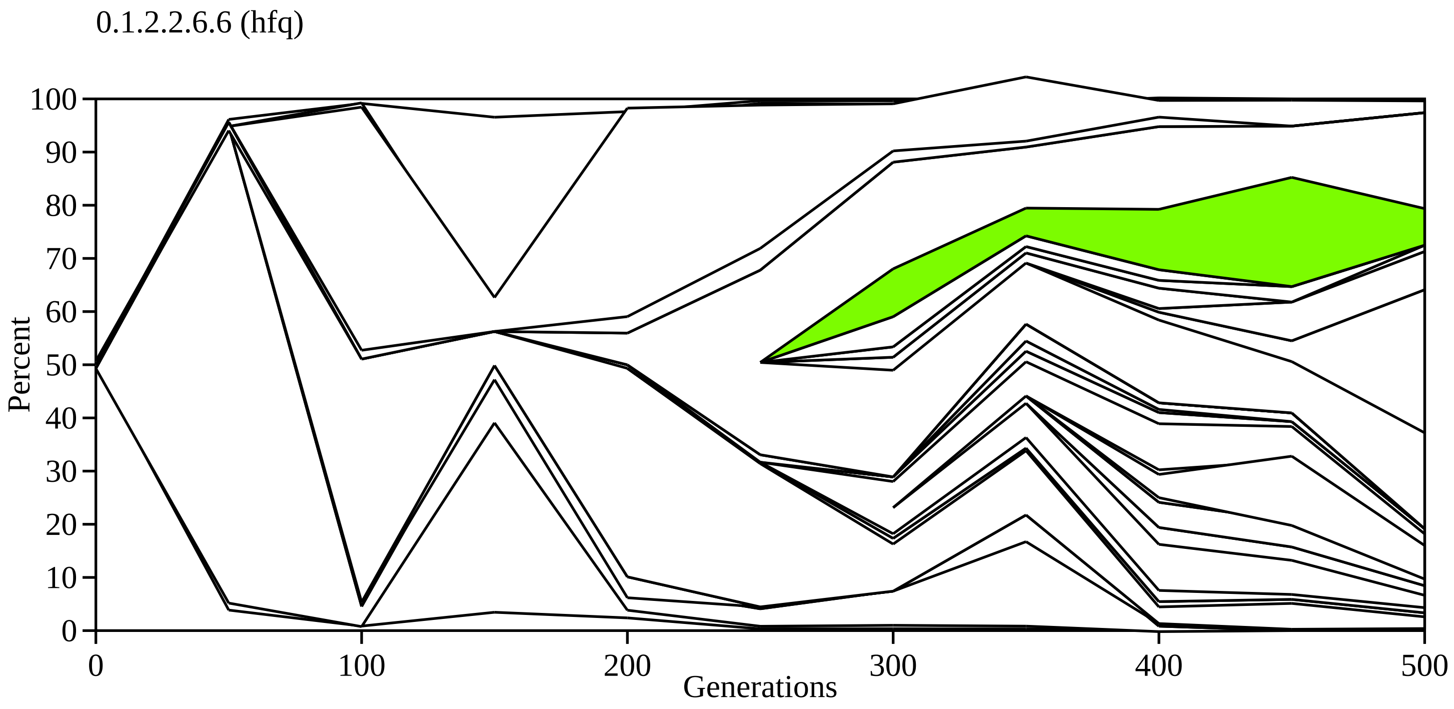
0.1.2.2.6.6 (hfq)
Percent
100
90
80
70
60
50
40
30
20
10
0
0
100
200
300
400
500
Generations

0.1.2.2.6.4.1 (rpoA)

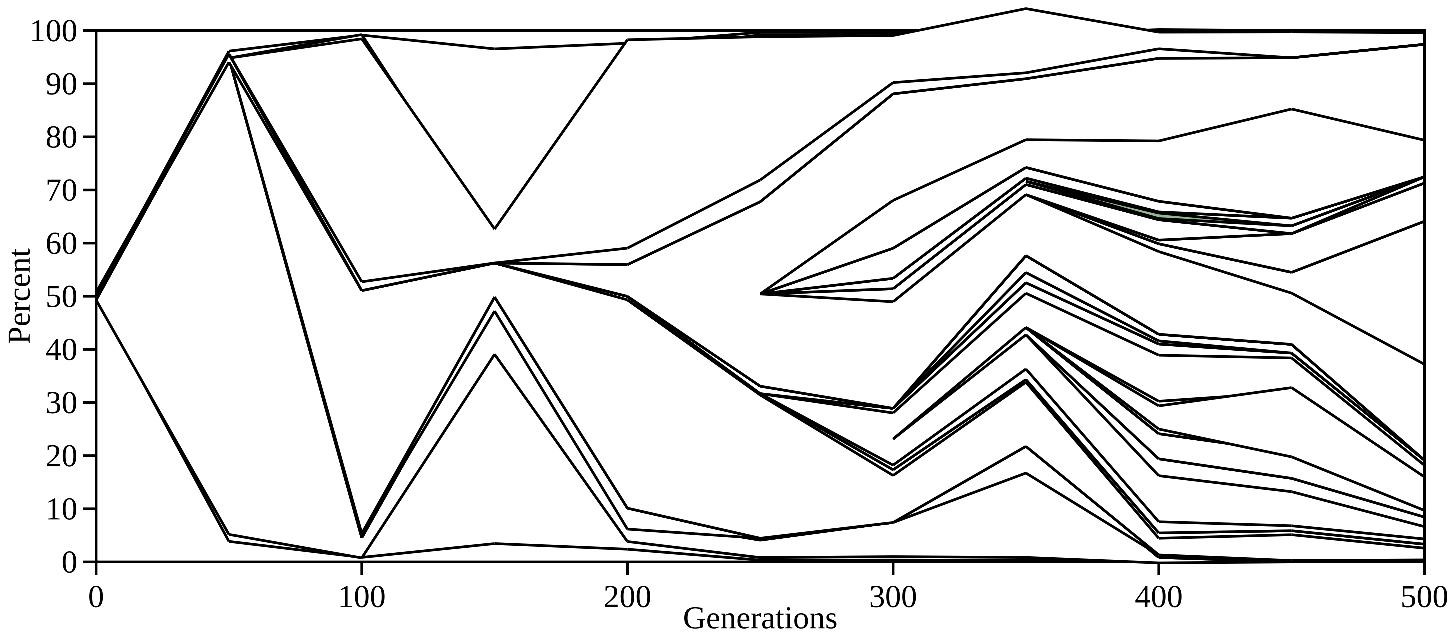

0.1.2.2.6.6.1 (pfkA)

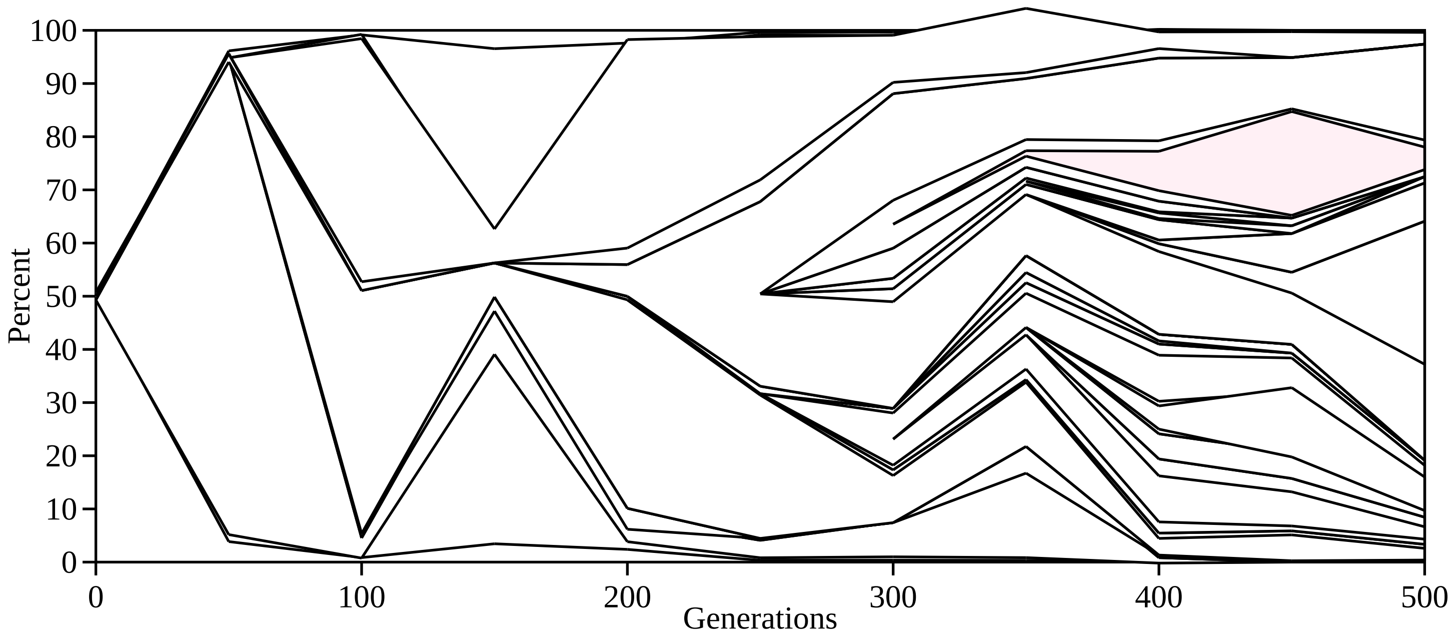
