## Supplementary Data File S4 for "Evolutionary dynamics of *de novo* mutations and mutant lineages arising in a simple, constant environment"

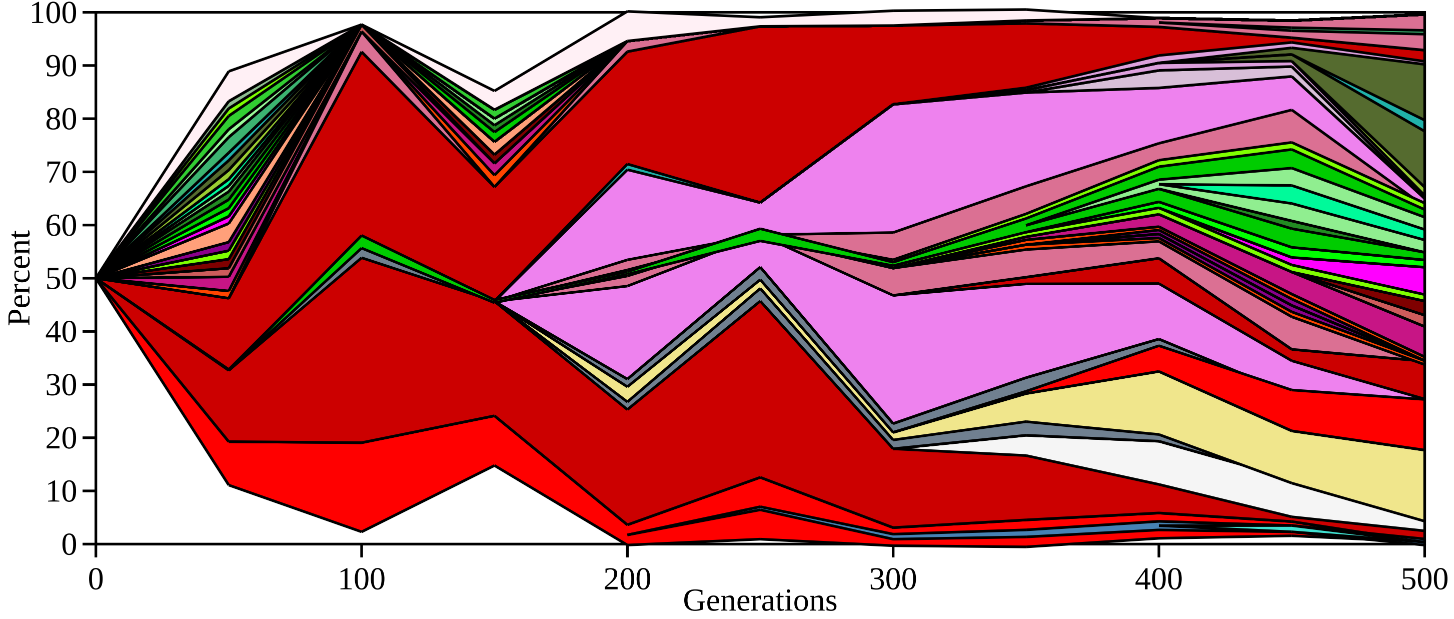
100
90
80
70
60
50
40
30
20
10
0
Percent
0
100
200
300
400
500
Generations

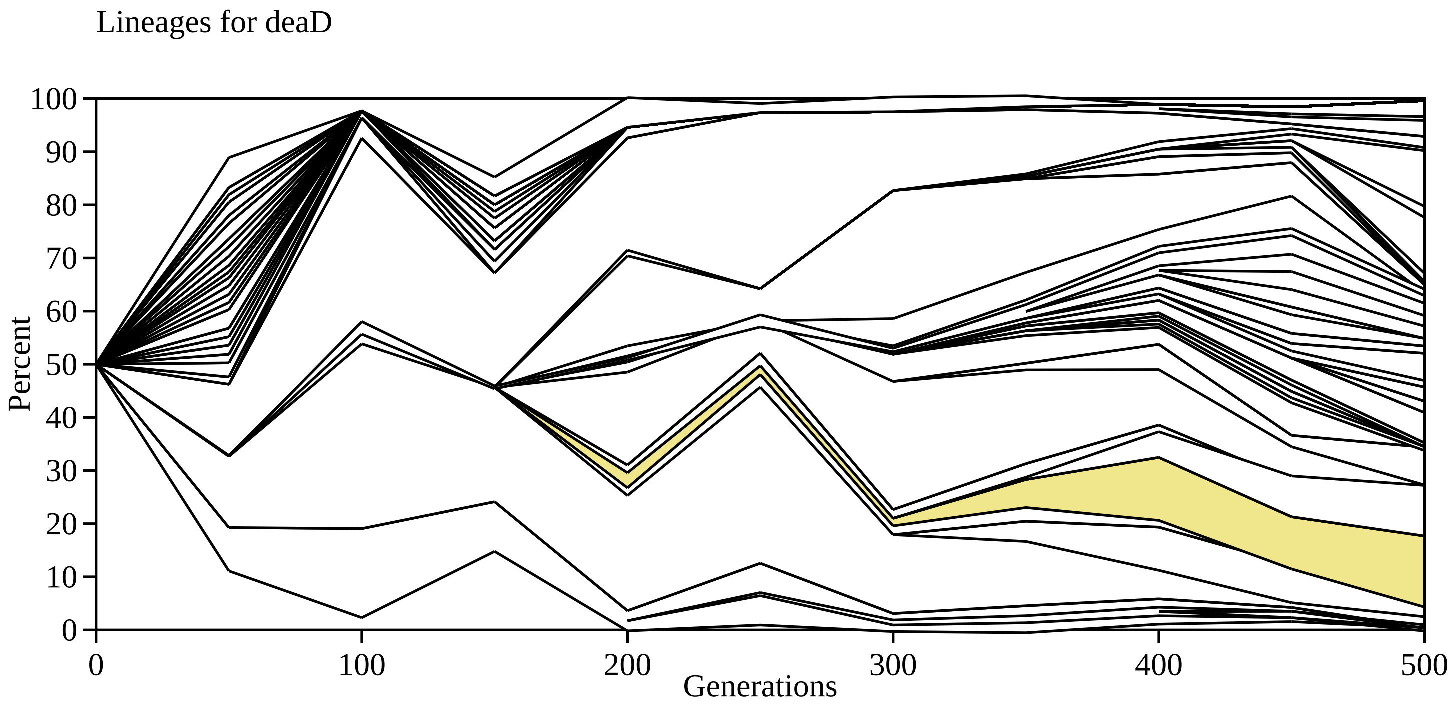

Lineages for deaD
Percent
0
10
20
30
40
50
60
70
80
90
100
0
100
200
300
400
500
Generations

Lineages for downstream fis

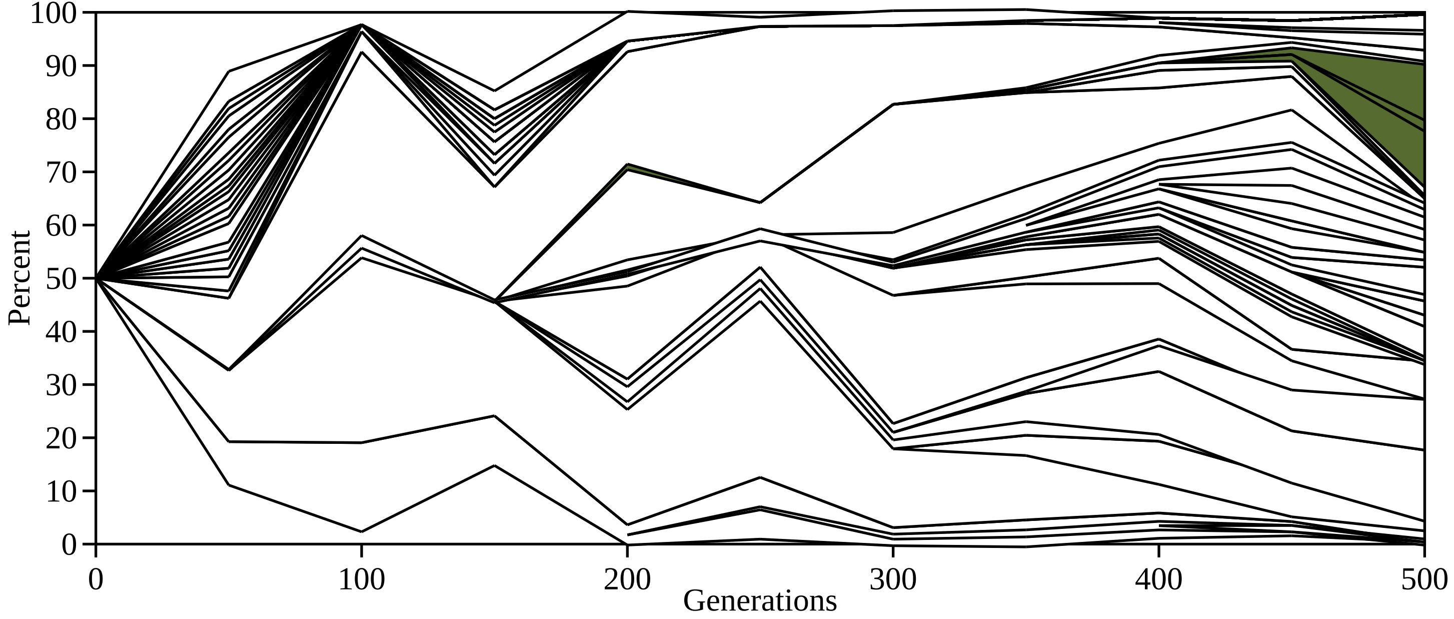

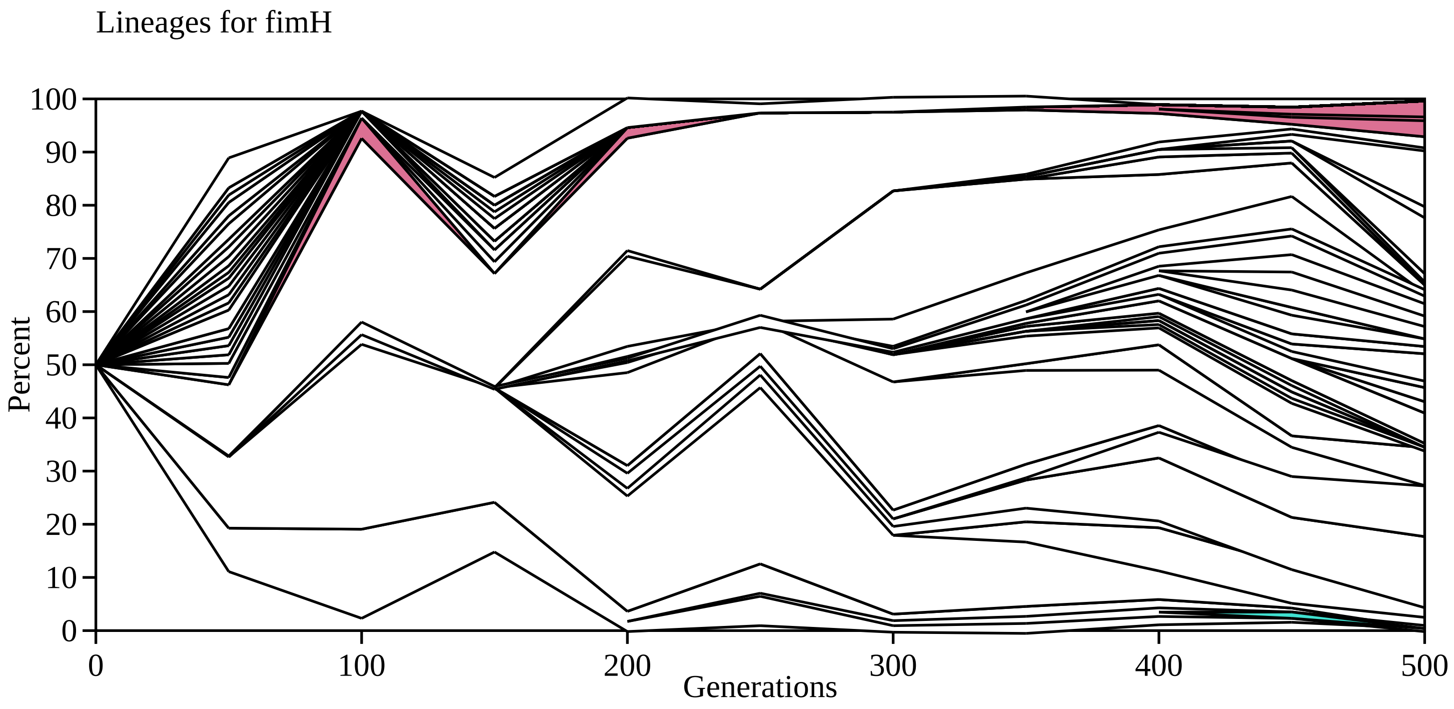
Lineages for fimH
Percent
Generations
0
10
20
30
40
50
60
70
80
90
100
0
100
200
300
400
500

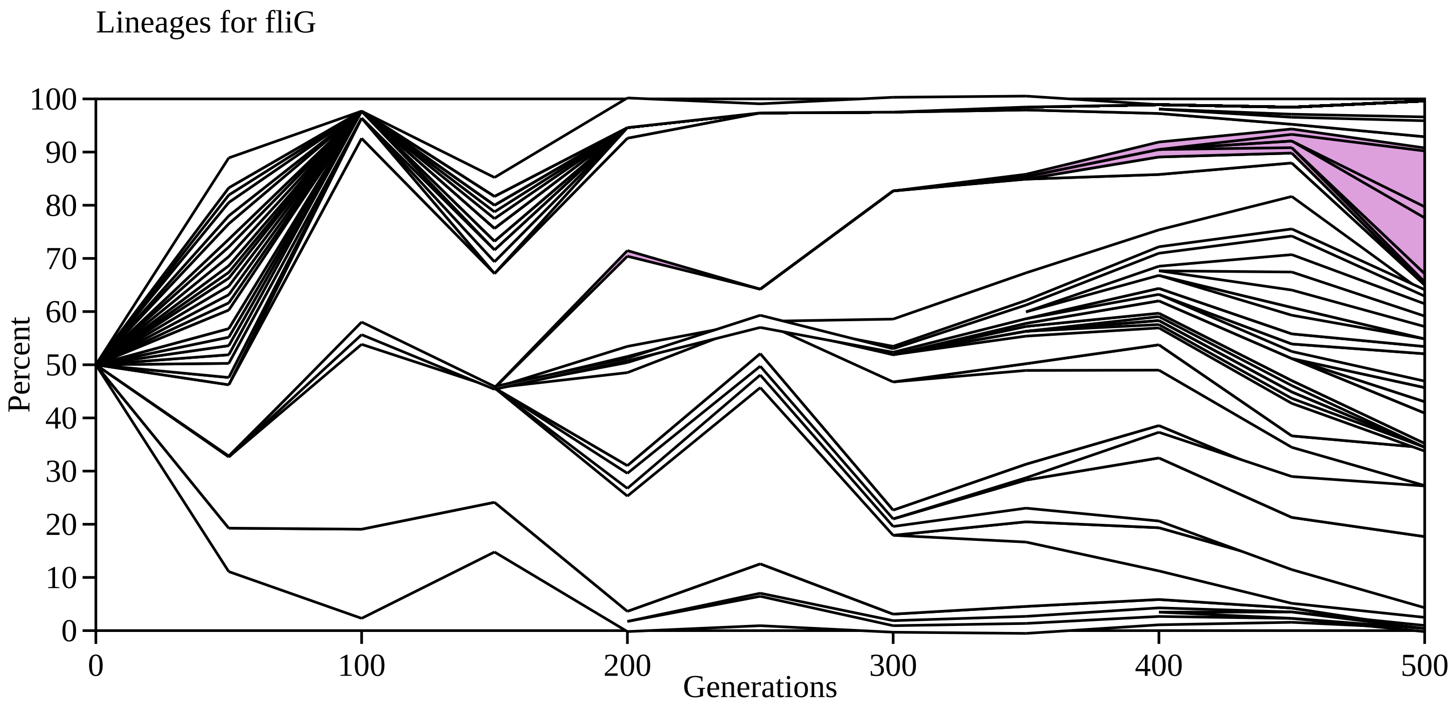

Lineages for fliG
Percent
100
90
80
70
60
50
40
30
20
10
0
0
100
200
300
400
500
Generations

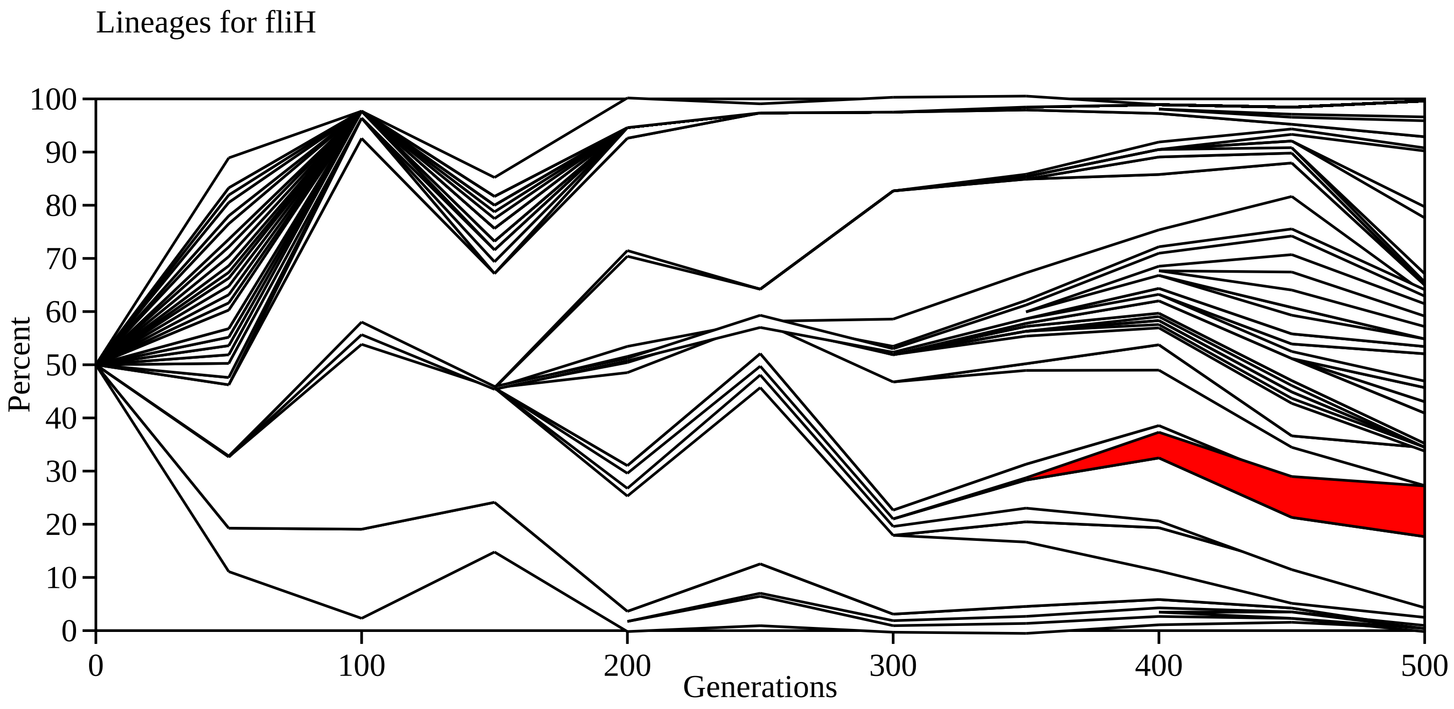

Lineages for fliH
Percent
100
90
80
70
60
50
40
30
20
10
0
0
100
200
300
400
500
Generations

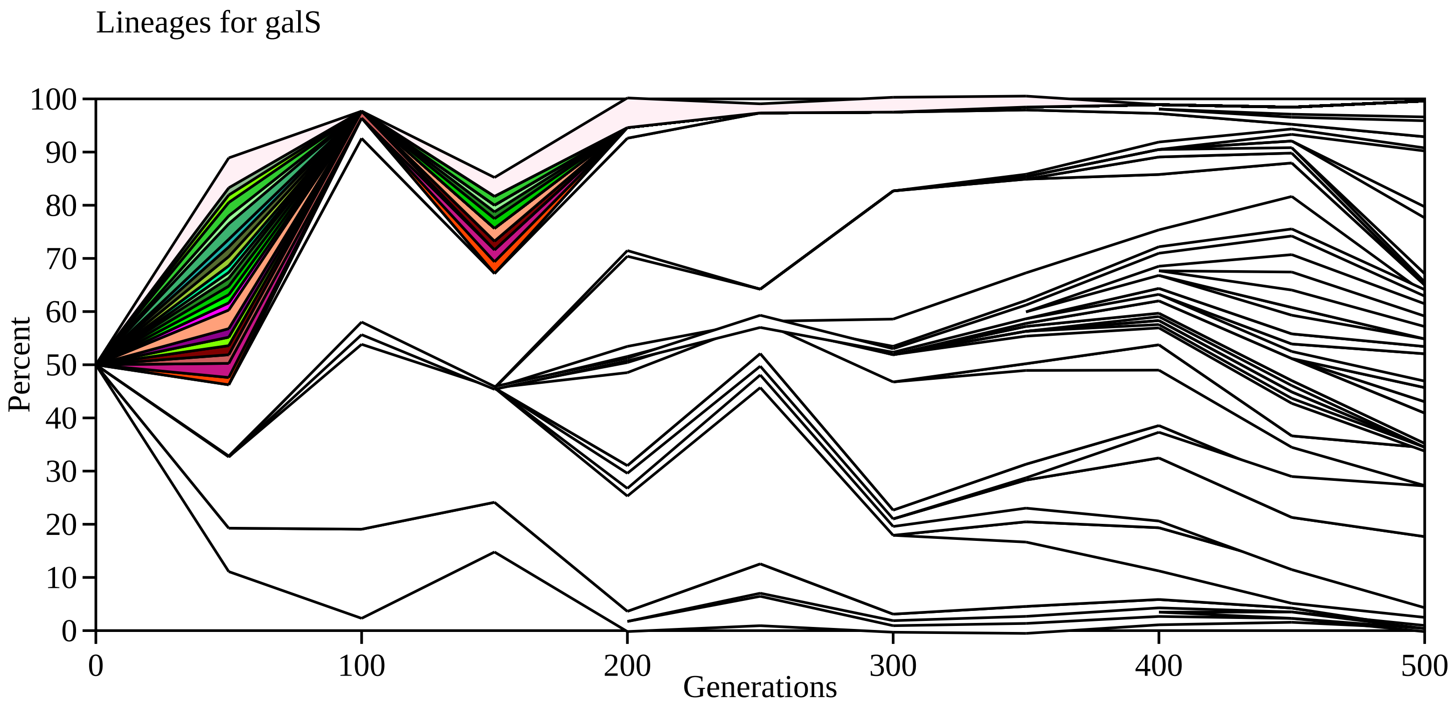
Lineages for galS
Percent
100
90
80
70
60
50
40
30
20
10
0
0
100
200
300
400
500
Generations

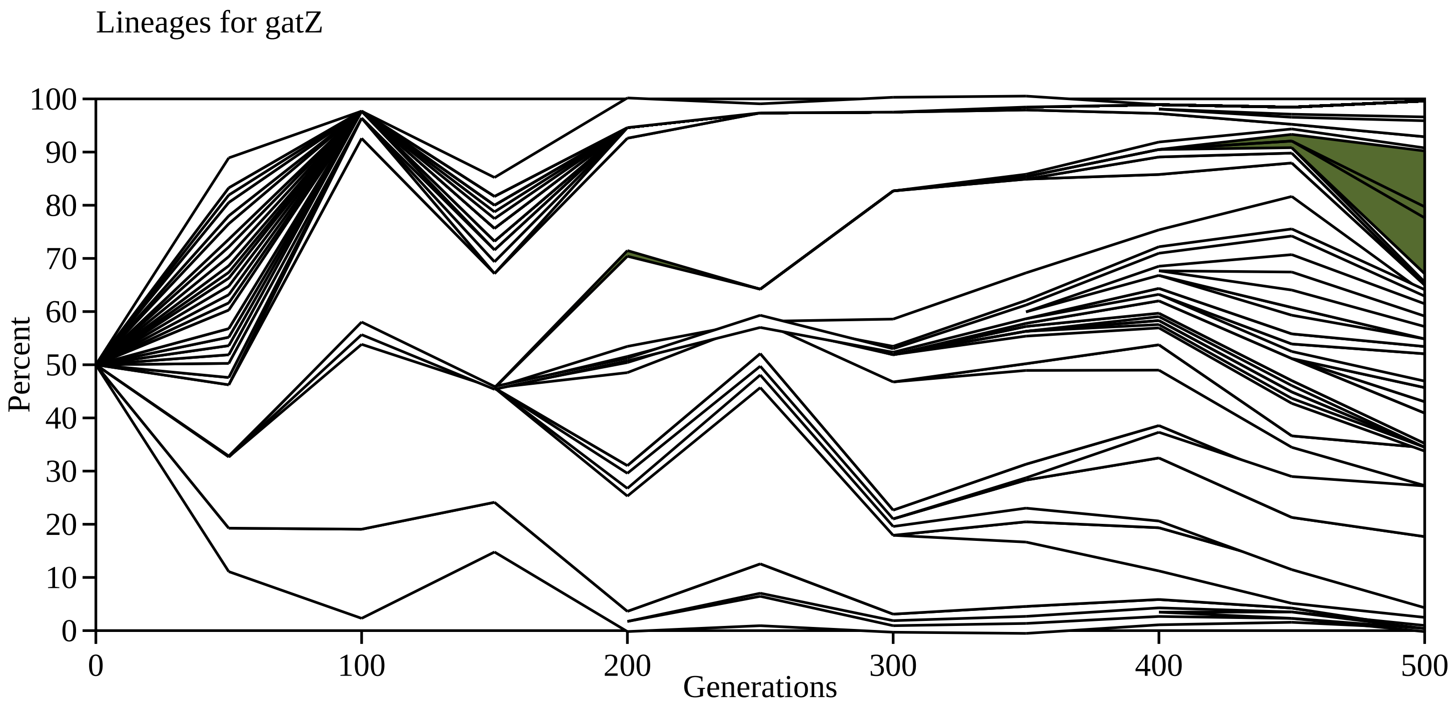

Lineages for gatZ
Percent
0
10
20
30
40
50
60
70
80
90
100
0
100
200
300
400
500
Generations

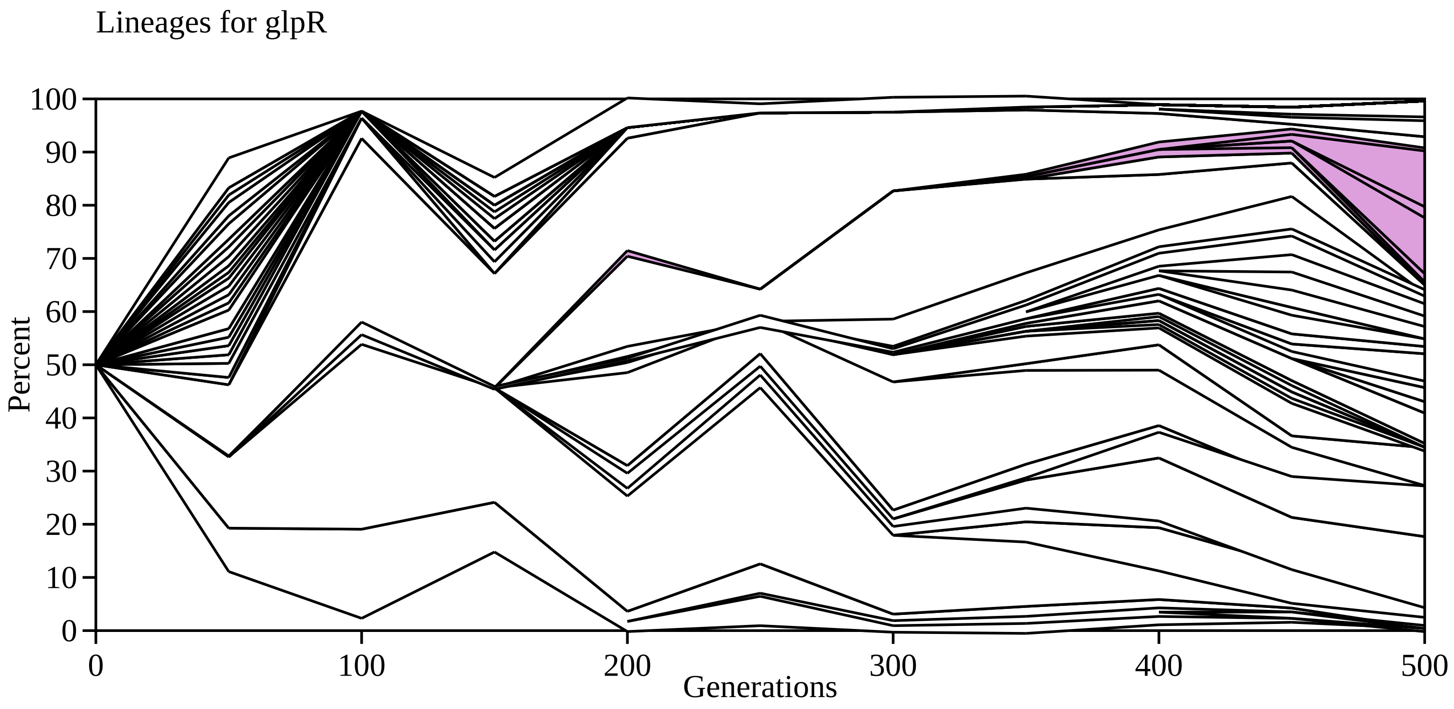
Lineages for glpR
Percent
Generations
0
10
20
30
40
50
60
70
80
90
100
0
100
200
300
400
500

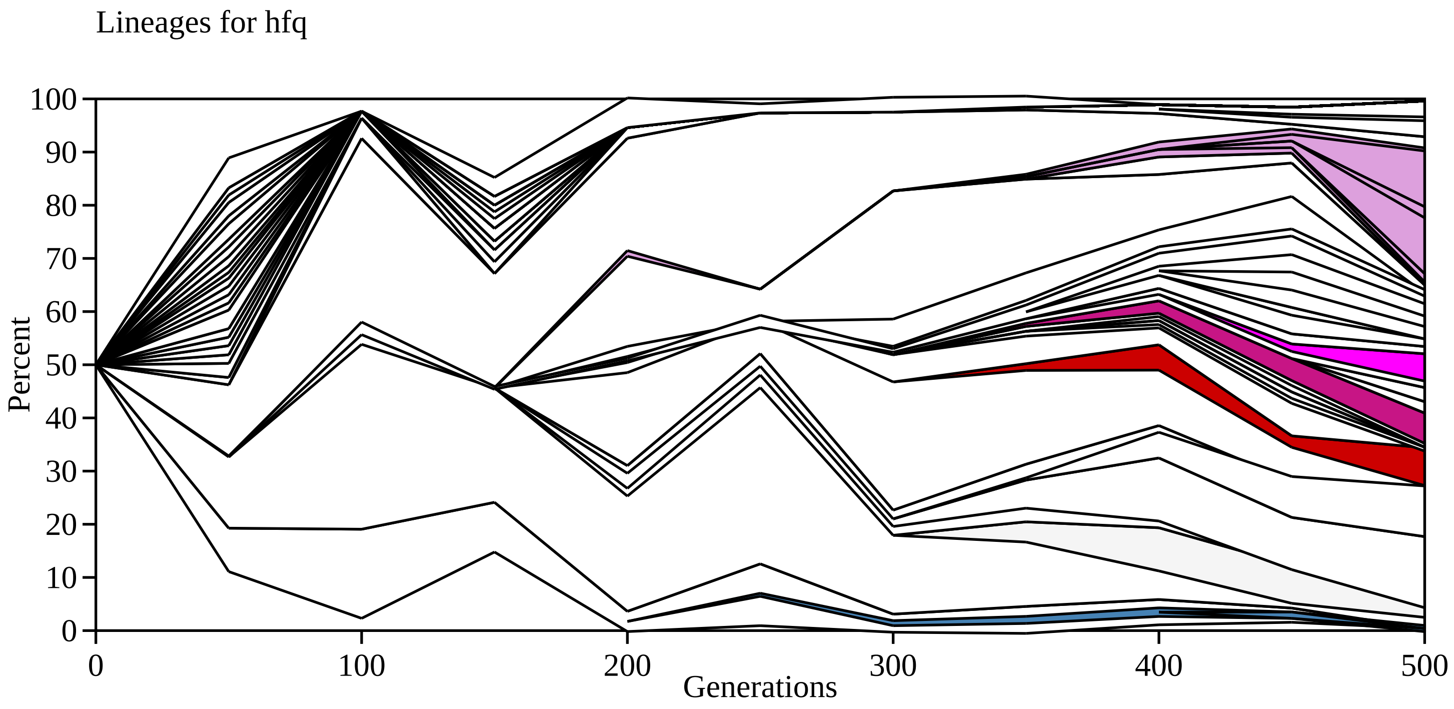
Lineages for hfq
Percent
100
90
80
70
60
50
40
30
20
10
0
0
100
200
300
400
500
Generations

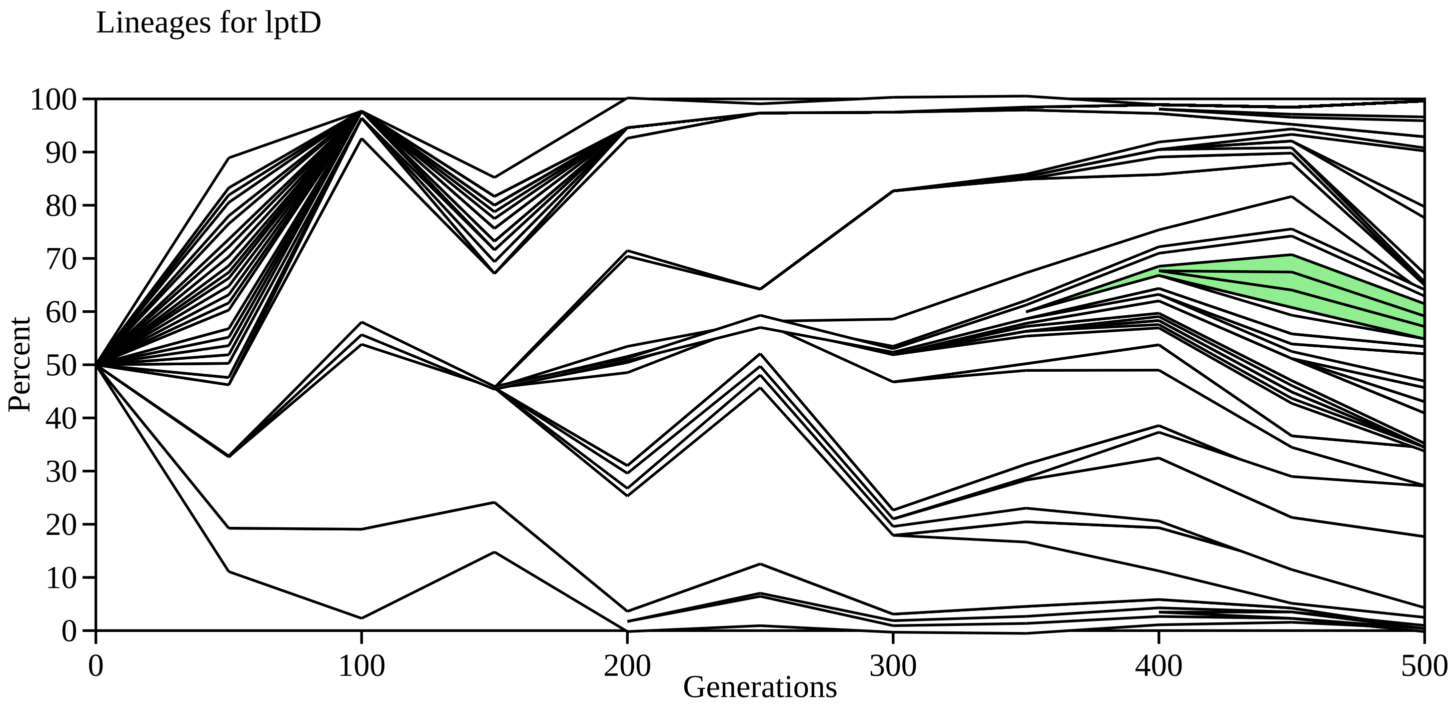

Lineages for lptD
Percent
0
10
20
30
40
50
60
70
80
90
100
0
100
200
300
400
500
Generations

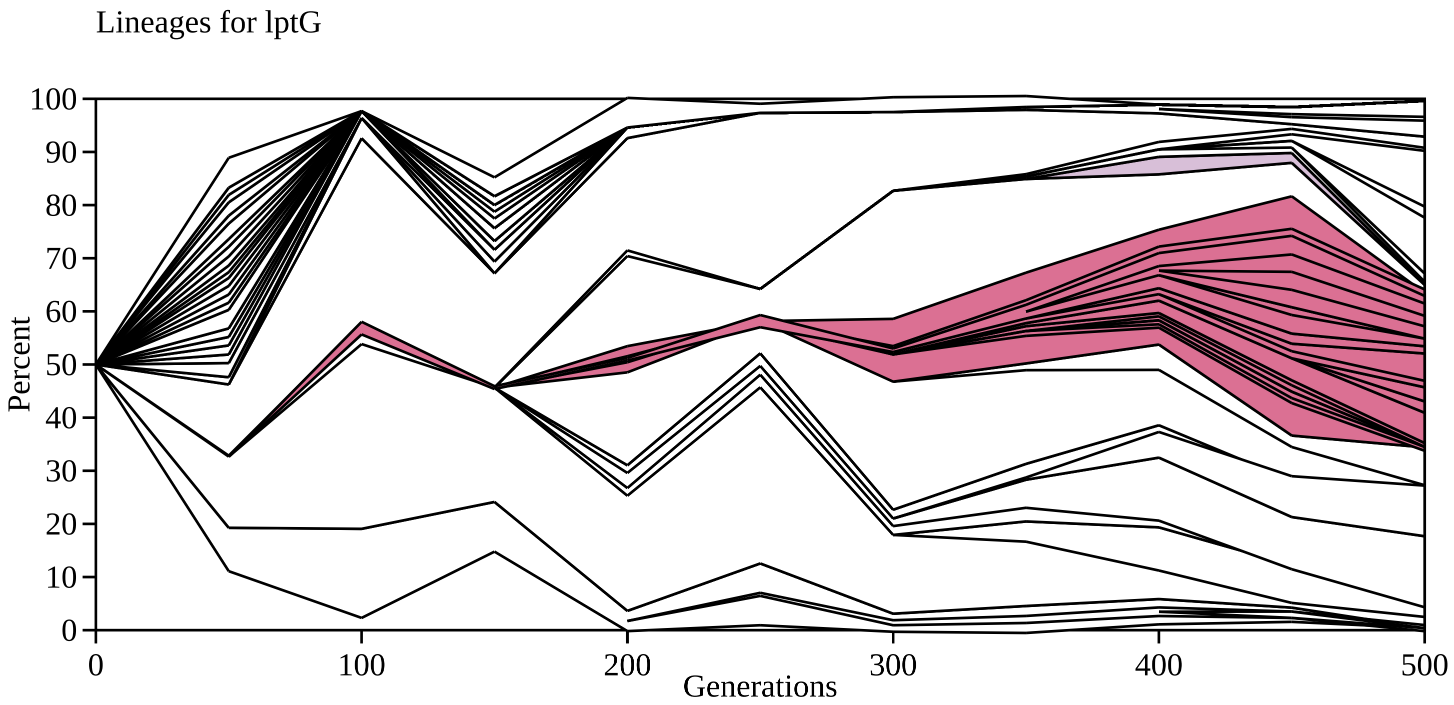

Lineages for lptG
Percent
Generations
100
90
80
70
60
50
40
30
20
10
0
0
100
200
300
400
500

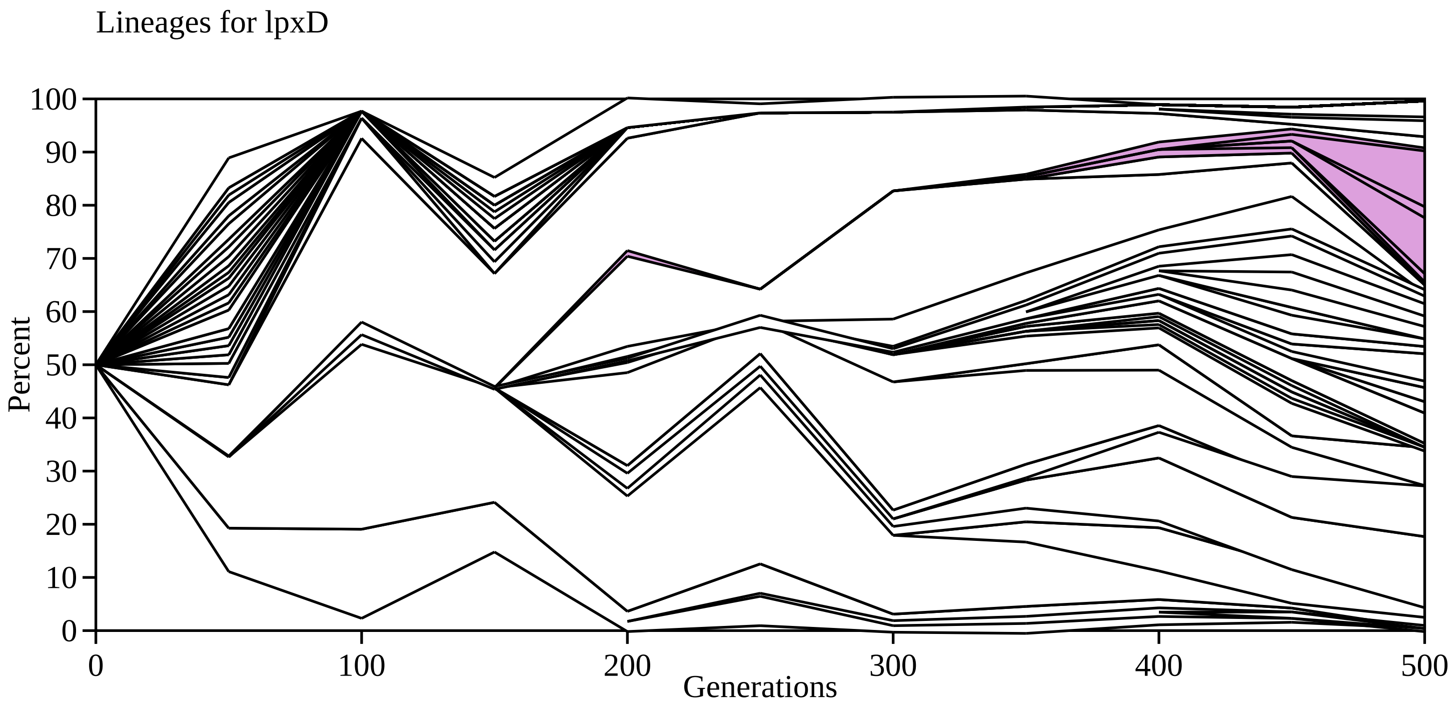

Lineages for lpxD
Percent
100
90
80
70
60
50
40
30
20
10
0
0
100
200
300
400
500
Generations

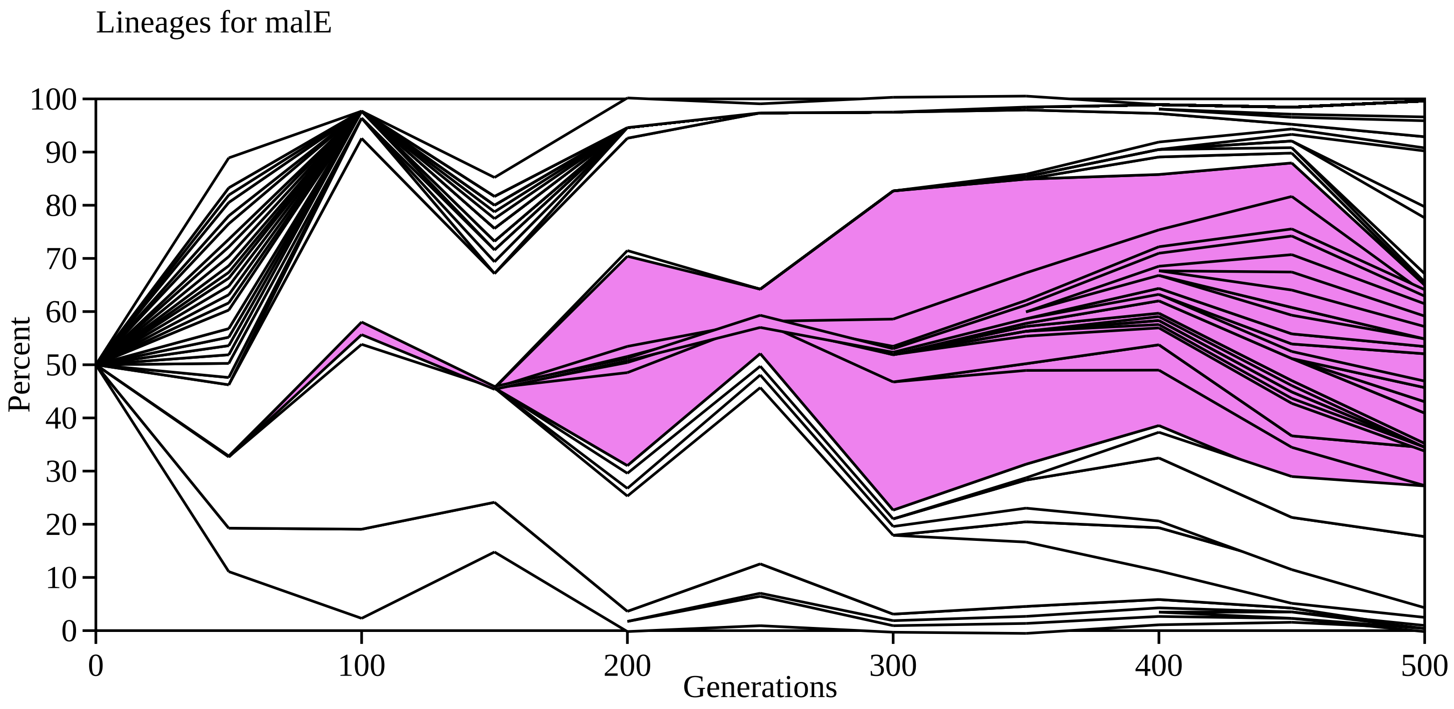

Lineages for malE
Percent
100
90
80
70
60
50
40
30
20
10
0
0
100
200
300
400
500
Generations

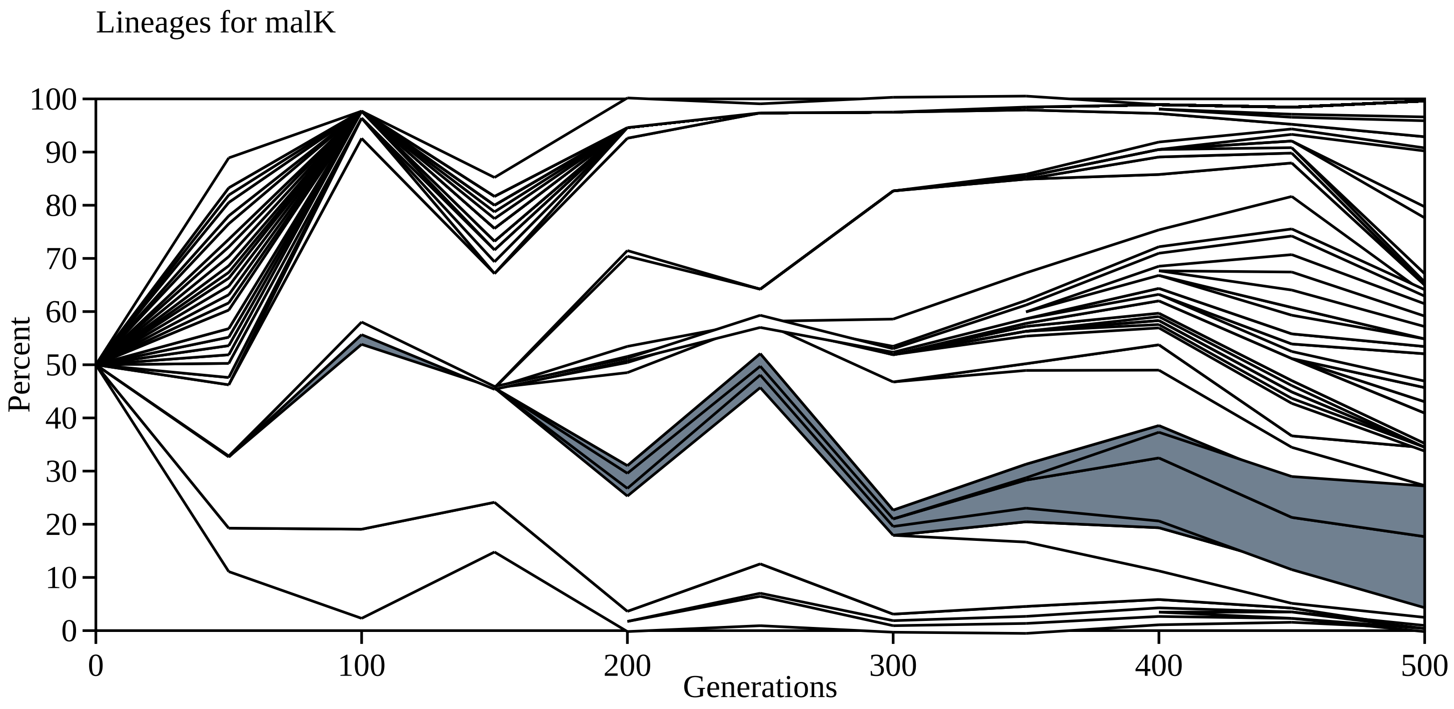
Lineages for malK
Percent
100
90
80
70
60
50
40
30
20
10
0
0
100
200
300
400
500
Generations

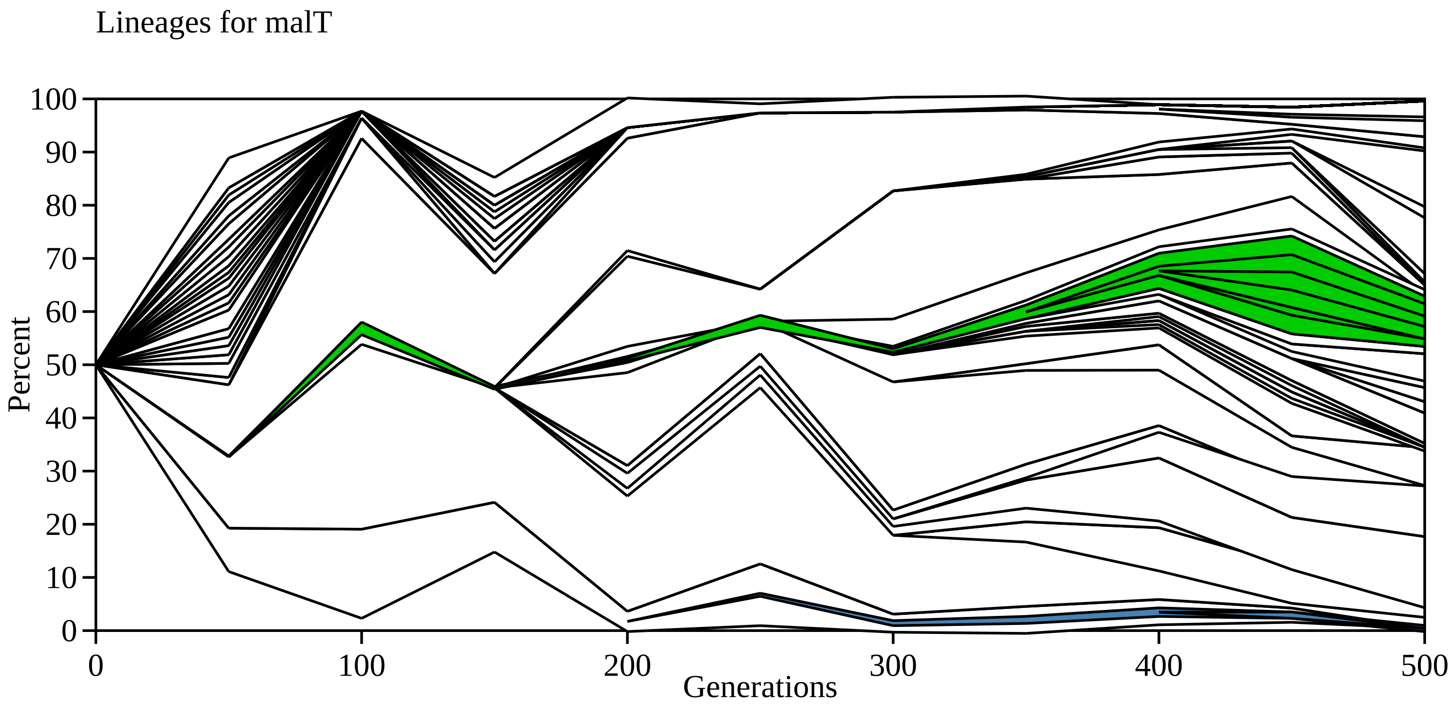

Lineages for malT
Percent
0
10
20
30
40
50
60
70
80
90
100
0
100
200
300
400
500
Generations

Lineages for ompR

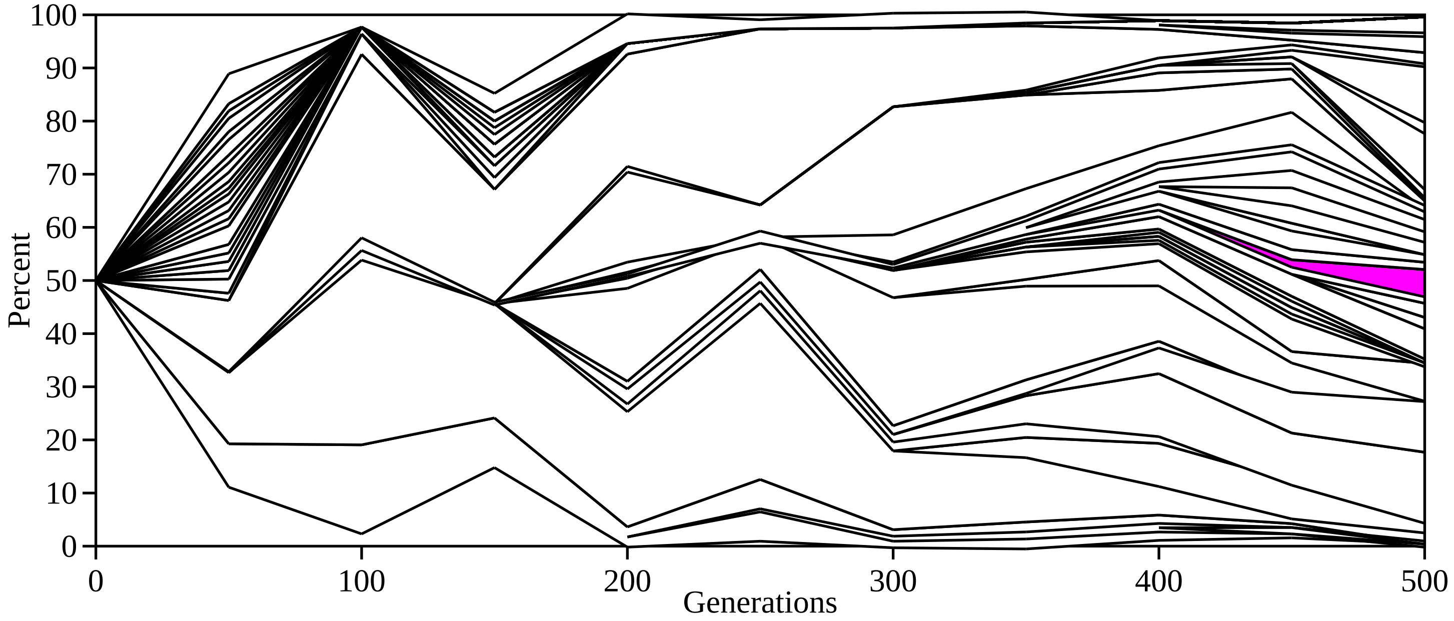

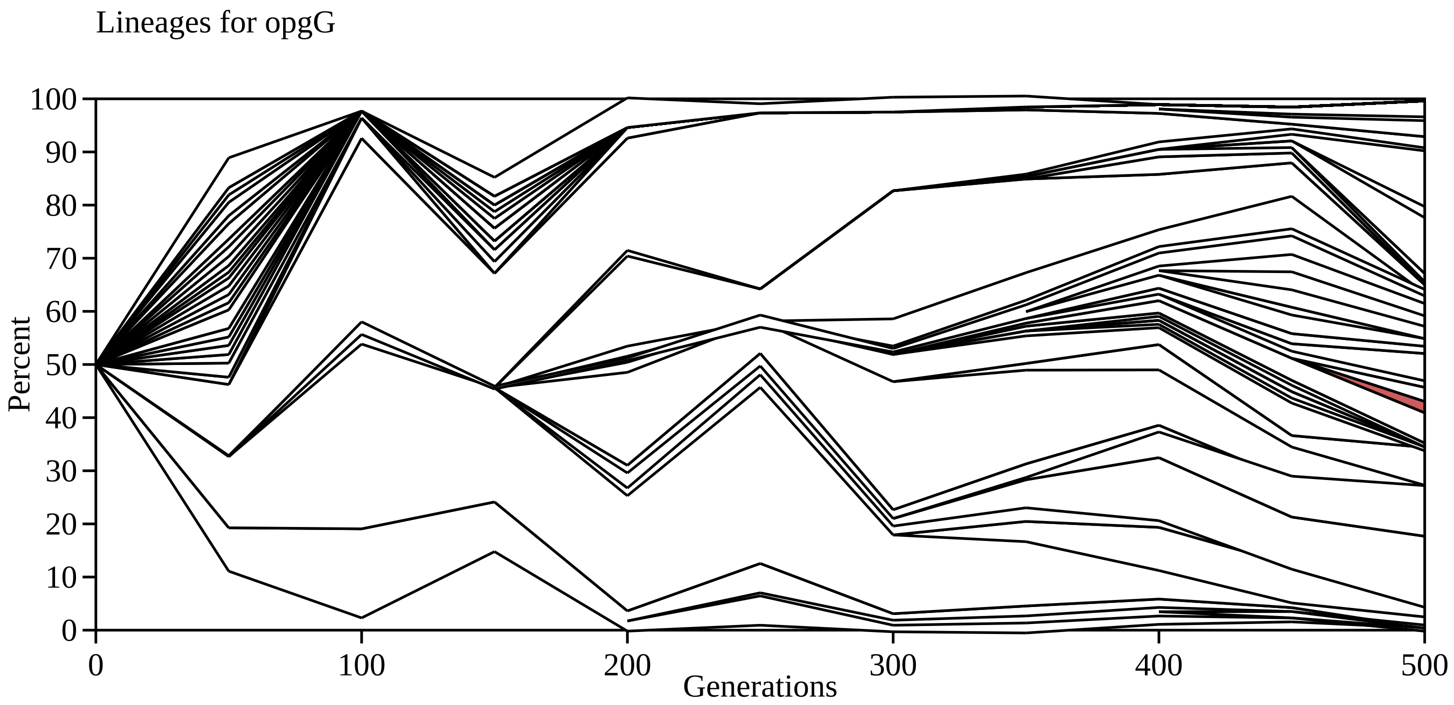

Lineages for opgG
Percent
0
10
20
30
40
50
60
70
80
90
100
0
100
200
300
400
500
Generations

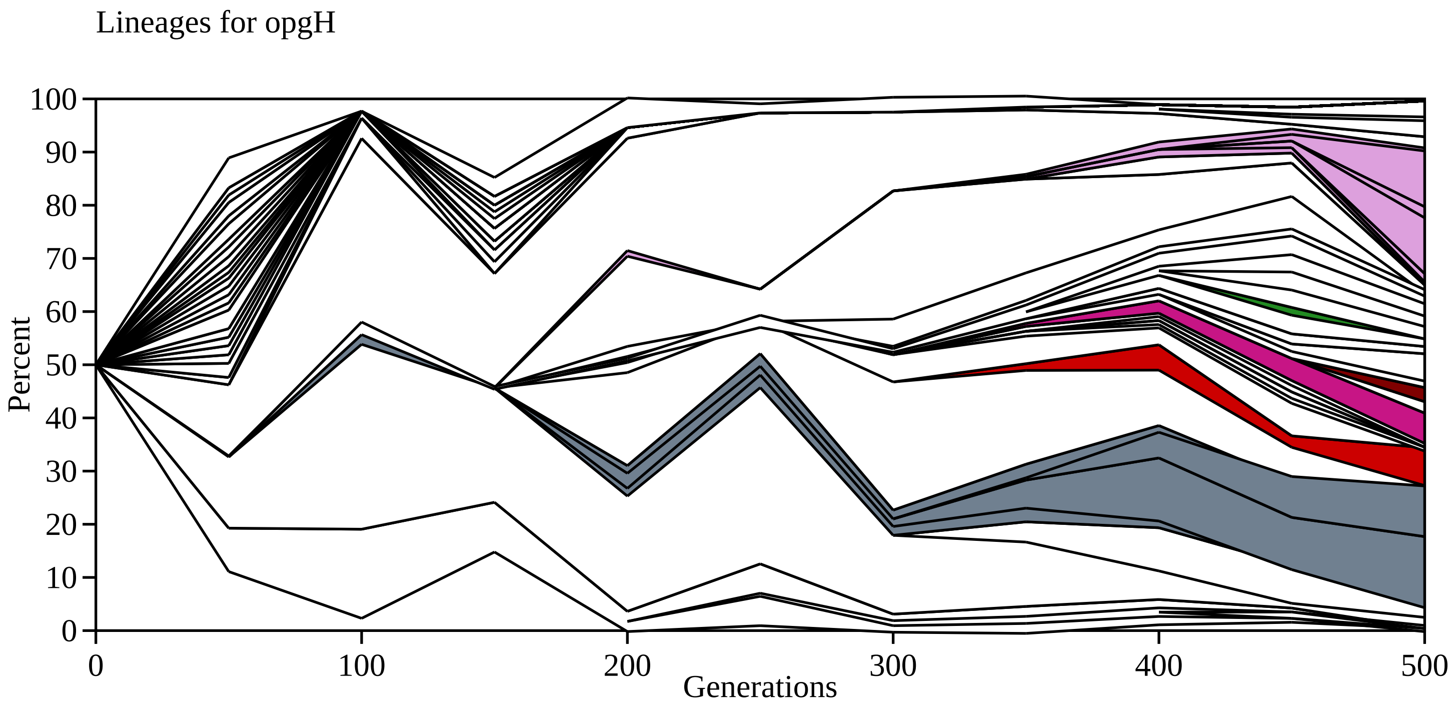

Lineages for opgH
Percent
Generations
0
10
20
30
40
50
60
70
80
90
100
0
100
200
300
400
500

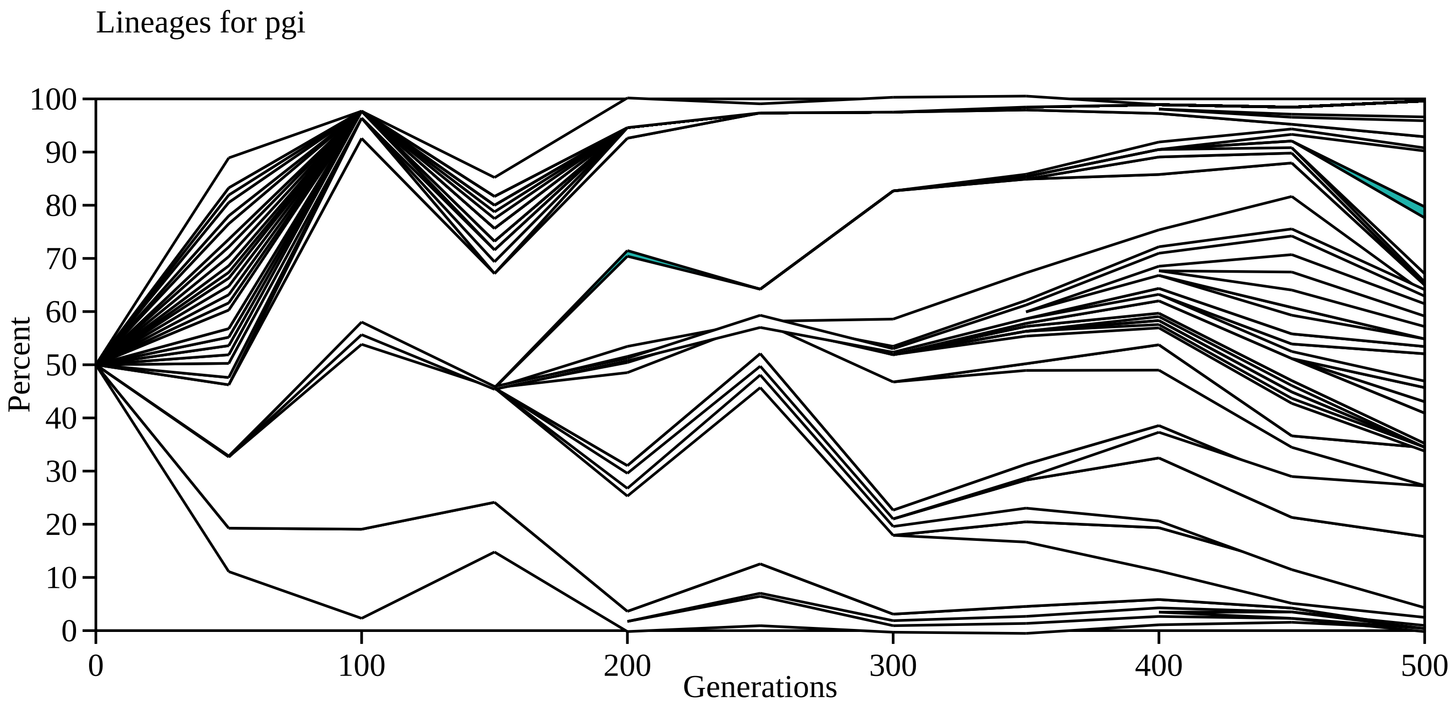

Lineages for pgi
Percent
Generations
0
10
20
30
40
50
60
70
80
90
100
0
100
200
300
400
500

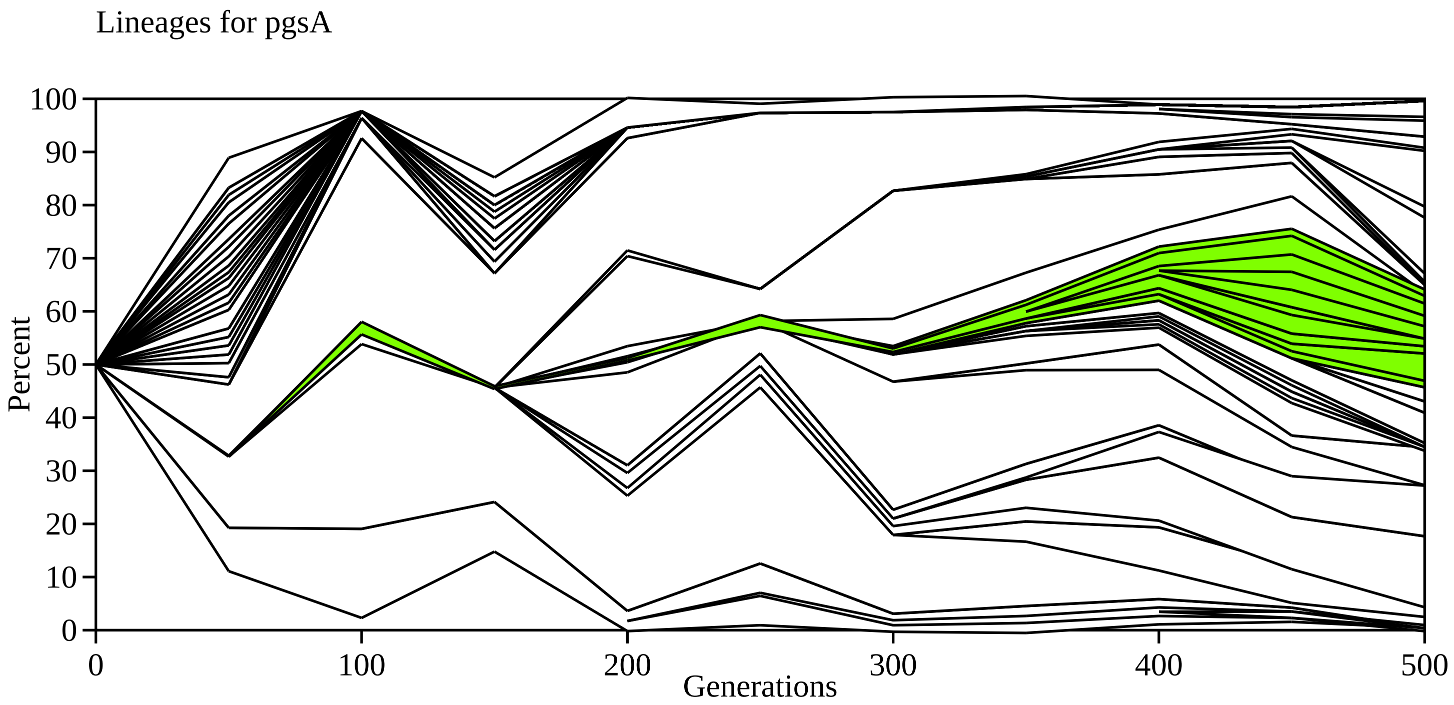
Lineages for pgsA
Percent
Generations
0
10
20
30
40
50
60
70
80
90
100
0
100
200
300
400
500

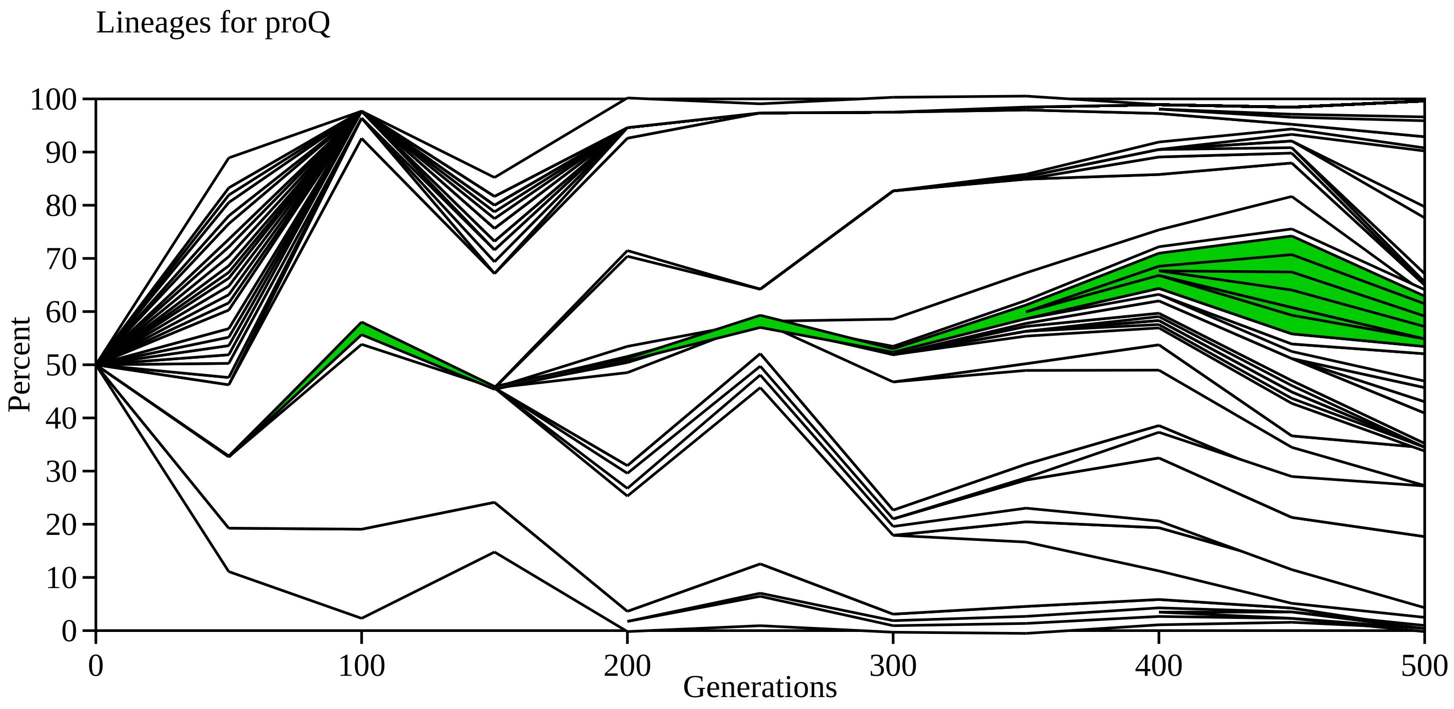

Lineages for proQ
100
90
80
70
60
50
40
30
20
10
0
Percent
0
100
200
300
400
500
Generations

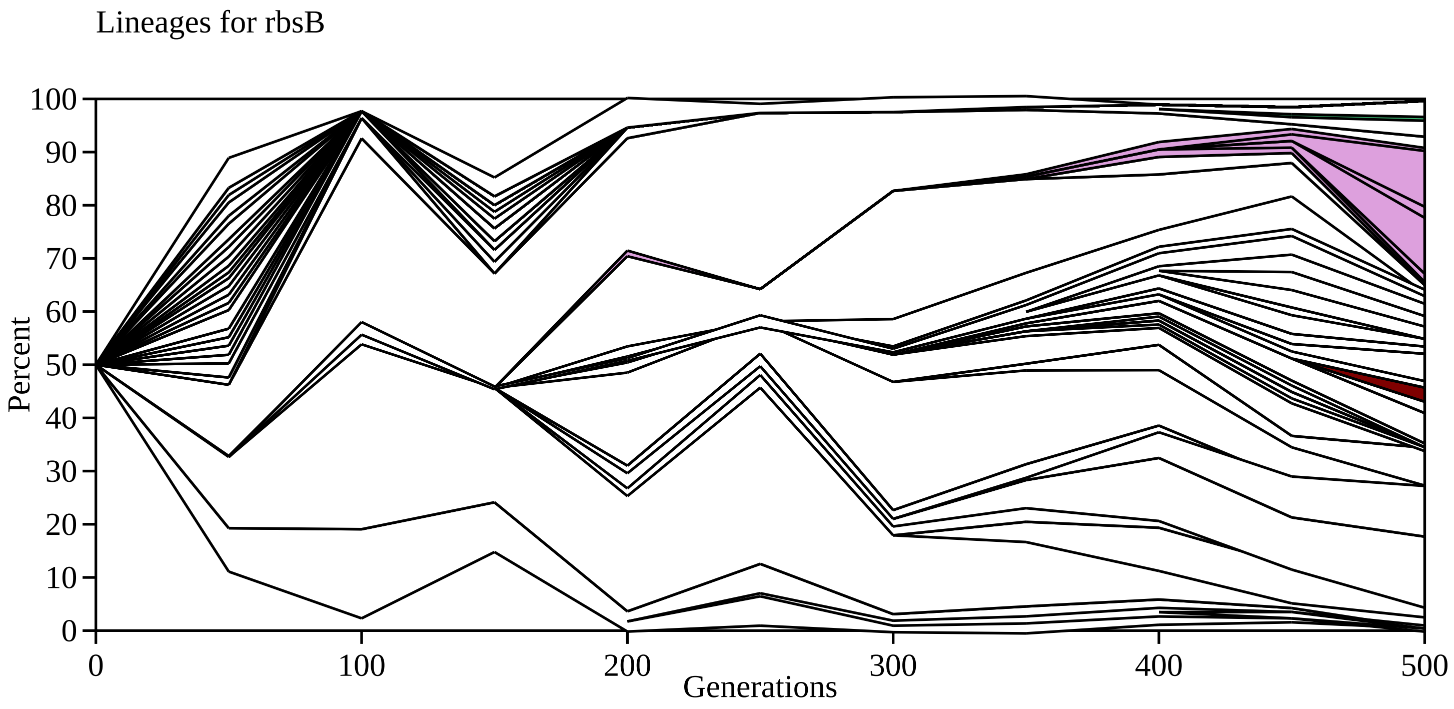

Lineages for rbsB
Percent
0
10
20
30
40
50
60
70
80
90
100
0
100
200
300
400
500
Generations

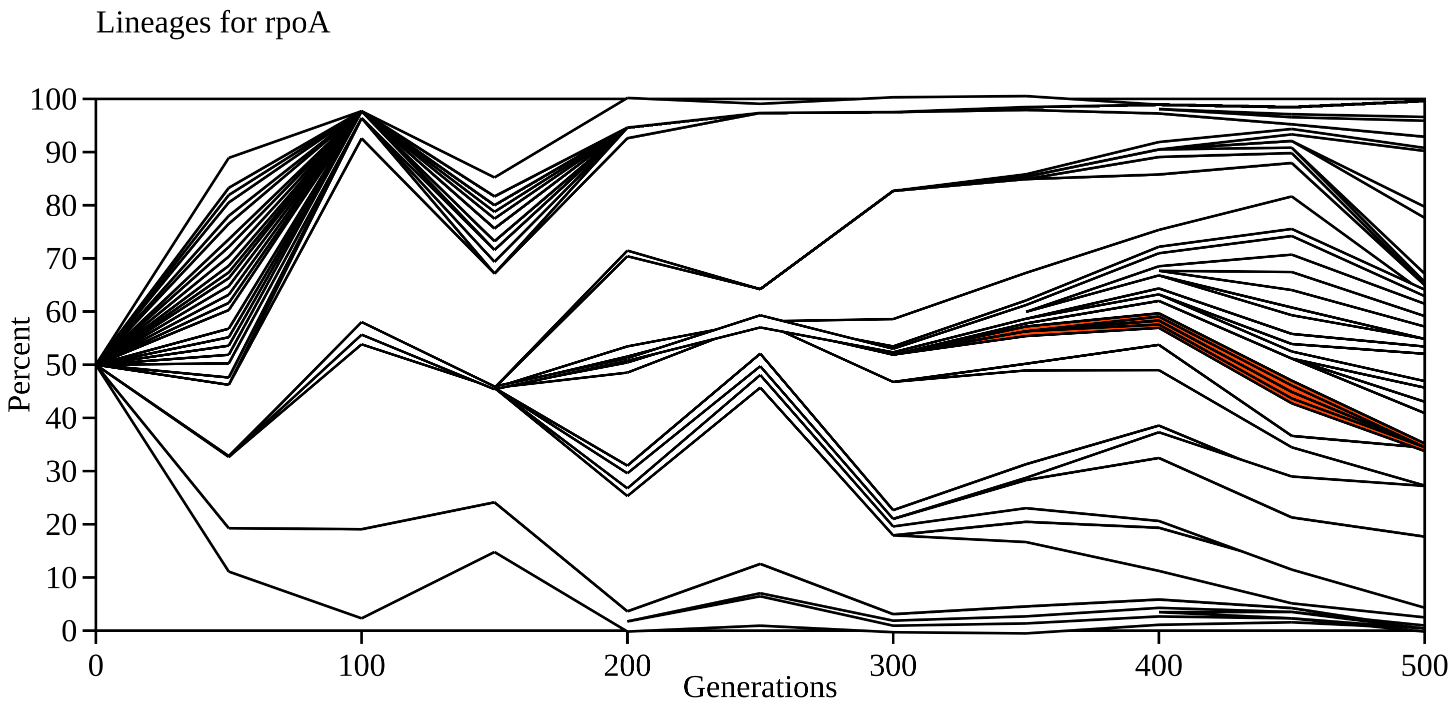

Lineages for rpoA
Percent
100
90
80
70
60
50
40
30
20
10
0
0
100
200
300
400
500
Generations

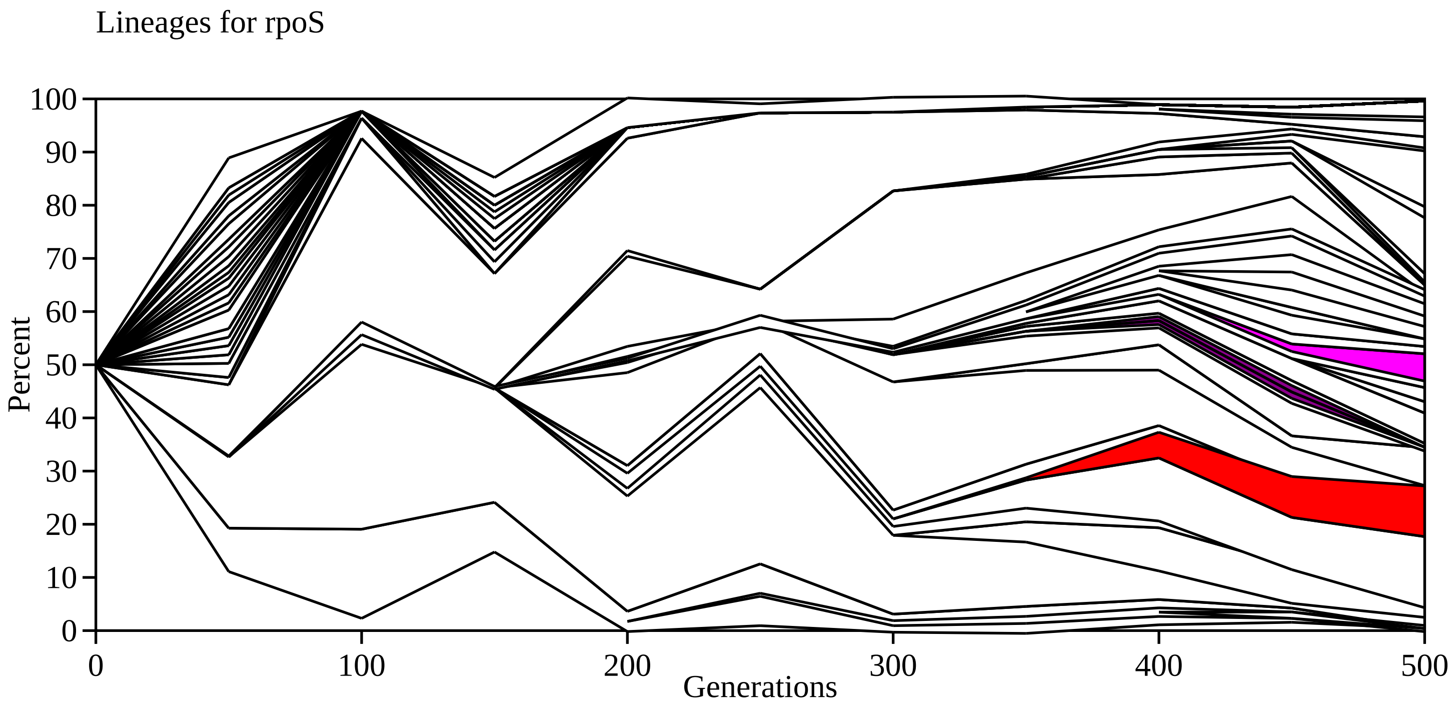

Lineages for rpoS
100
90
80
70
60
50
40
30
20
10
0
Percent
0
100
200
300
400
500
Generations

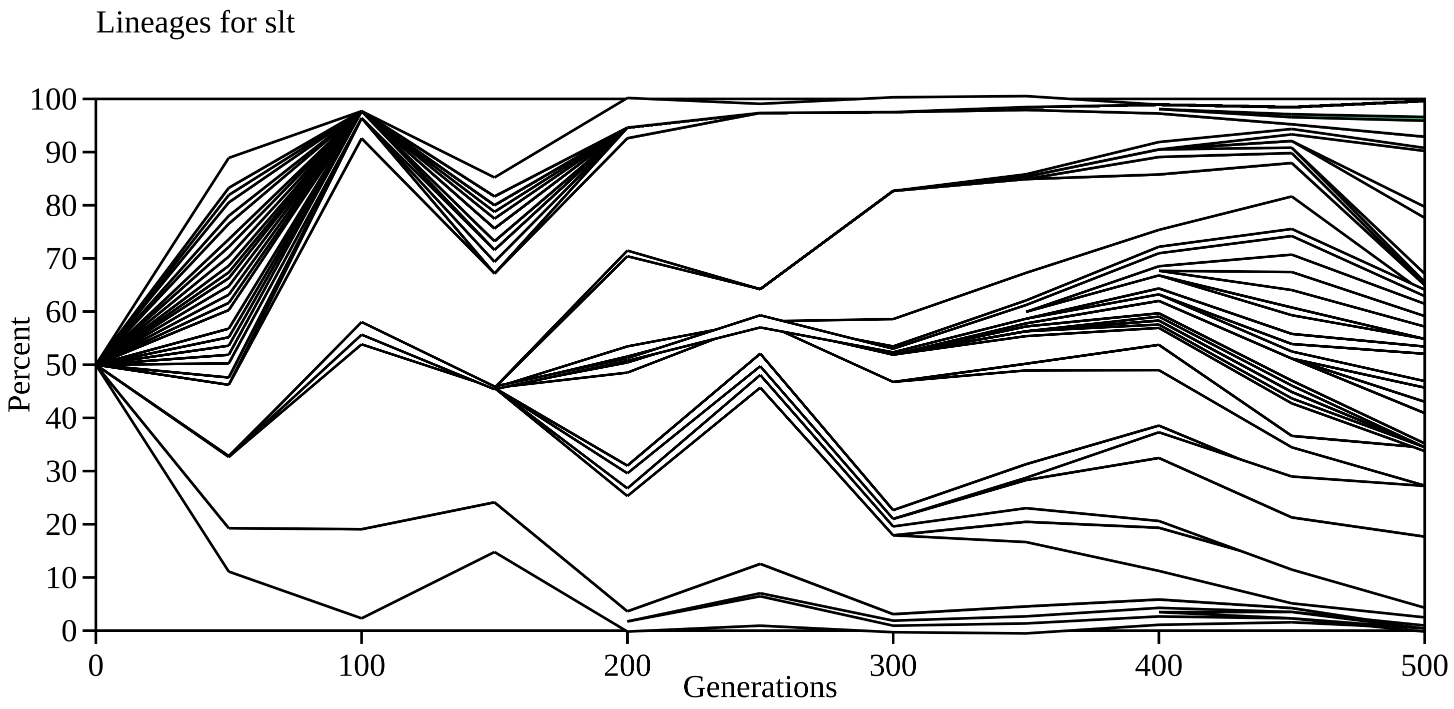
Lineages for slt
Percent
100
90
80
70
60
50
40
30
20
10
0
0
100
200
300
400
500
Generations

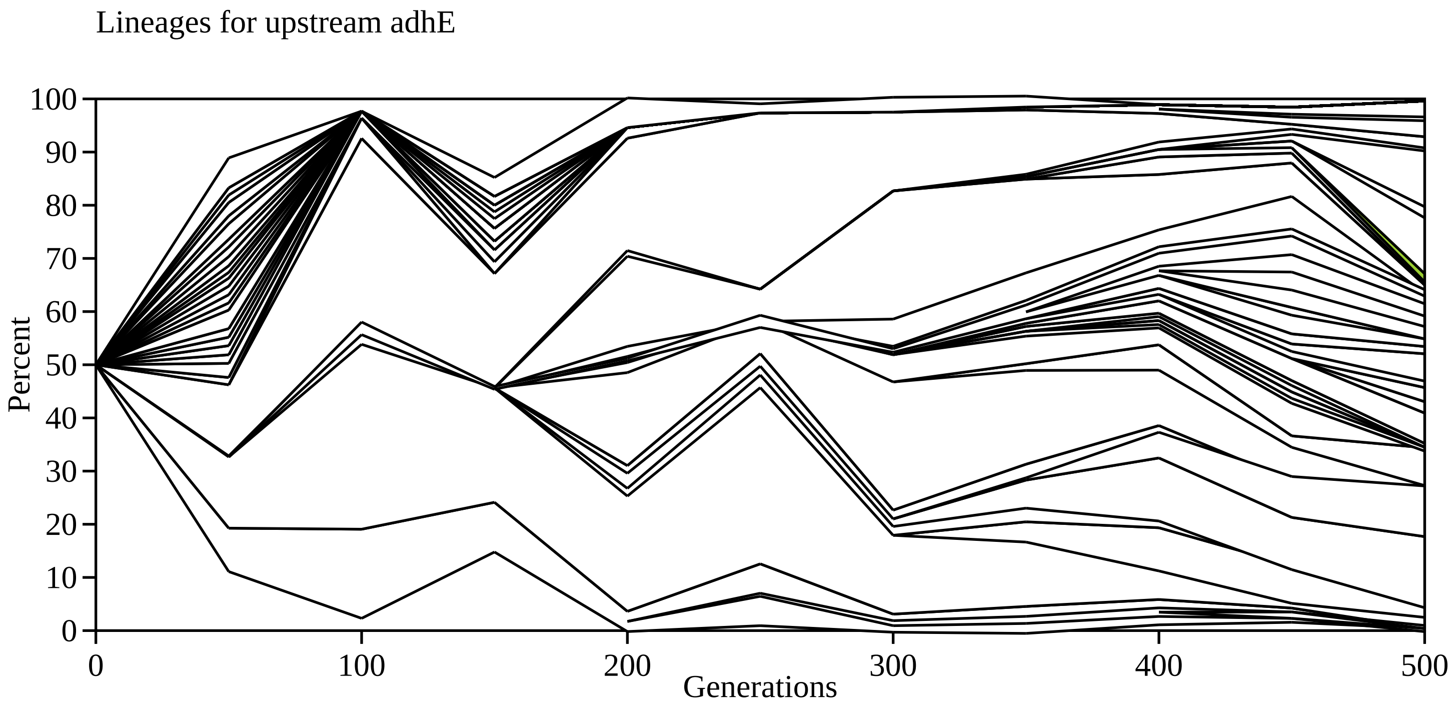

Lineages for upstream adhE
Percent
0
10
20
30
40
50
60
70
80
90
100
0
100
200
300
400
500
Generations

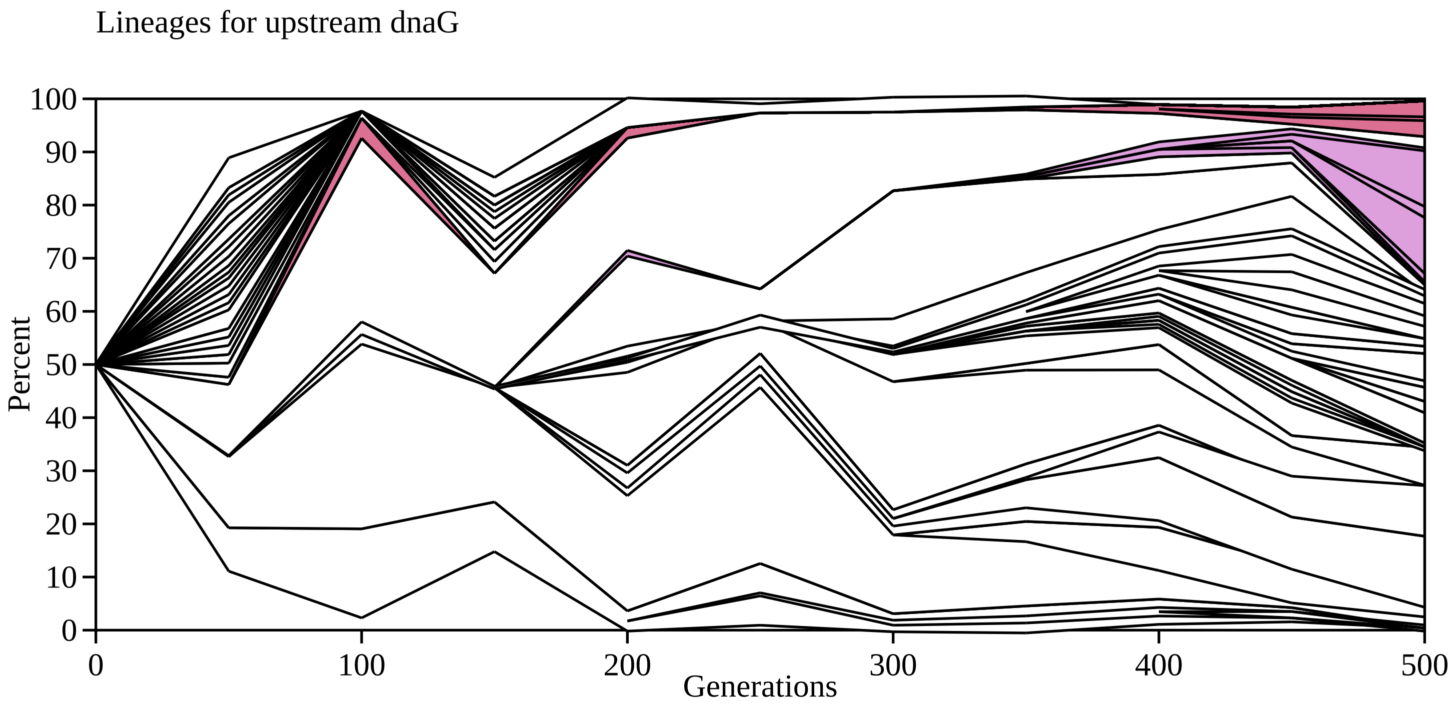

Lineages for upstream dnaG
Percent
100
90
80
70
60
50
40
30
20
10
0
0
100
200
300
400
500
Generations

Lineages for upstream mglB

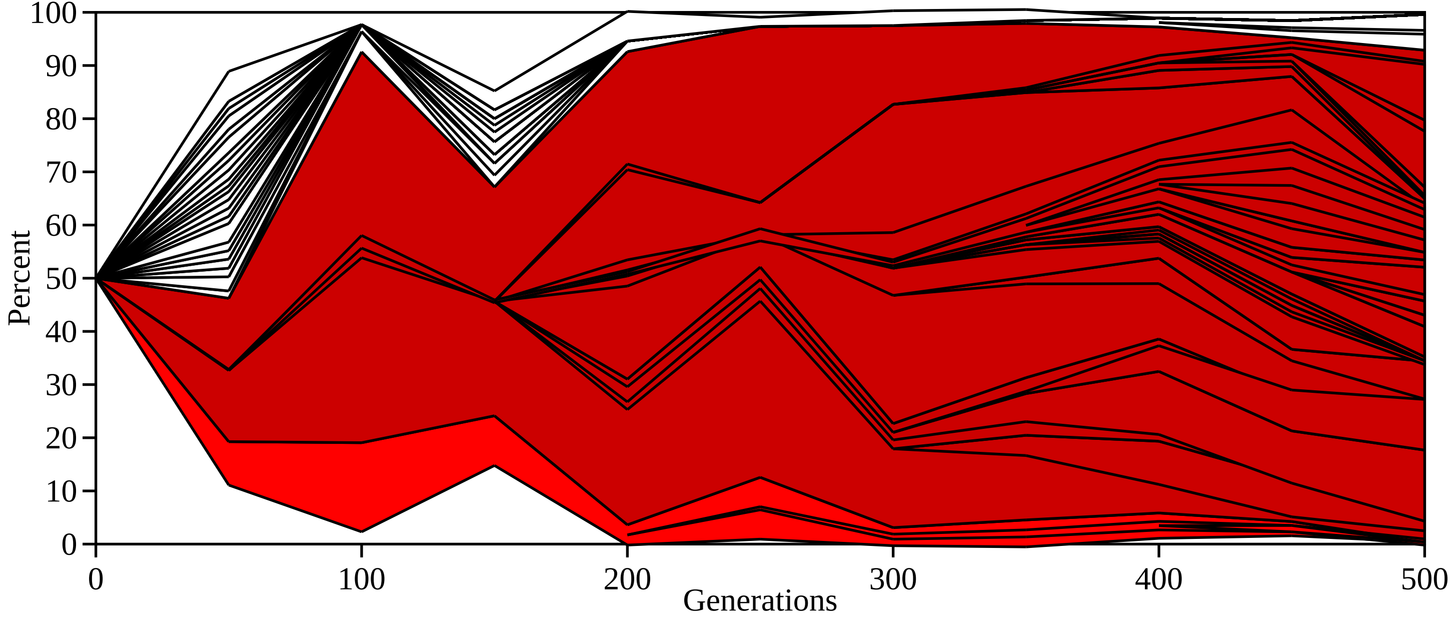

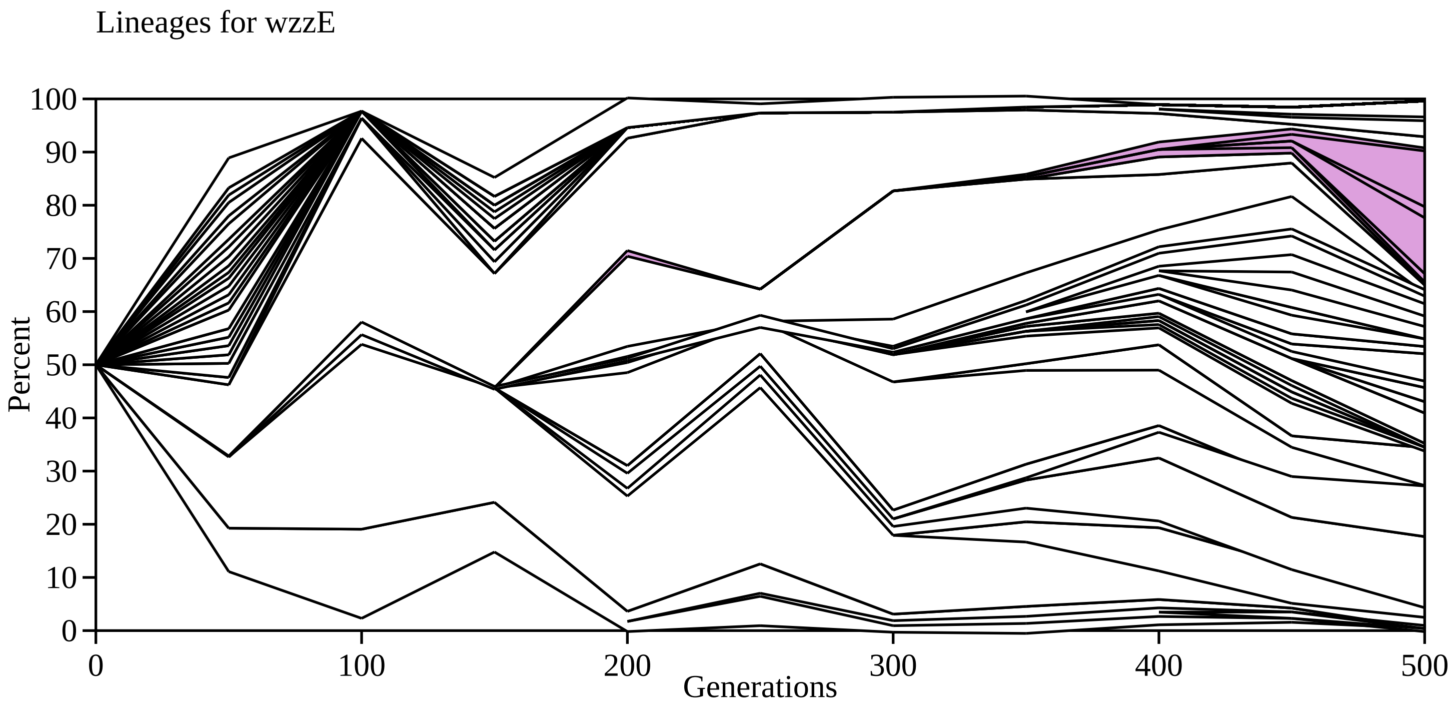

Lineages for wzzE
Percent
0
10
20
30
40
50
60
70
80
90
100
0
100
200
300
400
500
Generations

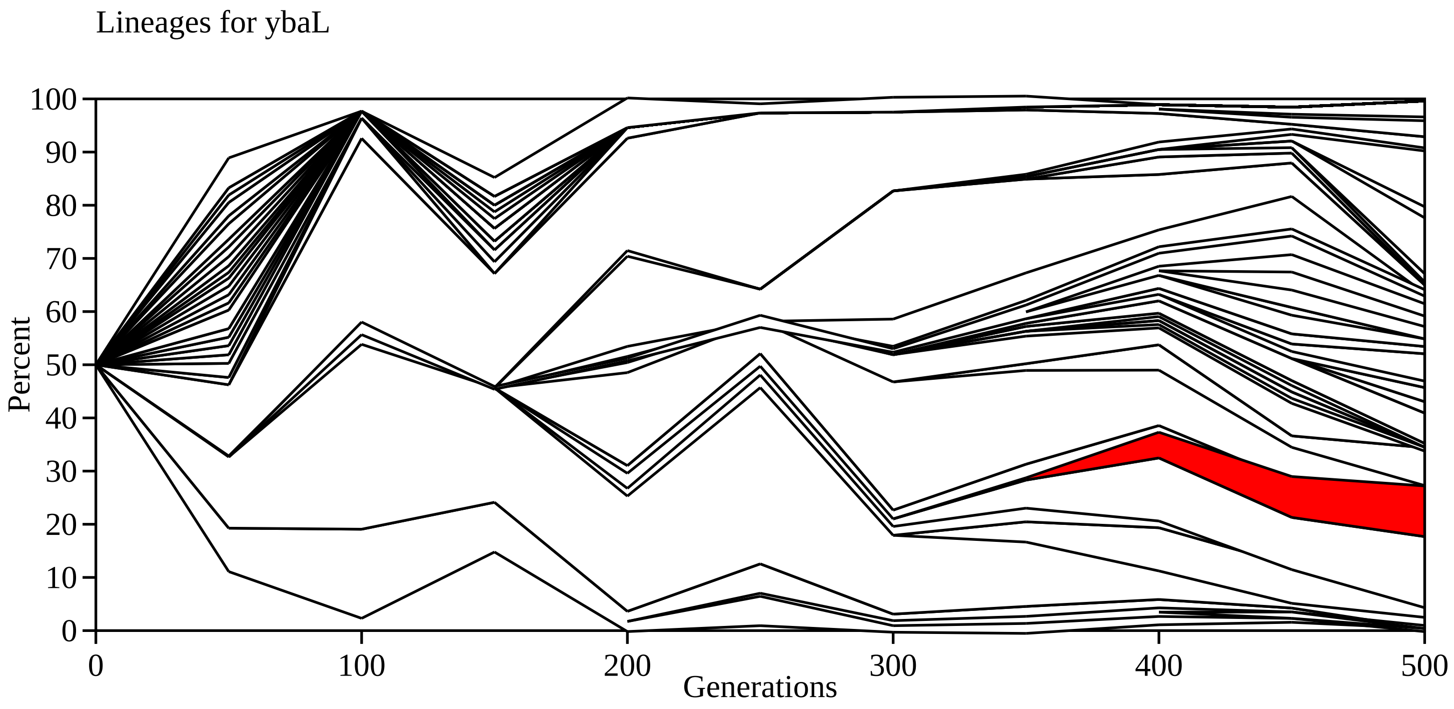

Lineages for ybaL
Percent
Generations
0
10
20
30
40
50
60
70
80
90
100
0
100
200
300
400
500

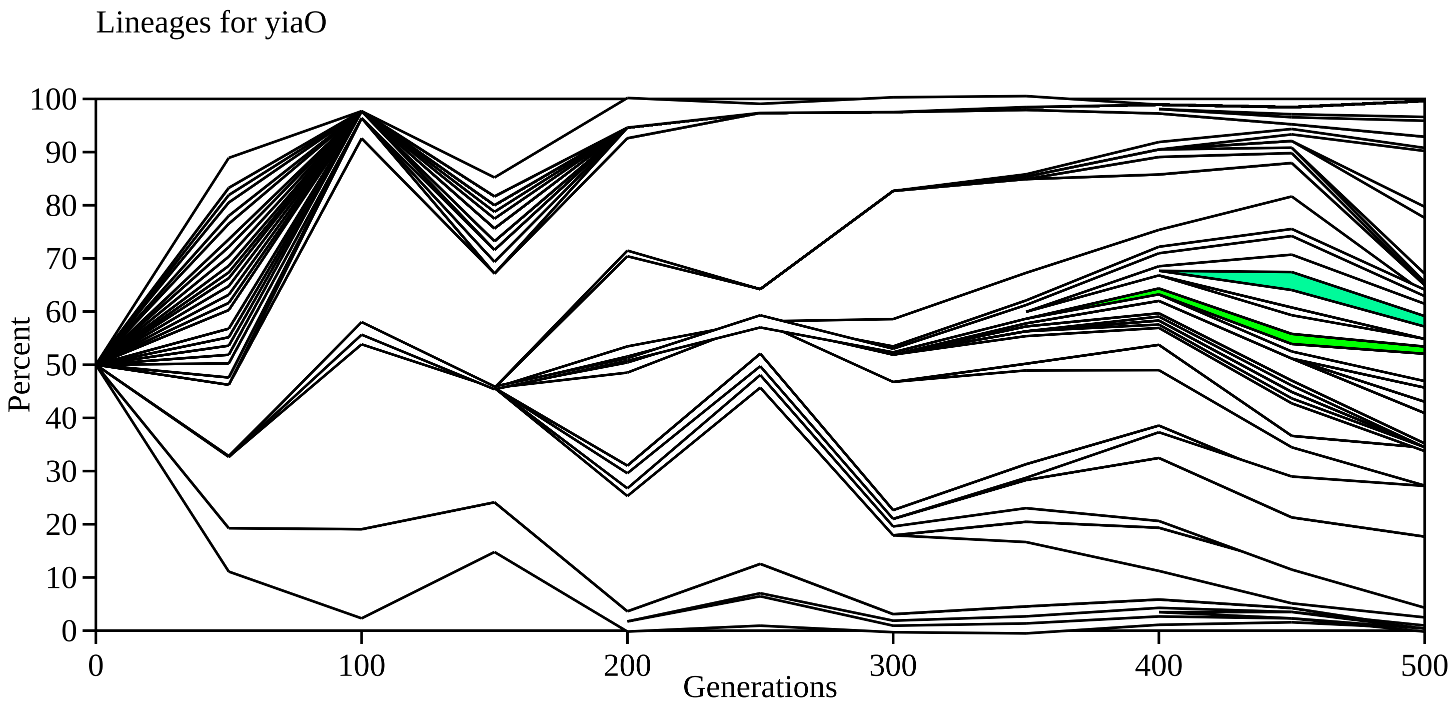

Lineages for yiaO
Percent
Generations
0
10
20
30
40
50
60
70
80
90
100
0
100
200
300
400
500

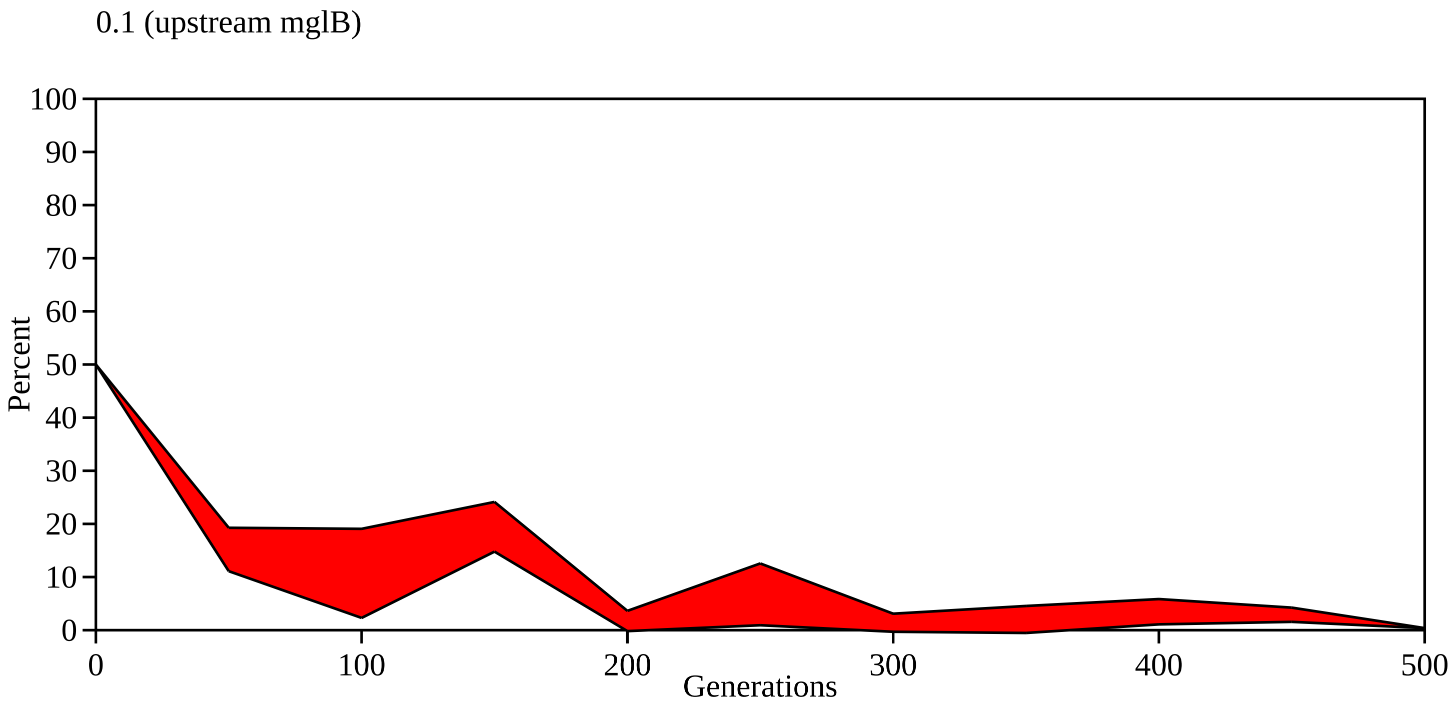

0.1 (upstream mglB)
100
90
80
70
60
50
40
30
20
10
0
Percent
0
100
200
300
400
500
Generations

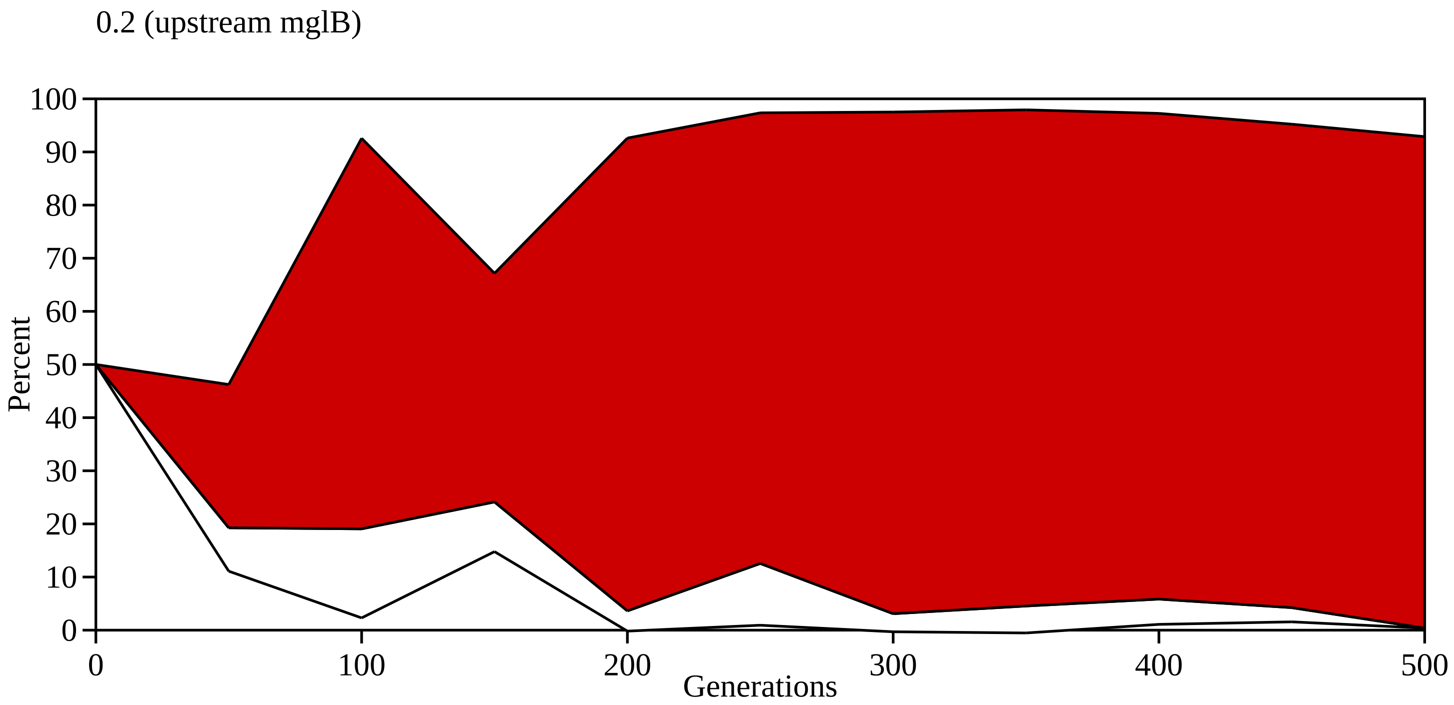
0.2 (upstream mglB)
Percent
100
90
80
70
60
50
40
30
20
10
0
0
100
200
300
400
500
Generations

0.3 (fimH, fimH, upstream dnaG)

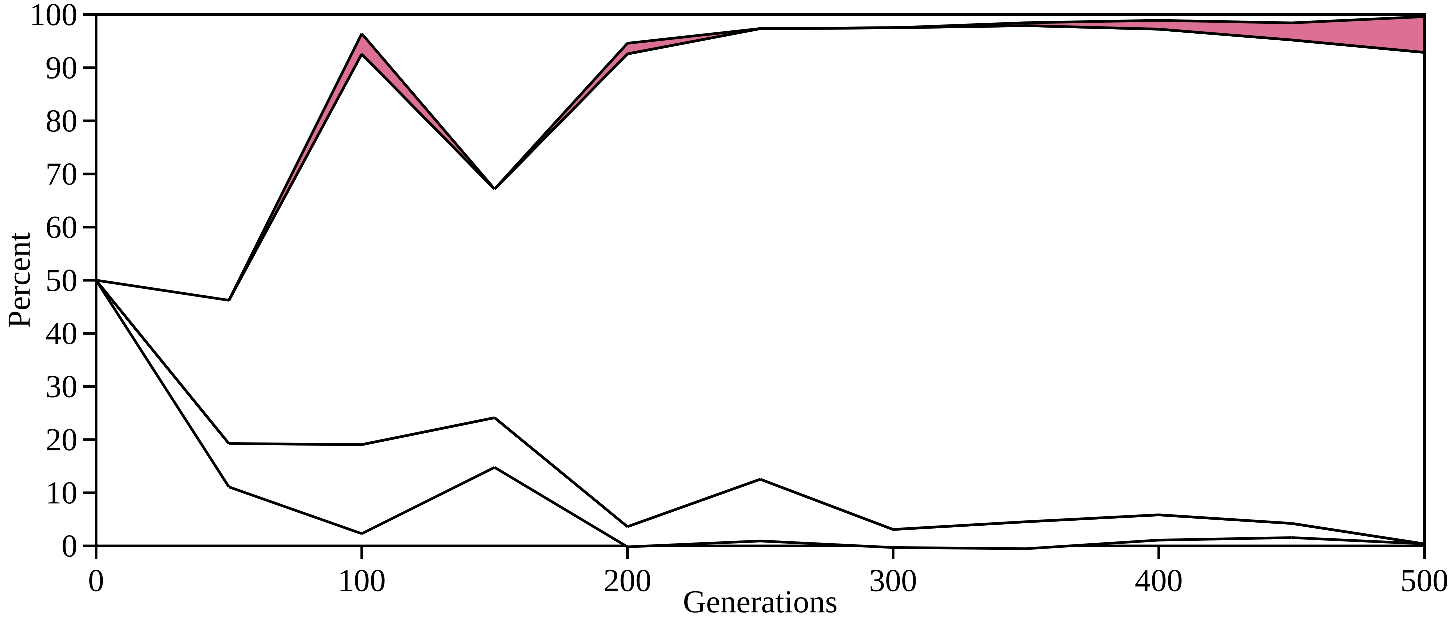

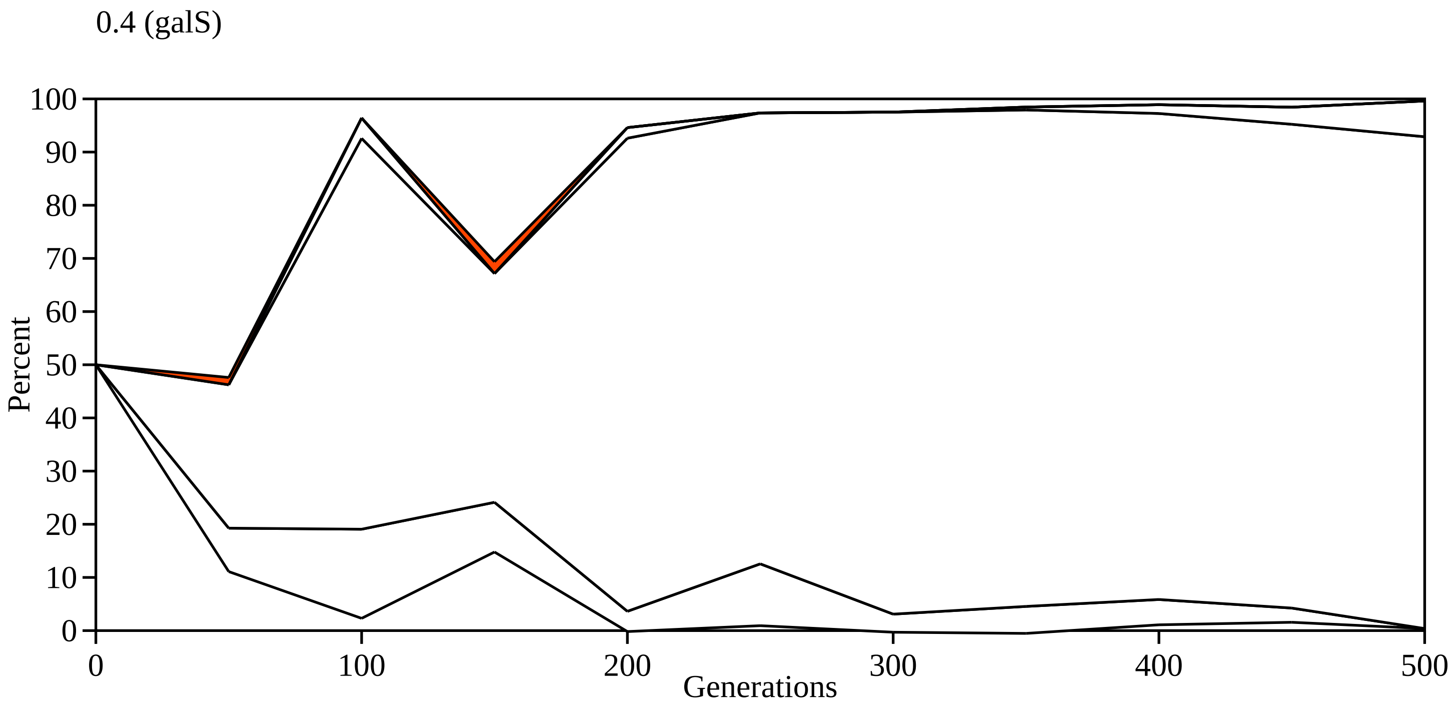

0.4 (galS)
Percent
100
90
80
70
60
50
40
30
20
10
0
0
100
200
300
400
500
Generations

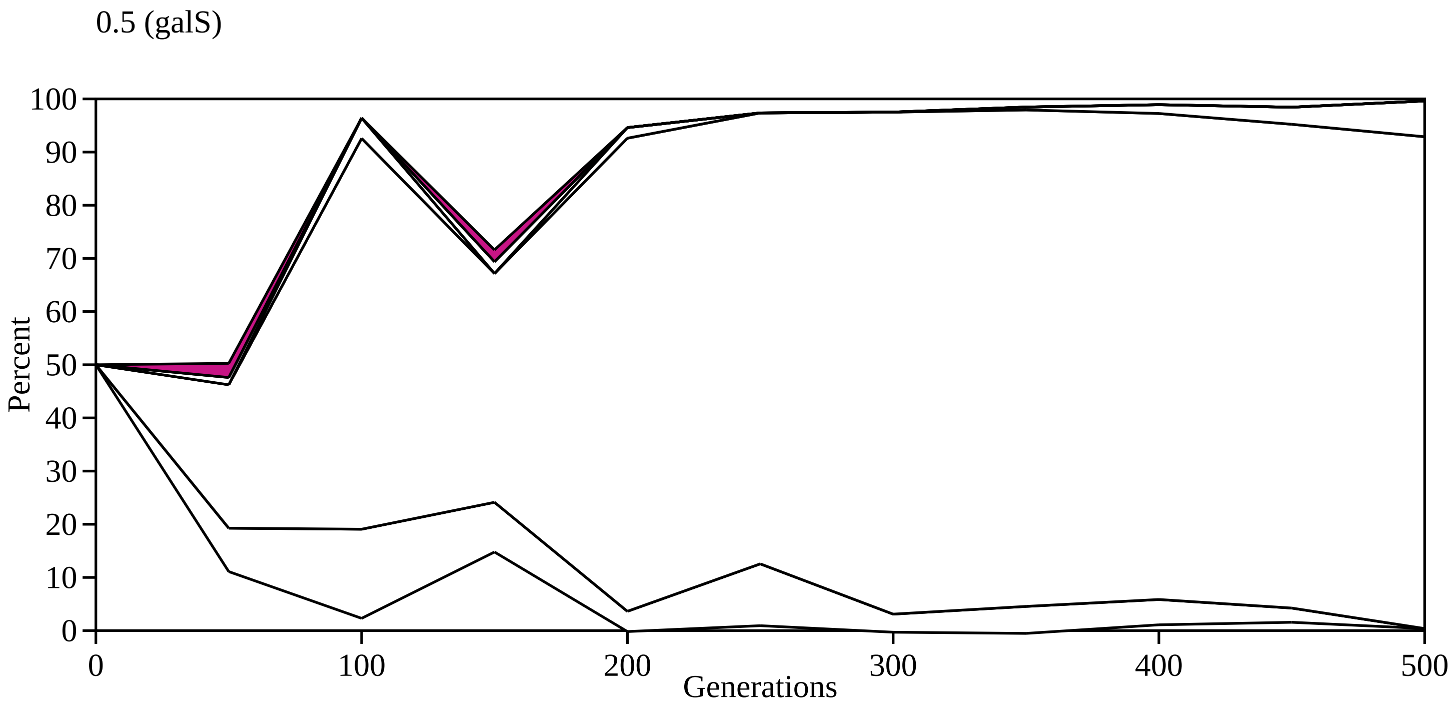
0.5 (galS)
Percent
Generations
100
90
80
70
60
50
40
30
20
10
0
0
100
200
300
400
500

0.6 (galS)

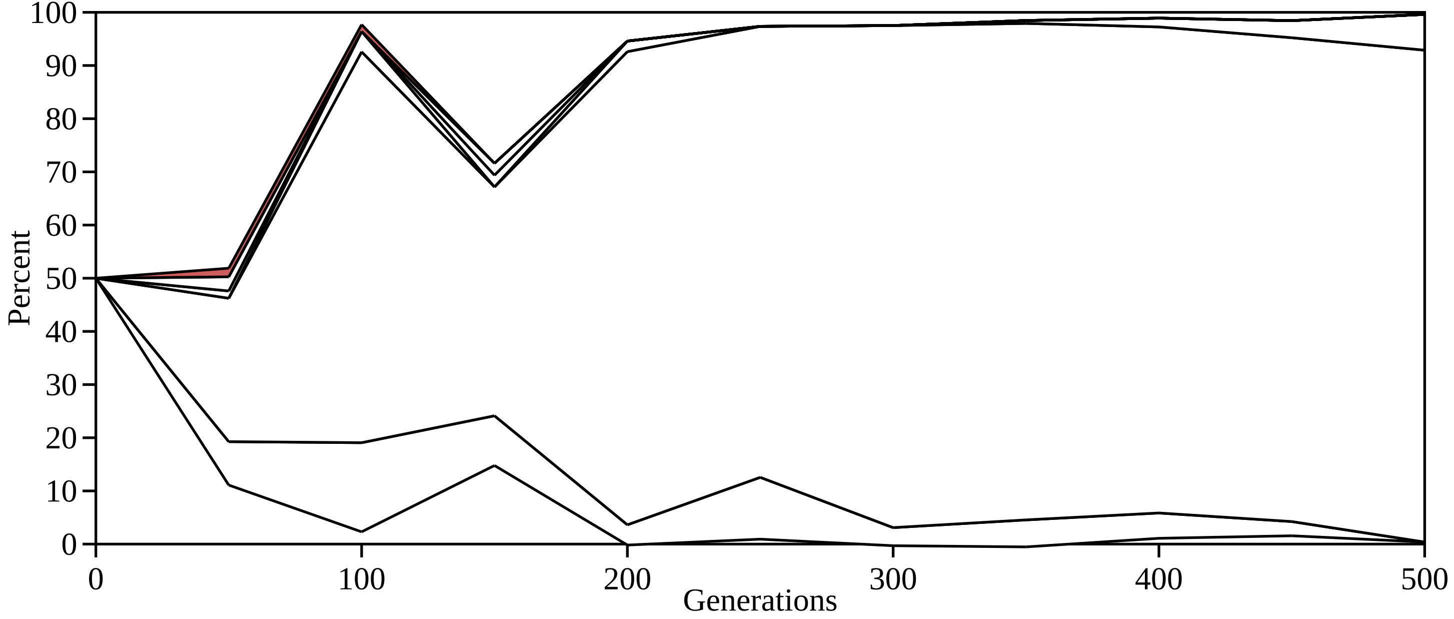

100
90
80
70
60
50
40
30
20
10
0
Percent
0
100
200
300
400
500
Generations

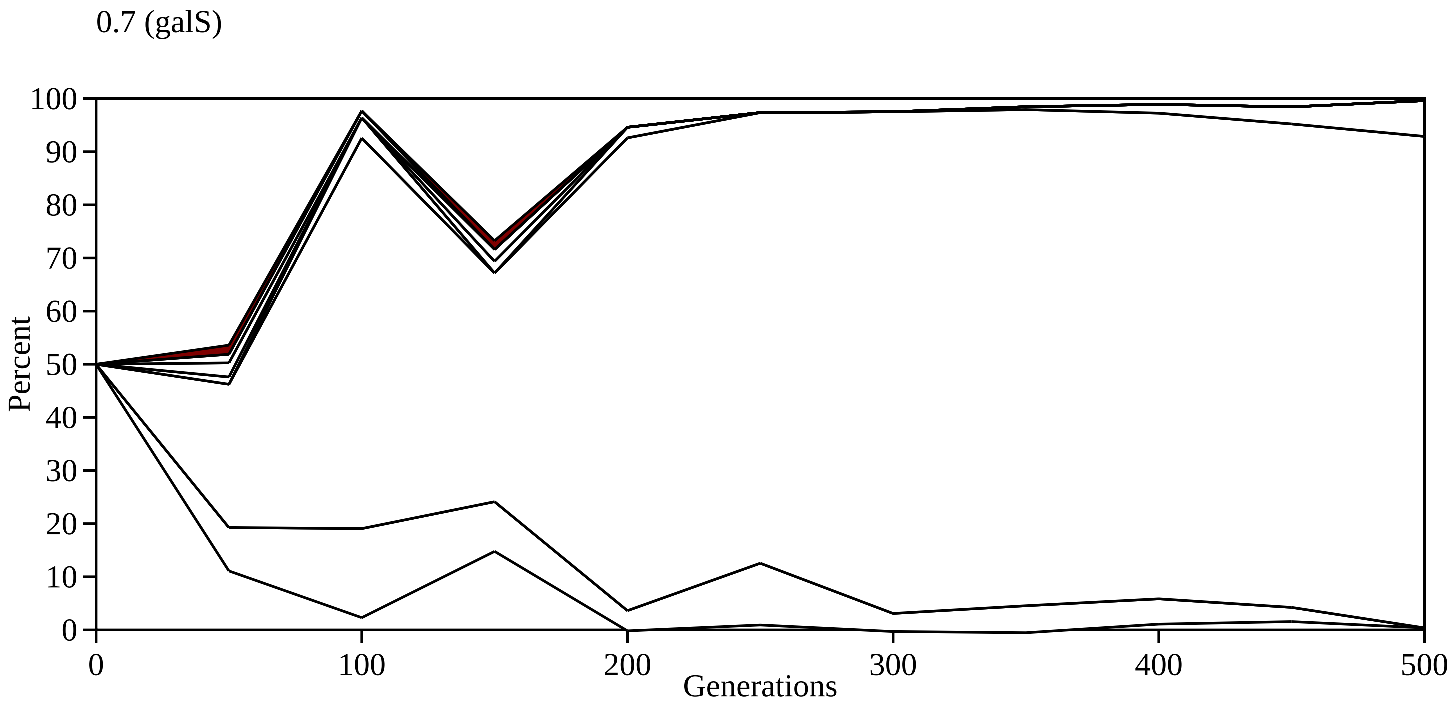
0.7 (galS)
Percent
100
90
80
70
60
50
40
30
20
10
0
0
100
200
300
400
500
Generations

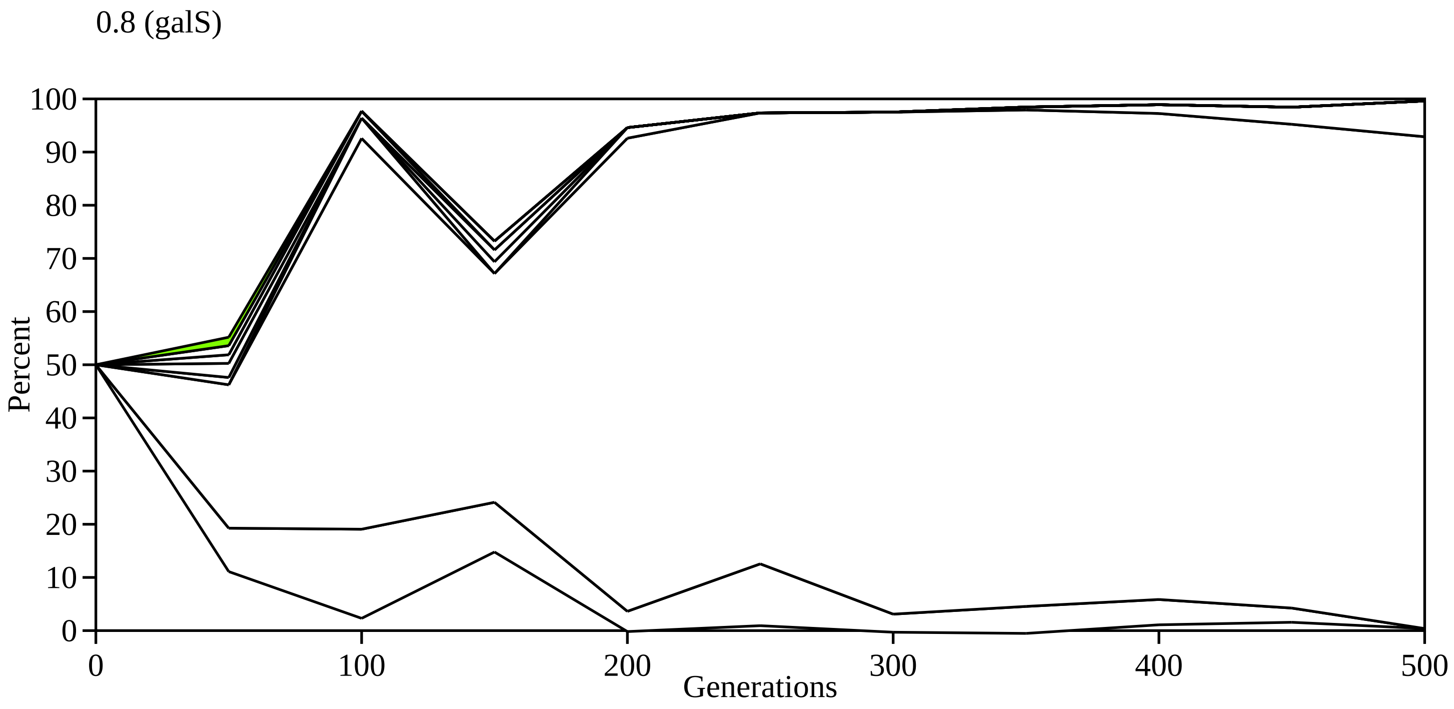
0.8 (galS)
100
90
80
70
60
50
40
30
20
10
0
Percent
0
100
200
300
400
500
Generations

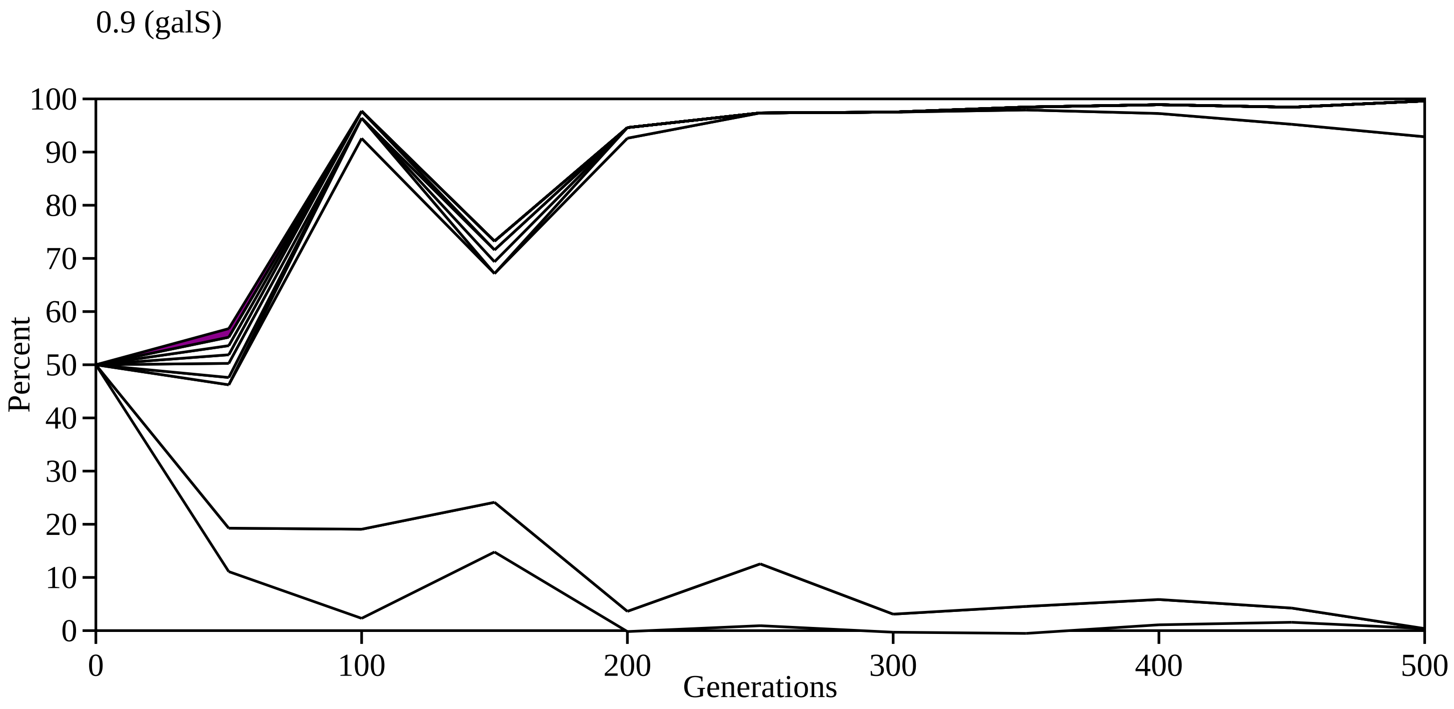

0.9 (galS)
100
90
80
70
60
50
40
30
20
10
0
Percent
0
100
200
300
400
500
Generations

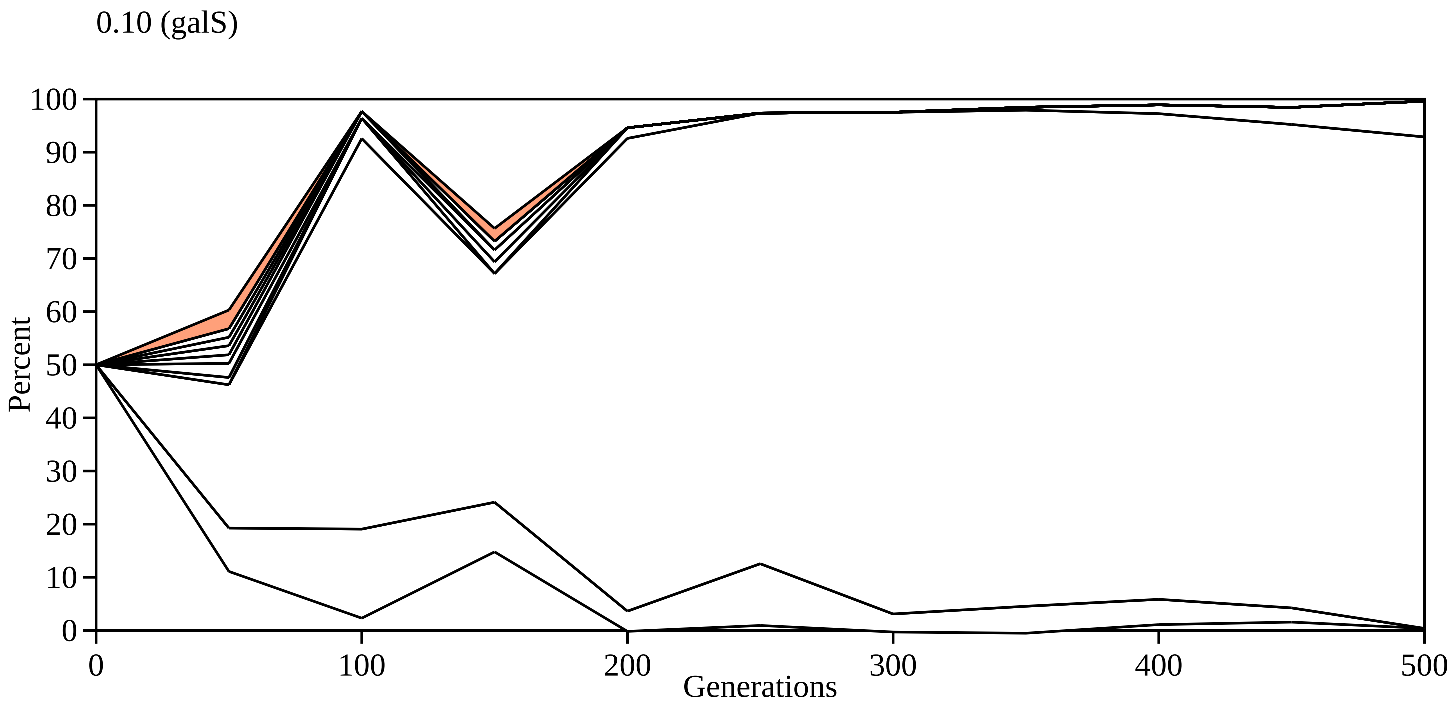
0.10 (galS)
100
90
80
70
60
50
40
30
20
10
0
Percent
0
100
200
300
400
500
Generations

0.11 (galS)

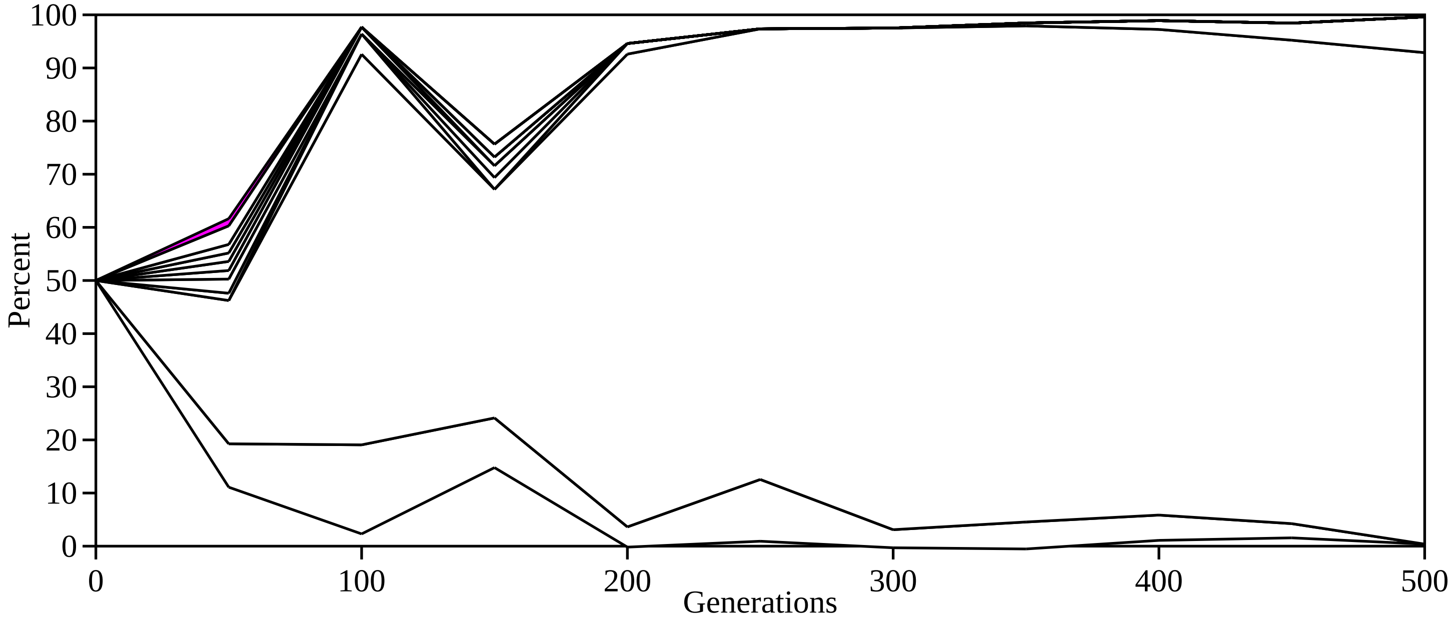
100
90
80
70
60
50
40
30
20
10
0
Percent
0
100
200
300
400
500
Generations

0.12 (galS)

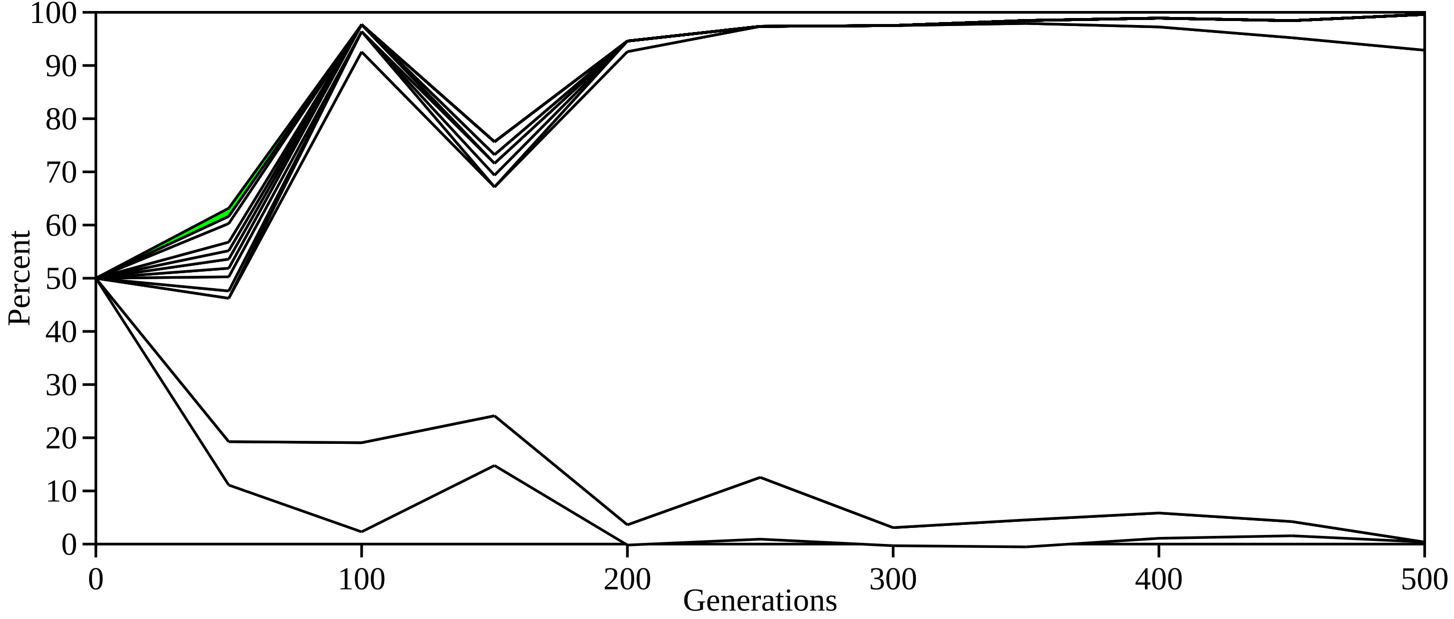

100
90
80
70
60
50
40
30
20
10
0
Percent
0
100
200
300
400
500
Generations

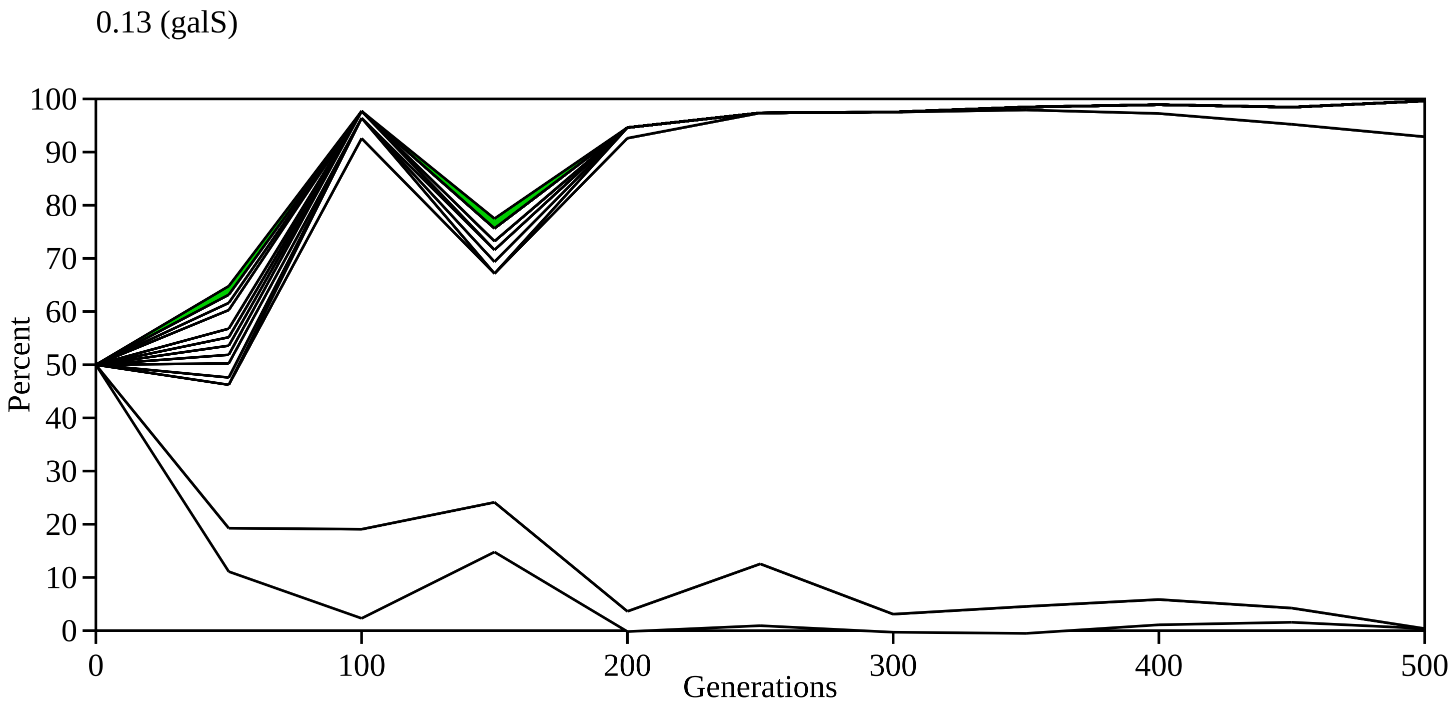
0.13 (galS)
Percent
100
90
80
70
60
50
40
30
20
10
0
0
100
200
300
400
500
Generations

0.14 (galS)

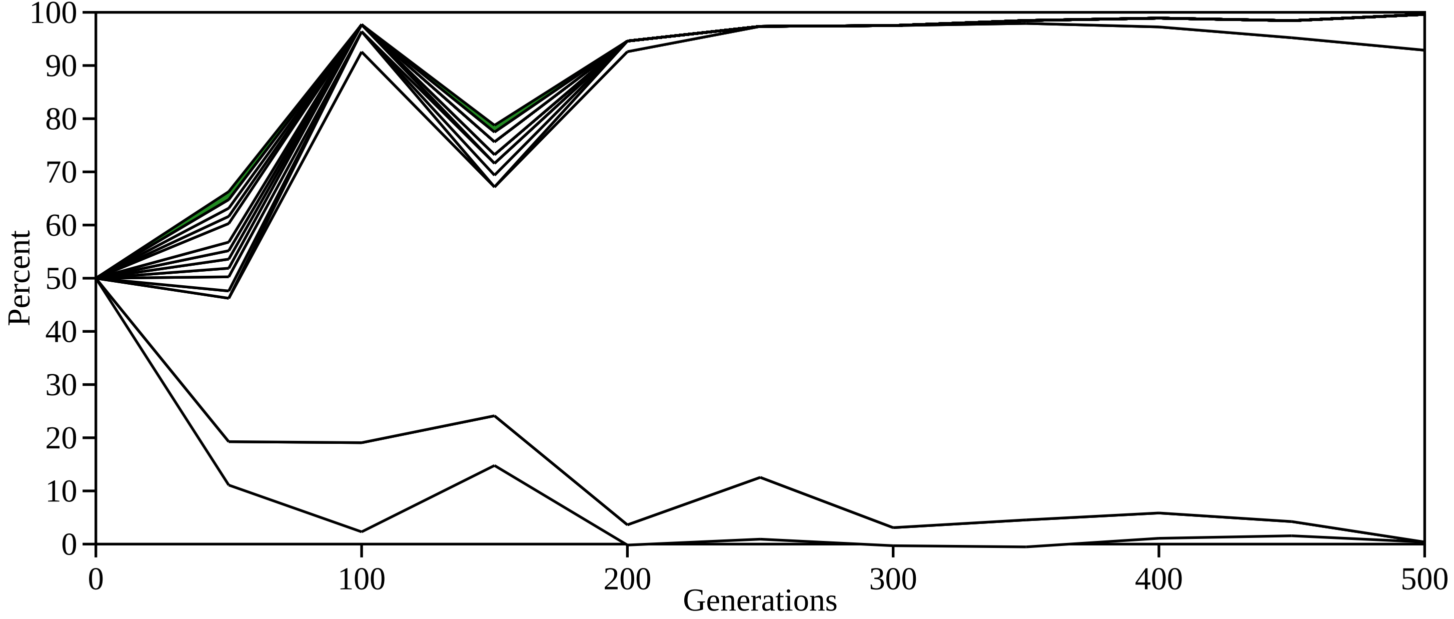

100
90
80
70
60
50
40
30
20
10
0
Percent
0
100
200
300
400
500
Generations

0.15 (galS)

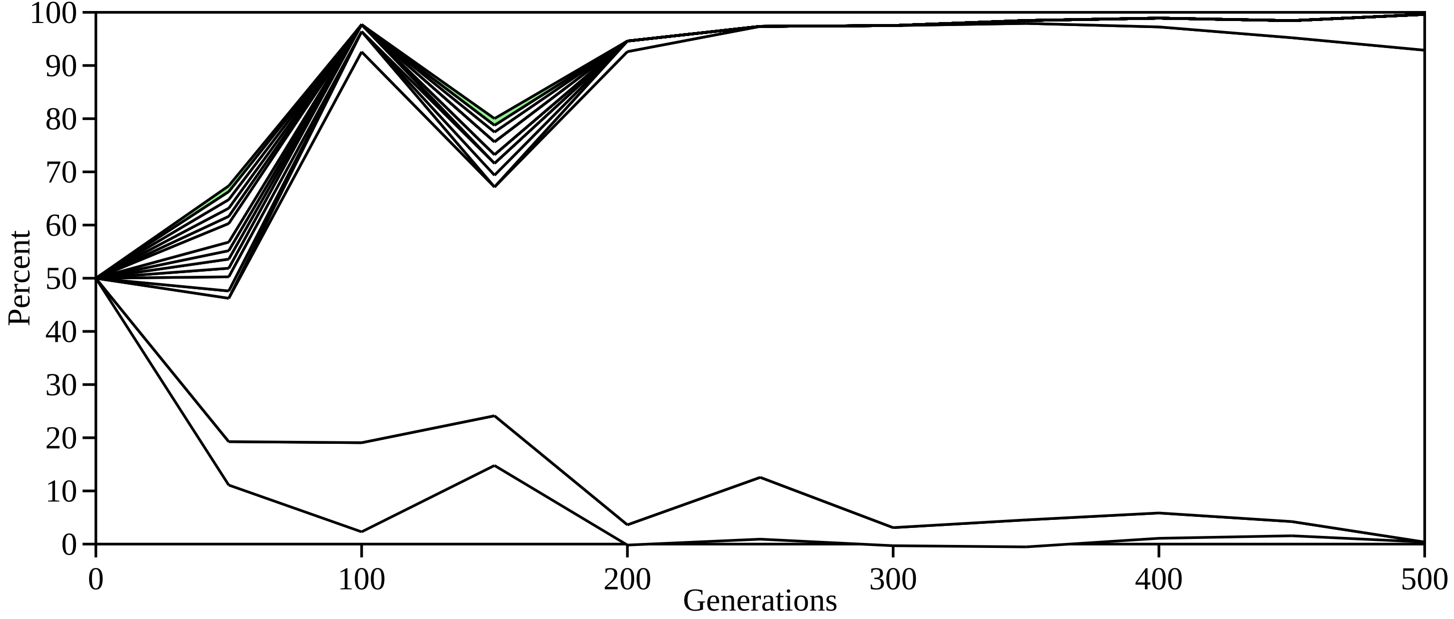

Percent
Generations
0
10
20
30
40
50
60
70
80
90
100
0
100
200
300
400
500

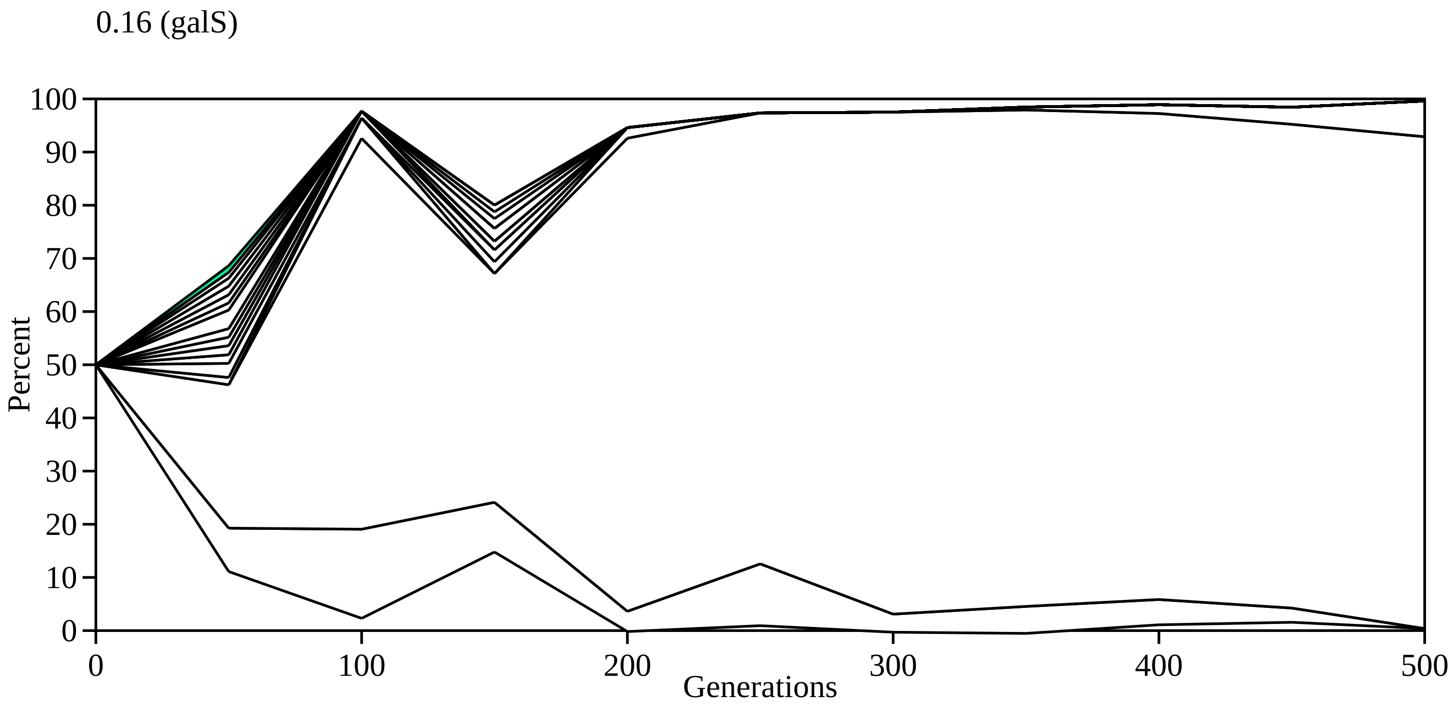
0.16 (galS)
Percent
100
90
80
70
60
50
40
30
20
10
0
0
100
200
300
400
500
Generations

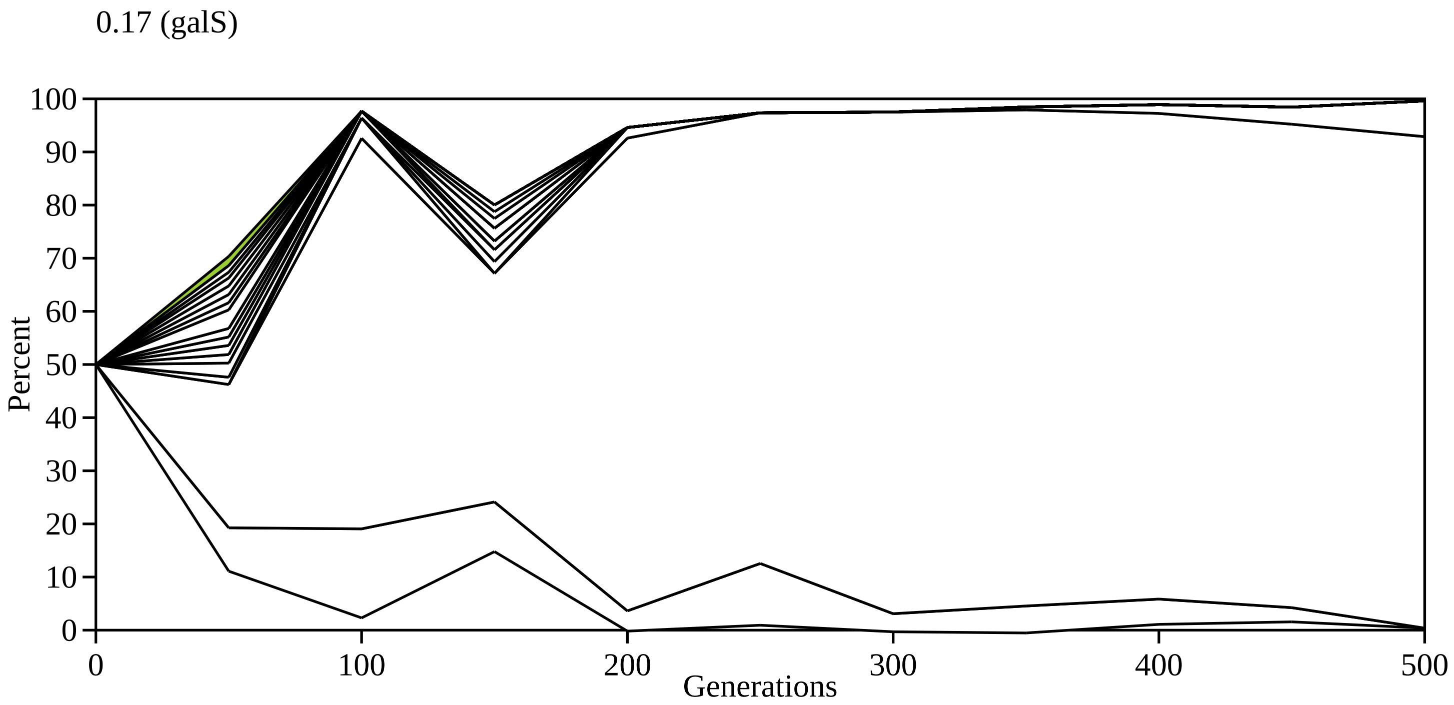

0.17 (galS)
Percent
Generations
0
10
20
30
40
50
60
70
80
90
100
0
100
200
300
400
500

0.18 (galS)

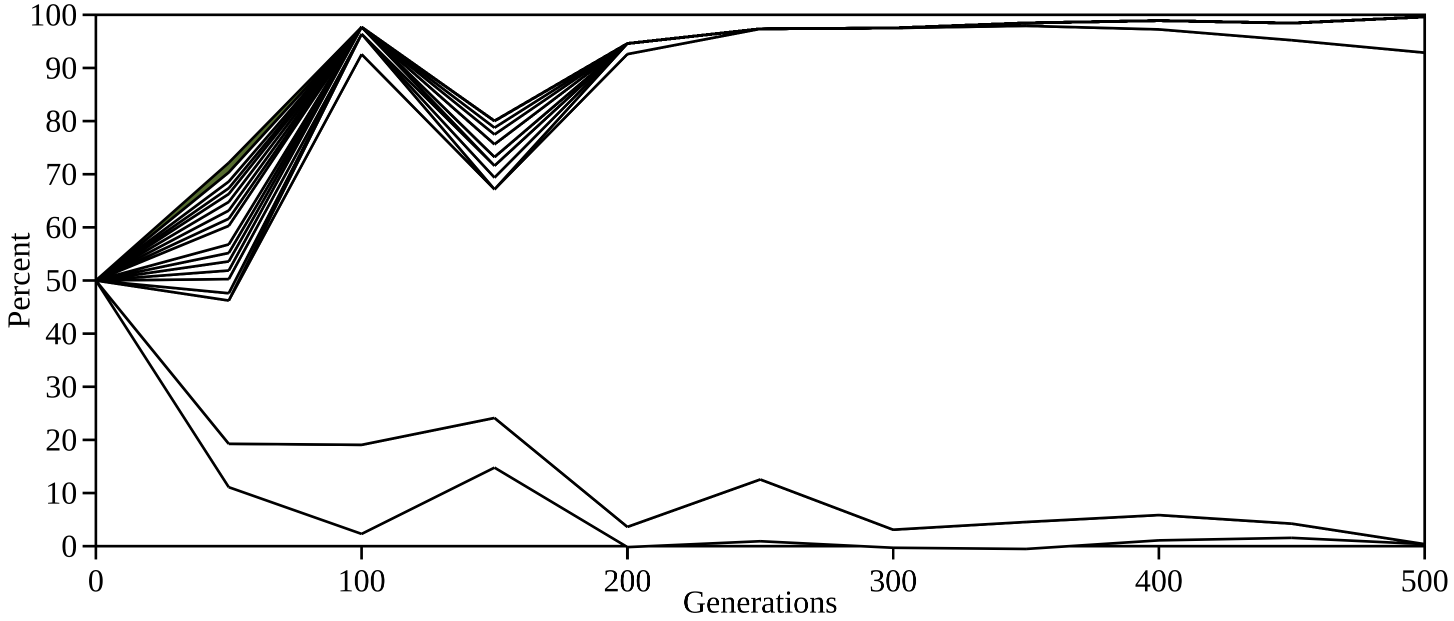
100
90
80
70
60
50
40
30
20
10
0
Percent
0
100
200
300
400
500
Generations

0.19 (galS)

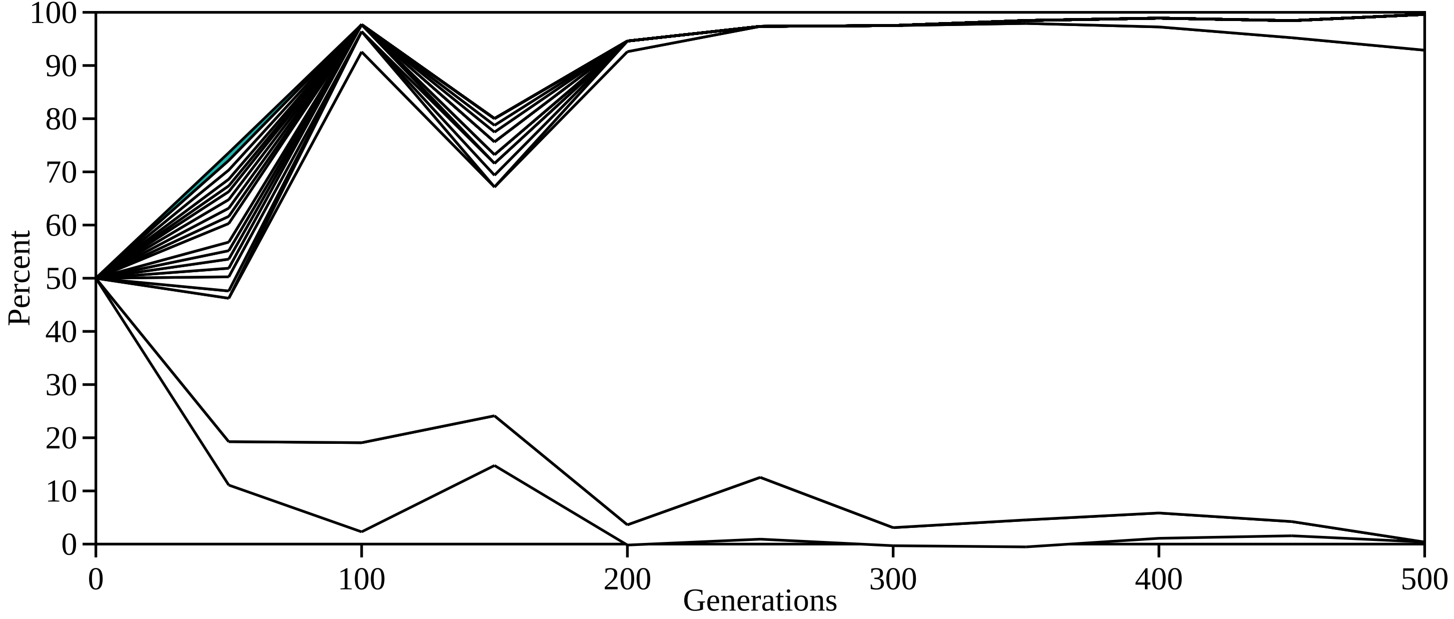

100
90
80
70
60
50
40
30
20
10
0
Percent
0
100
200
300
400
500
Generations

0.20 (galS)

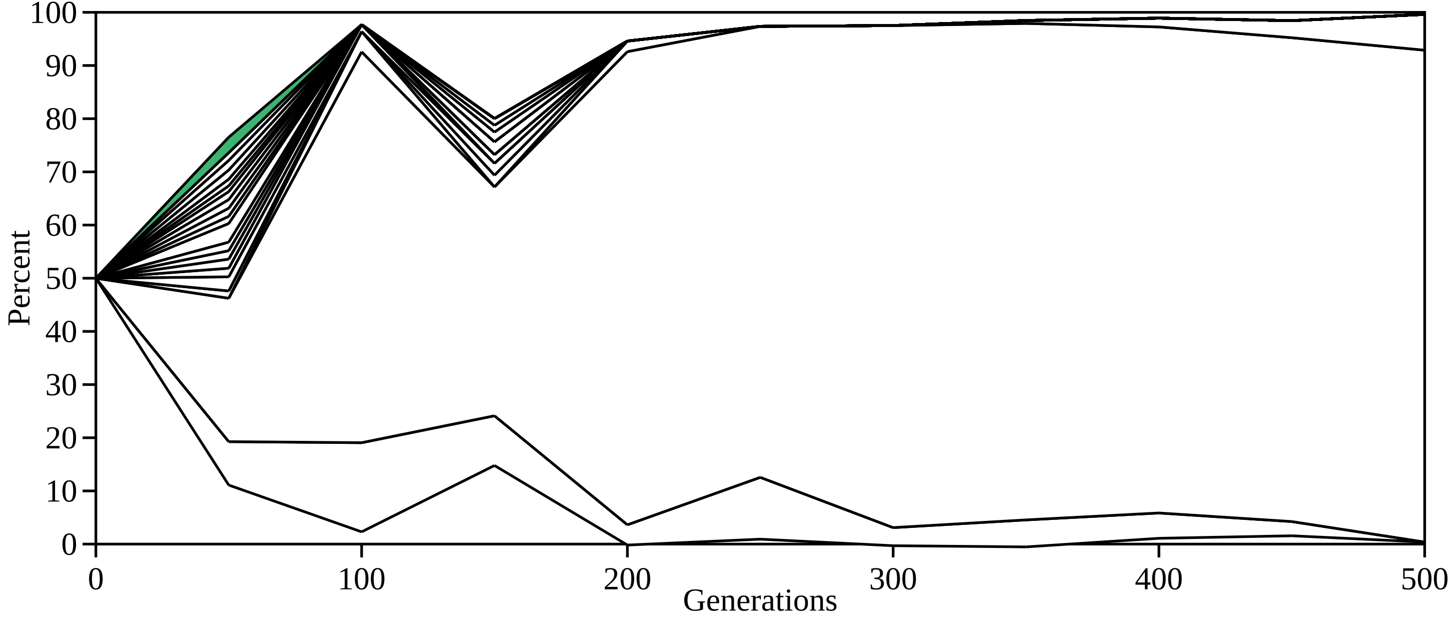

100
90
80
70
60
50
40
30
20
10
0
Percent
0
100
200
300
400
500
Generations

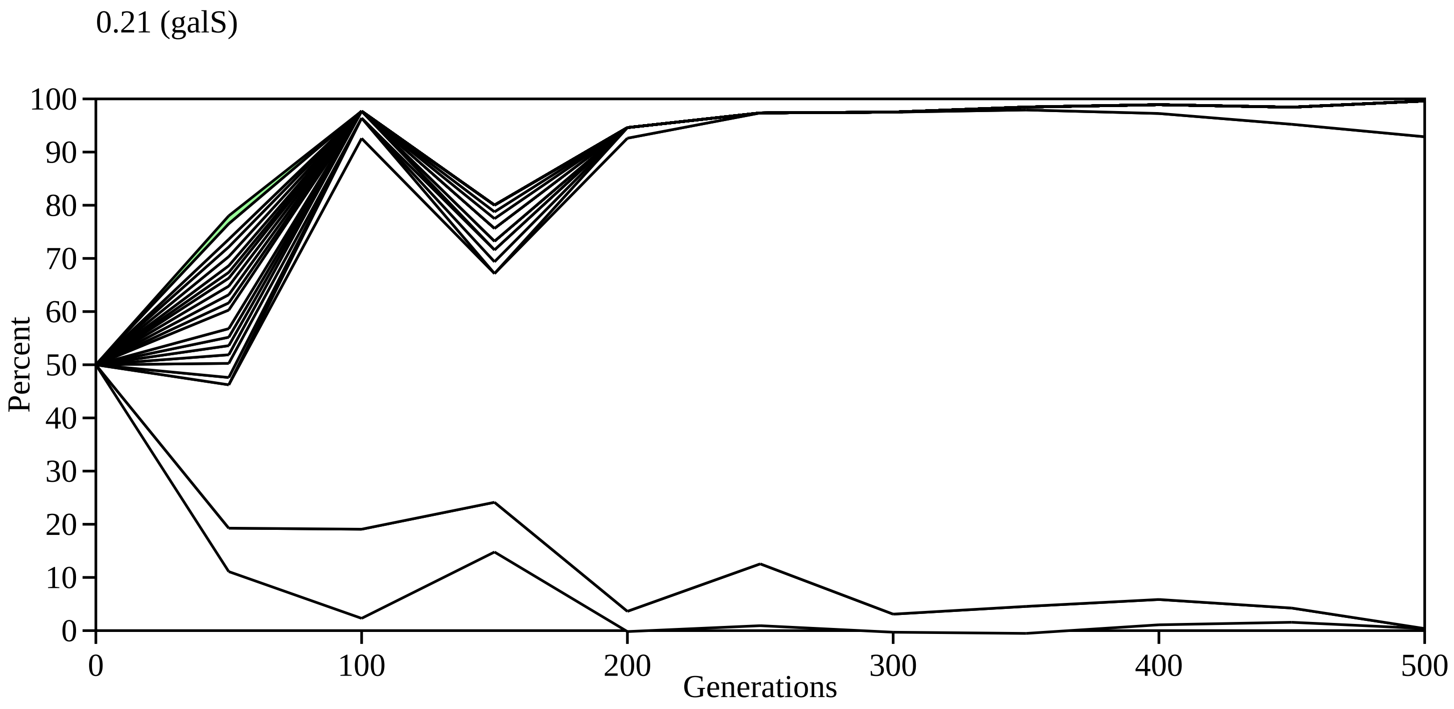

0.21 (galS)
Percent
Generations
0
10
20
30
40
50
60
70
80
90
100
0
100
200
300
400
500

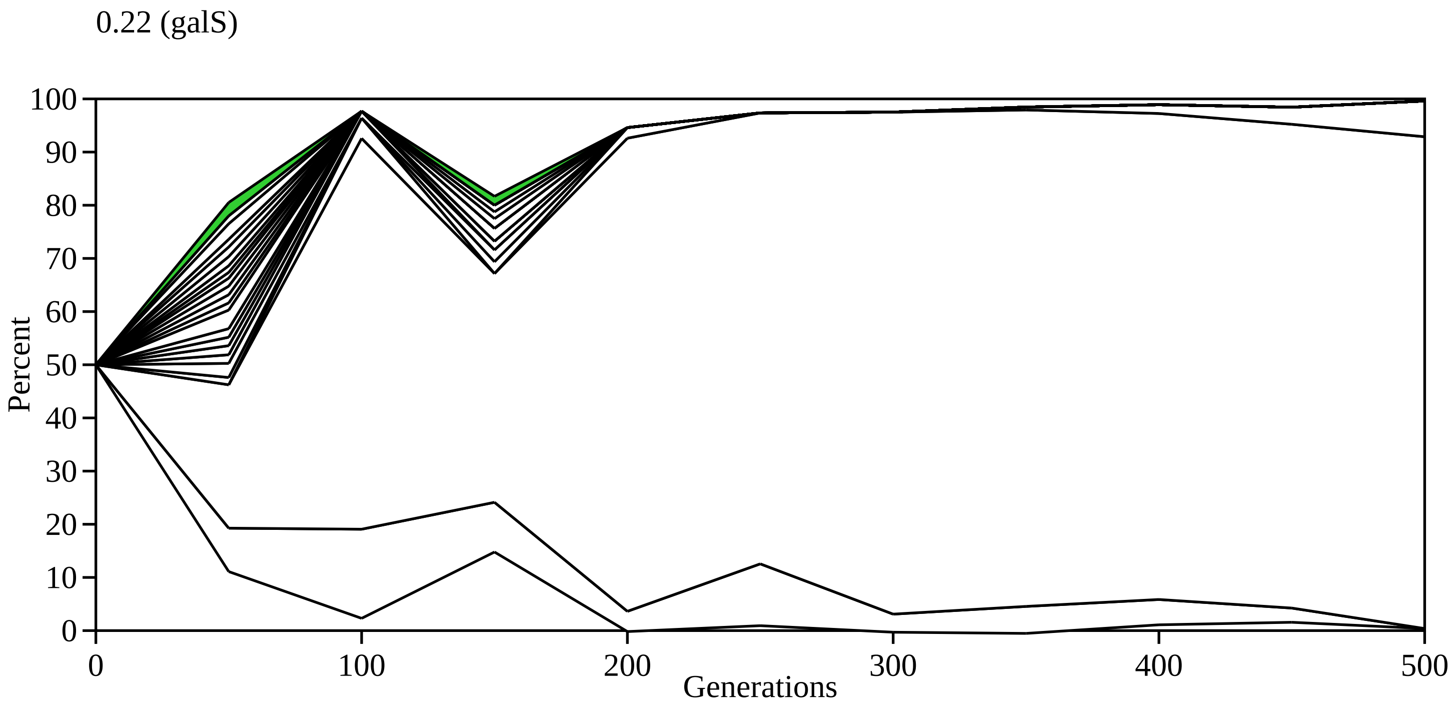
0.22 (galS)
Percent
100
90
80
70
60
50
40
30
20
10
0
0
100
200
300
400
500
Generations

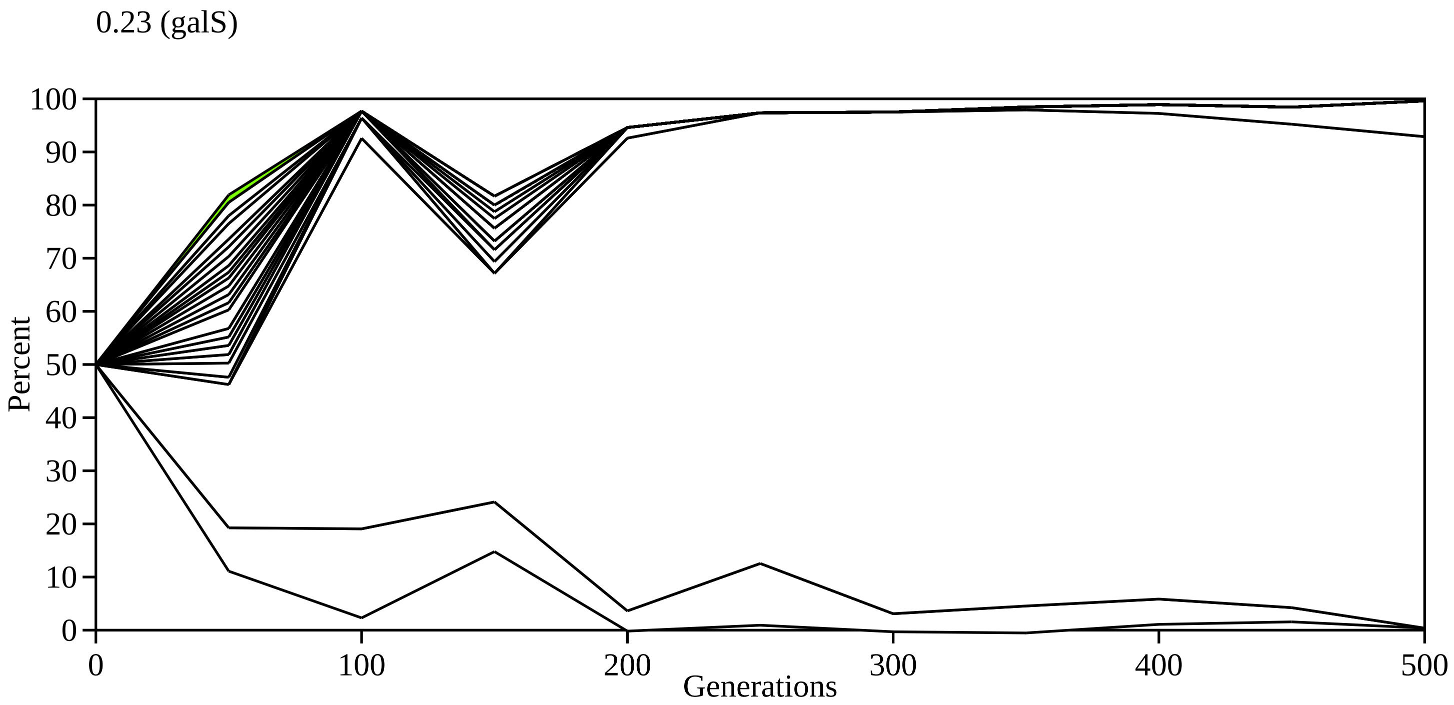
0.23 (galS)
100
90
80
70
60
50
40
30
20
10
0
Percent
0
100
200
300
400
500
Generations

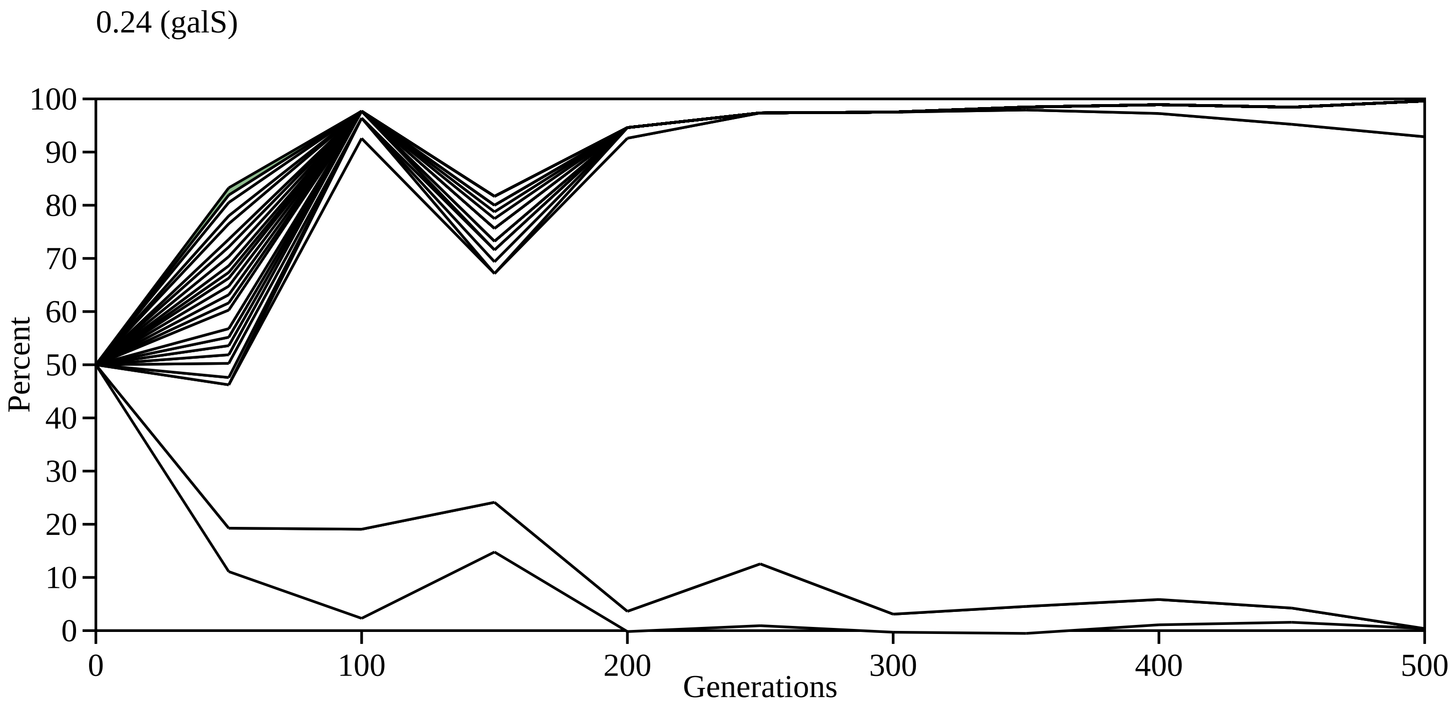
0.24 (galS)
100
90
80
70
60
50
40
30
20
10
0
Percent
0
100
200
300
400
500
Generations

0.25 (galS)

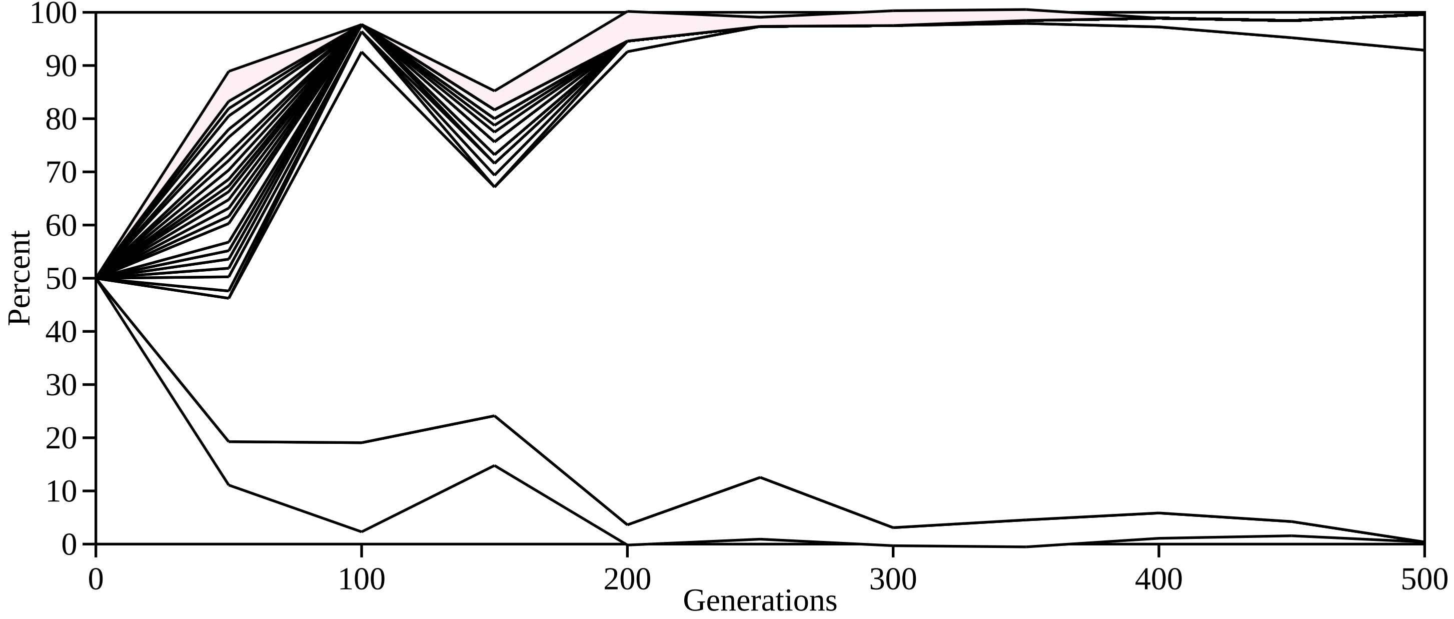

Percent
100
90
80
70
60
50
40
30
20
10
0
0
100
200
300
400
500
Generations

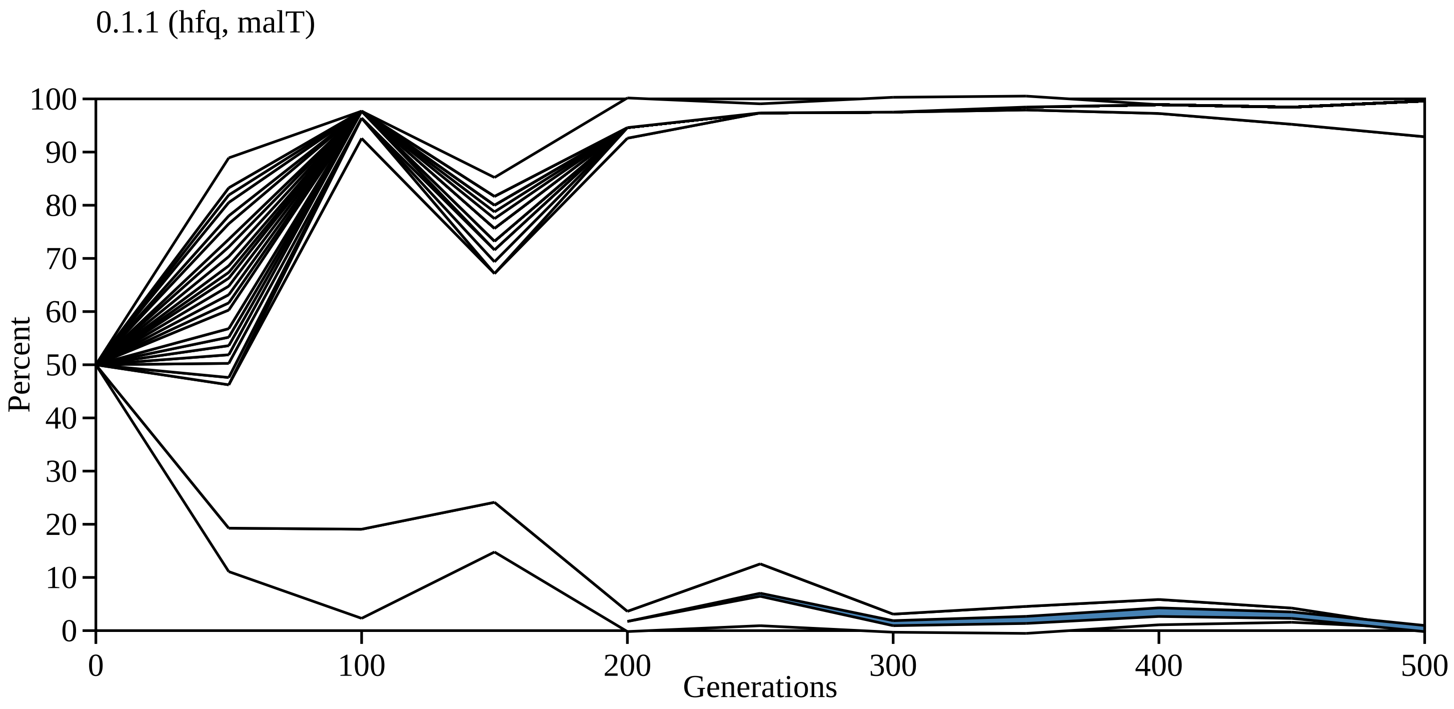

0.1.1 (hfq, malT)
Percent
Generations
0
10
20
30
40
50
60
70
80
90
100
0
100
200
300
400
500

0.1.1.1 (fimH)

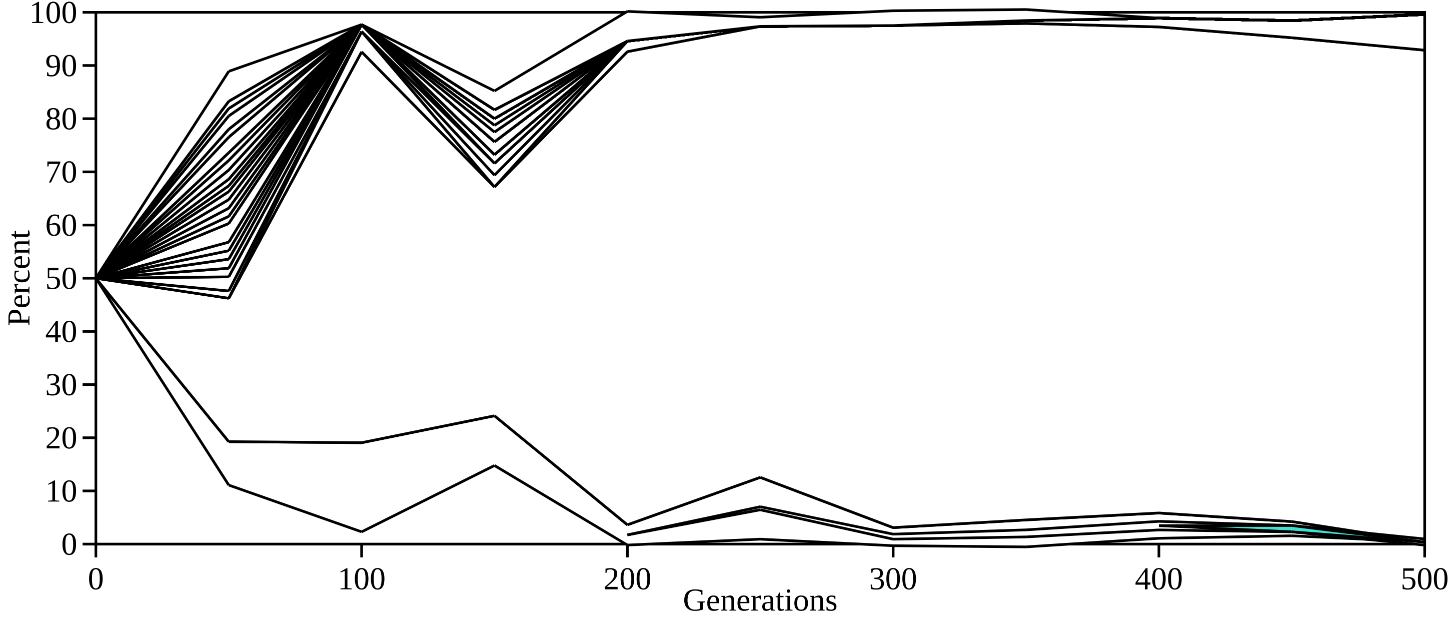

100
90
80
70
60
50
40
30
20
10
0
Percent
0
100
200
300
400
500
Generations

0.2.1 (hfq)

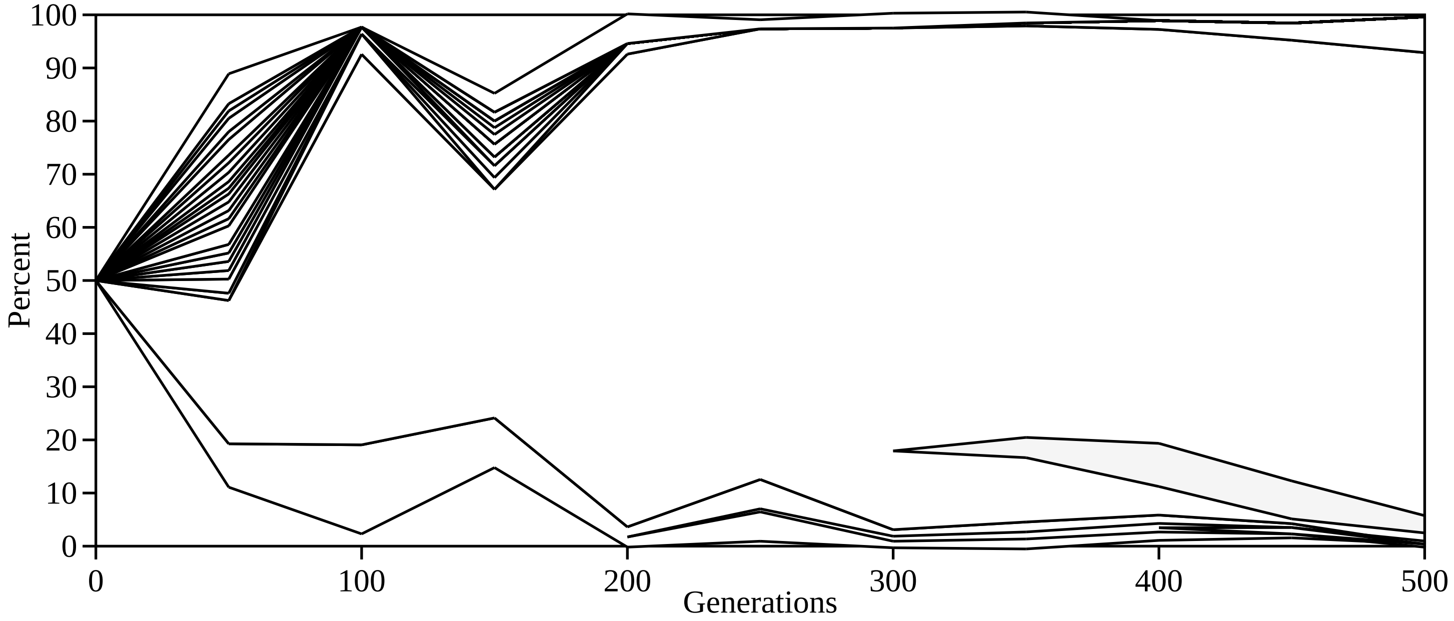

100
90
80
70
60
50
40
30
20
10
0
Percent
0
100
200
300
400
500
Generations

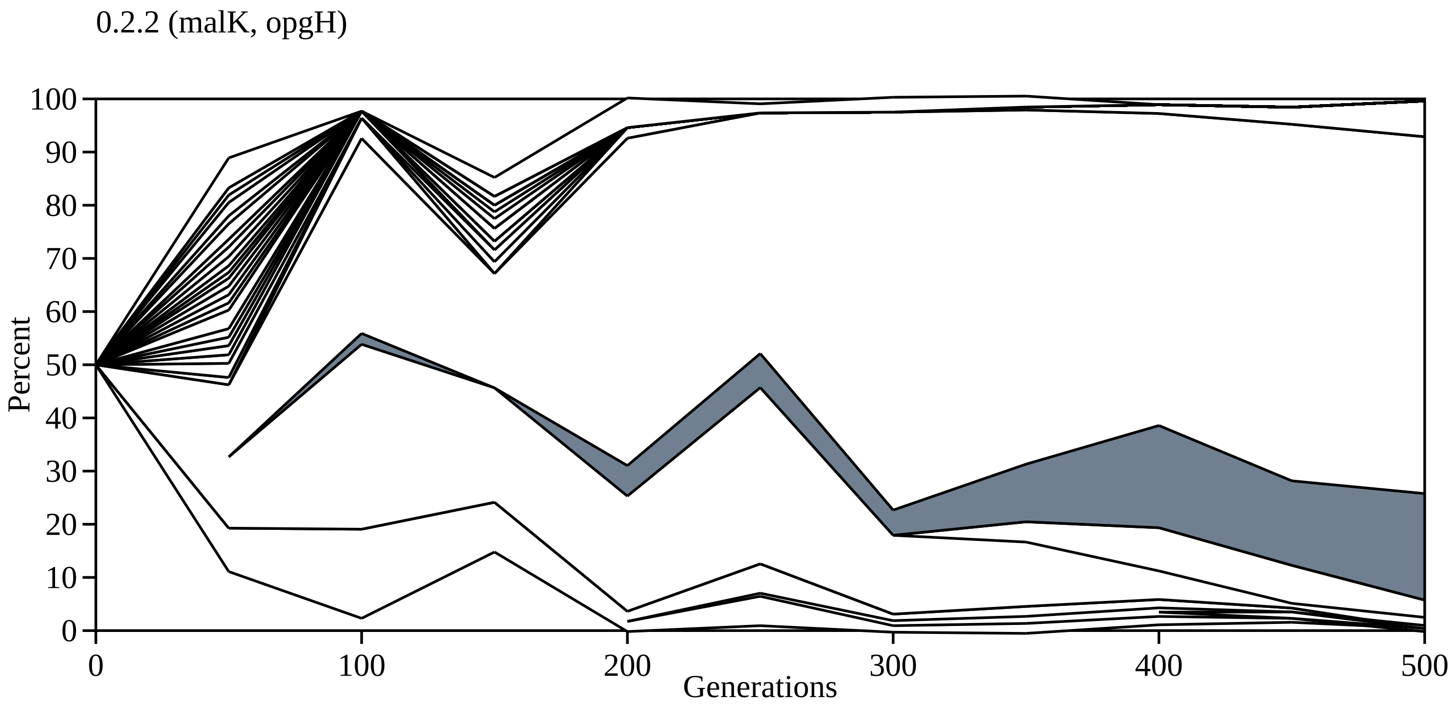
0.2.2 (malK, opgH)
Percent
Generations
0
10
20
30
40
50
60
70
80
90
100
0
100
200
300
400
500

0.2.3 (malE)

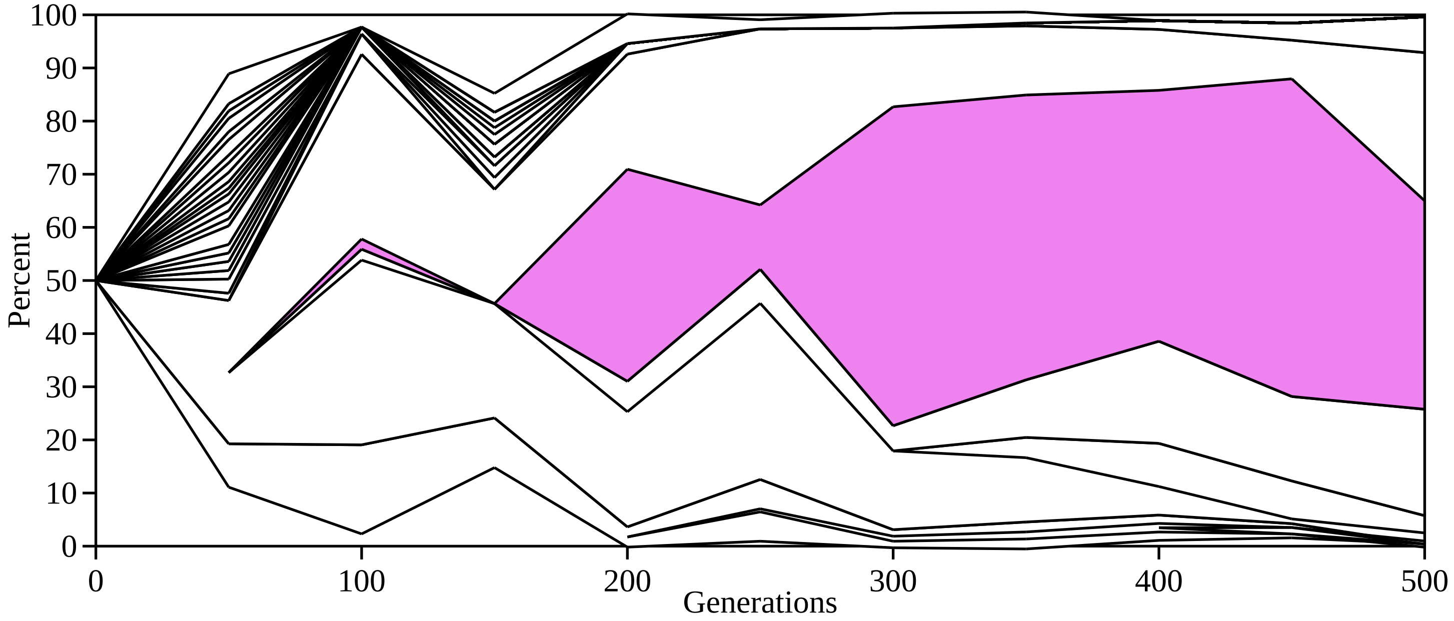
100
90
80
70
60
50
40
30
20
10
0
Percent
0
100
200
300
400
500
Generations

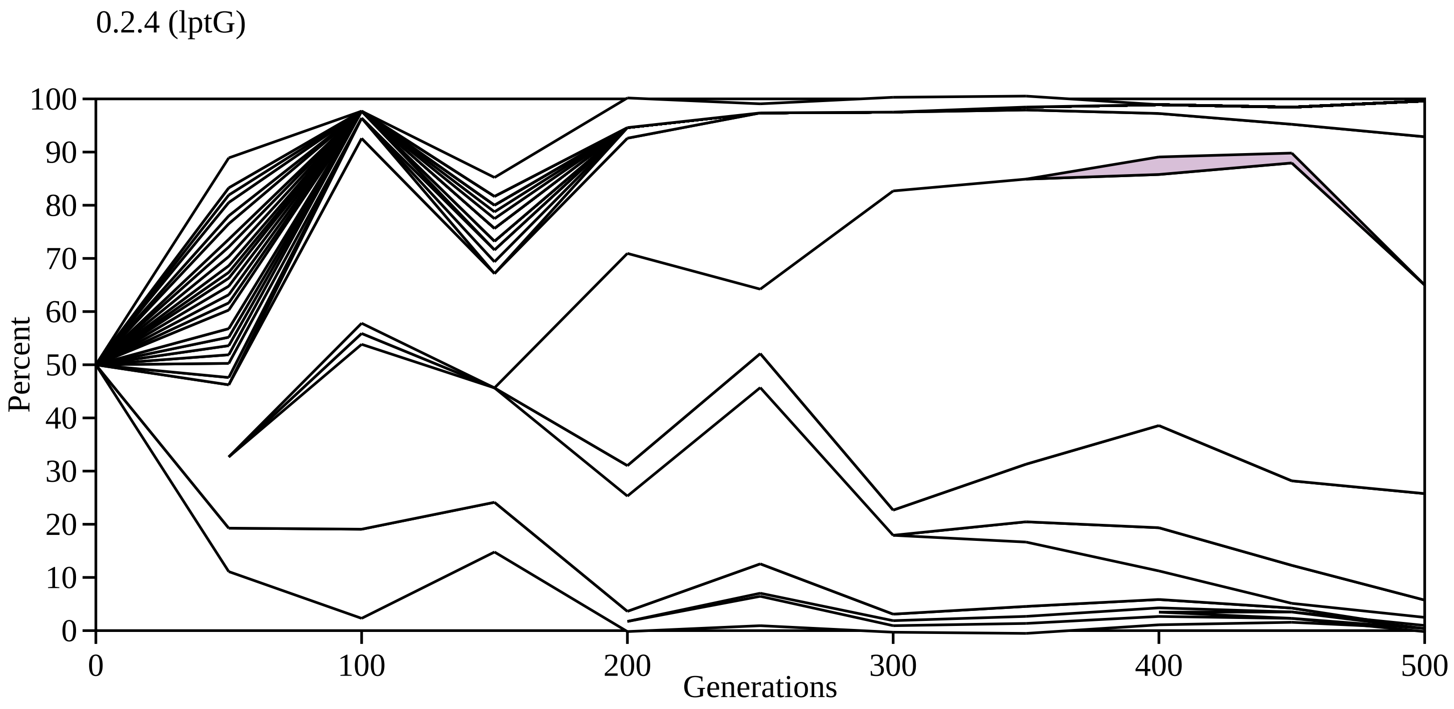

0.2.4 (lptG)
Percent
Generations
0
10
20
30
40
50
60
70
80
90
100
0
100
200
300
400
500

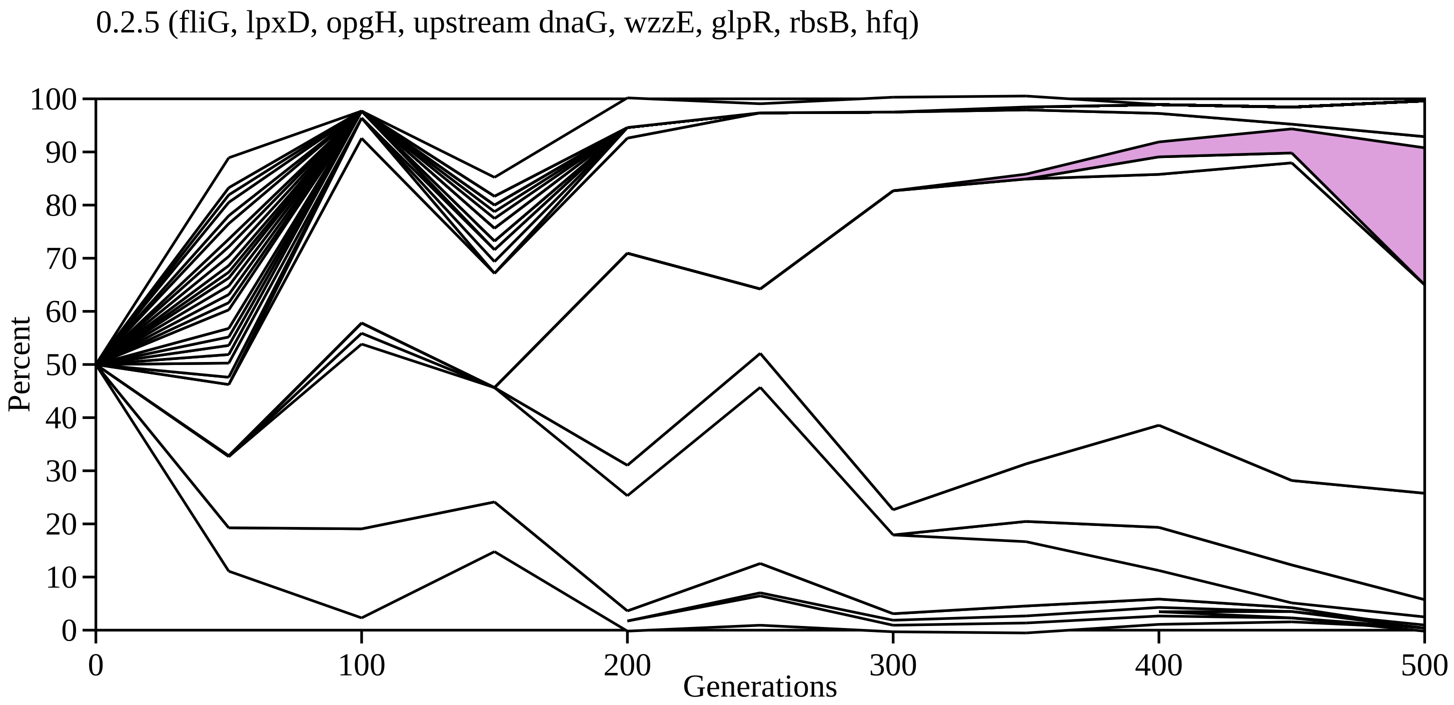

0.2.5 (fliG, lpxD, opgH, upstream dnaG, wzzE, glpR, rbsB, hfq)
Percent
0
10
20
30
40
50
60
70
80
90
100
Generations
0
100
200
300
400
500

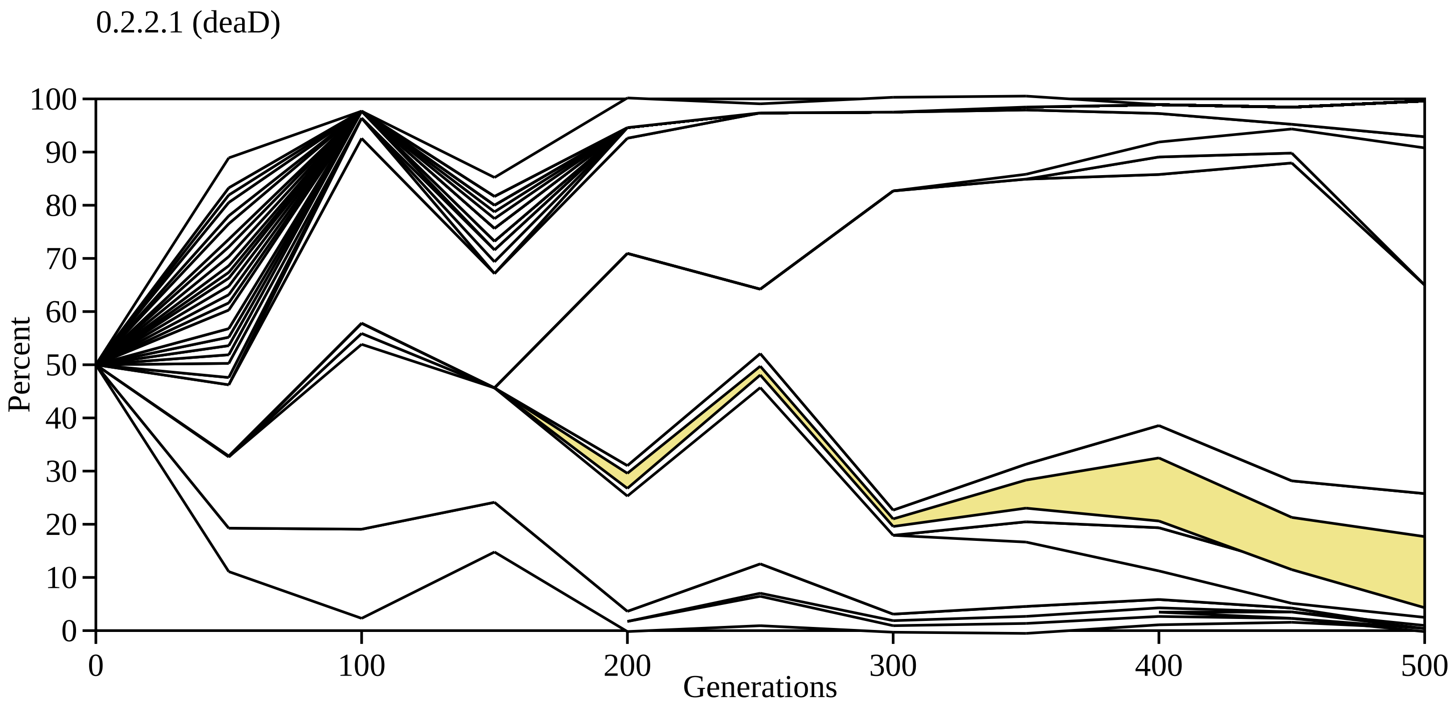
0.2.2.1 (deaD)
Percent
100
90
80
70
60
50
40
30
20
10
0
0
100
200
300
400
500
Generations

0.2.2.2 (fliH, rpoS, ybaL)

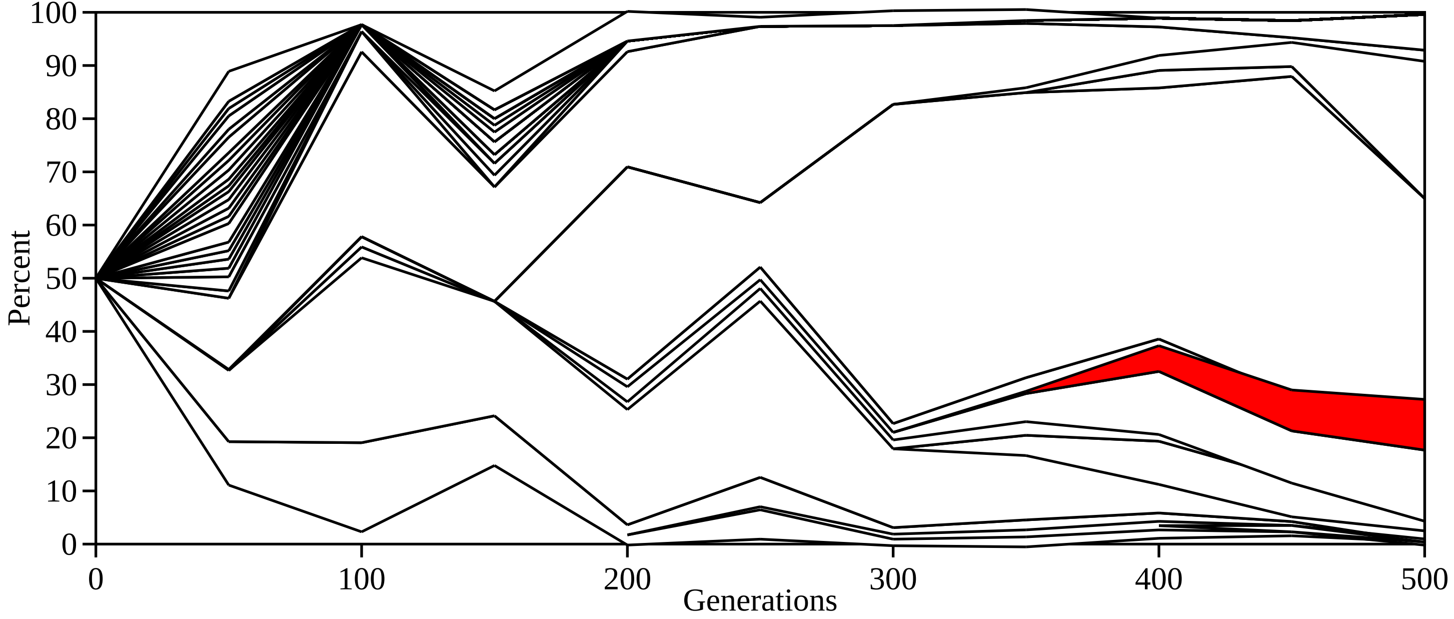

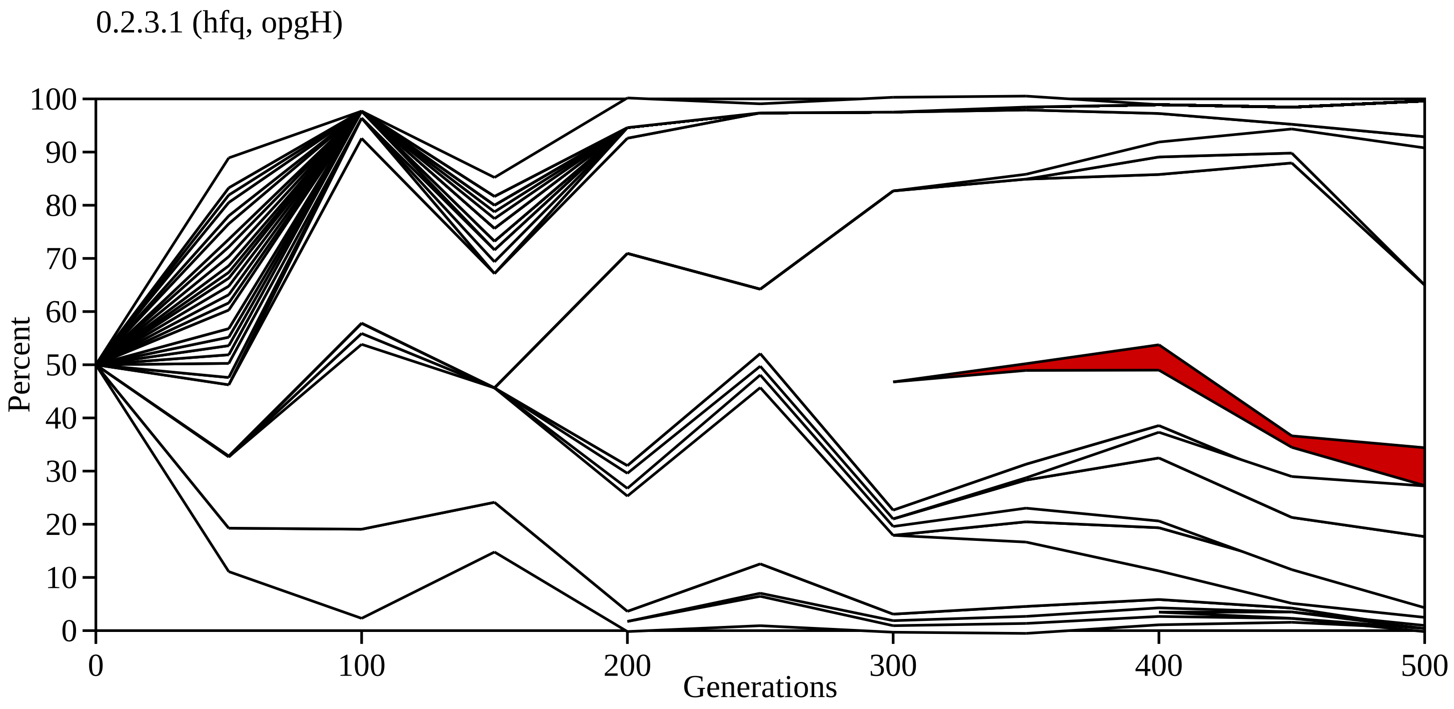
0.2.3.1 (hfq, opgH)
Percent
100
90
80
70
60
50
40
30
20
10
0
0
100
200
300
400
500
Generations

0.2.3.2 (lptG)

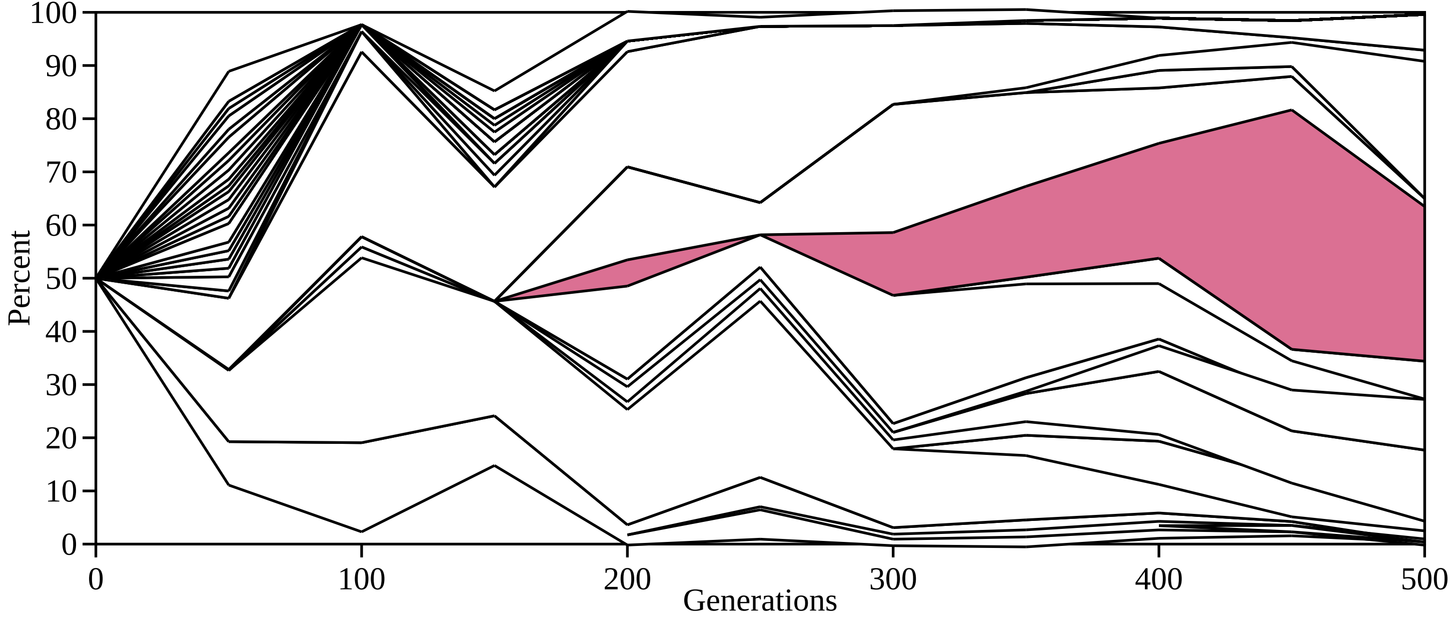

100
90
80
70
60
50
40
30
20
10
0
Percent
0
100
200
300
400
500
Generations

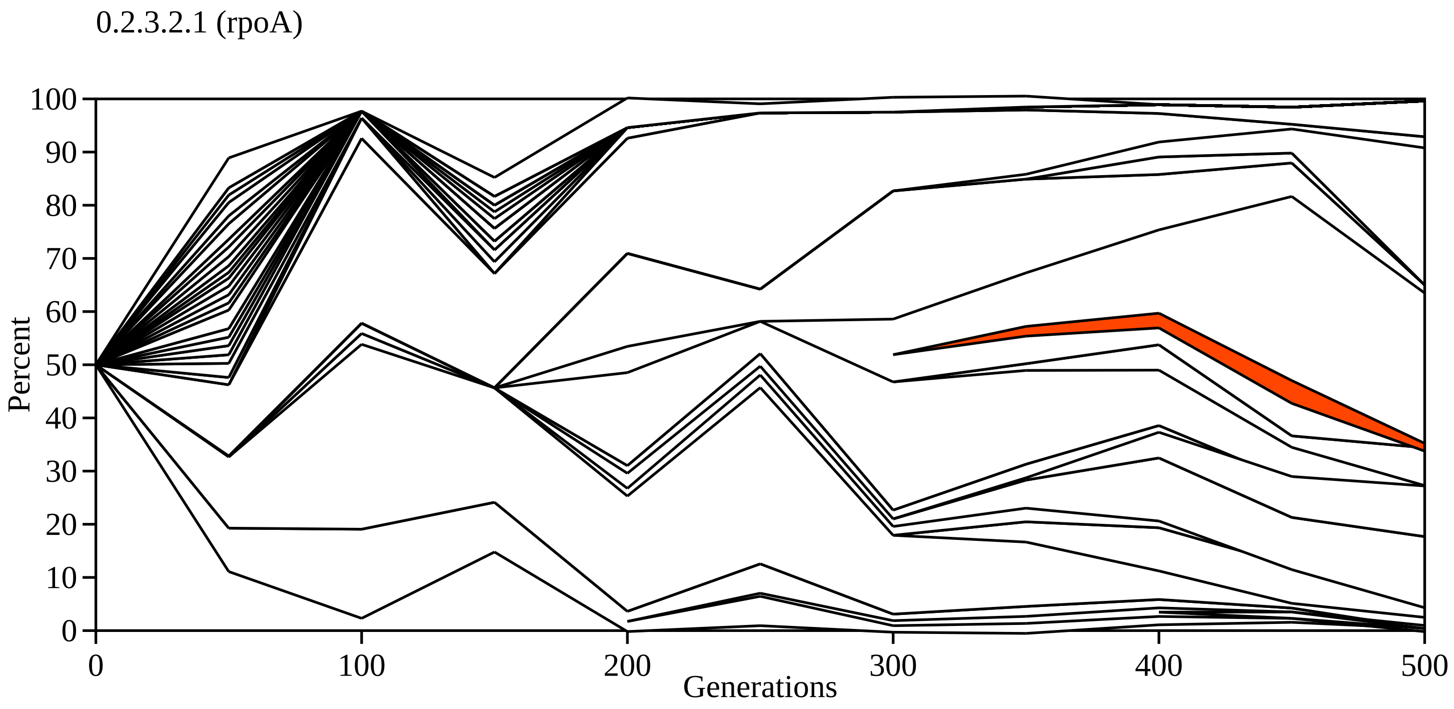
0.2.3.2.1 (rpoA)
Percent
100
90
80
70
60
50
40
30
20
10
0
0
100
200
300
400
500
Generations

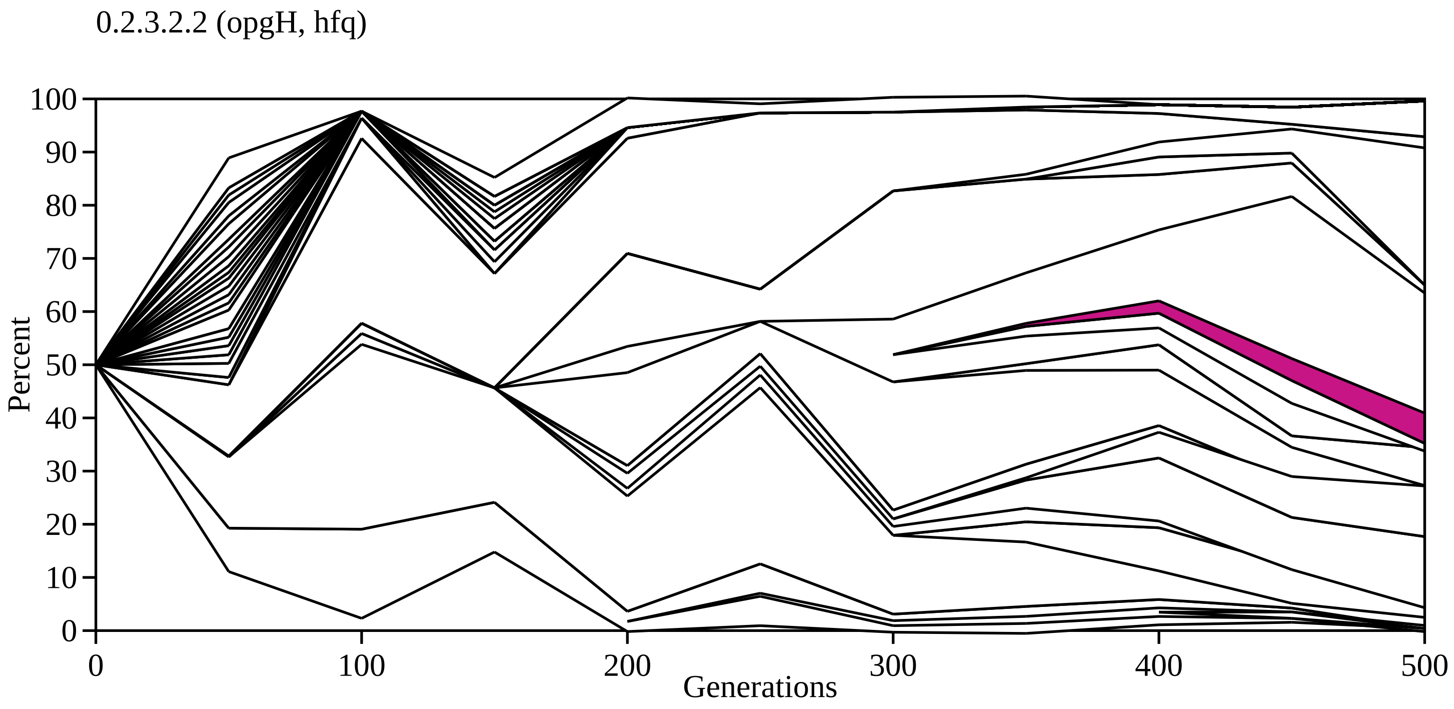

0.2.3.2.2 (opgH, hfq)
Percent
100
90
80
70
60
50
40
30
20
10
0
0
100
200
300
400
500
Generations

0.2.3.2.3 (opgG)

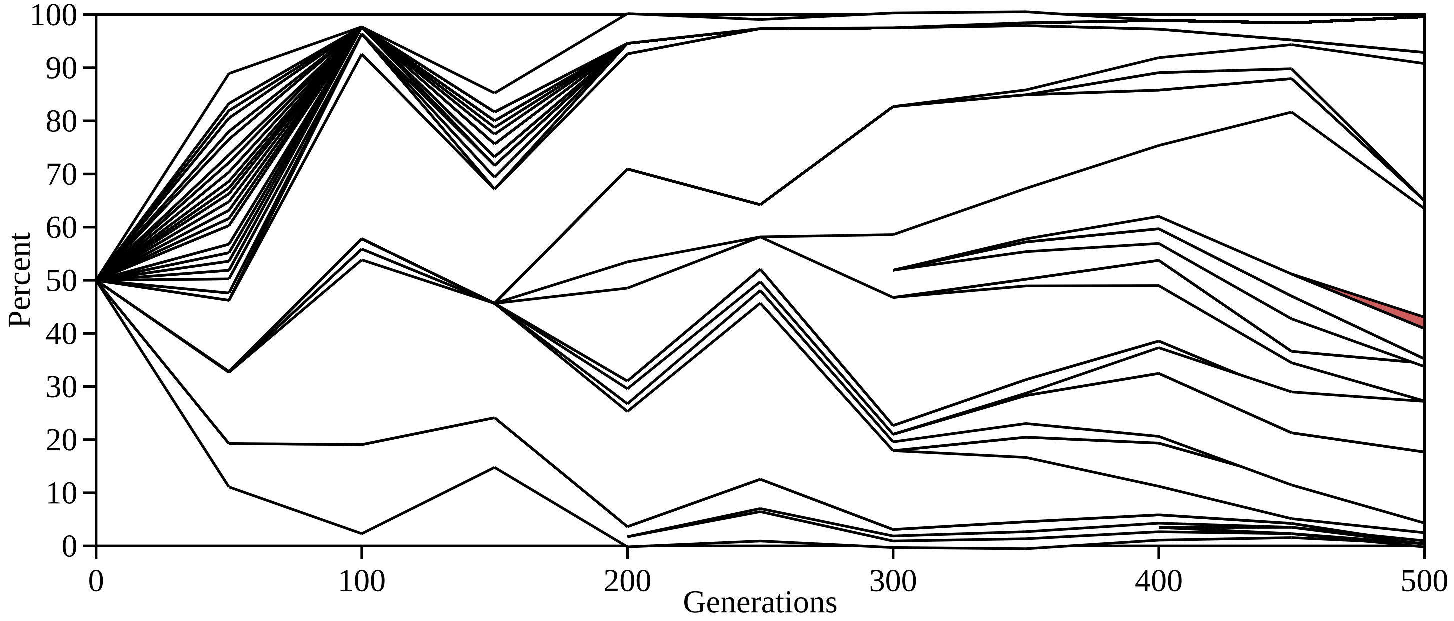

Percent
100
90
80
70
60
50
40
30
20
10
0
0
100
200
300
400
500
Generations

0.2.3.2.4 (rbsB, opgH)

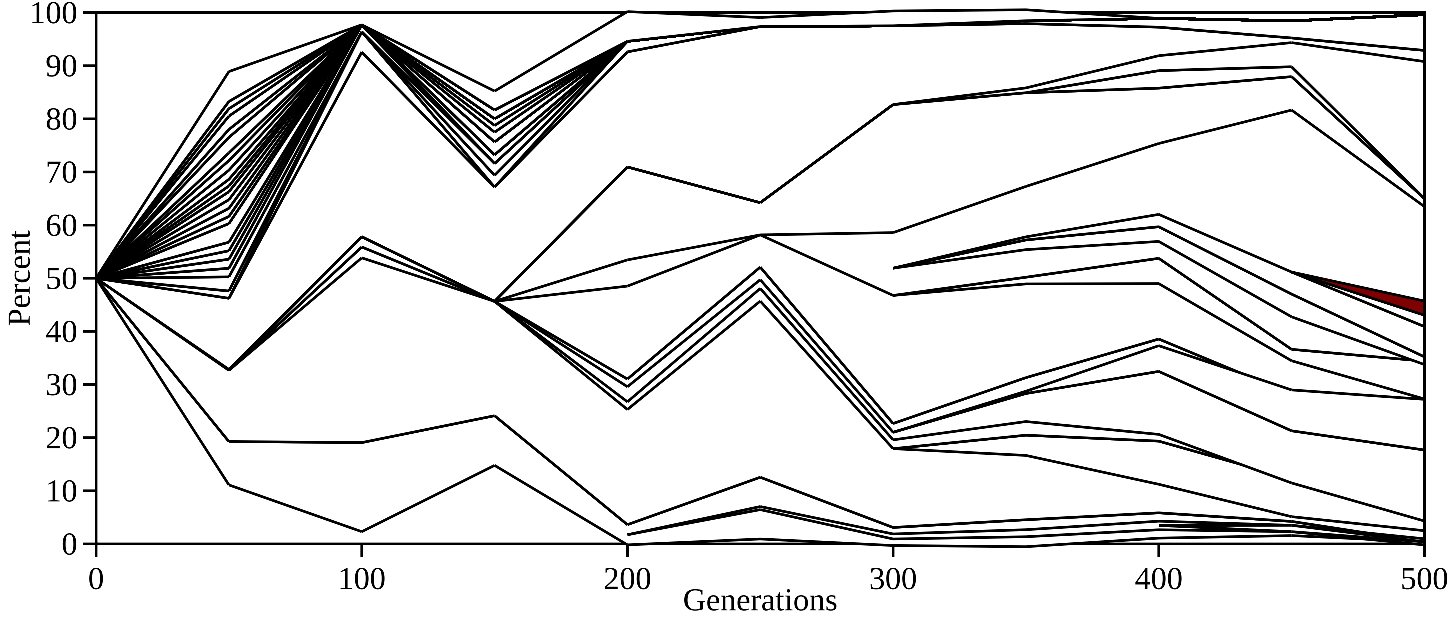

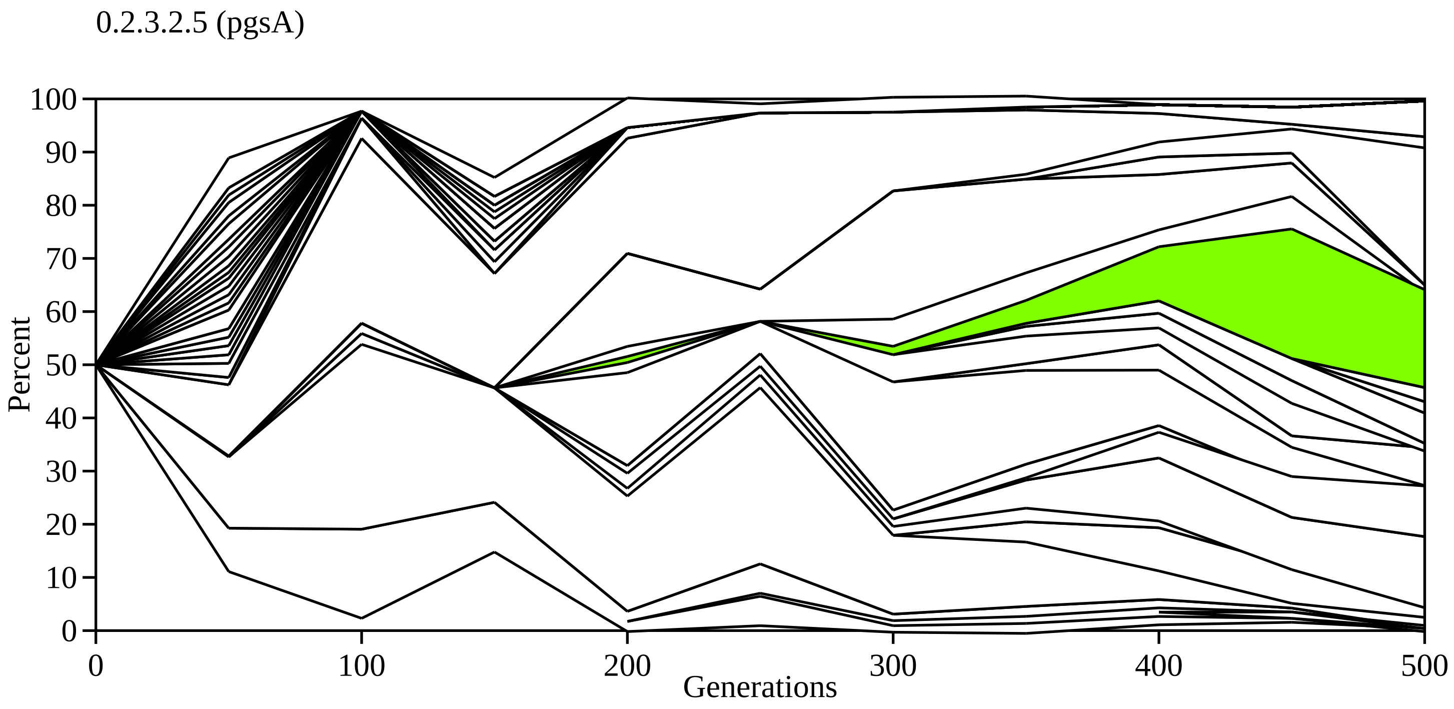
0.2.3.2.5 (pgsA)
Percent
100
90
80
70
60
50
40
30
20
10
0
0
100
200
300
400
500
Generations

0.2.3.2.1.1 (rpoS)

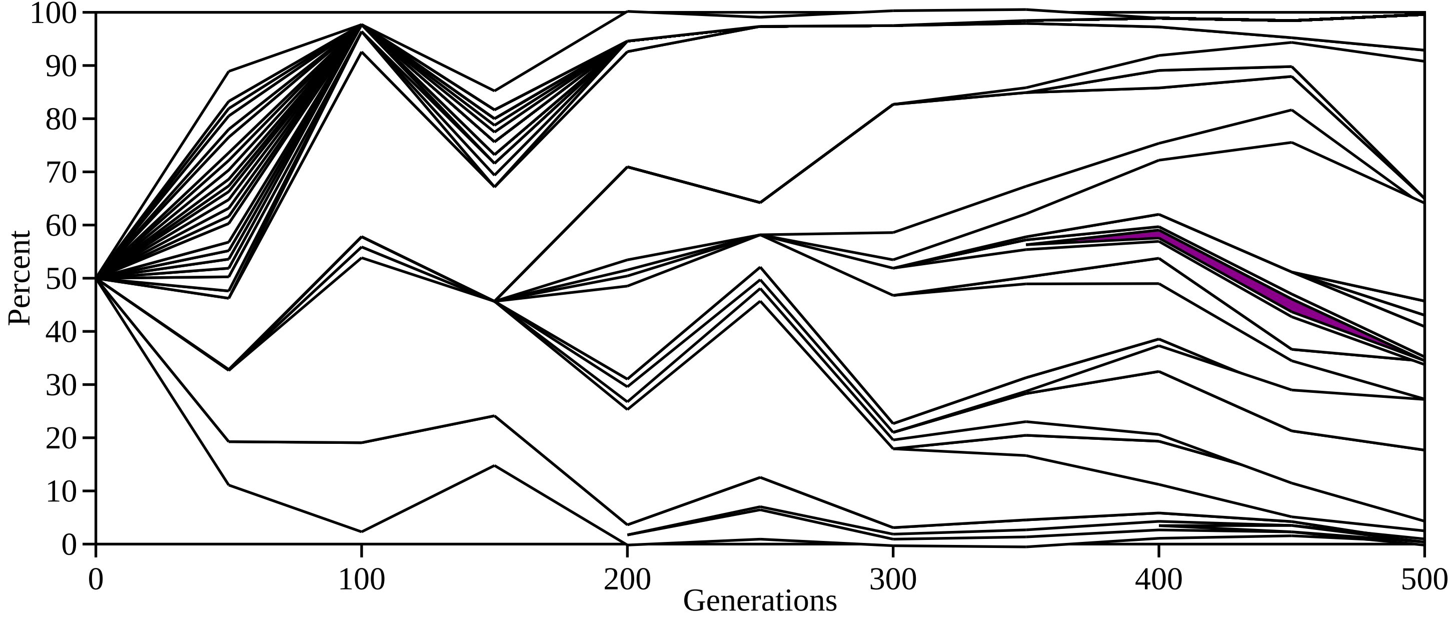

Percent
100
90
80
70
60
50
40
30
20
10
0
0
100
200
300
400
500
Generations

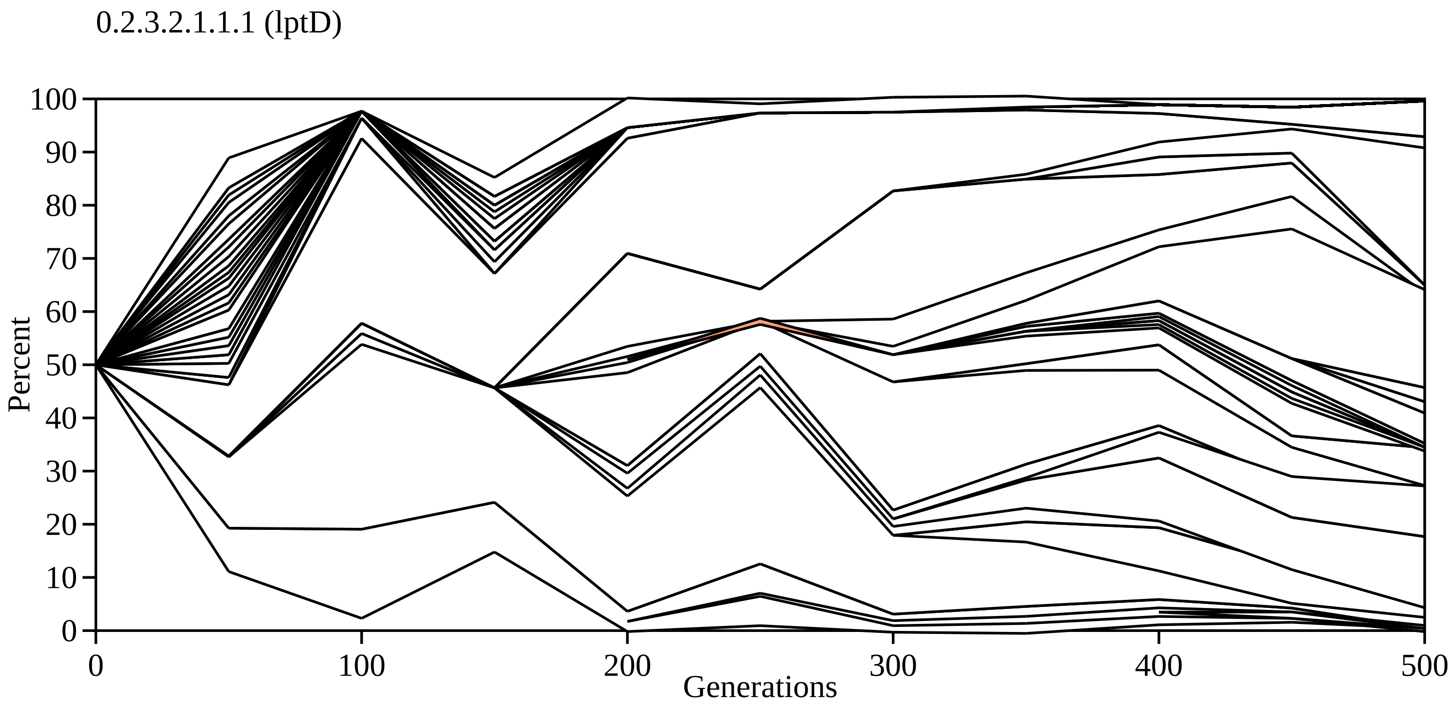
0.2.3.2.1.1.1 (lptD)
Percent
100
90
80
70
60
50
40
30
20
10
0
0
100
200
300
400
500
Generations

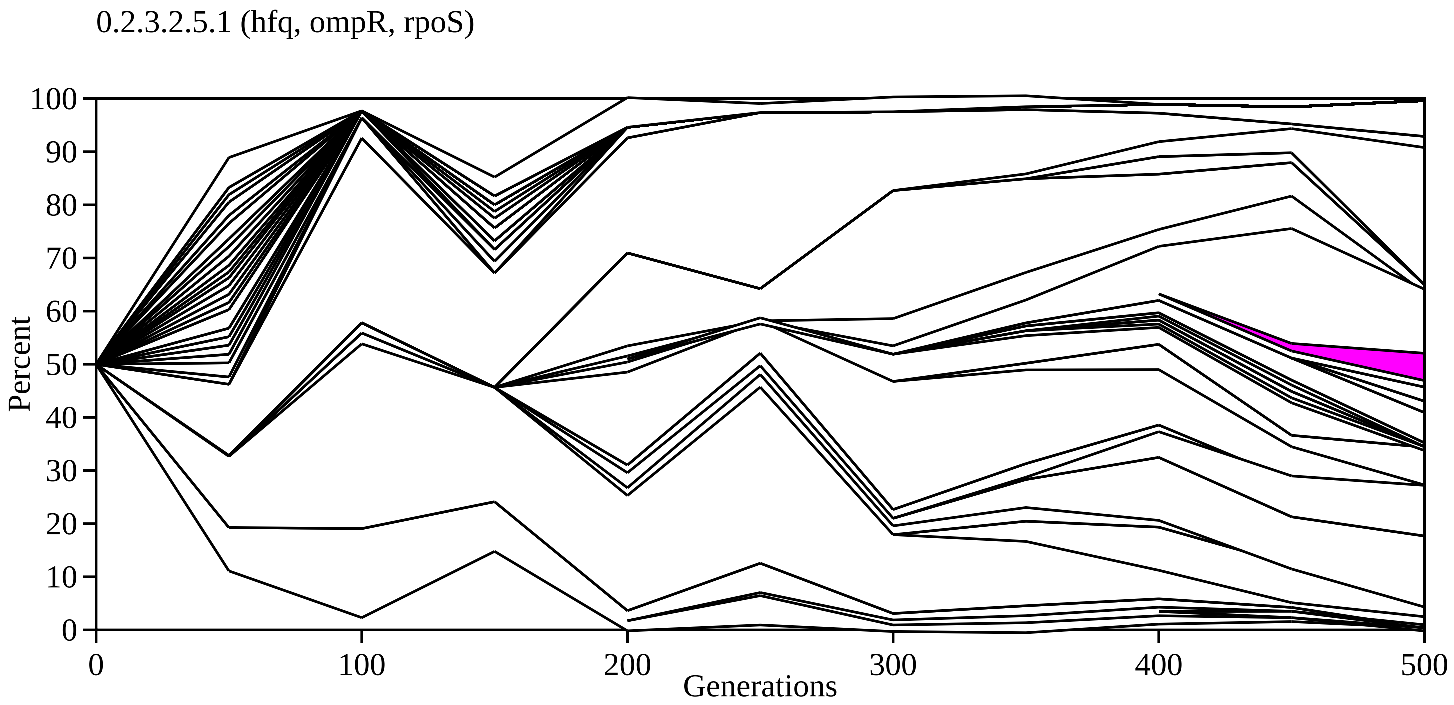
0.2.3.2.5.1 (hfq, ompR, rpoS)
Percent
0
10
20
30
40
50
60
70
80
90
100
0
100
200
300
400
500
Generations

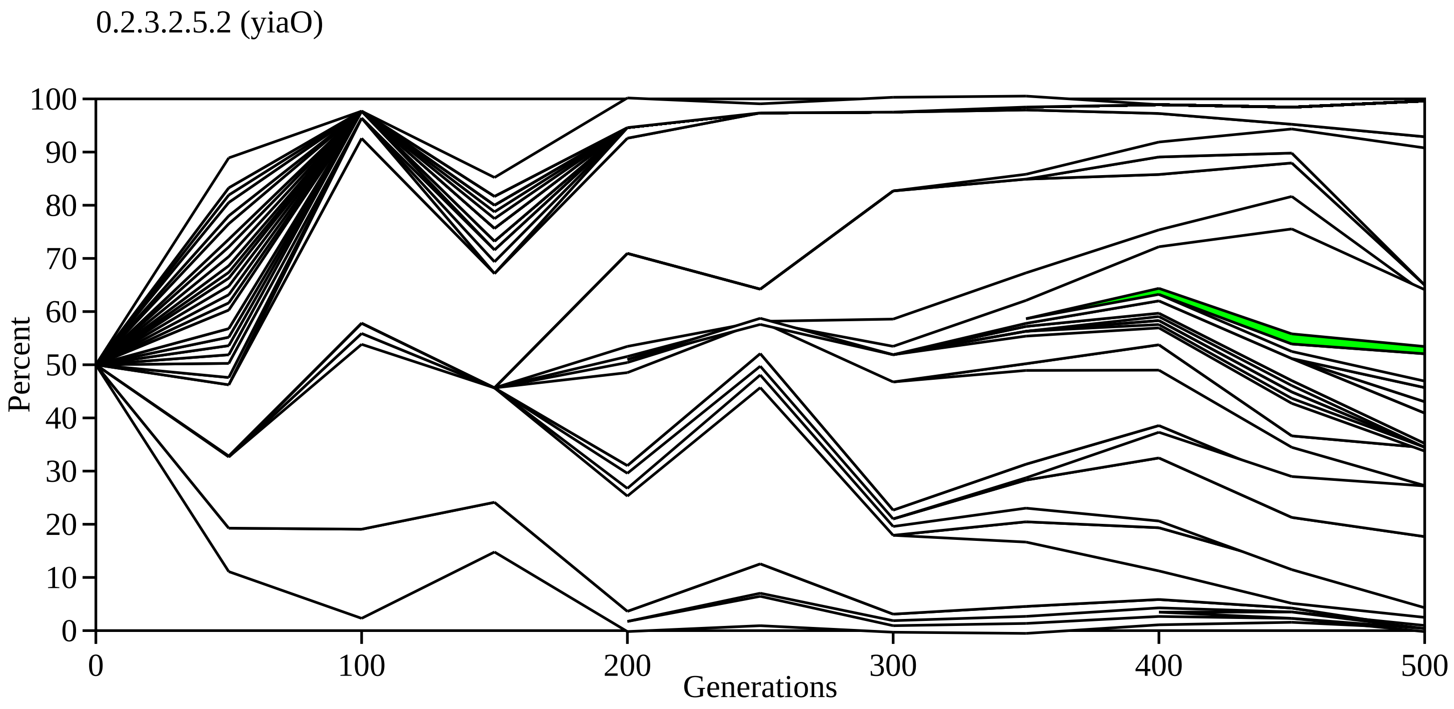

0.2.3.2.5.2 (yiaO)
Percent
100
90
80
70
60
50
40
30
20
10
0
0
100
200
300
400
500
Generations

0.2.3.2.5.3 (proQ, malT)

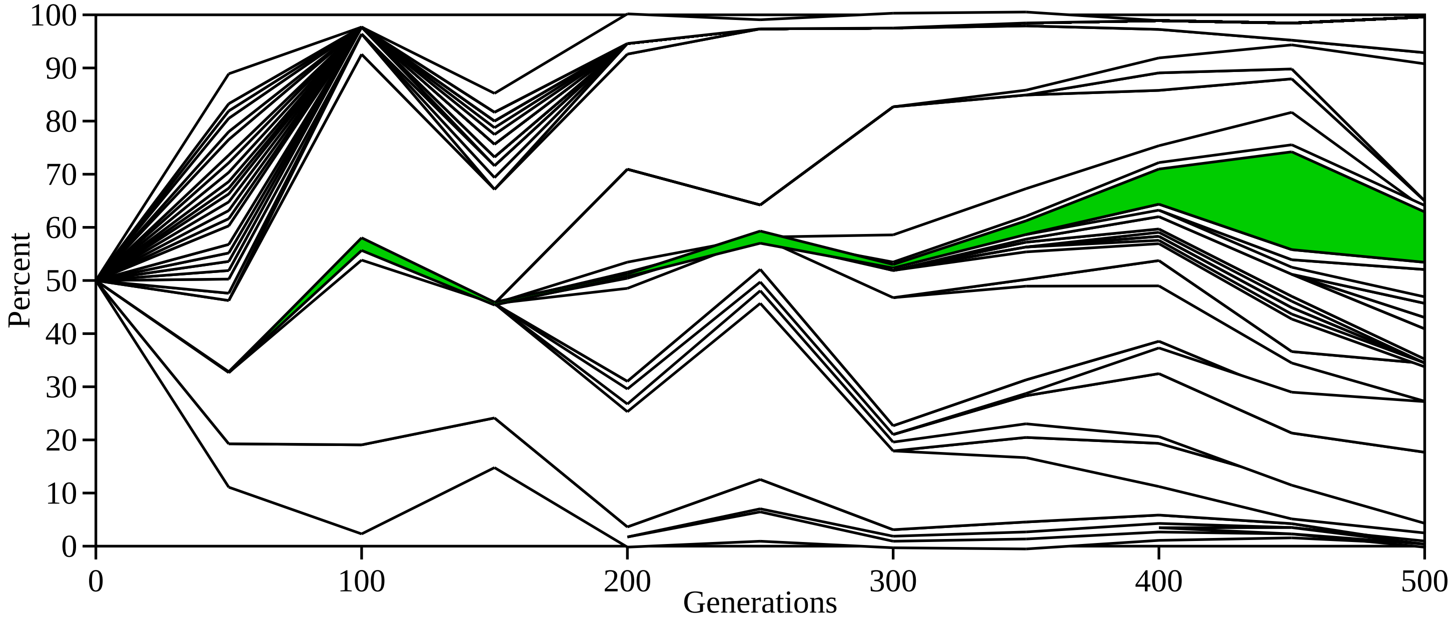

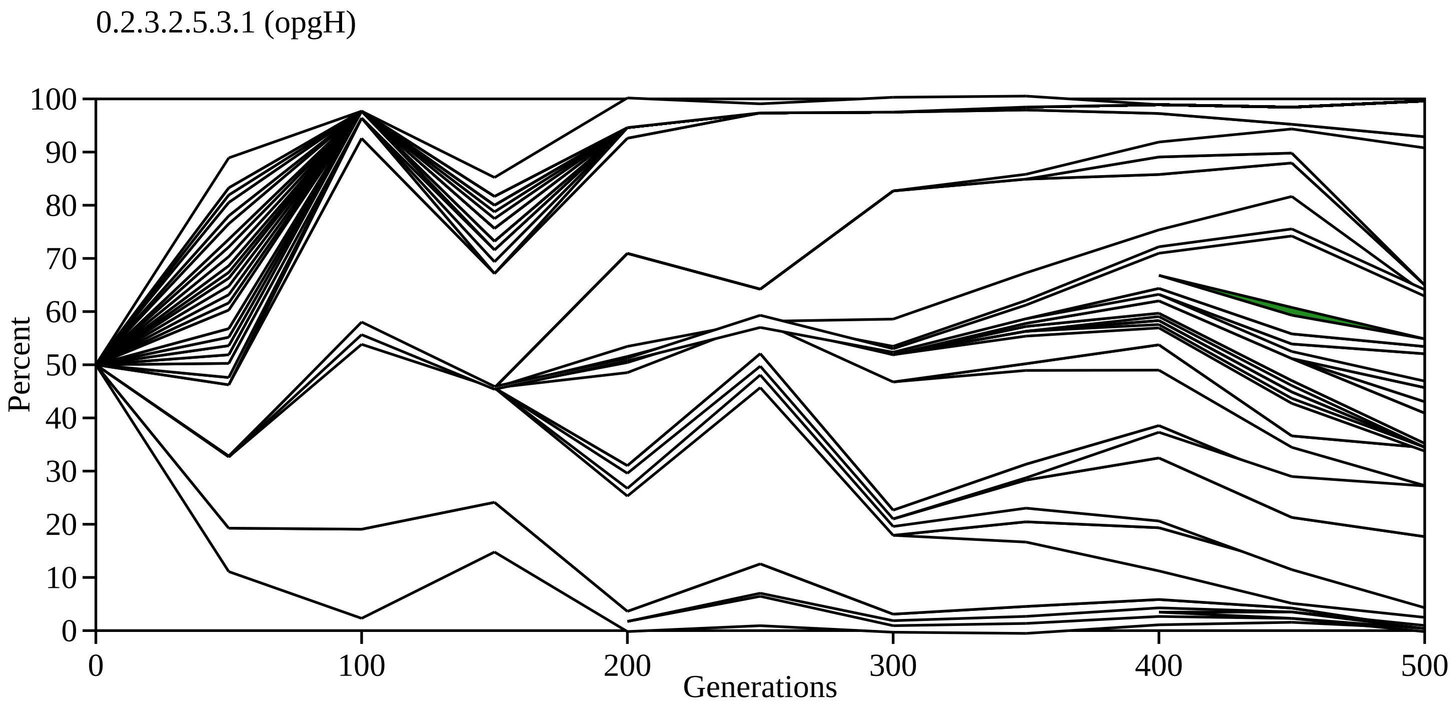

0.2.3.2.5.3.1 (opgH)
Percent
100
90
80
70
60
50
40
30
20
10
0
0
100
200
300
400
500
Generations

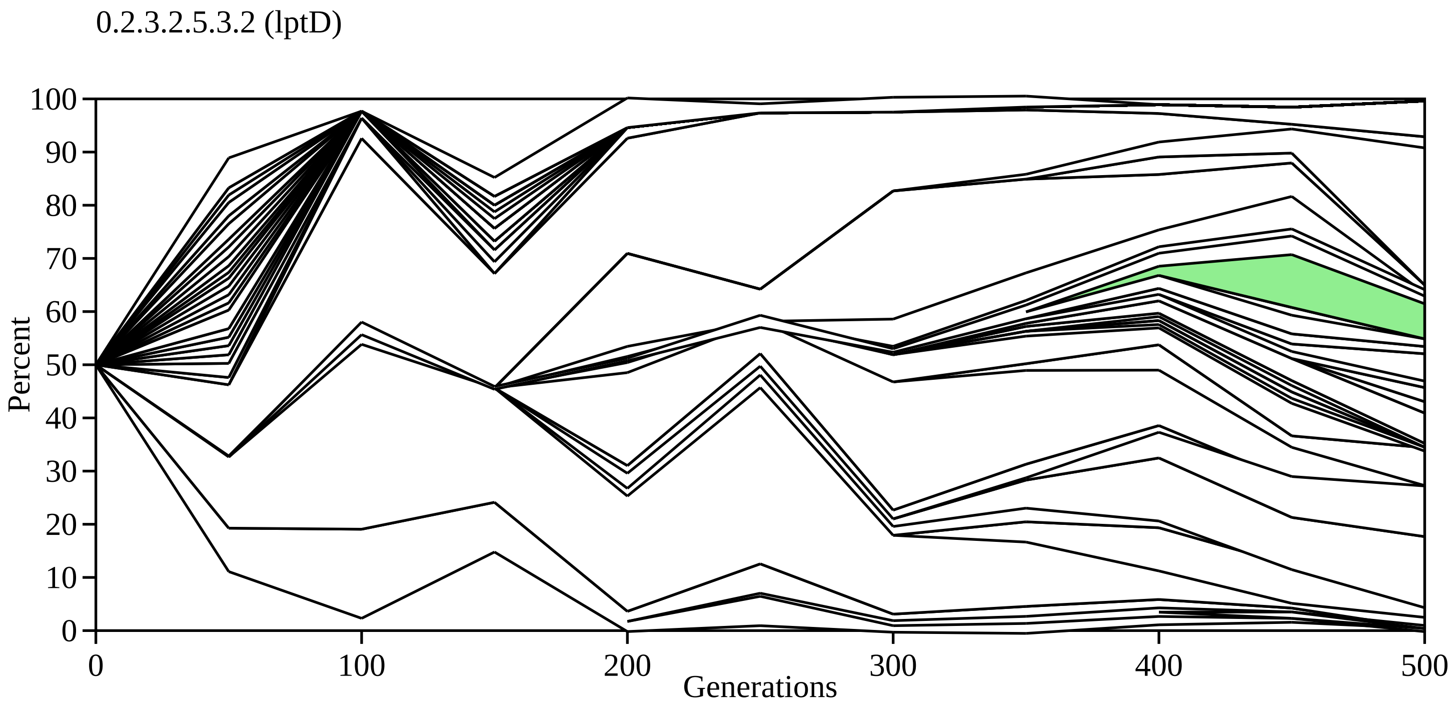

0.2.3.2.5.3.2 (lptD)
Percent
100
90
80
70
60
50
40
30
20
10
0
0
100
200
300
400
500
Generations

0.2.3.2.5.3.2.1 (yiaO)

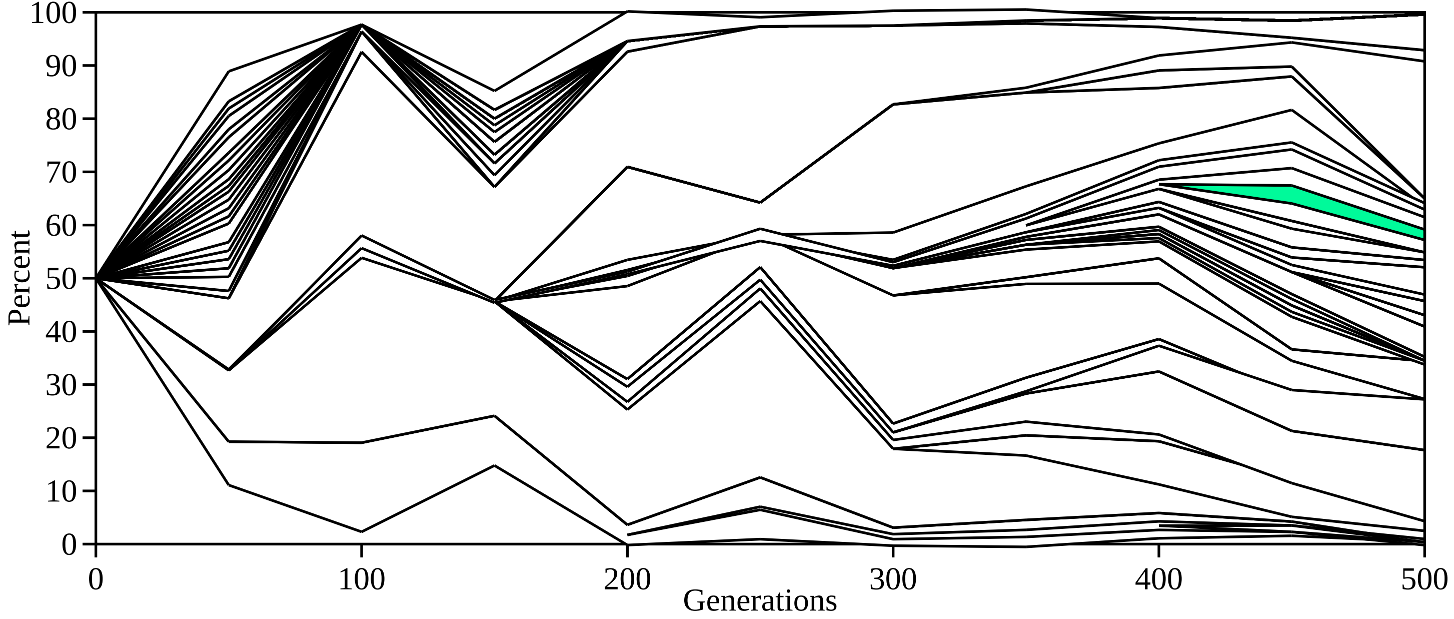

0.2.5.1 (upstream adhE)

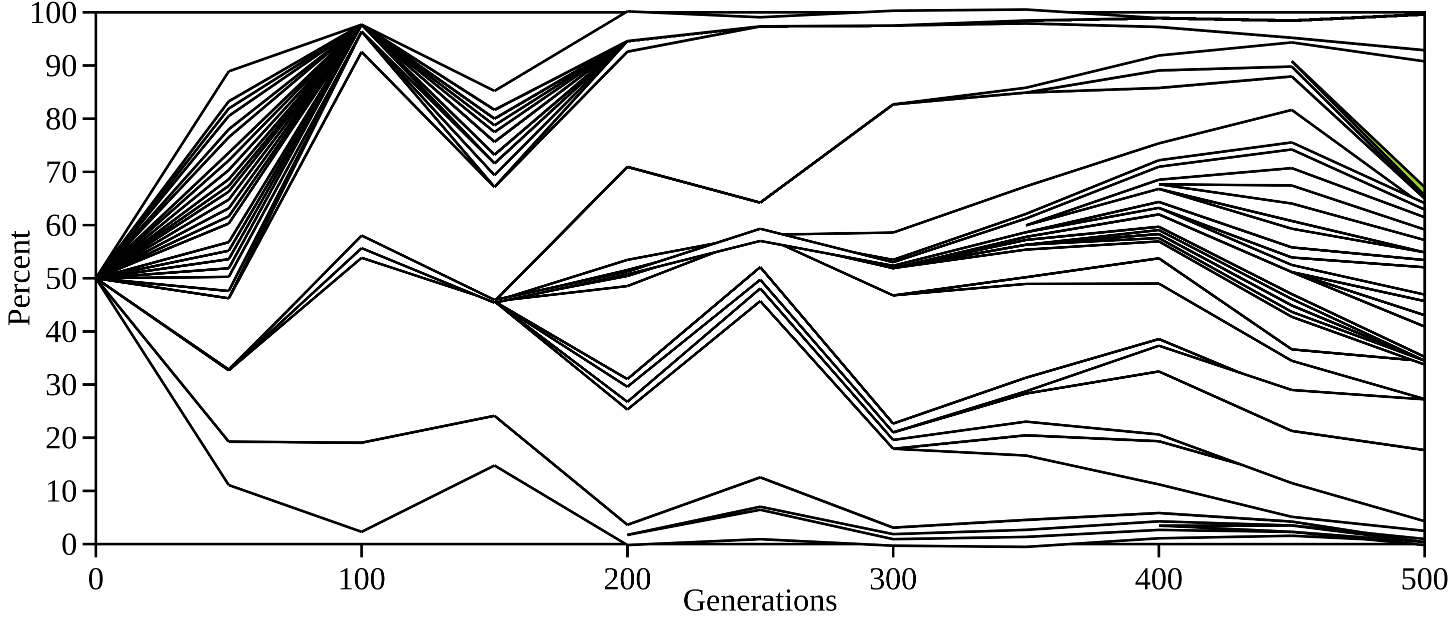

0.2.5.2 (gatZ, downstream fis)

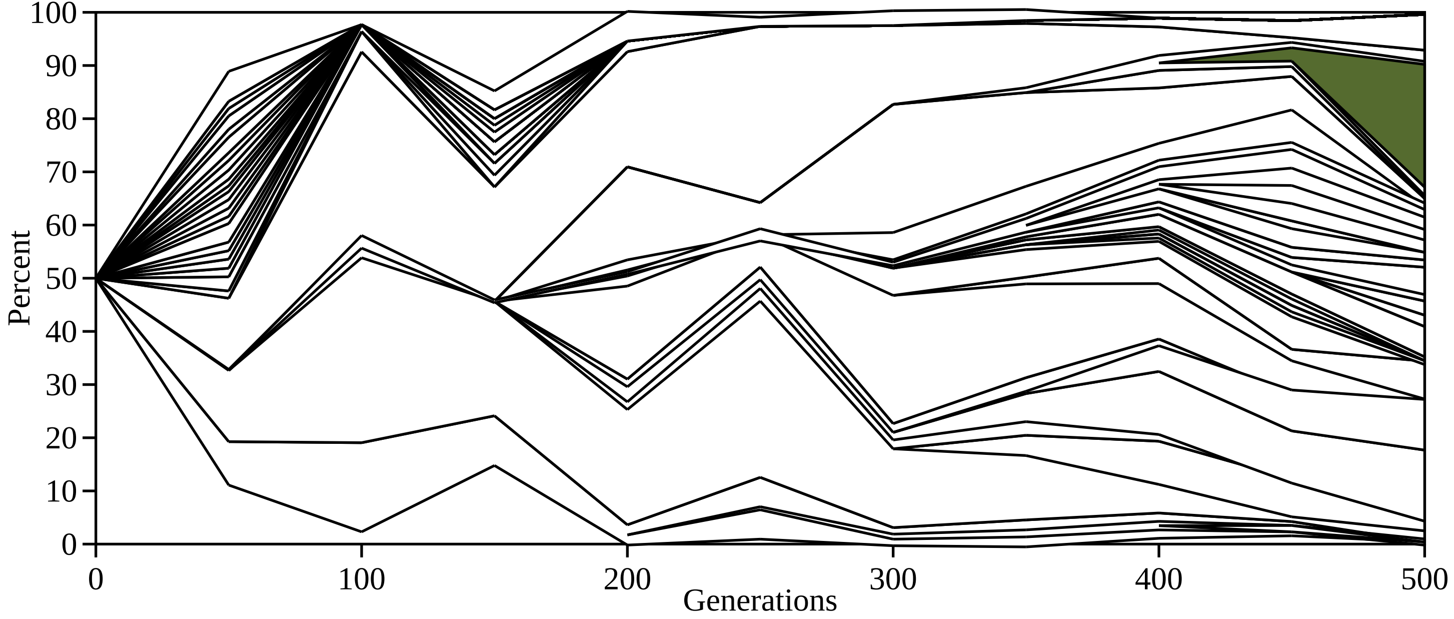

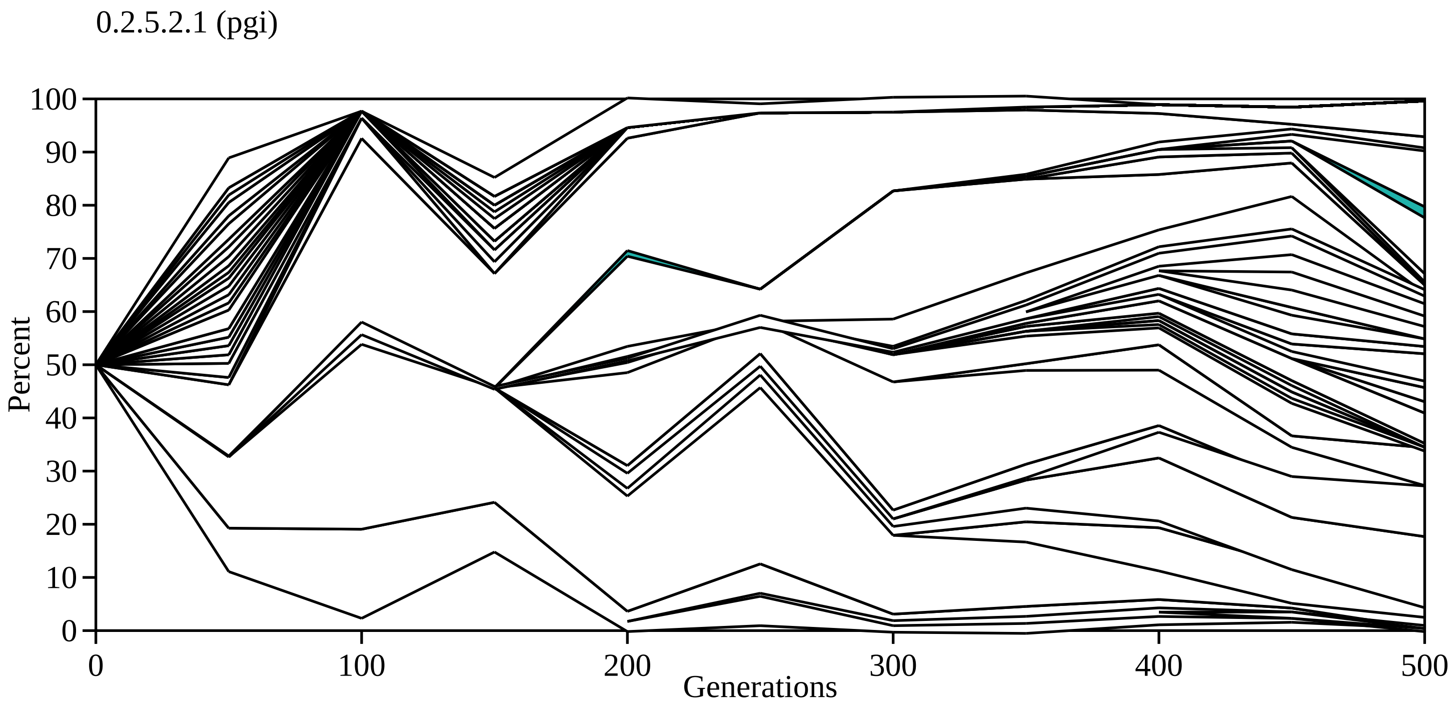

0.2.5.2.1 (pgi)
Percent
100
90
80
70
60
50
40
30
20
10
0
0
100
200
300
400
500
Generations

0.3.1 (rbsB, slt)

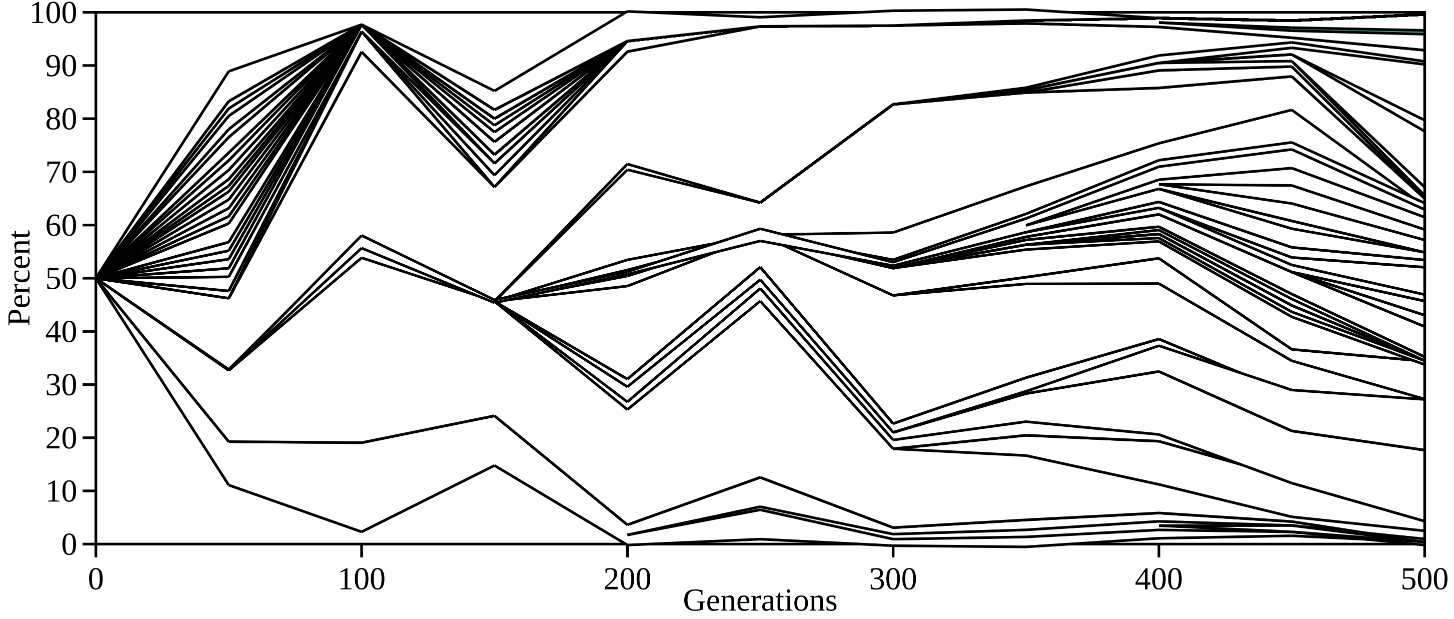

100
90
80
70
60
50
40
30
20
10
0
Percent
0
100
200
300
400
500
Generations
