## Supplementary Data File S5 for "Evolutionary dynamics of *de novo* mutations and mutant lineages arising in a simple, constant environment"

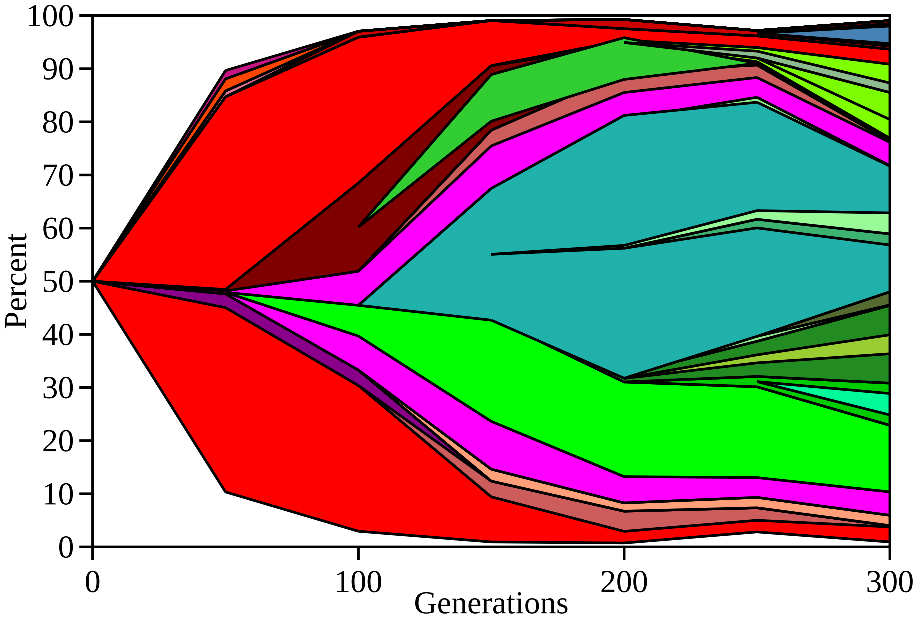

100
90
80
70
60
50
40
30
20
10
0
Percent
0
100
200
300
Generations

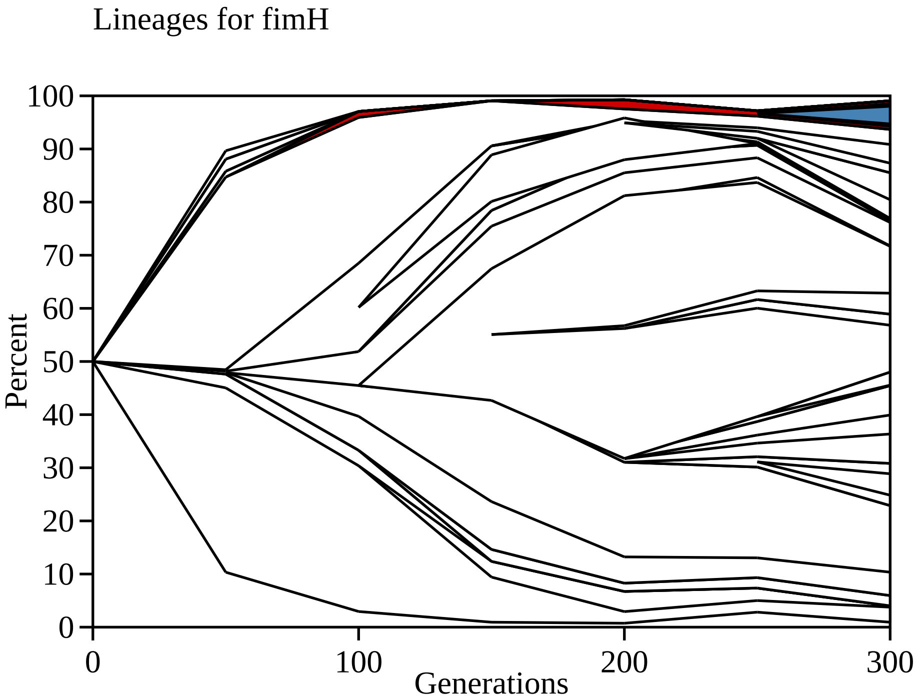
Lineages for fimH
Percent
100
90
80
70
60
50
40
30
20
10
0
0
100
200
300
Generations

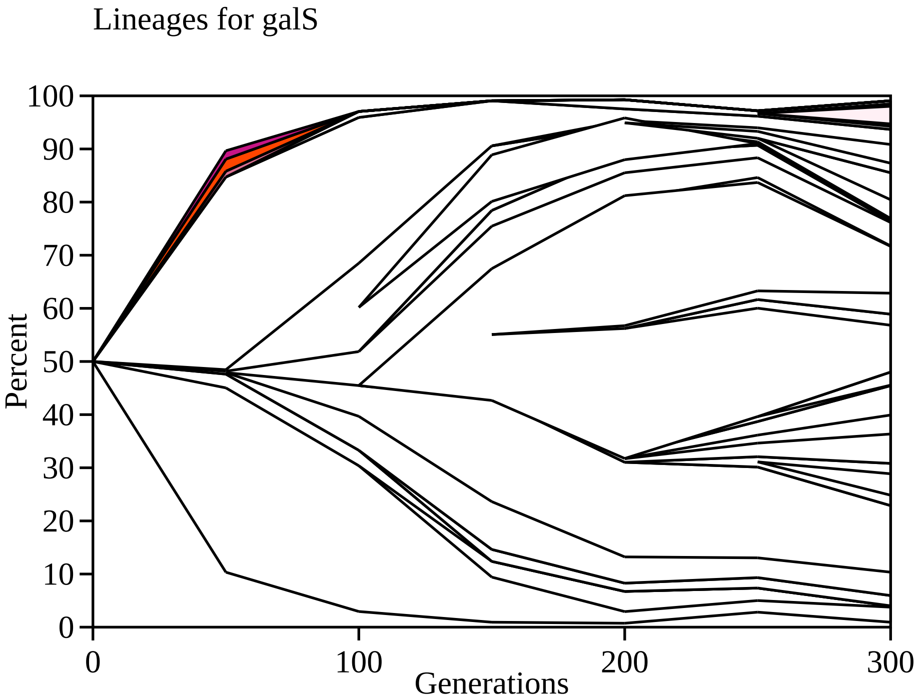
Lineages for galS
Percent
100
90
80
70
60
50
40
30
20
10
0
0
100
200
300
Generations

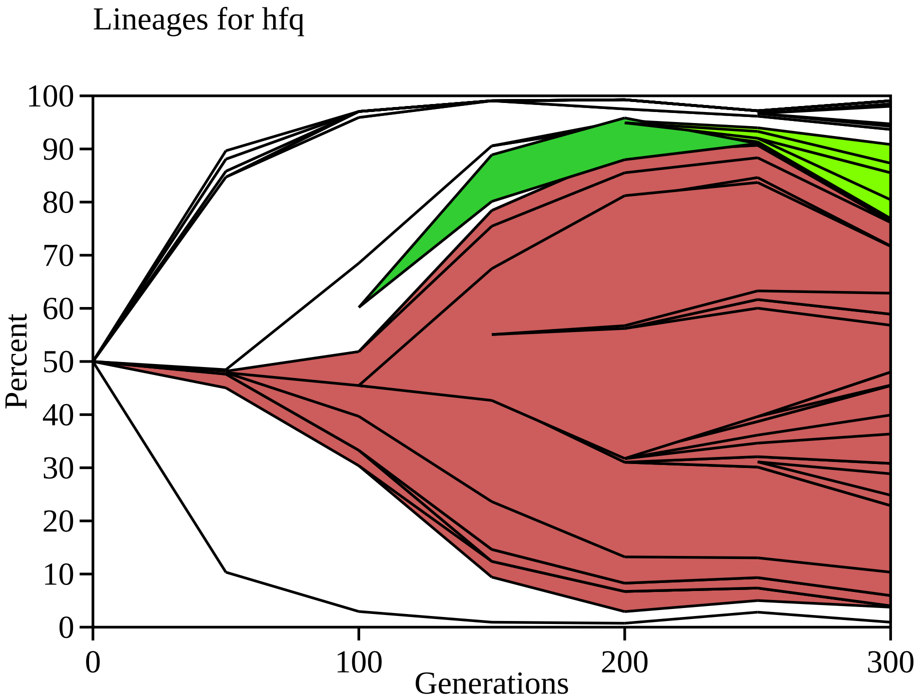
Lineages for hfq
Percent
100
90
80
70
60
50
40
30
20
10
0
0
100
200
300
Generations

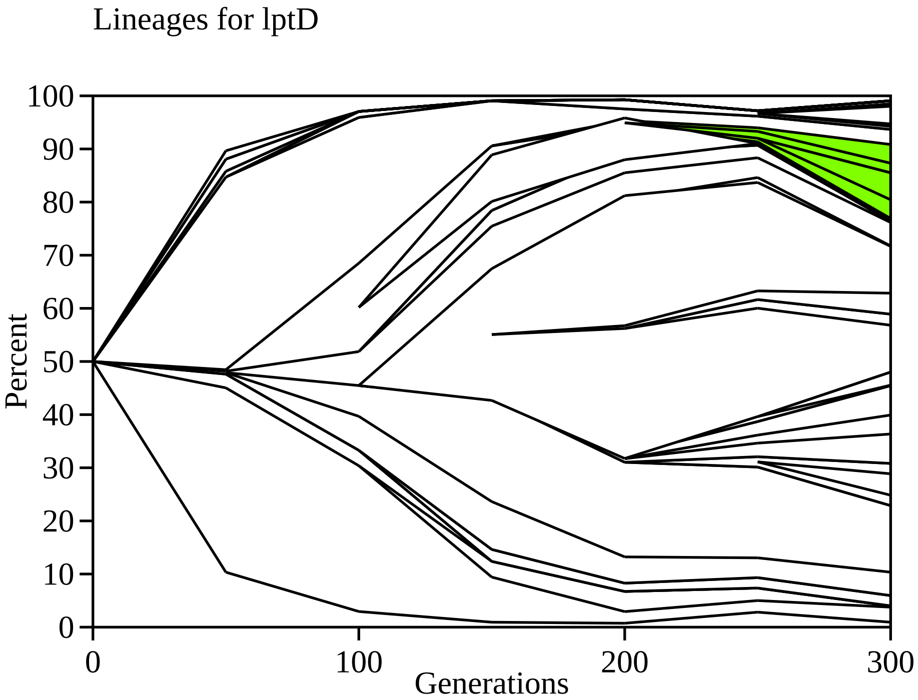

Lineages for lptD
Percent
100
90
80
70
60
50
40
30
20
10
0
0
100
200
300
Generations

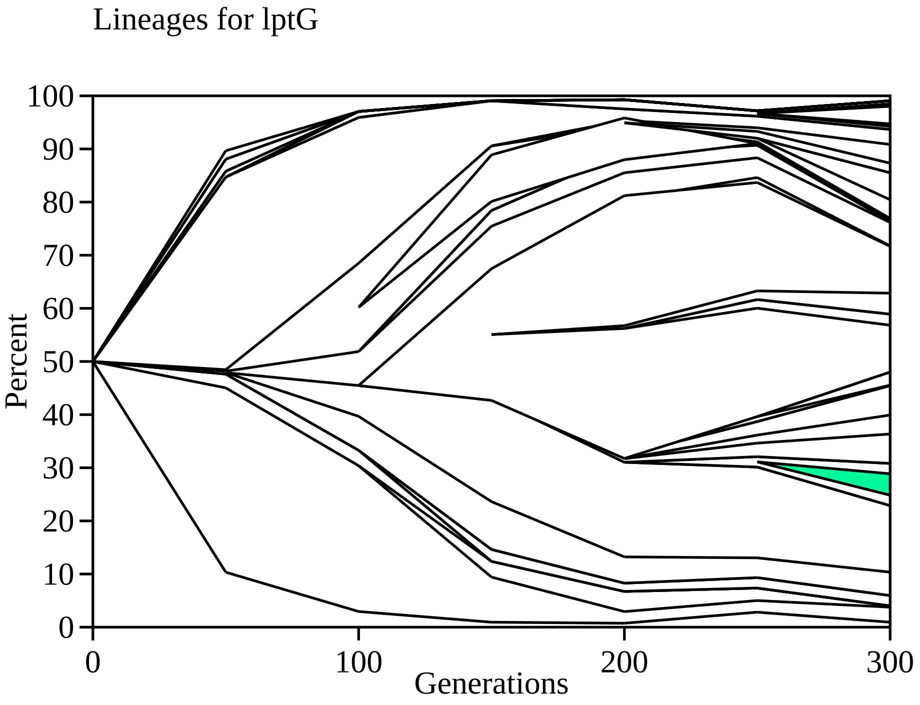
Lineages for lptG
100
90
80
70
60
50
40
30
20
10
0
Percent
0
100
200
300
Generations

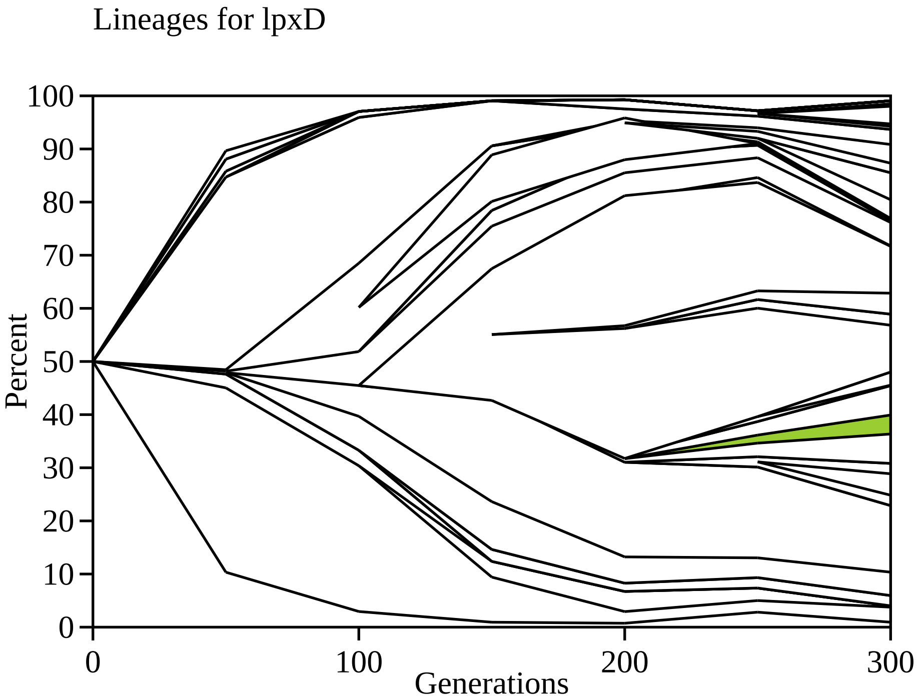

Lineages for lpxD
Percent
100
90
80
70
60
50
40
30
20
10
0
0
100
200
300
Generations

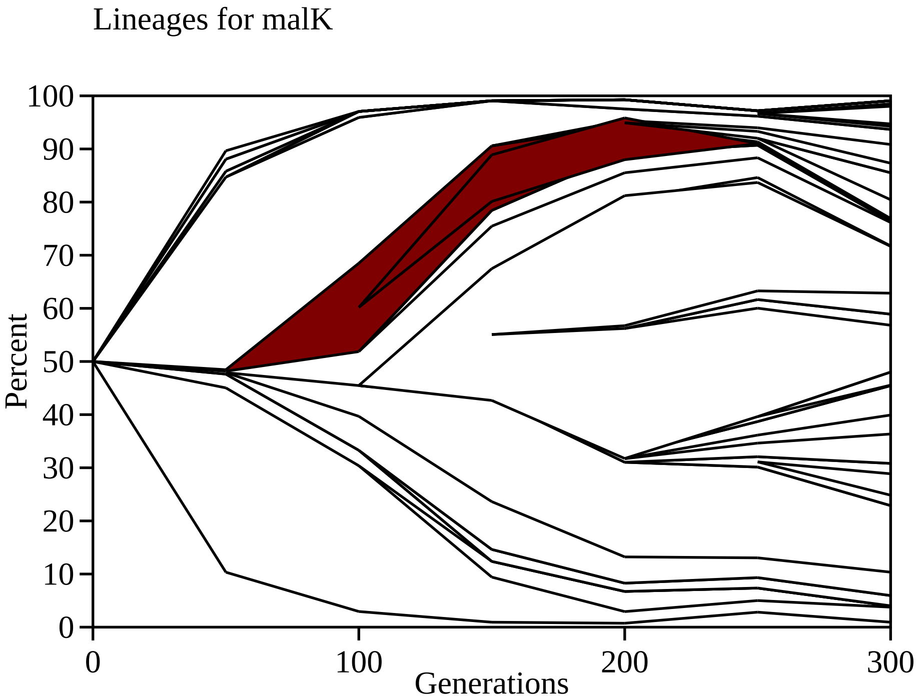
Lineages for malK
Percent
100
90
80
70
60
50
40
30
20
10
0
0
100
200
300
Generations

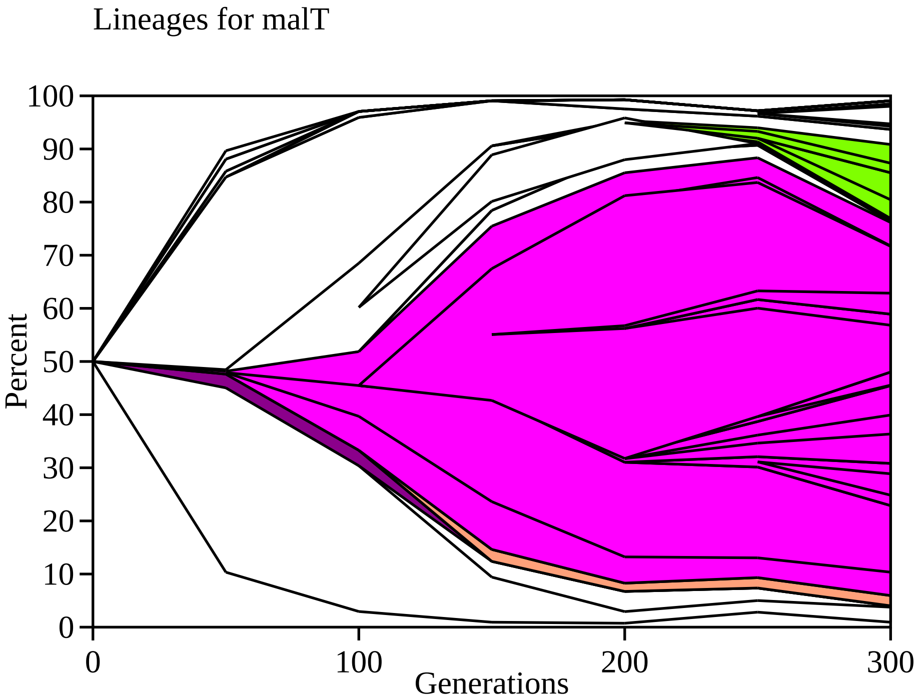

Lineages for malT
Percent
Generations
0
10
20
30
40
50
60
70
80
90
100
0
100
200
300

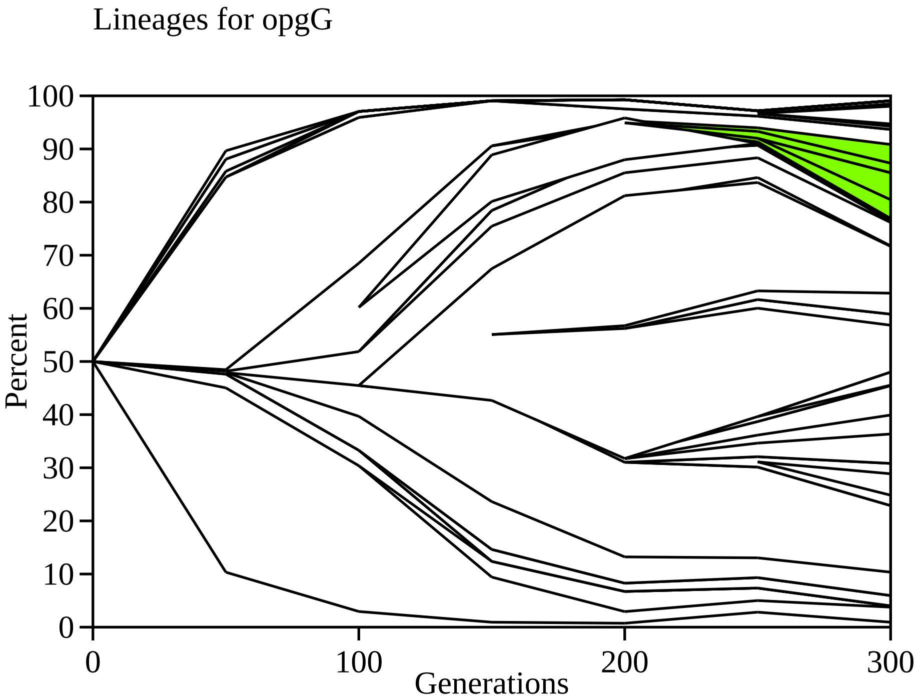
Lineages for opgG
100
90
80
70
60
50
40
30
20
10
0
Percent
0
100
200
300
Generations

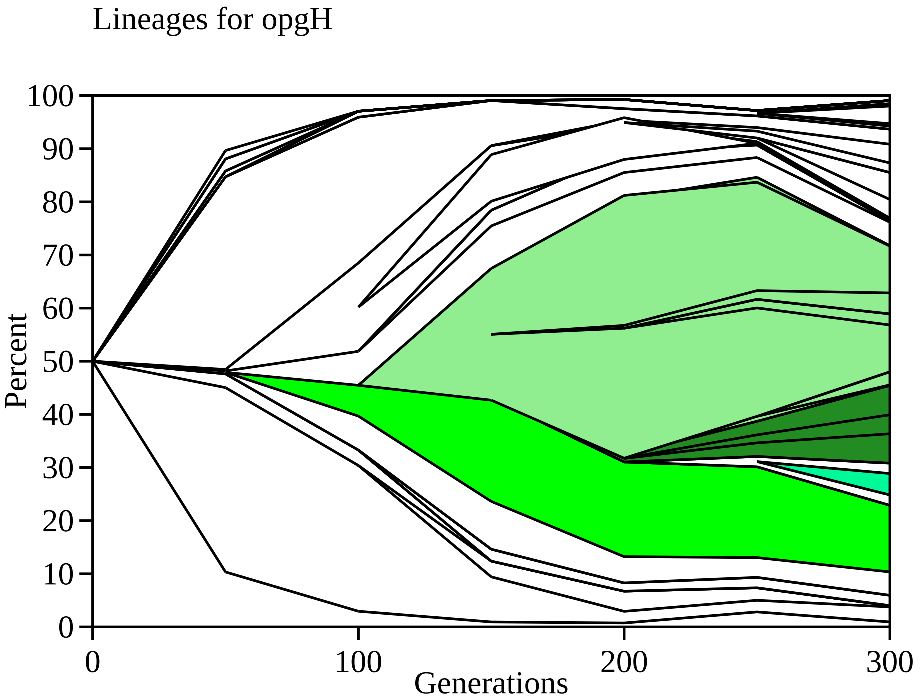
Lineages for opgH
Percent
100
90
80
70
60
50
40
30
20
10
0
0
100
200
300
Generations

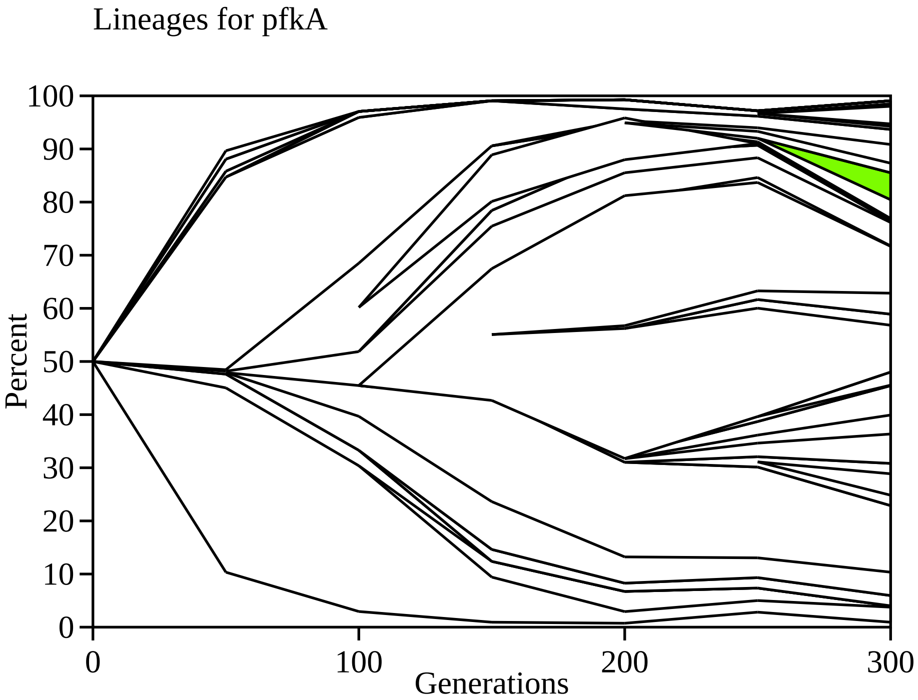

Lineages for pfkA
Percent
Generations
100
90
80
70
60
50
40
30
20
10
0
0
100
200
300

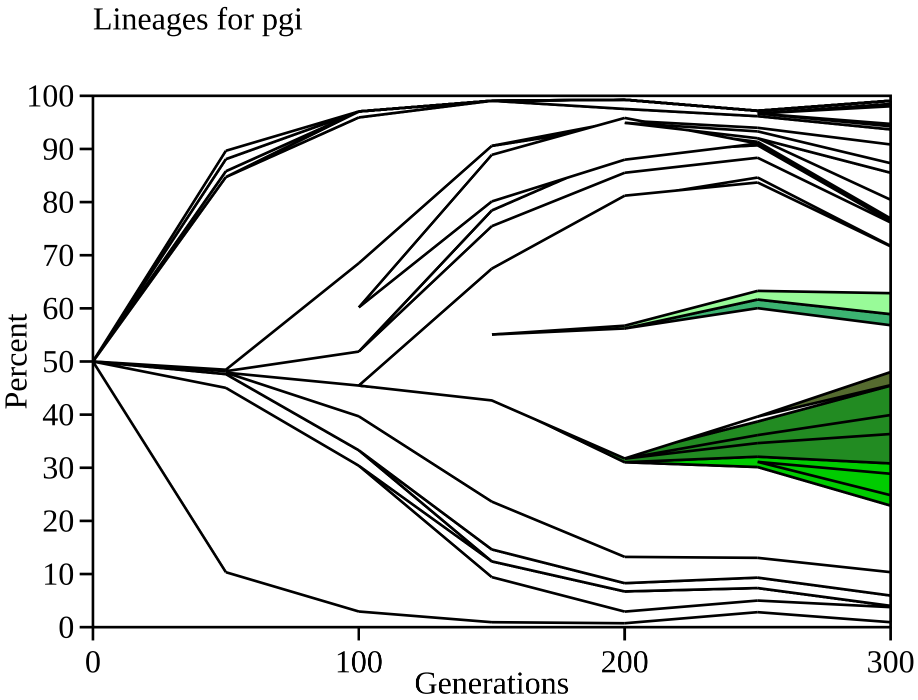
Lineages for pgi
Percent
0
10
20
30
40
50
60
70
80
90
100
0
100
200
300
Generations

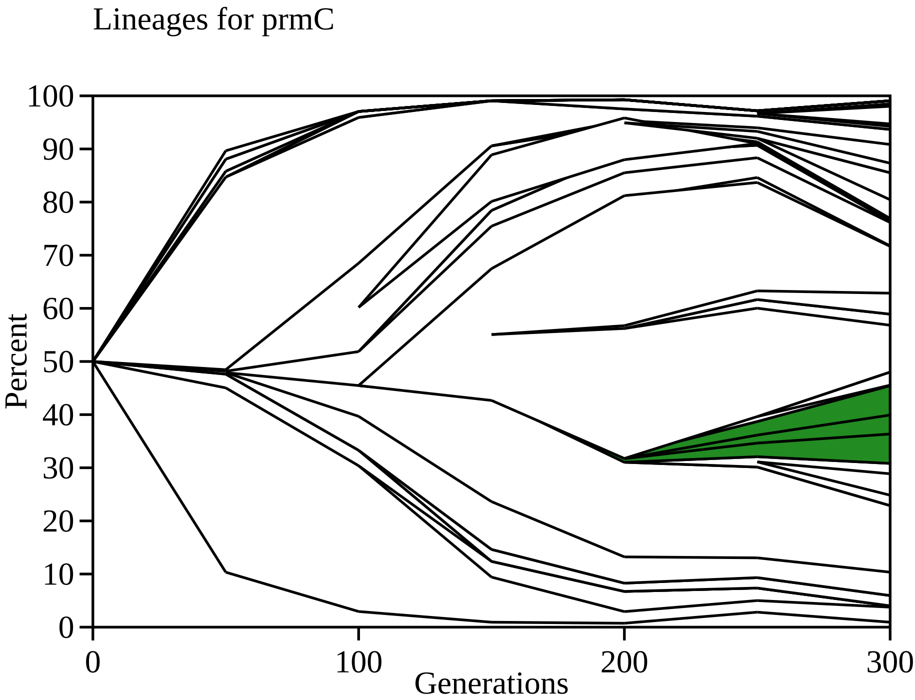

Lineages for prmC
Percent
Generations
0
10
20
30
40
50
60
70
80
90
100
0
100
200
300

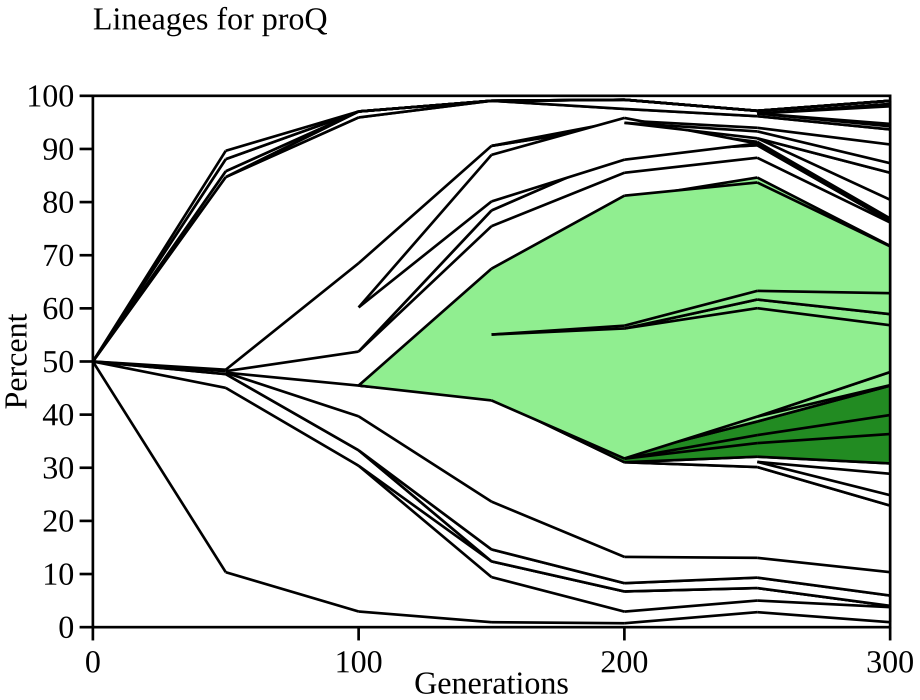
Lineages for proQ
Percent
Generations
100
90
80
70
60
50
40
30
20
10
0
0
100
200
300

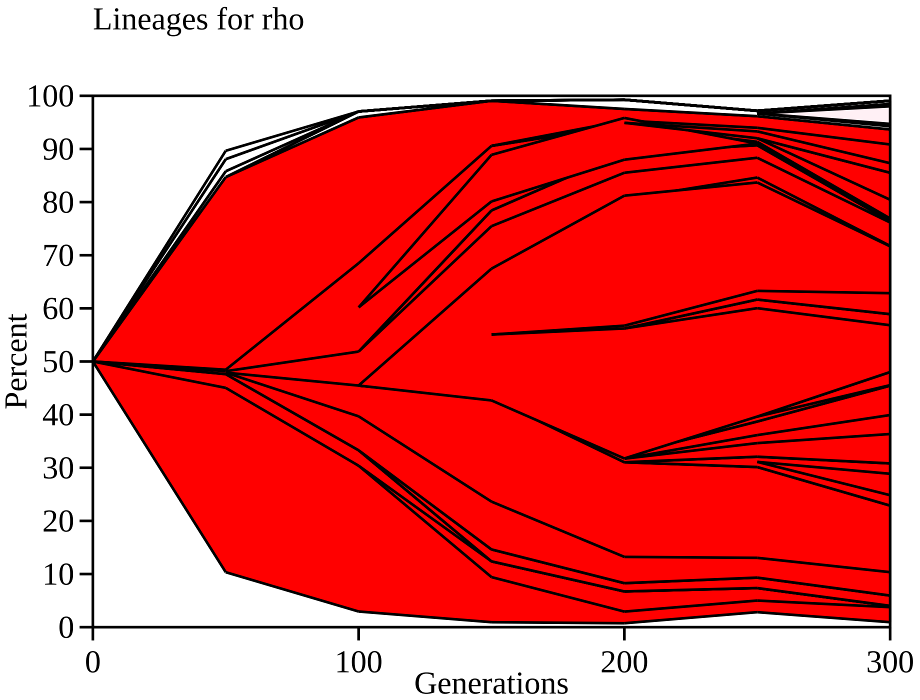

Lineages for rho
Percent
0
10
20
30
40
50
60
70
80
90
100
0
100
200
300
Generations

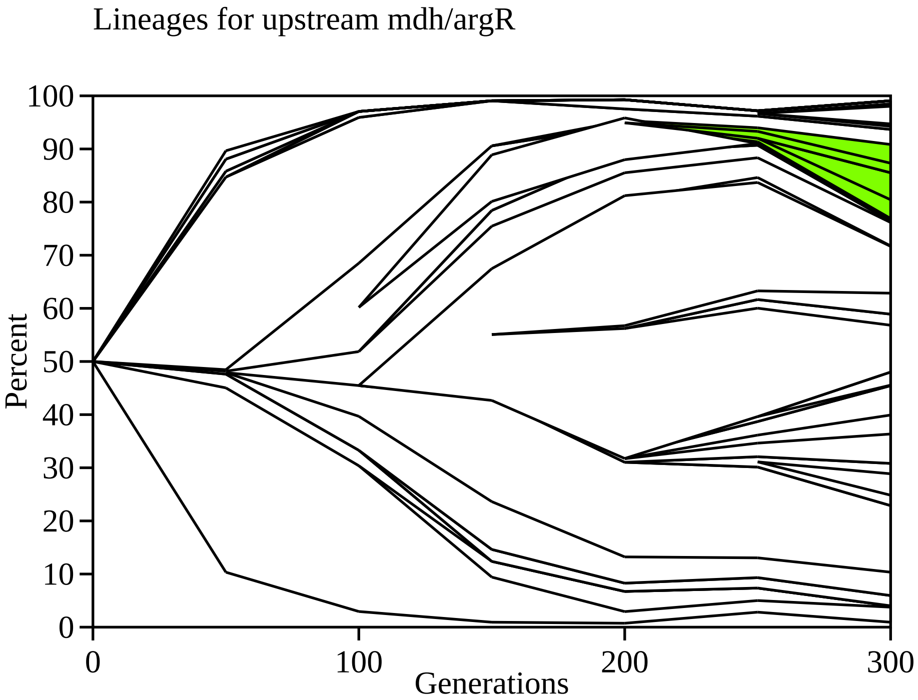
Lineages for upstream mdh/argR
Percent
100
90
80
70
60
50
40
30
20
10
0
0
100
200
300
Generations

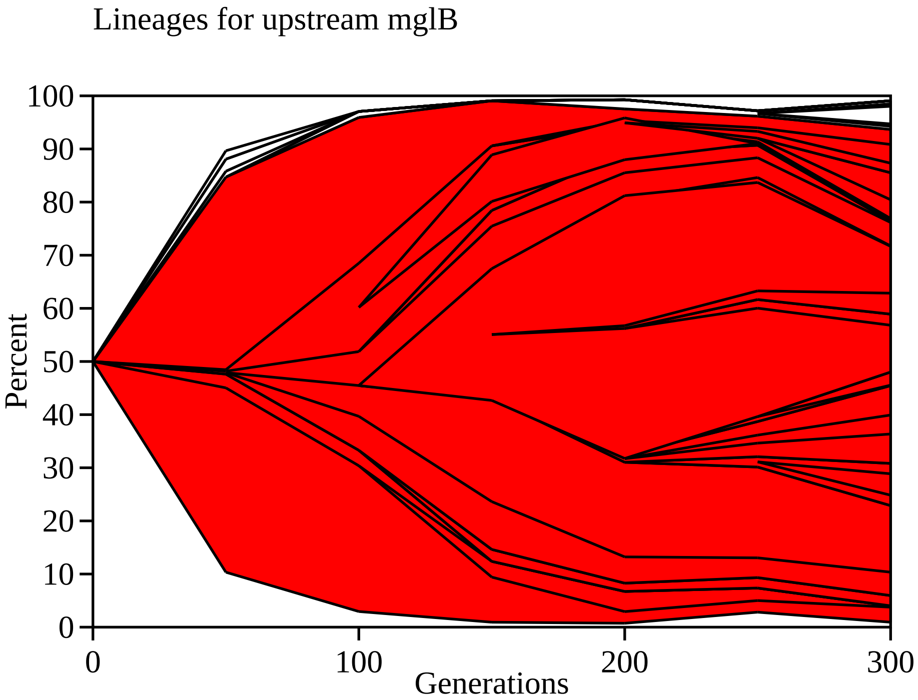
Lineages for upstream mglB
Percent
100
90
80
70
60
50
40
30
20
10
0
0
100
200
300
Generations

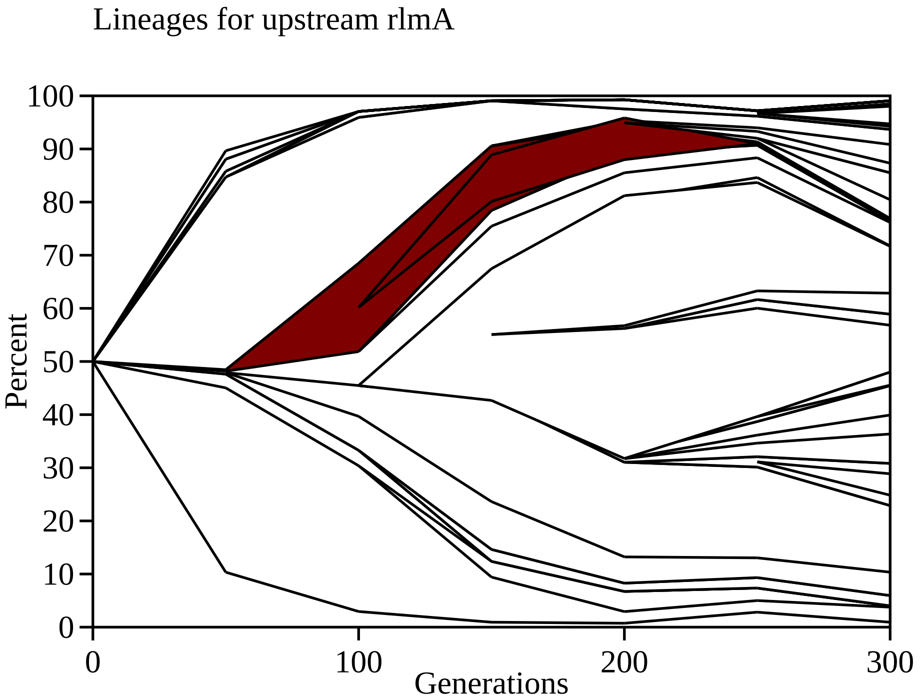
Lineages for upstream rlmA
Percent
100
90
80
70
60
50
40
30
20
10
0
0
100
200
300
Generations

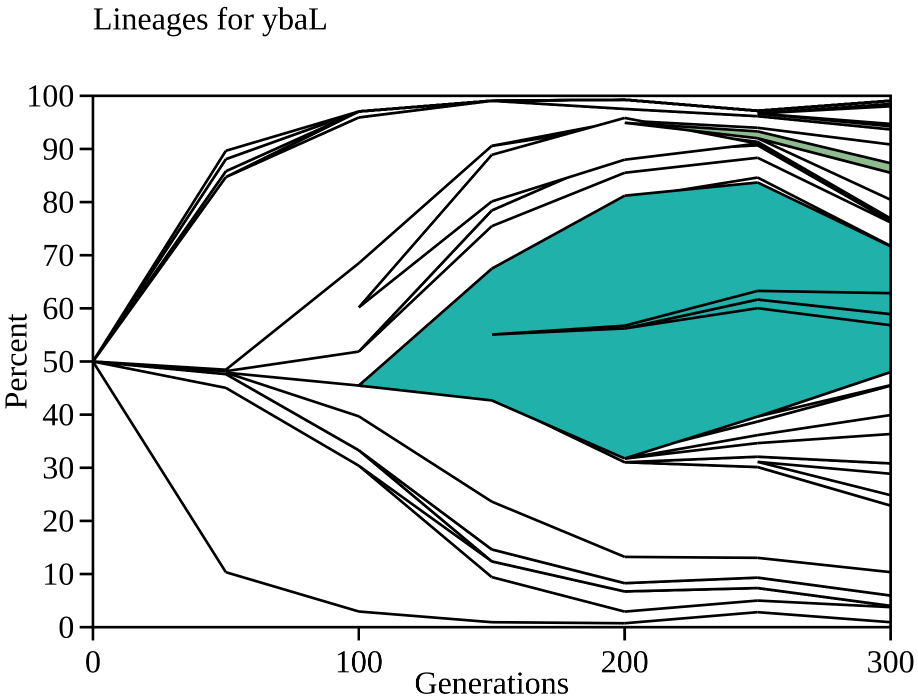

Lineages for ybaL
100
90
80
70
60
50
40
30
20
10
0
Percent
0
100
200
300
Generations

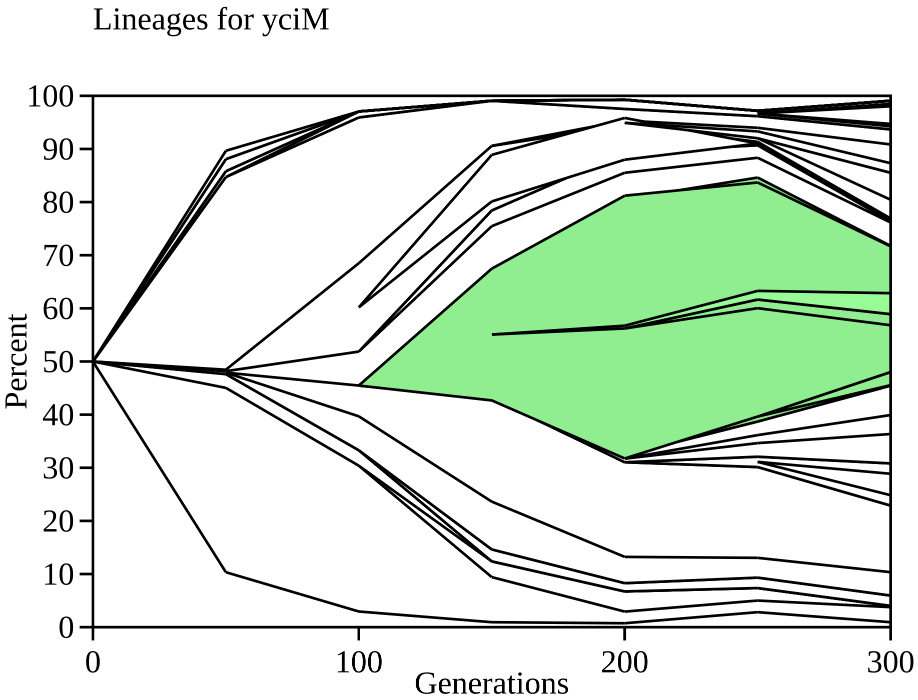

Lineages for yciM
Percent
100
90
80
70
60
50
40
30
20
10
0
0
100
200
300
Generations

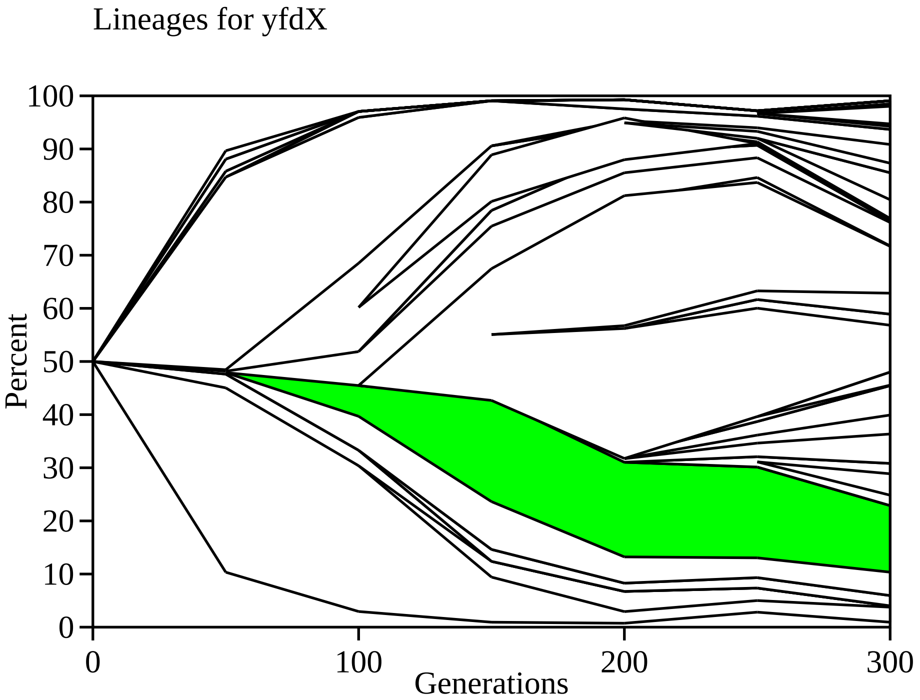

Lineages for yfdX
Percent
100
90
80
70
60
50
40
30
20
10
0
0
100
200
300
Generations

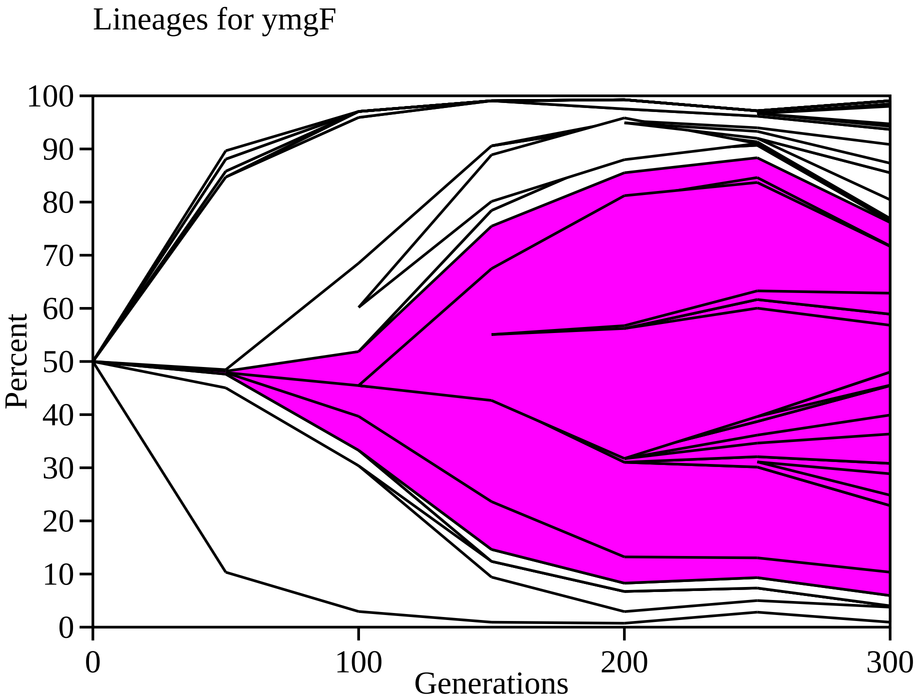
Lineages for ymgF
Percent
100
90
80
70
60
50
40
30
20
10
0
0
100
200
300
Generations

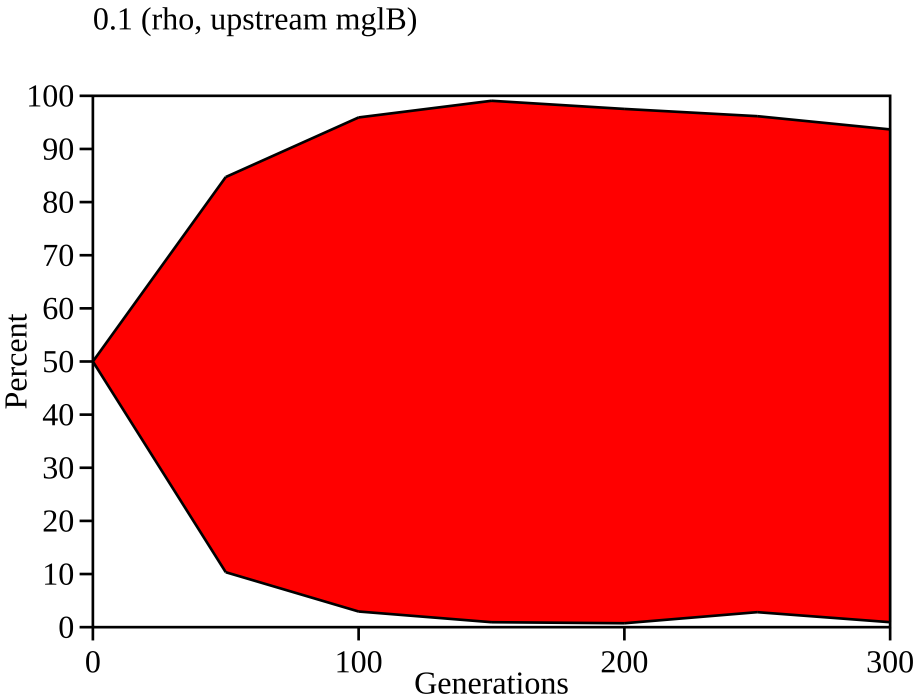

0.1 (rho, upstream mglB)
100
90
80
70
60
50
40
30
20
10
0
Percent
0
100
200
300
Generations

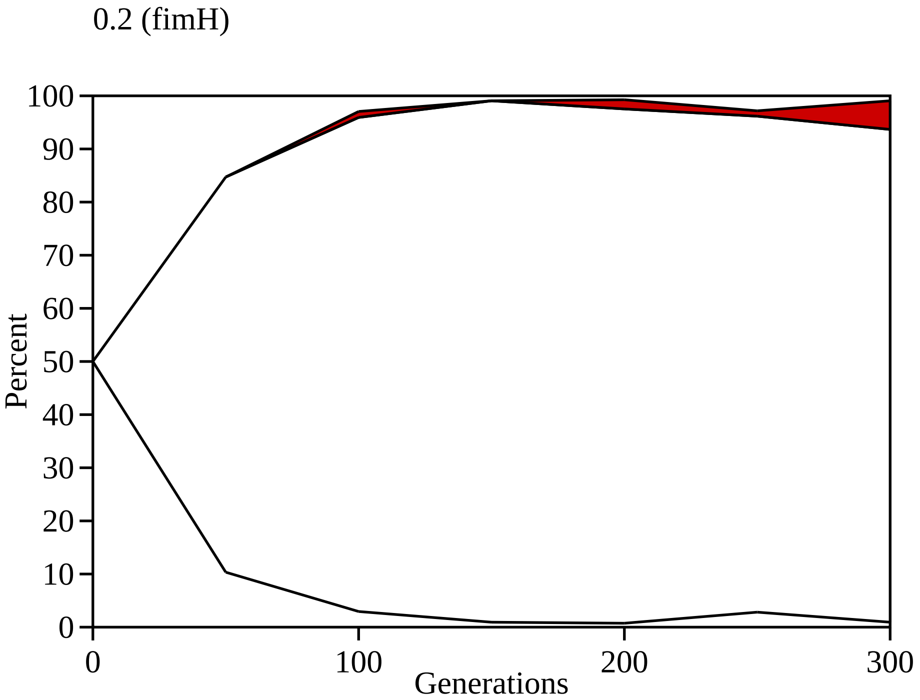
0.2 (fimH)
Percent
Generations
100
90
80
70
60
50
40
30
20
10
0
0
100
200
300

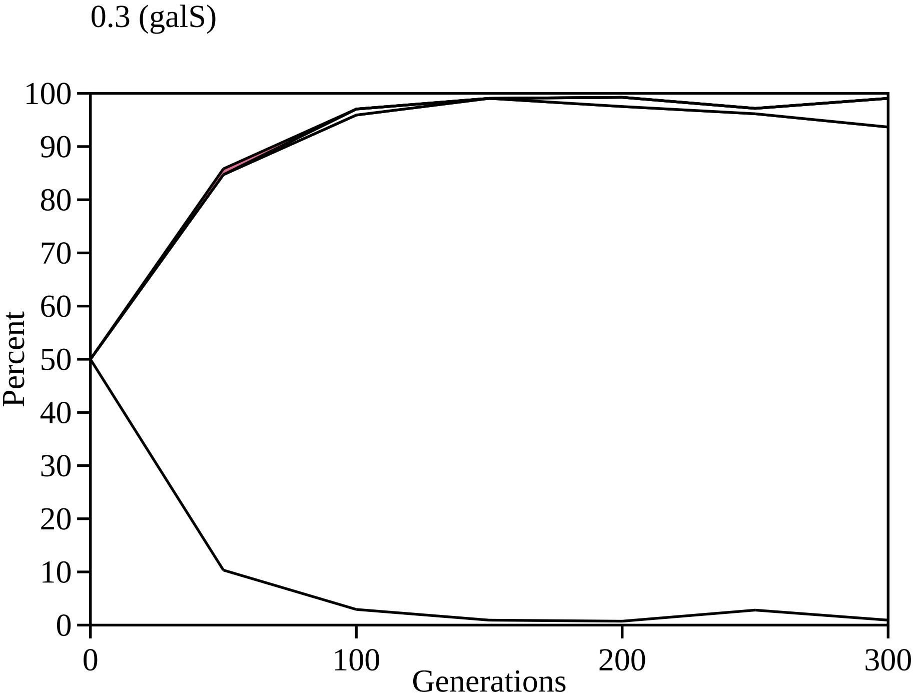
0.3 (galS)
Percent
100
90
80
70
60
50
40
30
20
10
0
0
100
200
300
Generations

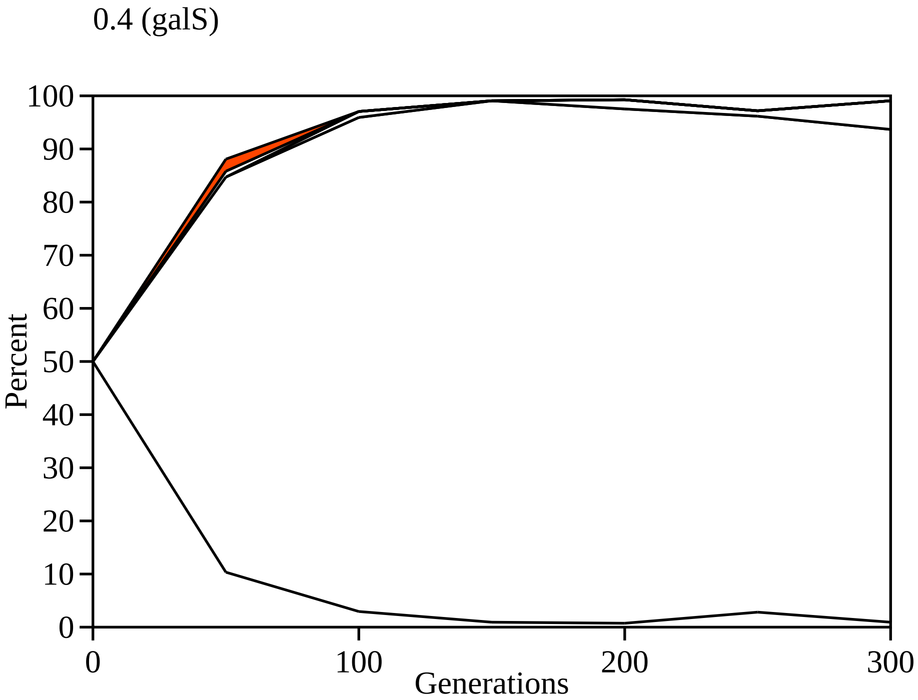
0.4 (galS)
Percent
100
90
80
70
60
50
40
30
20
10
0
0
100
200
300
Generations

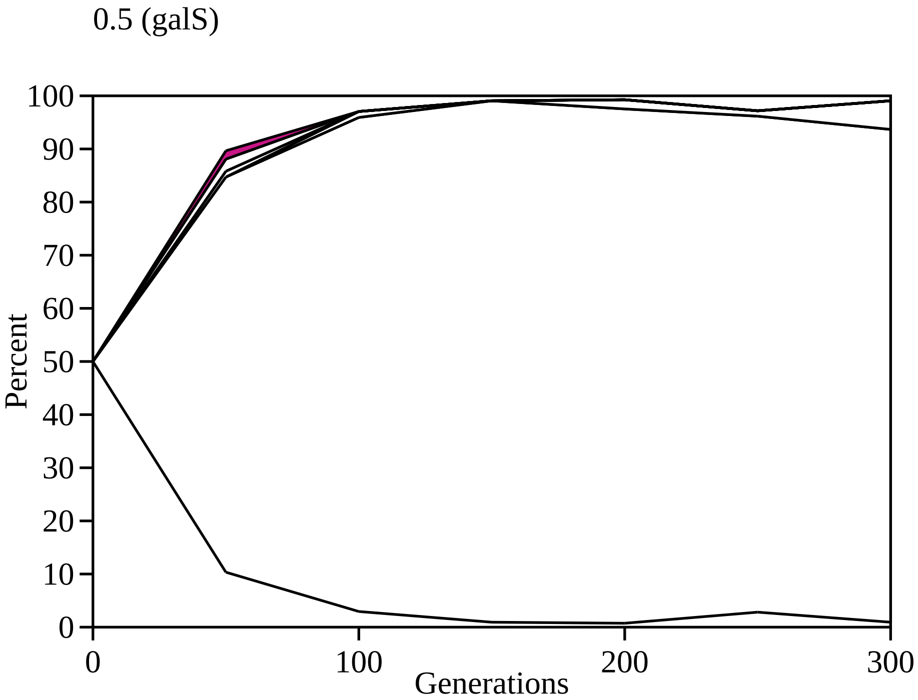

0.5 (galS)
Percent
100
90
80
70
60
50
40
30
20
10
0
0
100
200
300
Generations

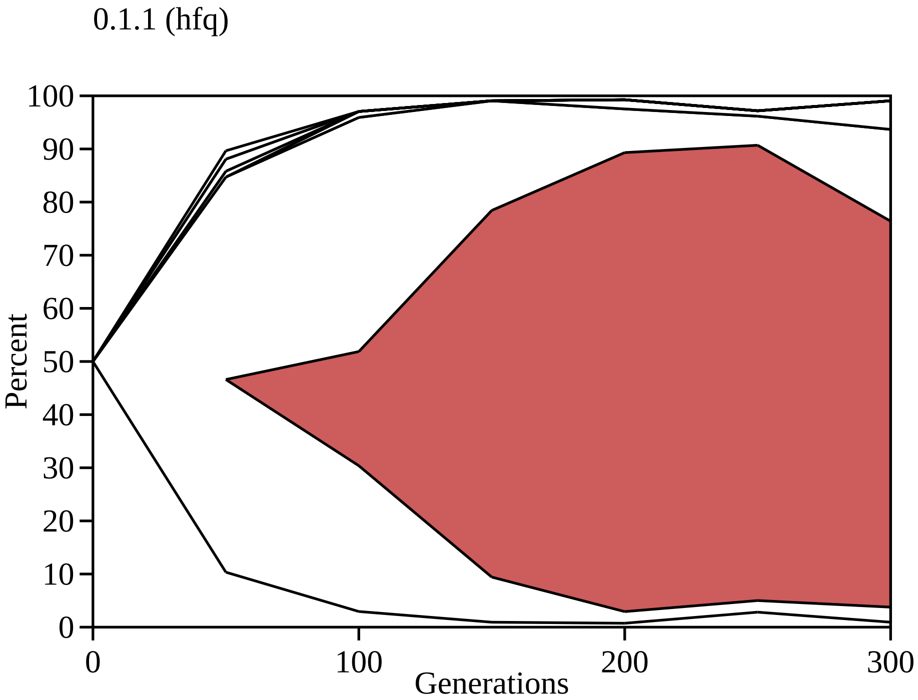

0.1.1 (hfq)
100
90
80
70
60
50
40
30
20
10
0
Percent
0
100
200
300
Generations

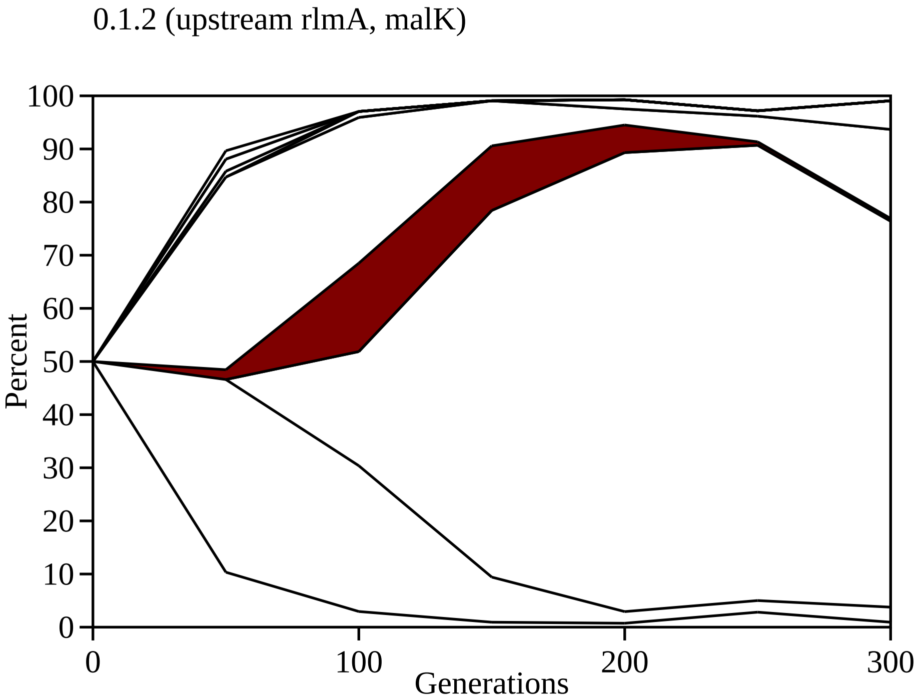

0.1.2 (upstream rlmA, malK)
Percent
Generations
0
10
20
30
40
50
60
70
80
90
100
0
100
200
300

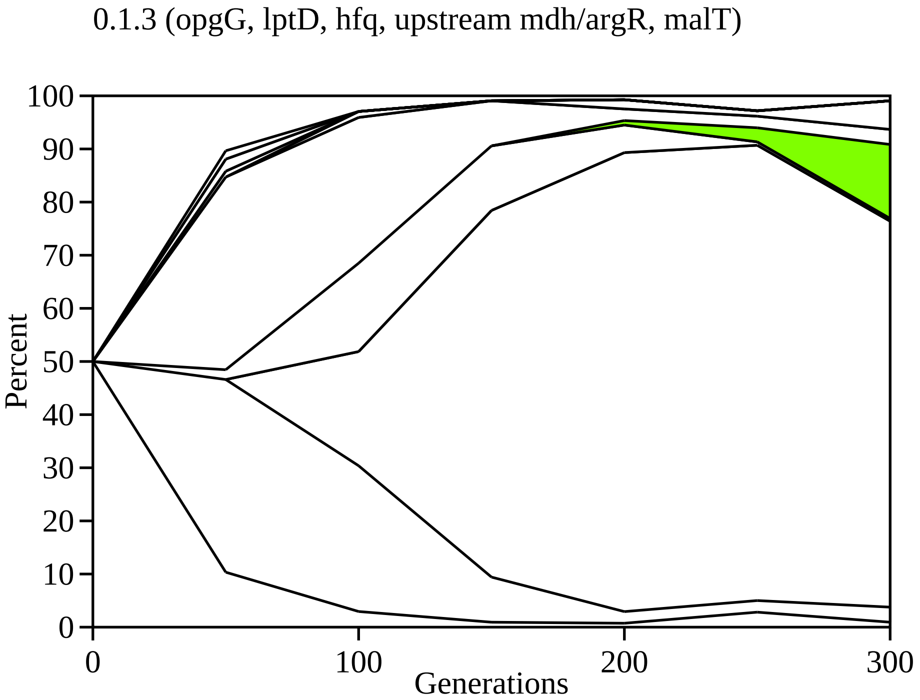
0.1.3 (opgG, lptD, hfq, upstream mdh/argR, malT)
100
90
80
70
60
50
40
30
20
10
0
Percent
0
100
200
300
Generations

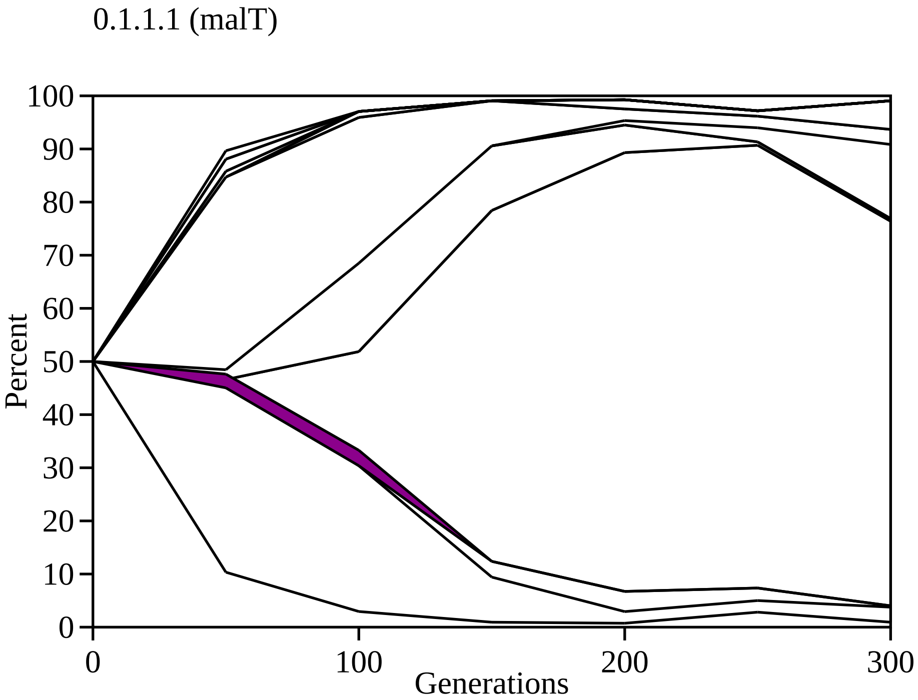
0.1.1.1 (malT)
Percent
100
90
80
70
60
50
40
30
20
10
0
0
100
200
300
Generations

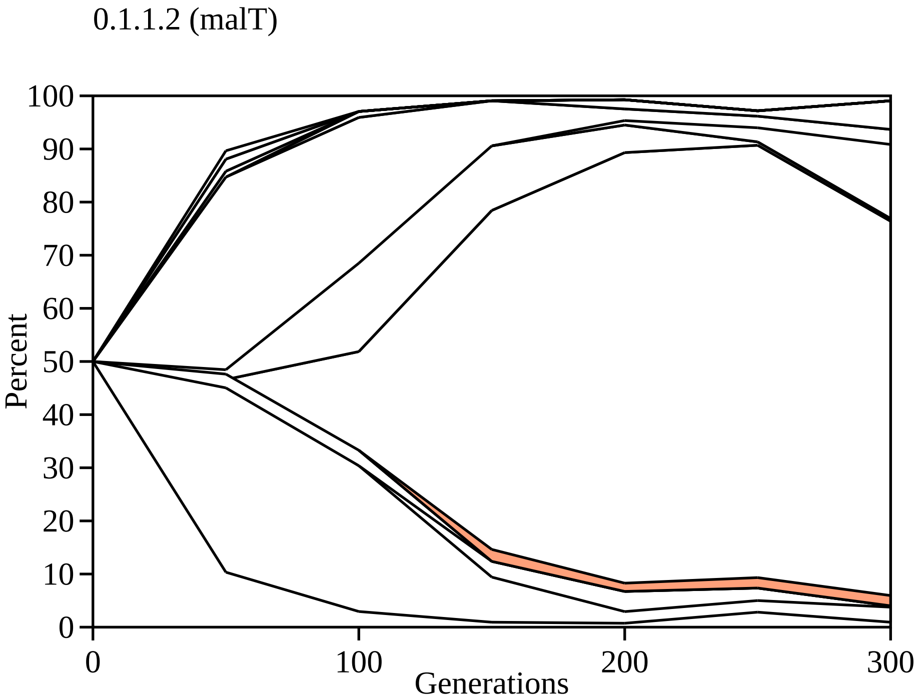

0.1.1.2 (malT)
Percent
100
90
80
70
60
50
40
30
20
10
0
0
100
200
300
Generations

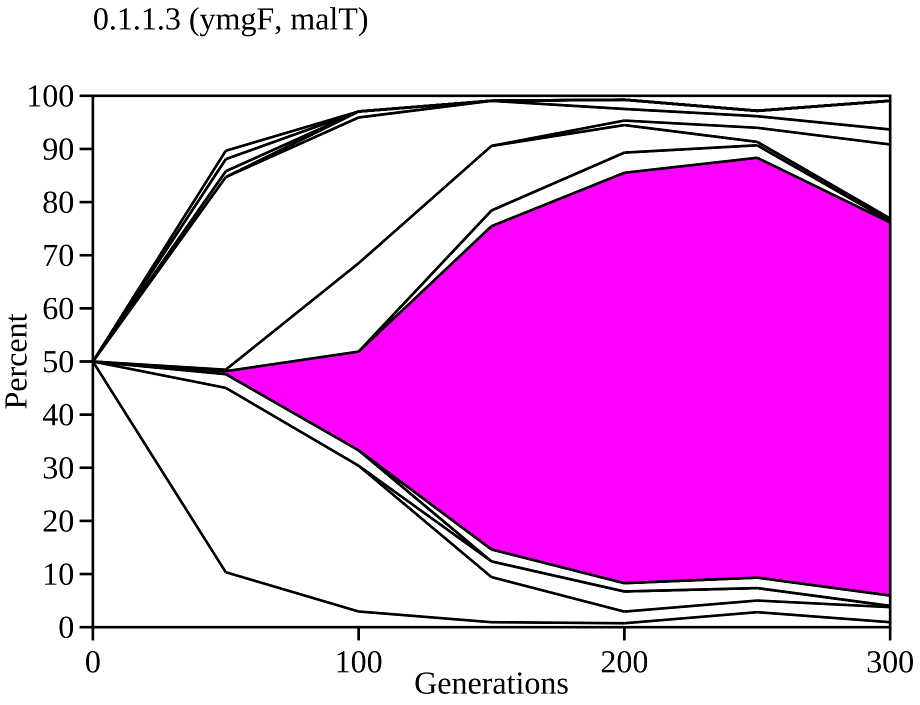

0.1.1.3 (ymgF, malT)
Percent
Generations
0
10
20
30
40
50
60
70
80
90
100
0
100
200
300

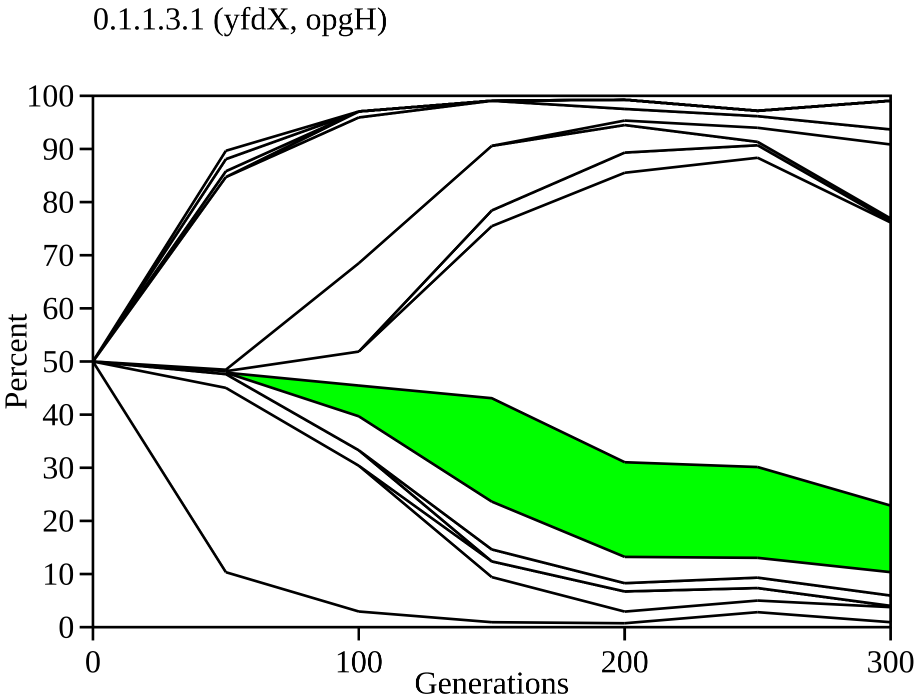
0.1.1.3.1 (yfdX, opgH)
100
90
80
70
60
50
40
30
20
10
0
Percent
0
100
200
300
Generations

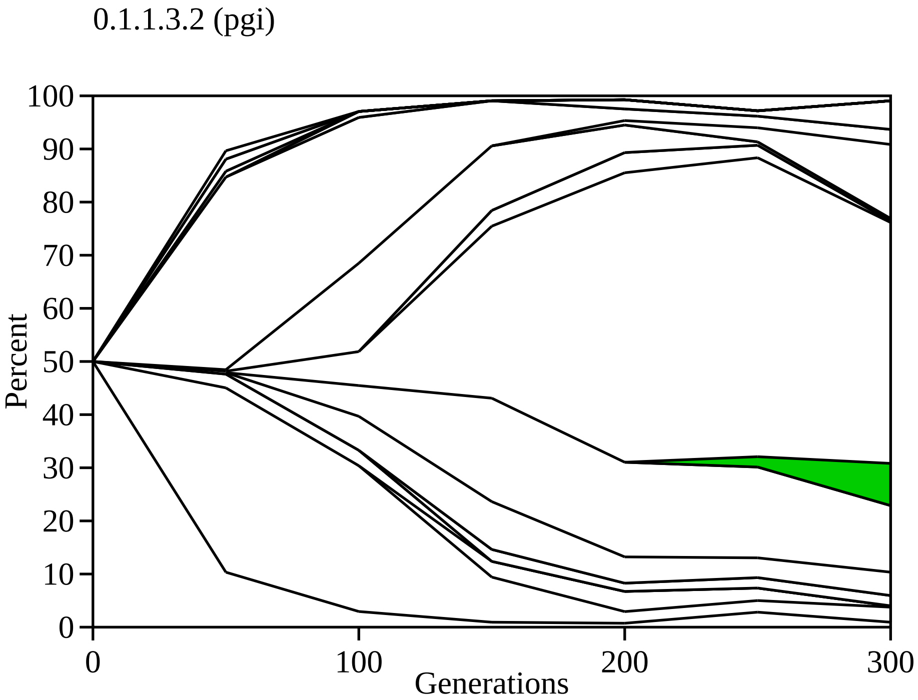
0.1.1.3.2 (pgi)
Percent
0
10
20
30
40
50
60
70
80
90
100
Generations
0
100
200
300

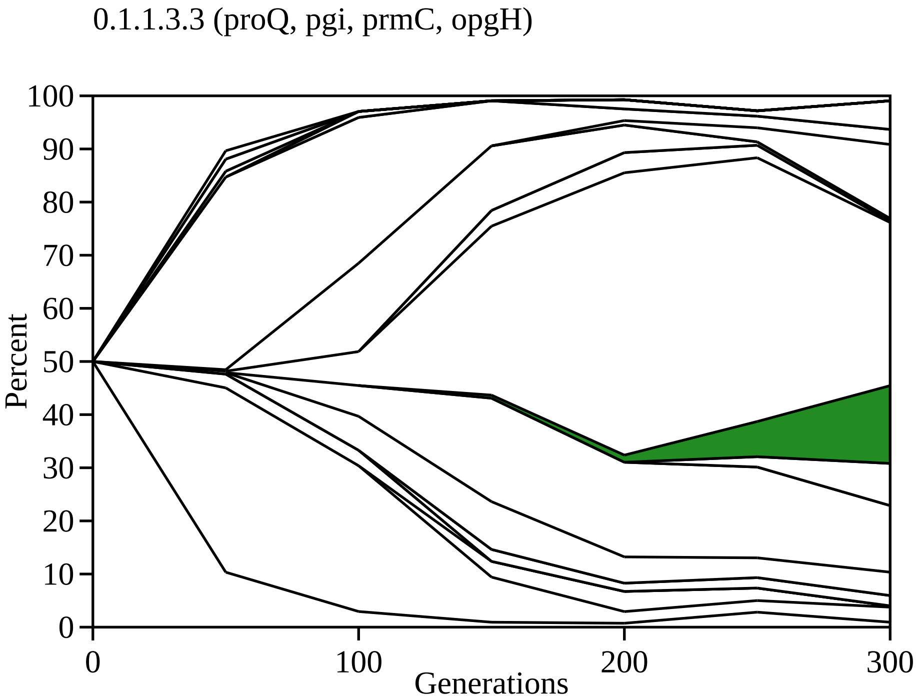

0.1.1.3.3 (proQ, pgi, prmC, opgH)
Percent
100
90
80
70
60
50
40
30
20
10
0
0
100
200
300
Generations

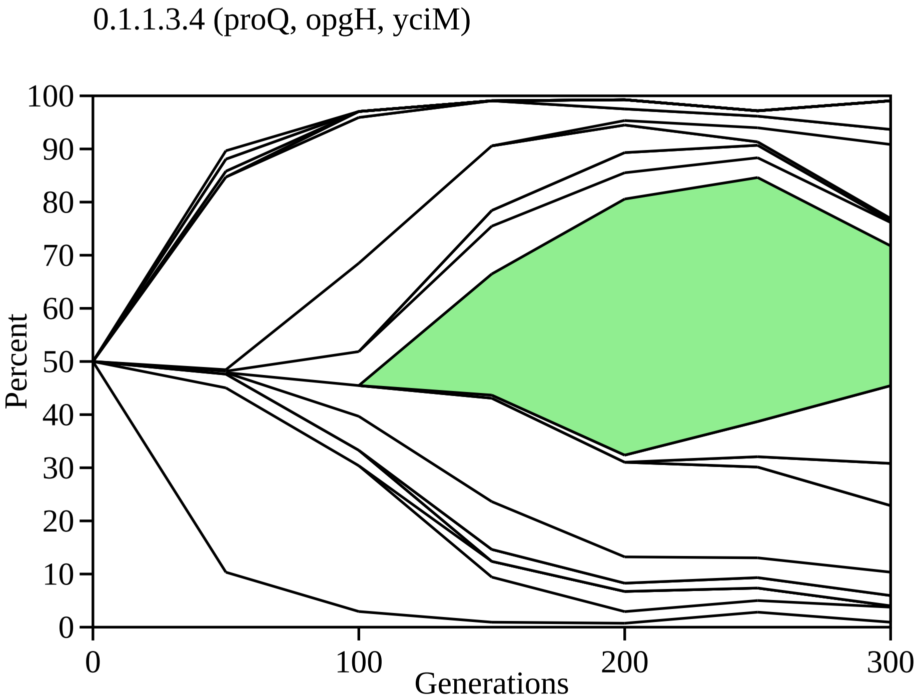
0.1.1.3.4 (proQ, opgH, yciM)
Percent
100
90
80
70
60
50
40
30
20
10
0
0
100
200
300
Generations

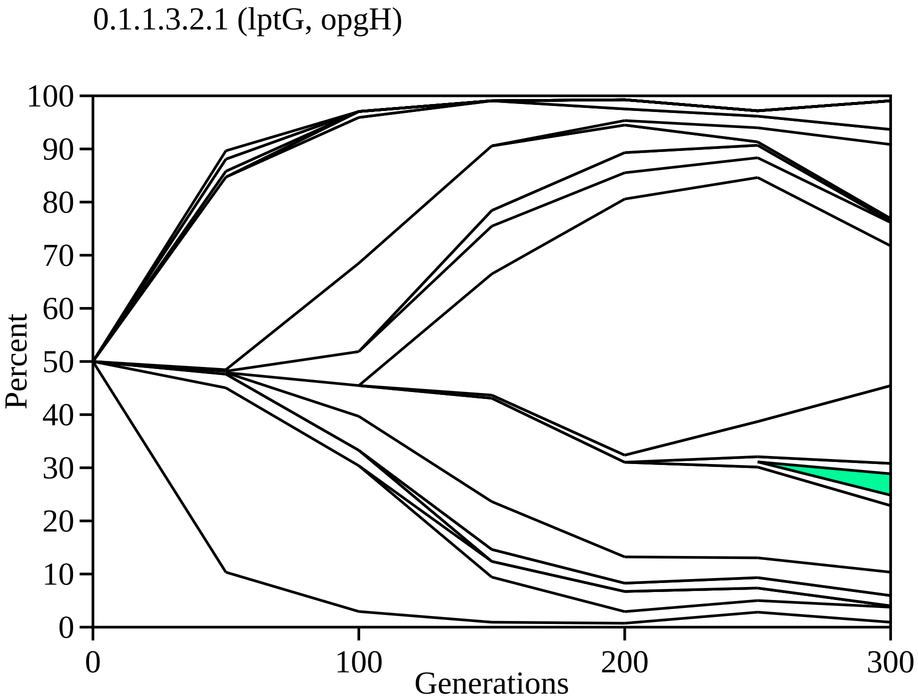
0.1.1.3.2.1 (lptG, opgH)
Percent
100
90
80
70
60
50
40
30
20
10
0
0
100
200
300
Generations

0.1.1.3.3.1 (lpxD)

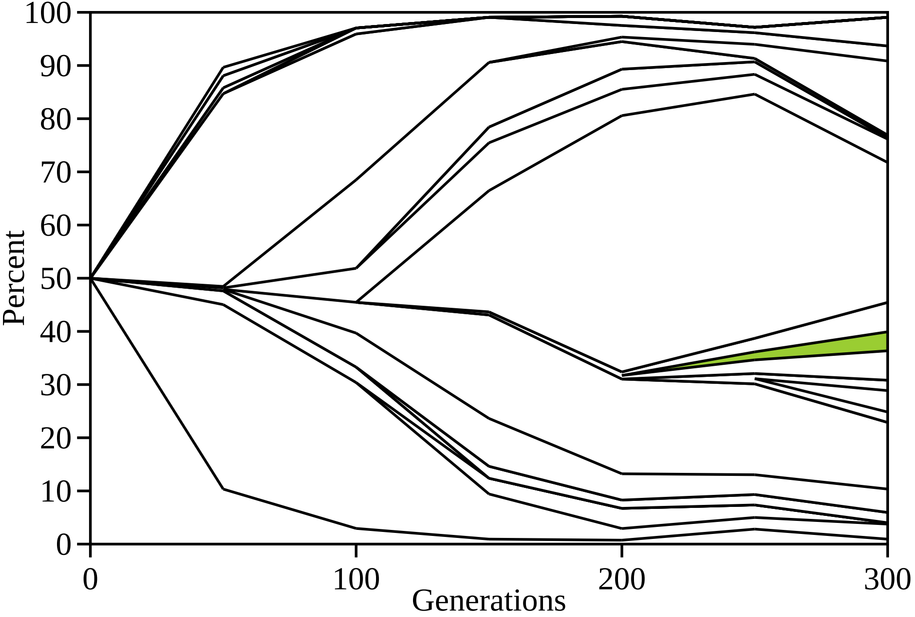
100
90
80
70
60
50
40
30
20
10
0
Percent
0
100
200
300
Generations

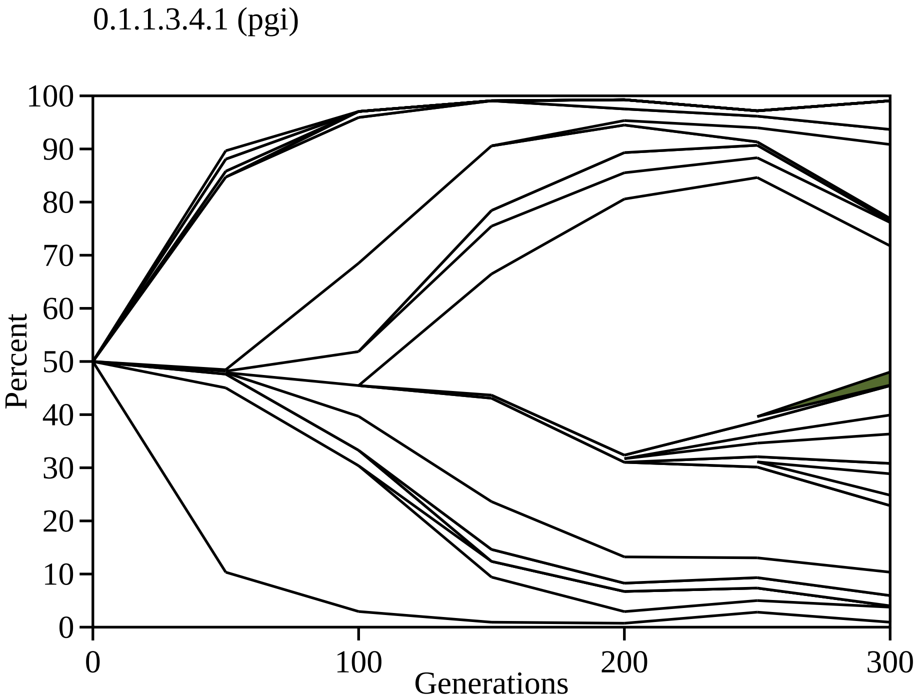
0.1.1.3.4.1 (pgi)
Percent
100
90
80
70
60
50
40
30
20
10
0
0
100
200
300
Generations

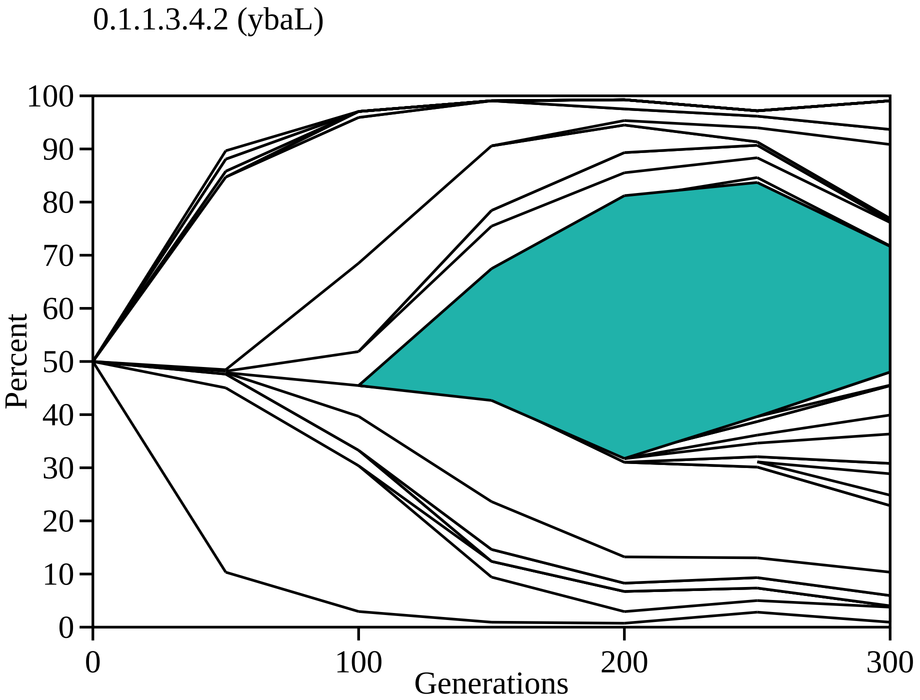
0.1.1.3.4.2 (ybaL)
Percent
100
90
80
70
60
50
40
30
20
10
0
0
100
200
300
Generations

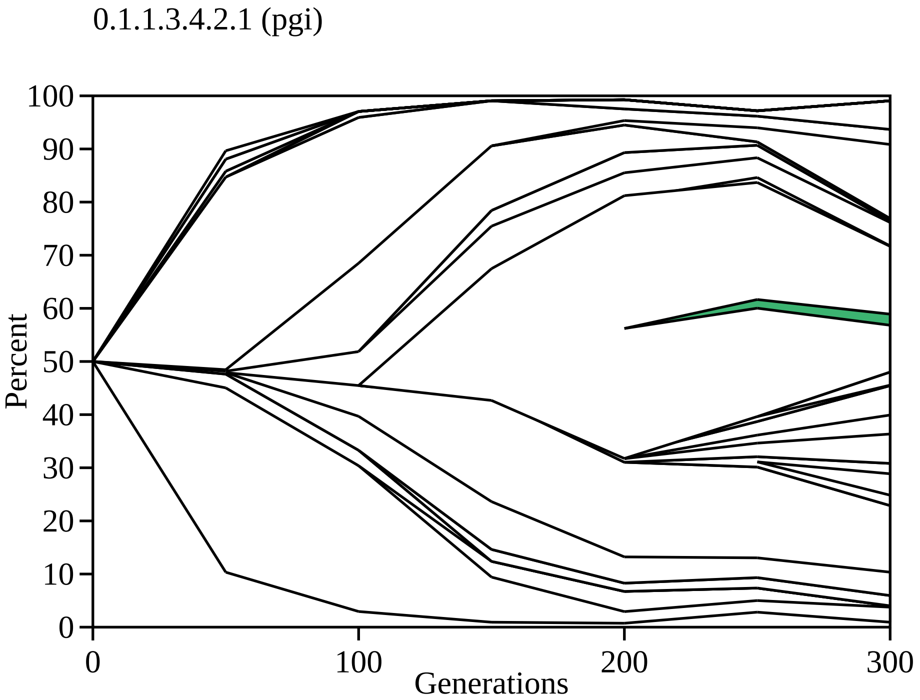

0.1.1.3.4.2.1 (pgi)
Percent
100
90
80
70
60
50
40
30
20
10
0
0
100
200
300
Generations

0.1.1.3.4.2.2 (pgi, yciM)

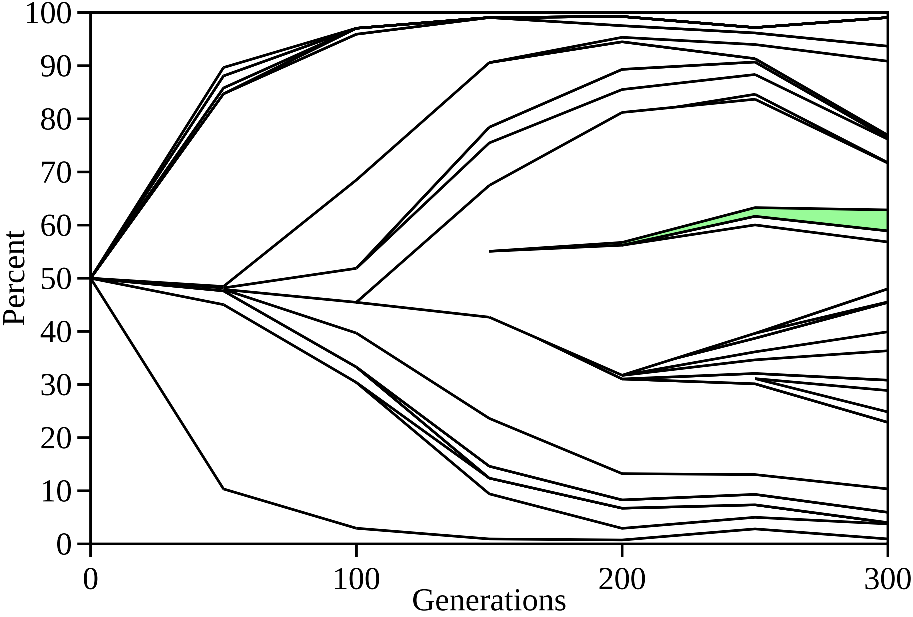

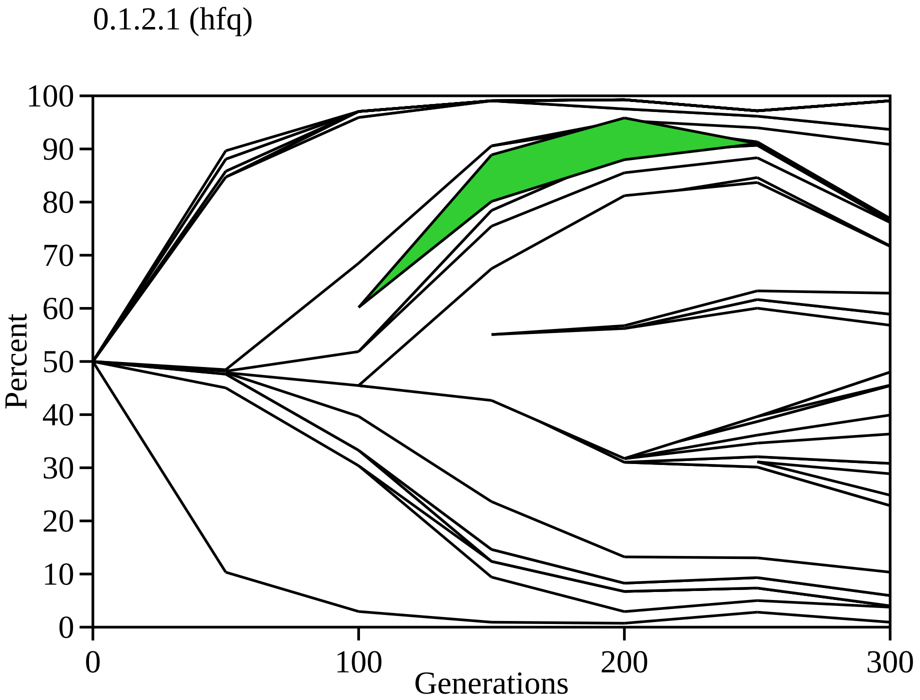

0.1.2.1 (hfq)
Percent
100
90
80
70
60
50
40
30
20
10
0
0
100
200
300
Generations

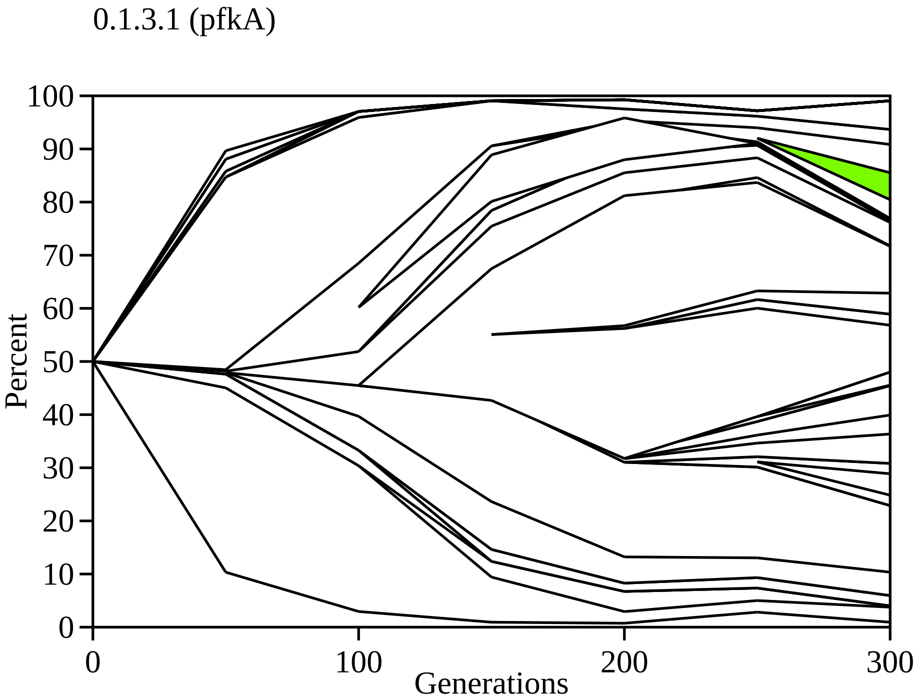
0.1.3.1 (pfkA)
Percent
Generations
100
90
80
70
60
50
40
30
20
10
0
0
100
200
300

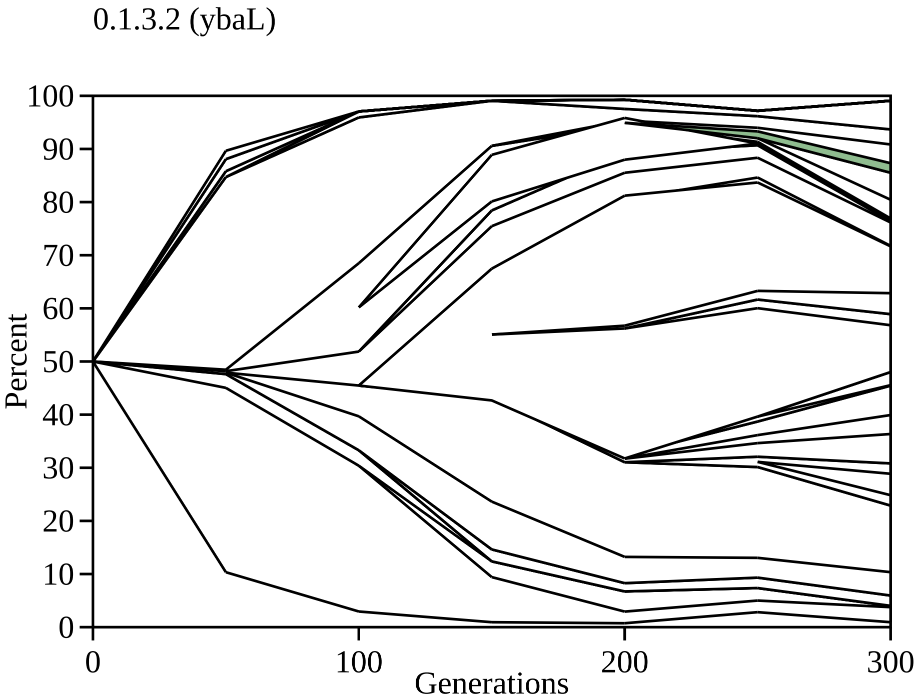
0.1.3.2 (ybaL)
Percent
100
90
80
70
60
50
40
30
20
10
0
0
100
200
300
Generations

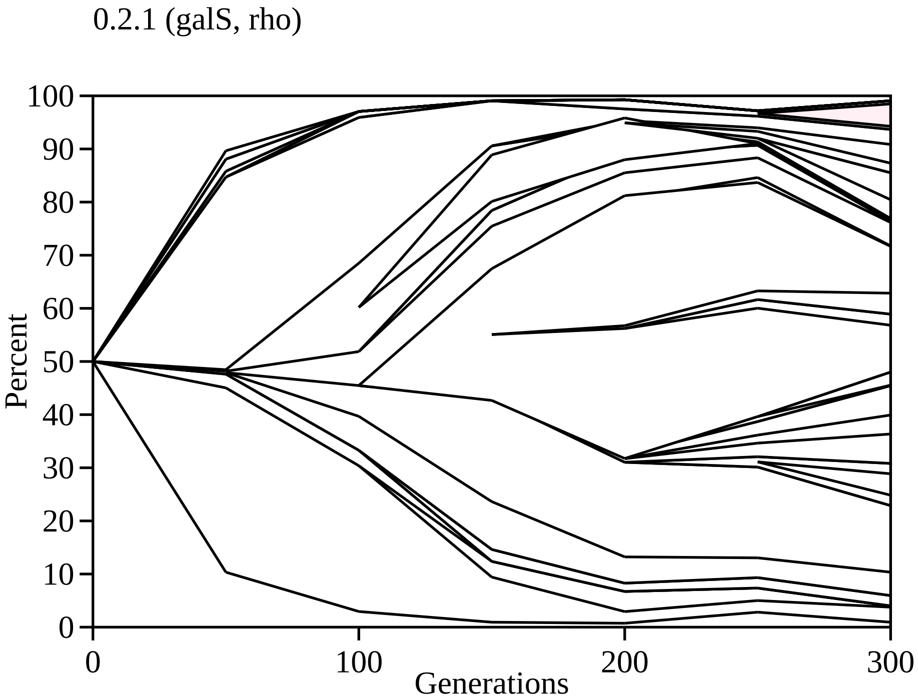
0.2.1 (galS, rho)
Percent
100
90
80
70
60
50
40
30
20
10
0
0
100
200
300
Generations

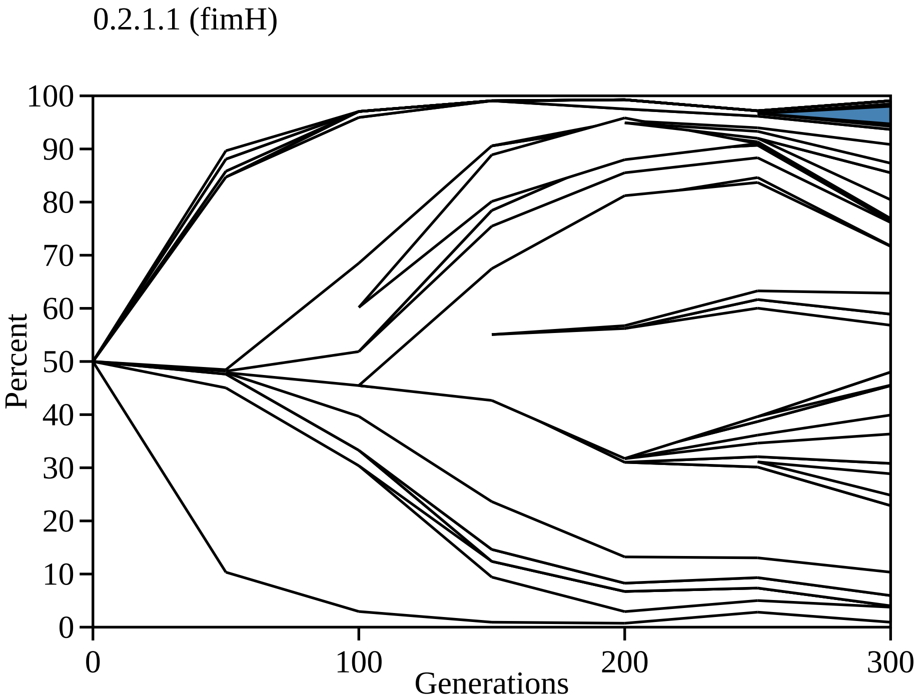
0.2.1.1 (fimH)
100
90
80
70
60
50
40
30
20
10
0
Percent
0
100
200
300
Generations

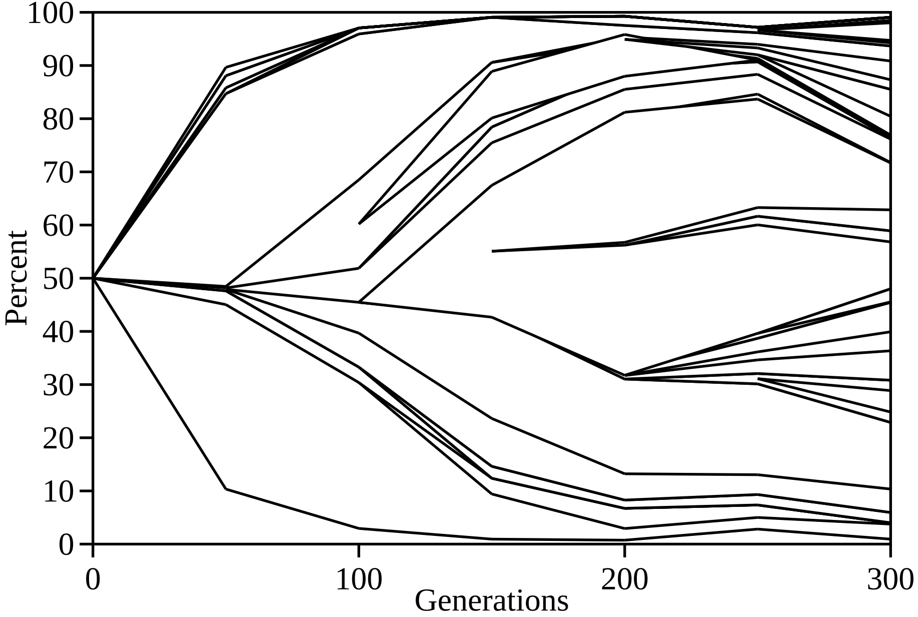

Percent
Generations
100
90
80
70
60
50
40
30
20
10
0
0
100
200
300
